# Functional cooperation, adaptation and synchronization of subcellular processes in cellular neighborhoods as inferred from spatial transcriptomics

**DOI:** 10.64898/2026.11.24.768887

**Authors:** Jens Hansen, Ravi Iyengar

## Abstract

Tissue-level functions emerge from pathway activities distributed across neighboring cells. Yet analyses of spatial transcriptomics mostly focus on cell type composition or signaling through ligand-receptor pairs, leaving spatial relationships between effector processes undescribed. We inferred networks of functionally interacting subcellular processes (SCPs) that are independent of shared gene products and lie beyond sibling relationships defined in parent-child hierarchies. Using those networks for the analysis of published spatial transcriptomics datasets from human kidney biopsies, we identified functionally related SCP activities between a tissue spot and its neighbors. Identified pairs of adjacent SCPs indicate functional cooperation, e.g. basement membrane assembly and cell-matrix adhesion, functional adaption, e.g. complement synthesis and inhibition, and functional synchronization, e.g. folate cycle and cobalamin metabolism or pre-mRNA splicing and nuclear mRNA export. Two independent null models, shuffling of SCP labels within the network and shuffling of spots on the tissue slice, distinguish pairs that are functionally non-random and pairs that are additionally enriched within the immediate neighborhood, separating local from tissue-wide coordination. Our results suggest that tissue physiology reflects not only cell type composition but also functional coordination between neighboring regions.

## Introduction

Cell biological functions arise from the interaction of multiple pathways, each representing a set of interacting molecular components. Pathways form functional networks that converge into physiological or pathophysiological responses. Within single cells networks of interacting pathways are well described and enable whole-cell functions and cell physiology. Whole-tissue functions and physiology are likely generated by interacting cells within spatial neighborhoods. Multiple studies have characterized spatially colocalizing cells and communication networks that orchestrate their activities, while, to our knowledge, cross-cellular interactions between effector pathways have not been systematically analyzed yet.

Understanding whole-cell functions and examining potential trans-cellular pathway interactions depends on knowledge of the topology of functionally interacting pathways and the resultant functional networks. To enable the development of algorithms for the analysis of functional pathway interactions, we have generated, optimized and extended the Molecular Biology of the Cell Ontology (MBCO),^1^ a strictly cell biological ontology that organizes subcellular processes (SCPs) following the structure of cell biology and biochemistry textbooks.^2–4^ MBCO SCPs are organized across four levels of increasing granularity, with minimized functional redundancies within the same level. The annotated hierarchy that connects parent and child SCPs of adjacent levels is enriched by inferred networks that connect functionally interacting pathways within either level 2 or level 3. These weighted interactions that are predicted from text mining and statistical analyses span all areas of cell biology and exceed those predicted from shared gene annotations. SCPs are connected, if genes and metabolites annotated to one SCP are found in the abstracts that were mined for the other SCP. Consequently, identified functional relationships within inferred MBCO networks include SCPs whose gene products physically interact, that process each other’s substrates and products, represent sequential steps within a biological program, regulate the other SCP’s gene expression, or generate co-factors for the other SCP’s enzymes. Inferred SCP networks can be investigated using our desktop and command line tool MBC PathNet.^5^

Generated comprehensive networks connecting SCPs across all areas of cell biology allowed us to systematically identify functionally related SCPs from genes expressed in adjacent tissue spots, using published spatial transcriptomic experiments.^6^ Our approach addresses a gap in current spatial analysis methods. Ligand-receptor inference tools identify communication events between neighboring spots,^7–9^ but do not investigate functional relationships between adjacent effector pathways. Niche and domain detection methods group spots by cellular composition or by overall expression similarity, so they describe where tissue regions differ without specifying which cellular functions are shared or not between neighboring regions. Machine learning approaches that predict a center spot’s features from its surroundings identify statistically significant associations between features ^10,11^ but do not reveal how functional relationships between local and tissue-wide neighboring features together enable tissue physiology.

Our algorithm identified SCP interactions between neighboring spots that extend beyond pairs describing intercellular interactions and instead connect SCPs whose relationships are traditionally considered intracellular. The latter include SCP pairs involved in mitochondrial structural dynamics and energy generation, mRNA maturation as well as endogenous antigen processing and presentation. Since our algorithm prioritizes mapping of identical over functionally related SCP pairs, identified pairs exclude cases that would simply reflect both SCPs being co-expressed in neighboring spots. Adjacent SCPs are mapped according to their enrichment scores, and each predicted SCP pair is assigned a magnitude that reflects potential relevance. Our results are derived from two independent null models, i.e. shuffling of SCP labels within the network and of spot positions on the slice. Both allow us to identify relationships that are functionally non-random and to further separate them by local versus tissue-wide coordination, a distinction that, to our knowledge, other approaches do not make.

Building on the combination of prior-knowledge-based functional SCP networks and graph theoretical algorithms, our analysis provides a deeper understanding of tissue organization in terms of adjacency of cell types and subtypes and their interacting pathways that give rise to emergent tissue level functions.

## Results

### Enhancement and optimization of MBCO SCPs and SCP networks

Optimizing and enhancing MBCO, we increased the number of SCPs, genes and gene SCP annotations from 743, 5357 and 18488 to 1710, 7695 and 31518, respectively (Table 1). New SCPs were added to increase the coverage of cell biological functions, existing SCPs were redesigned to emphasize the functional consequences of gene expression changes, split to cover different functional aspects that can be active independently from each other and combined to merge functions that are normally linked to each other. We also populated level 3 SCPs with 1515 metabolites or metabolite groups. Though the number of metabolites is not yet sufficient for enrichment analysis of metabolite lists by themselves, they enable better predictions of networks of functionally interacting SCPs (see below).

Table 1.

| **Level** | **# SCPs in v1** | **# SCPs in v2** | **# genes in v1** | **# genes in v2** | **# SCP gene annotations in v1** | **# SCP gene annotations in v2** | **# new & renamed SCPs** | **# removed & renamed SCPs** |
|---|---|---|---|---|---|---|---|---|
| **1** | 29 | 29 | 5357 | 7695 | 6302 | 9349 | 2 | 2 |
| **2** | 125 | 136 | 5322 | 7539 | 6658 | 9748 | 51 | 40 |
| **3** | 483 | 768 | 4177 | 6498 | 4897 | 8122 | 511 | 226 |
| **4** | 106 | 777 | 597 | 3336 | 631 | 4299 | 718 | 47 |
| **Total** | 743 | 1710 | 5357 | 7695 | 18488 | 31518 | 1282 | 315 |

Our desktop and command line application MBC PathNet can be used to explore parent-child hierarchy and functional SCP networks as well as for enrichment analysis of custom datasets.^5^

As an example of the capability of this enhanced MBCO, we show all populated offsprings of the level 2 SCP ‘Enzymatic extracellular matrix degeneration and processing’ (Figure 1A). We replaced SCPs that grouped enzymes based on their biochemical classification by new SCPs that describe the degradation target, shifting the focus from enzymatic mechanisms to the functional consequences of SCP activities.

**Figure 1.**
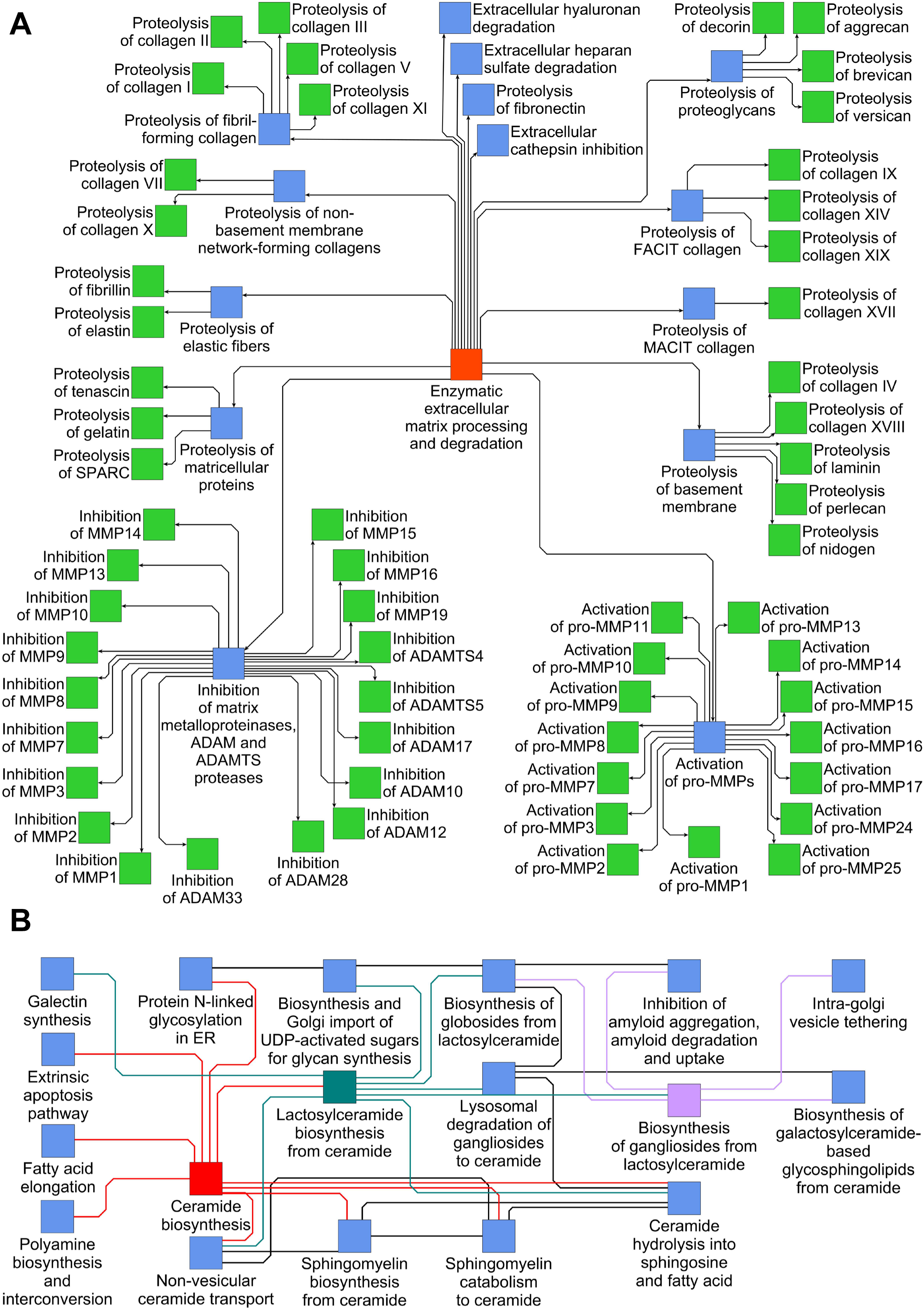
Hierarchal and inferred SCP interactions. **(A)** All populated SCPs that are children of the level 2 SCP ‘Enzymatic extracellular matrix processing and degradation’ have been redesigned. Instead of grouping enzymes with the same biochemical mechanism into the same SCPs, we now group enzymes with the same substrates into the same SCPs, increasing the focus on the functional consequences of expression changes. **(B)** Among the new SCPs are multiple mitochondrial SCPs that are connected within our inferred network of functional relationships. The network shows all direct neighbor SCPs (light blue) of the selected SCPs (red, dark cyan, purple), based on the top 25% of the inferred SCP interactions.

Matching the new structure of the ECM degradation branch, the other SCPs involved in ECM homeostasis were organized accordingly (Suppl. Figure 1A). Other examples of newly added and redesigned level 3 SCPs are SCPs involved in redox homeostasis (Suppl. Figure 1B), organelle organization, e.g. mitochondrial structural dynamics and repair as well as plasma membrane organization (Suppl. Fig. 1C), intracellular degradation, e.g. selective autophagy (Suppl. Figure 1D), amino acid metabolism, e.g. polyamine metabolism (Suppl. Figure 1E), lipoprotein metabolism (Suppl. Fig. 1F), protein folding and post-translational protein modification (Suppl. Figure 1G), intracellular cascades shared by multiple signaling pathways (Suppl. Figure 1H), vesicle traffic (Suppl. Figure 1I), actin (Supl. Fig. 1J) and microtubule dynamics (Suppl. Fig. 1K), interphase centrosome dynamics (Suppl. Fig. 1L), cellular protrusion and cilium dynamics (Suppl. Fig. 1M), electric transmission (Suppl. Fig. 1N), transmembrane ion movement (Suppl. Fig. 1O), cell cycle (Suppl. Figure 1P), DNA replication, recombination and repair (Suppl. Figure 1Q), cholesterol metabolism (Suppl. Figure 1R), sphingolipid metabolism (Suppl. Figure 1S) and signaling pathways involved in cellular proliferation and differentiation (Suppl. Fig. 1T).

Functional SCP interactions were independently predicted based on genes or metabolites and then averaged across both entity classes. Resultant networks contained many interactions between SCPs that are well known to functionally interact with each other, e.g. SCPs involved in sphingolipid metabolism and their interaction partners (Fig. 1B). Other examples connect SCPs involved in mitochondrial energy generation (Suppl. Fig. 2A), reactive oxygen species generation and detoxification (Suppl. Fig. 2B), mitochondrial dynamics and lipid droplet biology (Suppl. Fig. 2C), the mevalonate and cholesterol synthesis pathway (Suppl. Fig. 2D), protein lipidation pathways (Suppl. Fig. 2E), polyamine, folate and methionine metabolism (Suppl. Fig. 2F), eicosanoid metabolism (Suppl. Fig. 2G), heme and hemoglobin metabolism (Suppl. Fig. 2H), lysosomal degradation pathways (Suppl. Fig. 2I), autophagy pathways (Suppl. Fig. 2J), G-protein signaling pathways (Suppl. Fig. 2K), steroid hormone homeostasis (Suppl. Fig. 2L), signaling pathways regulating metabolism (Suppl. Fig. 2M), immune response signaling (Suppl. Fig. 2N), microtubule nucleation pathways (Suppl. Fig. 2O), cellular actin structures (Suppl. Fig. 2P), cellular contraction and force transmission (Suppl. Fig. 2Q), cellular adhesion (Suppl. Fig. 2R) as well as mRNA processing and nuclear export (Suppl. Fig. 2S). These findings indicate our ability to capture molecular networks for higher level biochemical (e.g., energy metabolism), cell biological (e.g., autophagy) and physiological (e.g. contractility) functions.

### SCP networks document functional integrity for multiple cell types from single nucleus transcriptomic data sets

To test the relevance of inferred SCP networks for integrated whole cell level functions, we analyzed if SCPs expressed in a cell type of interest are functionally more related to each other than expected by chance. The underlying assumption of this approach is that SCPs expressed in the same cell type should be functionally interacting with each other to orchestrate a cell-type-selective whole-cell-function. Marker genes of liver,^12^ cardiac,^13^ and kidney ^14^ cell types were subjected to pathway enrichment analysis using Fisher’s exact test and level 3 SCPs (Suppl. Table 1A, B, C, respectively). For each cell type, we mapped the predicted SCPs onto the inferred functional interaction networks, followed by calculation of all shortest paths connecting any two SCPs (Suppl. Fig. 3A). The associated network distances were averaged after weighting them by the –log_10_ p-values of the connected SCPs (Suppl. Table 2A-C). To test significance, we shuffled the labels of SCPs predicted for at least one cell type of the same study within the SCP networks. Very significant, significant and marginally significant empirical p-values were obtained for four, two and zero of seven liver cell types (Fig. 2), four, three and two of 12 heart cell types (Suppl. Fig. 3B) and eleven, two and one of 21 kidney cell types (Suppl. Fig. 3C). Among the few cell types without significance were endothelial cells and the descending and ascending limb epithelial cells of the kidney, whose marker genes describe comparatively few and functionally narrow processes. In summary, this analysis documents functional relevance of inferred SCP networks, since they indicate functional integrity for most analyzed cell types from multiple organ systems.

**Figure 2.**
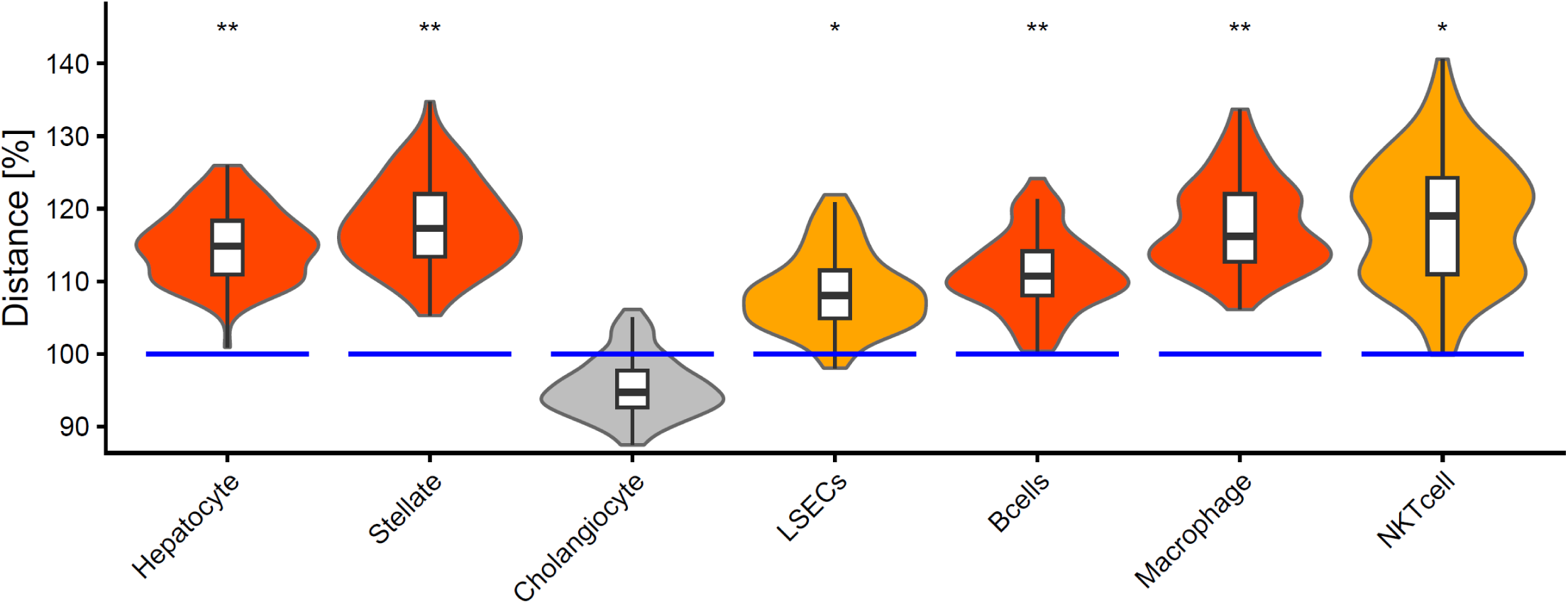
Functional integrity of SCPs expressed in the same cell type. Marker genes of liver cell types were subjected to enrichment analysis using MBCO level 3 SCPs and Fisher’s exact test. For each cell type we calculated the pairwise distances between all significant SCPs within the inferred network of functional relationships. All distances were averaged after weighting them by –log_10_ p-values of the connected SCPs and defined as 100% (blue lines). Labels of SCPs predicted in at least one cell type were shuffled 100 times in the SCP network, followed by recalculation of the weighted average distance, normalizing the results towards the real average distances (violins) and calculation of empirical p-values. red, orange, gray: (*), *, **: 0.1 ≥ p > 0.05, 0.05 ≥ p > 0.01, p ≤ 0.01.

### Functionally related SCPs expressed in adjacent tissue spots suggest functional cooperation, synchronization and adaptation

In an earlier analysis of SCPs expressed in subtypes of the major kidney cell types, we predicted extracellular matrix and collagen synthesis activities for cortical myofibroblasts, while marker genes of failed-repair proximal tubule epithelial cells were enriched for genes involved in ECM and collagen processing.^15^ Since both cell subtypes show at least partial spatial colocalization, we hypothesized that cortical myofibroblasts and failed-repair proximal tubule cells cooperate during scar formation. Using our prior expertise to the analysis of spatial transcriptomics datasets, here we systematically investigated whether neighboring tissue spots exhibit functional cooperation, synchronization, or adaptation of their activities (Fig. 3A). This led us to ask whether correlated expression of two SCPs across a tissue section reflects a functional relationship between those SCPs. We investigated whether SCPs that are coordinately upregulated in adjacent spots are identical, functionally related, or functionally independent, using spatial transcriptomic data from human reference and diseased kidney biopsies.^6^ We defined a neighborhood by the center spot and all peripheral spots that lie within 220 μm (two spots in each direction). Upregulated genes in each spot were subjected to pathway enrichment analysis using level 3 SCPs (Suppl. Table 3). **E**nrichment **S**cores (ES) of each spot’s top 30 SCPs were normalized to sum to 200 ES units. Using the networks of functionally interacting SCPs, we mapped the ES units of each center spot’s SCPs to those of its peripheral spots so that the total network distance between mapped SCPs was minimal (Fig. 3B). Since an SCP’s network distance to itself is zero, this approach maps an SCP to itself before mapping it to other SCPs. The number of mapped ES units for each SCP pair was averaged across peripheral and center spots, weighted by spatial distance, to obtain a tissue section (i.e. slice) –level score. We then identified functionally and spatially significant SCP pairs by comparing the slice-level scores with those obtained after shuffling the labels of SCPs expressed on the same slice in the MBCO networks or spot positions on the slice, respectively (empirical p-value ≤ 0.1).

**Figure 3.**
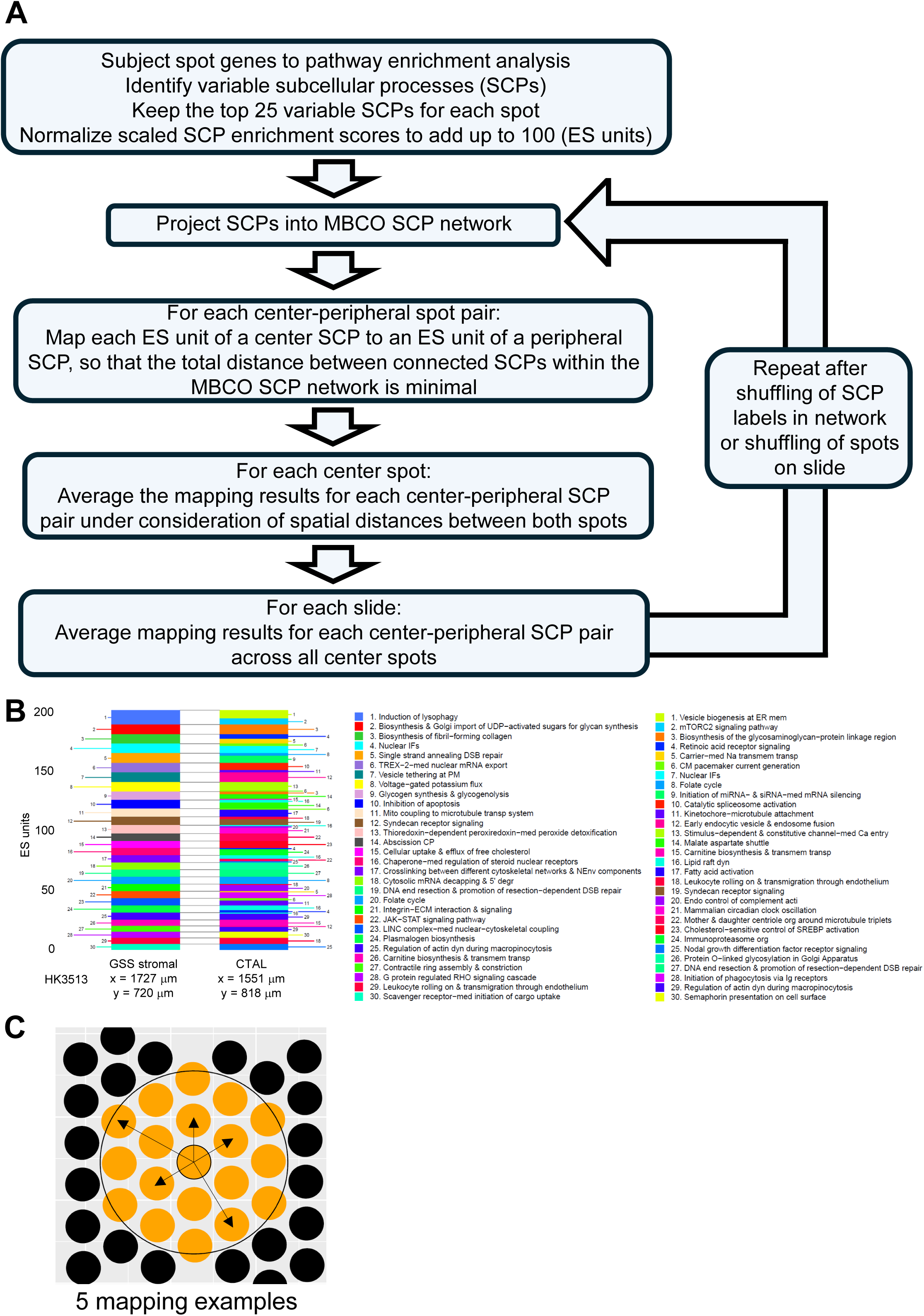
Functional relationships between SCPs expressed in neighboring spots. **(A)** Our pipeline compares functional relationships between SCPs expressed in a center spot and all peripheral spots in its neighborhood with and without shuffling of SCP labels in MBCO SCP networks and of spatial spots on the slice. **(B)** To map SCPs expressed in two neighboring spots to each other, our pipeline first normalizes the enrichment scores (ES) of the top 30 SCPs expressed in the same spot to add up to 200 ES units. SCPs of the two spots are mapped to MBCO SCP networks, followed by calculation of all shortest pathways between the SCPs of both spots. The Hungarian algorithm is used to connect each ES unit of one spot with one ES unit of the other spot, so that the sum of all connected SCPs is minimal. Colors and numbers label SCPs as indicated in the left and right legend for the left and right bar. **(C)** Mappings between any two SCPs of the center spot and each peripheral spot are averaged under consideration of spatial distances. Mapping between any two SCPs are then averaged across all center spots that expressed the related center SCP under consideration of the number and distances of the peripheral spots of each center spot (not shown).

Before we searched for functionally related SCPs in adjacent spots, we first investigated if variable SCPs are selectively upregulated in spots annotated to the same cell type (Suppl. Table 4). Our results document varying SCP activities within cell types, as exemplified by selected SCPs (Suppl. Fig. 4A). For most SCPs, the standard variations in enrichment scores across spots annotated to the same cell type and across all spots on the same slide were similar (Suppl. Fig. 4B), indicating that differences in SCP scores are not explained by cell type annotation alone.

We used our algorithm to find functionally significant center-peripheral SCP mappings. For an easier comparison of mapping results between different center SCPs, we normalized all averaged ES units for each center SCP to add up to 100 (Fig. 4A). We use the term normalized ES units to describe these values in the following analyses. Pairwise correlation analysis followed by hierarchical clustering of center (source) and peripheral (target) SCPs groups center and peripheral SCPs by functional relationships, as shown for selected SCPs on a selected slice (Fig. 4A). Since this analysis clusters center or peripheral SCPs by similar mapping profiles, it documents that functionally related center SCPs are mapped to similar functionally related peripheral SCPs. Individually, the highest normalized ES units are obtained for identical center and peripheral SCPs, as expected from the zero-network distance of an SCP to itself. Summed across all mappings of a center SCP, however, the ES units carried by pairs of different but functionally significant SCPs exceed those carried by self-mappings, and in turn exceed those carried by pairs without functional significance (Fig. 4B).

**Figure 4.**
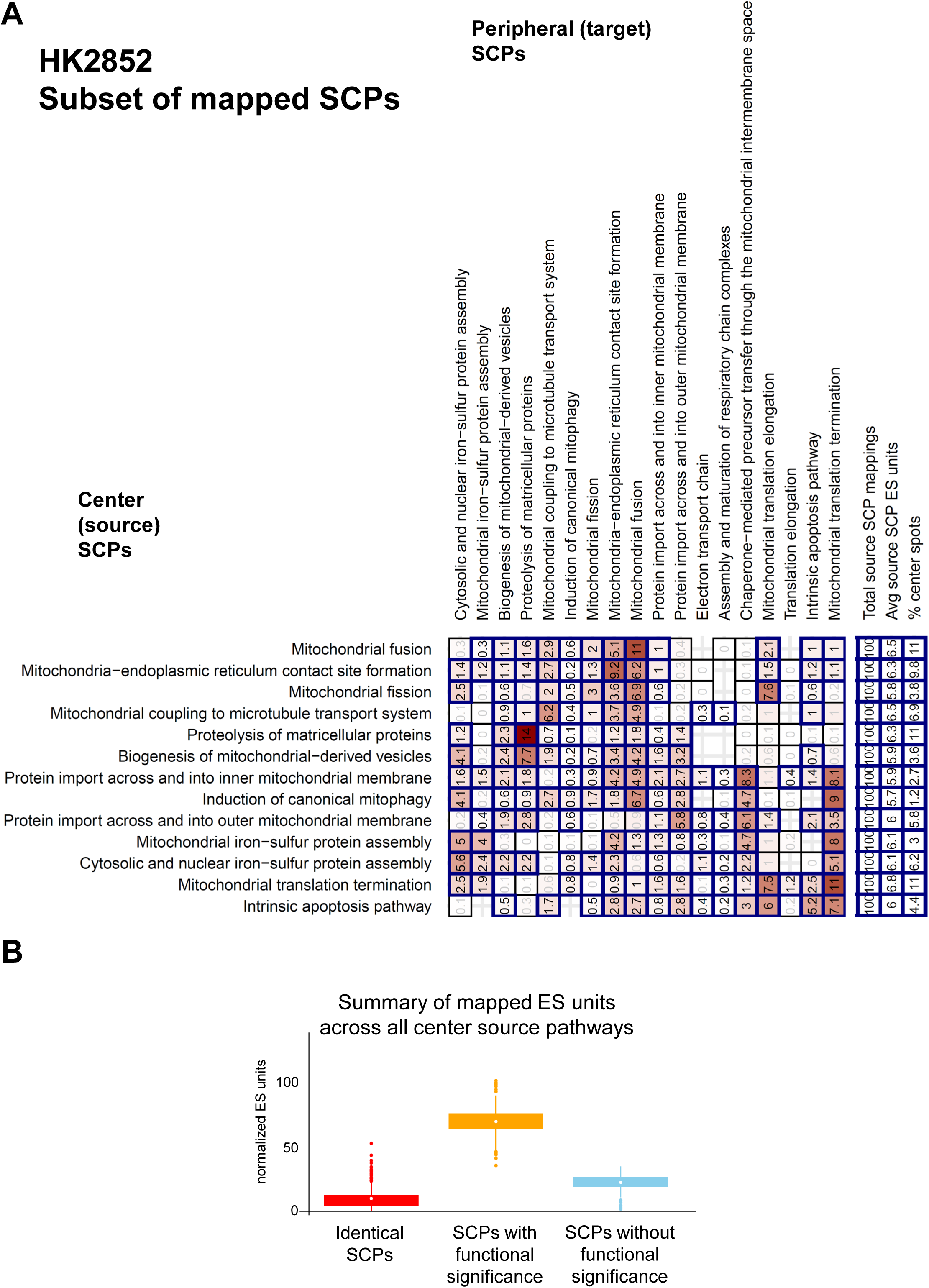
Example mappings between center and peripheral SCPs. **(A)** ES units between each center source SCP (rows) and all identified peripheral target SCPs (columns) were normalized to add up to 100 (fields show normalized ES units). Pairwise correlation analysis, followed by hierarchical clustering, grouped center and peripheral SCPs by shared functional relationships. The heatmap shows selected interactions for the slice HK2852. Framed fields with black numbers show interactions between the same center and peripheral SCPs or between SCPs with functional significance, as identified by comparing real results with results obtained after shuffling of SCP labels in the MBCO network 300 times (empirical p-value ≤ 0.1). Only labels of SCPs that were predicted in at least one spot on the same slice were shuffled. Three columns on the right show the sum of all normalized, initial ES units and the percentages of center spots that expressed the related center SCP. **(B)** For each center SCP we calculated how many normalized ES units map to the same peripheral SCP or SCPs that are or are not functionally related to the center SCP, as defined by empirical significance after SCP label shuffling.

The relationships described were consistently observed for all 13 slices representing biopsies from different patients and reference donors (Fig. 5A). Investigation of spatial significance documents that more than half of the self-mappings are spatially significant, while only a fraction of the functionally significant mappings between different SCPs are spatially significant as well (compare Figure 5A to 5B). In absolute numbers, however, spatially significant mappings are observed to a similar extent for self-mappings and mappings between functionally related SCPs across all slices.

**Figure 5.**
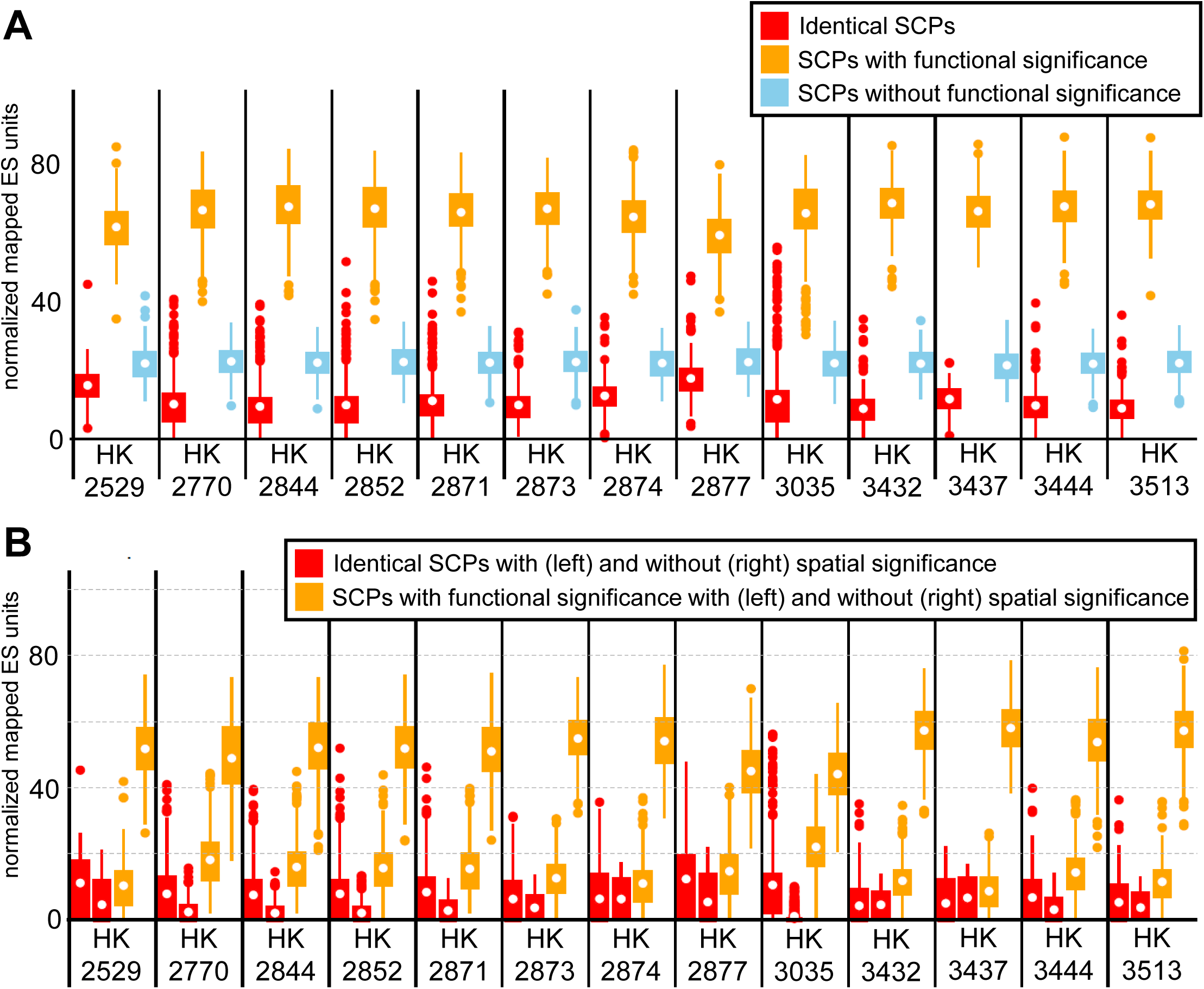
Functional and spatial relationships between mapped center and peripheral SCPs. **(A)** For each slice, we identified interactions between the same center and peripheral SCP (red), between SCPs with (orange) and without (blue) functional significance. White dots show mean values. **(B)** Spatial spots on each slice were shuffled 300 times. SCP interactions with spatial significance were identified by comparing ES units obtained with and without shuffling (empirical p-value ≤ 0.1). Results are only shown for interactions between the same center and peripheral SCP (red) or between SCPs with functional significance (orange). Left/right red and orange boxplots show ES units of SCPs with/without spatial significance.

Investigation of the functionally (Suppl. Tables 5) and spatially (Suppl. Tables 6) significant SCP-mappings or all functionally significant SCP-mappings with the highest normalized ES units (Suppl. Fig. 5A and B, respectively) identifies SCP pairs that describe ligand-receptor interactions, structural interactions and interactions suggesting cooperative or adaptive activities between similar and different cell types in neighboring spots (Fig. 6). At the same time, most predicted SCP pairs describe interactions that standard cell biology labels as intracellular interactions (Fig. 6), potentially indicating synchronized pathway activities between cells within neighboring spots. SCP pairs of both groups show spatial significance on some, but not all slices.

**Figure 6.**
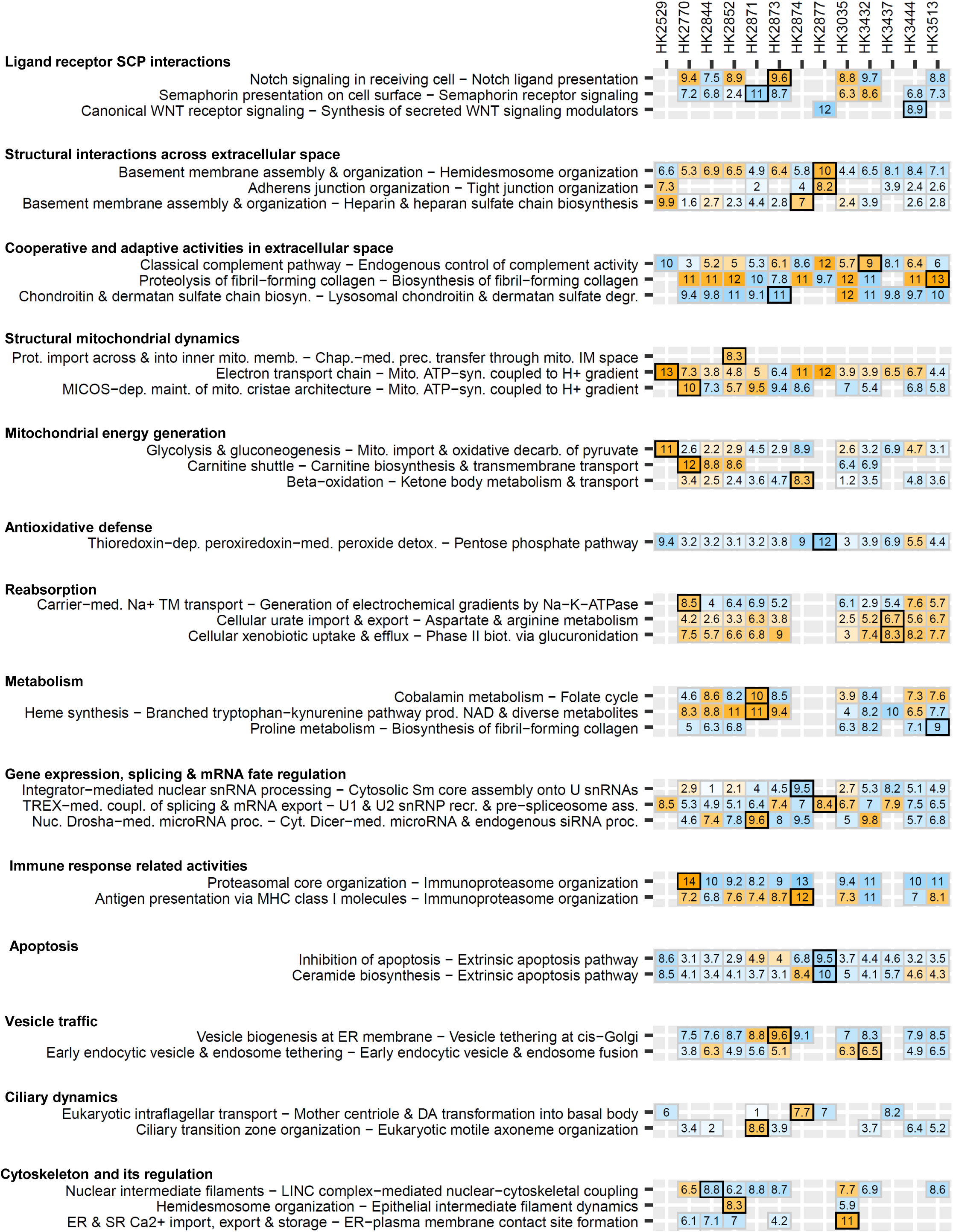
Selected Interactions between SCPs with functional and potential spatial significance. Identified interactions between center and peripheral SCPs with functional significance were ranked by mapped ES units. Selected interactions were grouped into six groups. Numbers show (normalized) ES units, numbers in brackets (normalized) ES unit ranks. Orange/Blue fields show interactions between SCPs with functional and spatial or only functional significance, respectively. Framed fields indicate selections for follow-up visualizations, e.g., SCP pair expression on the given slice. Ass.: assembly, biosyn.: biosynthesis, chap.: chaperone, coupl.: coupling, cyt.: cytosolic, degr.: degradation, dep.: dependent, det.: detoxification, ER: endoplasmic reticulum, IM: intermembrane, maint.: maintenance, med.: mediated, memb.: membrane, mito.: mitochondrial, nuc.: nuclear, org.: organization, prec.: precursor, proc.: processing, prod.: producing, prot.: protein, recr.: recruitment, SR: sarcoplasmic reticulum

The first group includes SCP pairs related to WNT signaling (‘Canonical WNT receptor signaling’ and ‘Synthesis of secreted WNT signaling modulators’), basement membrane and cell-matrix adhesion (‘Basement membrane assembly and organization’ and ‘Hemidesmosome organization’), complement regulation (‘Classical complement pathway’ and ‘Endogenous control of complement activity’), collagen turnover (‘Proteolysis of fibril-forming collagen’ and ‘Biosynthesis of fibril-forming collagen’), and chondroitin/dermatan sulfate turnover (‘Chondroitin sulfate and dermatan sulfate chain biosynthesis’ and ‘Lysosomal degradation of chondroitin and dermatan sulfate’). While some of these pairs suggest direct cellular interactions (e.g., Notch signaling) that are unlikely to reflect true cell-cell contacts across neighboring spots, most involve secreted or extracellular components, such as matrix proteins, proteoglycans and complement factors, whose effects may therefore extend across adjacent spots.

SCP pairs describing intercellular interactions span a broad range of cellular functions and often represent sequential steps within broader biological processes. These include mitochondrial dynamics and energy generation, e.g., ‘Protein import across and into inner mitochondrial membrane’ and ‘Chaperone-mediated precursor transfer through the mitochondrial intermembrane space’, ‘Glycolysis and gluconeogenesis’ and ‘Mitochondrial import and oxidative decarboxylation of pyruvate’ as well as ‘Carnitine shuttle’ and ‘Carnitine biosynthesis and transmembrane transport’. Other SCP pairs are involved in oxidative stress defense, such as ‘Thioredoxin-dependent peroxiredoxin-mediated peroxide detoxification’ and the NADPH regenerating ‘Pentose phosphate pathway’ or in reabsorption, e.g., ‘Carrier-mediated sodium transmembrane transport’ and ‘Generation of electrochemical gradients by Na-K-ATPase’. Metabolic SCP pairs included ‘Cobalamin metabolism’ and ‘Folate cycle’, which are functionally linked through folate-mediated one-carbon metabolism.^16^ ‘Heme synthesis’ was linked to ‘Branched tryptophan-kynurenine pathway producing NAD and diverse metabolites’, as it supplies heme prosthetic groups required for the catalytic activity of involved enzymes.^17^ The functional relationship between ‘Proline metabolism’ and ‘Biosynthesis of fibril-forming collagen’ is supported by the observation that dietary arginine contributes to proline synthesis and its subsequent incorporation into collagen in patients with proteinuria.^18^ Multiple SCP pairs were involved in gene expression, e.g. ‘Integrator-mediated nuclear snRNA processing’ and ‘Cytosolic Sm core assembly onto U snRNAs’, ‘TREX-mediated coupling of splicing and mRNA export’ and ‘U1 and U2 snRNP recruitment and pre-spliceosome assembly’ as well as ‘Nuclear Drosha-mediated microRNA processing’ and ‘Cytosolic Dicer-mediated microRNA and endogenous siRNA processing’. Examples of SCP pairs involved in immune responses and vesicular trafficking included ‘Antigen presentation via MHC class I molecules’ and ‘Immunoproteasome organization’, as well as ‘Vesicle biogenesis at ER membrane’ and ‘Vesicle tethering at cis-Golgi’, respectively. A pair related to ciliary dynamics linked ‘Eukaryotic intraflagellar transport’ with ‘Mother centriole and distal appendage transformation into basal body’. Our algorithm also identified SCP pairs associated with cytoskeletal organization and its regulation, including ‘Nuclear intermediate filaments’ and ‘LINC complex-mediated nuclear-cytoskeletal coupling’, ‘Hemidesmosome organization’ and ‘Epithelial intermediate filament dynamics’ as well as ‘Endoplasmic and sarcoplasmic calcium import, export and storage’ and ‘Endoplasmic reticulum-plasma membrane contact site formation’.

Visual inspection of the expression of SCP mappings (Fig. 7A, Suppl. Fig. 6) suggests that the mapped center and peripheral SCPs are mostly upregulated in neighboring, but not in the same spots. For each spot expressing a center SCP we quantified how many of its ES units could be mapped to the related target SCP within the center spot itself (Fig. 7B, Suppl. Fig. 6). Here, we included those SCPs that were not among the top 30 SCPs, by keeping them for further analysis and not using any significance cutoff. We compared those spots with the ES units averaged across all peripheral spots or averaged only across those peripheral spots that express the related target SCP. For all five example pairs (Fig. 7B), the mapped ES units averaged only across those peripheral spots that express the target SCP are at least twice as high than the ES units mapped within the center spot itself. Averaged across all peripheral spots, irrespective of whether they express the target SCP, the mapped ES units fall below the levels of the center spots. The signal therefore does not arise from the whole neighborhood, but specifically from a few neighboring spots, suggesting ultra local specialization that does not necessarily align with our empirical definition of a neighborhood by a circle of 220 μm.

**Figure 7.**
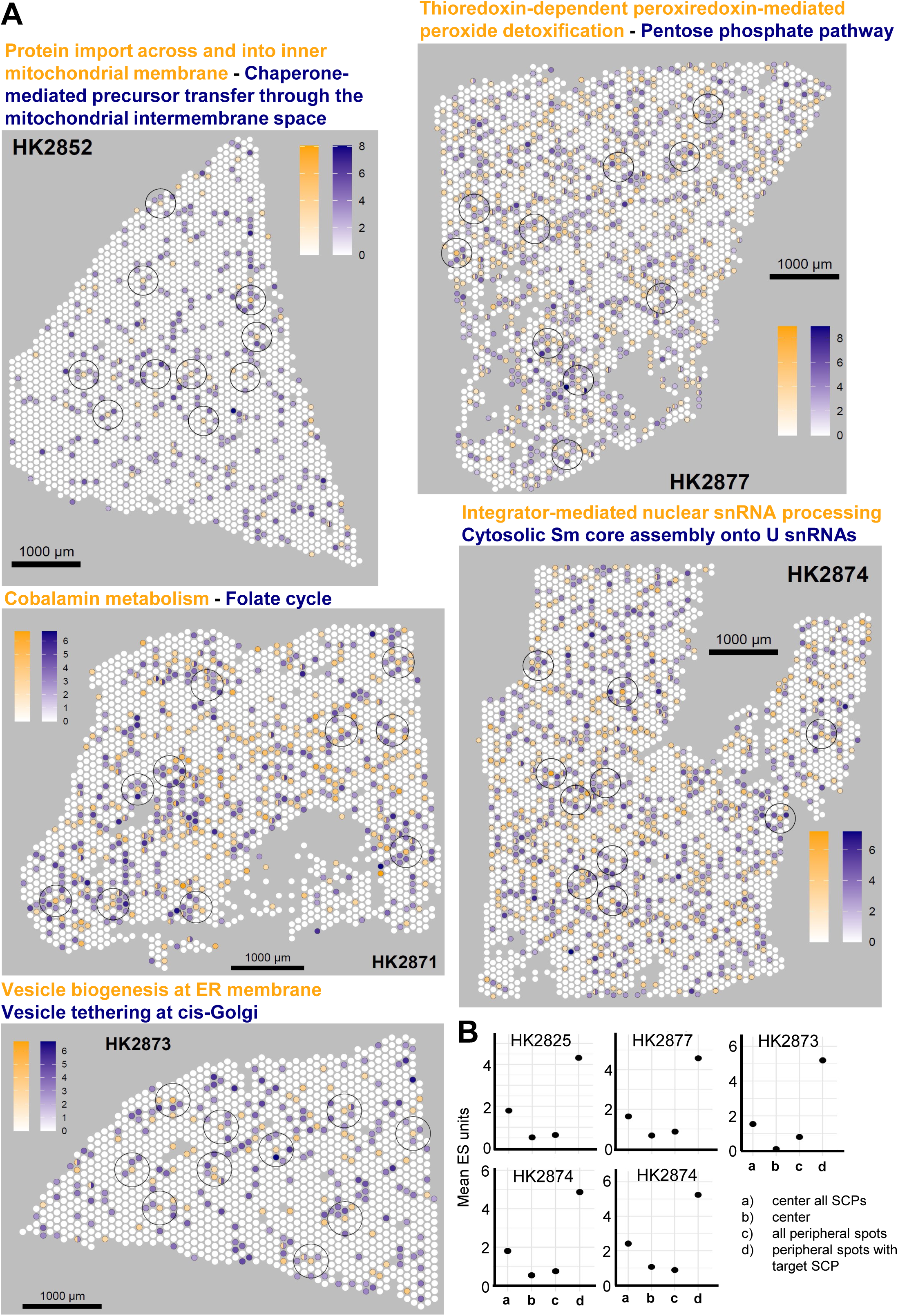
Spatial expression of selected SCP interactions with functional and spatial significance. Spots on selected slices are colored by the (not normalized, averaged) ES scores for given center (orange) and their identified peripheral interaction partner SCP (blue). Spots expressing both SCPs are split into halves and colored with both colors. Circles frame sites with the highest mapped ES score units. **(B)** For each center spot expressing the source SCP within a neighborhood that expresses the target SCP, we calculated how many of its ES units could be mapped to the target SCP within the same spot (‘center’). Because each ES unit maps to exactly one other ES unit, this value equals the smaller of the two SCPs’ ES units in that spot. We additionally calculated this potential mapping with all expressed SCPs considered instead of only the top 30 (‘center all SCPs’). For the top 30 SCPs, we also show the mapped ES units between center and peripheral spots, averaged either across all peripheral spots (as in the analysis pipeline) or only across peripheral spots expressing the target SCP. All four values were averaged across all center spots expressing the center SCP on the same slice.

Calculation of the same metrics across all SCP mappings with functional and spatial significance (Fig. 8A) and with functional significance irrespective of spatial significance (Fig. 8B) across all slices, confirms these results. Part of this difference is due to the algorithm itself. Because self-mappings carry zero network distance, a center SCP will generally only be assigned to a different peripheral SCP when that peripheral SCP is not already expressed in the center. Additionally, to avoid reporting SCP pairs that would not have been mapped had all SCPs in the center spot been considered, our algorithm removes any SCP pair whose potential mapping strength within the center spot exceeds 80% of the observed mapping strength between the center spot and the peripheral spots expressing the target SCP. For simplicity, we implemented this by setting both the functional and spatial permutation results for these pairs to non-significant, even though considering the target SCP in the center spot would probably have led to a cross-spot self-mapping. Potential mappings in the center for selected and discussed SCP pairs lie well below the 80% cutoff (Fig. 7, Suppl. Fig. 6B).

**Figure 8.**
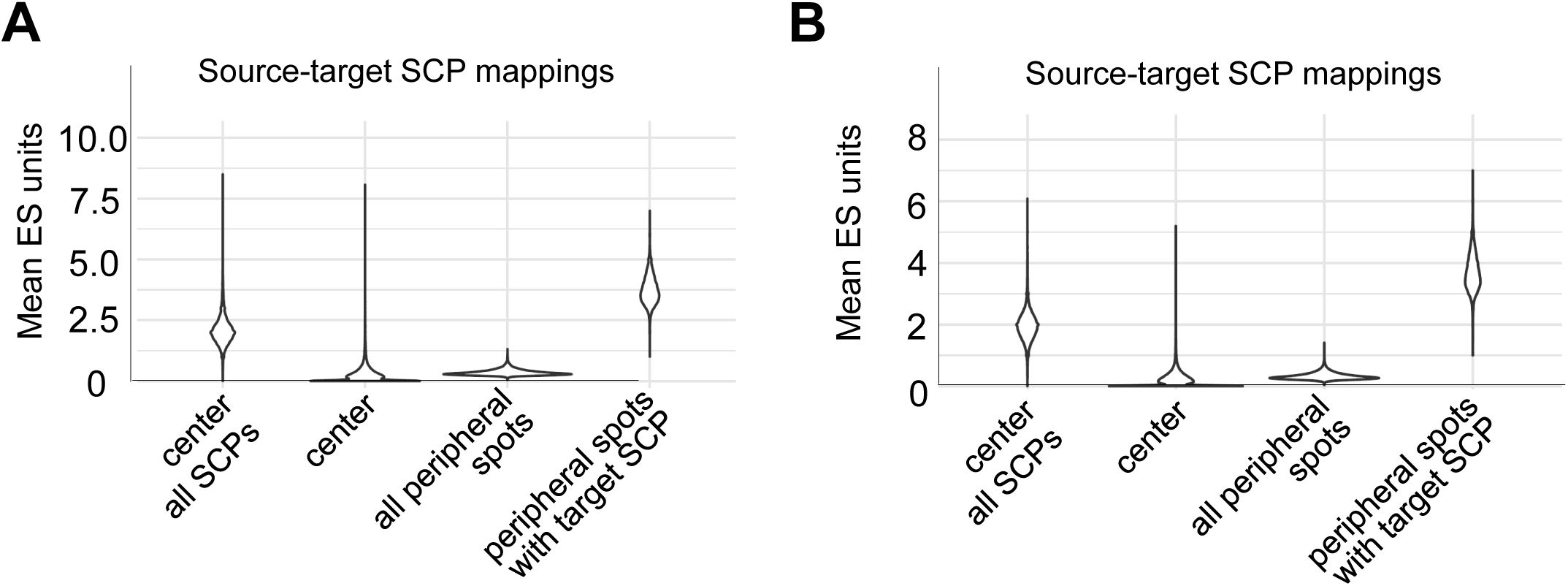
ES units of source-target SCP interactions across all slices. As outlined in figure 7, we calculated the number of ES units that could be mapped between both SCPs when considering either the top 30 SCPs (‘center’) or all expressed SCPs (‘center all SCPs’). Additionally, we calculated the mapped ES units between center and peripheral SCPs (based on the top 30 SCPs), averaged either across all peripheral spots or only across peripheral spots expressing the target SCP. Results are visualized separately for SCP interactions with **(A)** spatial and functional or **(B)** functional significance (with and without spatial significance).

Taken together, our analyses document that upregulated SCP activities across a tissue section show a higher correlation between functionally related SCPs than between identical SCPs. Though most SCP pairs did not exhibit spatial significance, SCP pairs that showed spatial significance reflected mappings between identical and functionally related SCPs to similar degrees. Our results suggest that functionally related SCPs rather than identical SCPs are upregulated in neighboring spots with similar or higher likelihood in the case of locally restricted or whole-tissue biological programs, respectively. Consequently, the characterization of physiological and pathological responses embedded in organization of cell types within tissues should benefit from the consideration of networks of interacting SCPs rather than building on lists of unrelated SCPs alone.

### Higher selectivity of SCP pairs for microenvironments than cell type pairs

To investigate which adjacent cell types express SCP pairs that were identified as functionally significant we repeated the analysis but averaged the results for each combination of center and peripheral cell type annotations independently. Cross-comparison of the mapped ES units for example SCP pairs suggests that they describe interactions between multiple cell type pairs, instead of being restricted to a single pair (Suppl. Fig. 7A, Suppl. Table 7). For example, the mapping of heme synthesis and branched tryptophan degradation SCPs is observed between a range of different cell types (Fig. 9A), suggesting that heme synthesis is not simply predicted within vascular erythrocytes.

**Figure 9.**
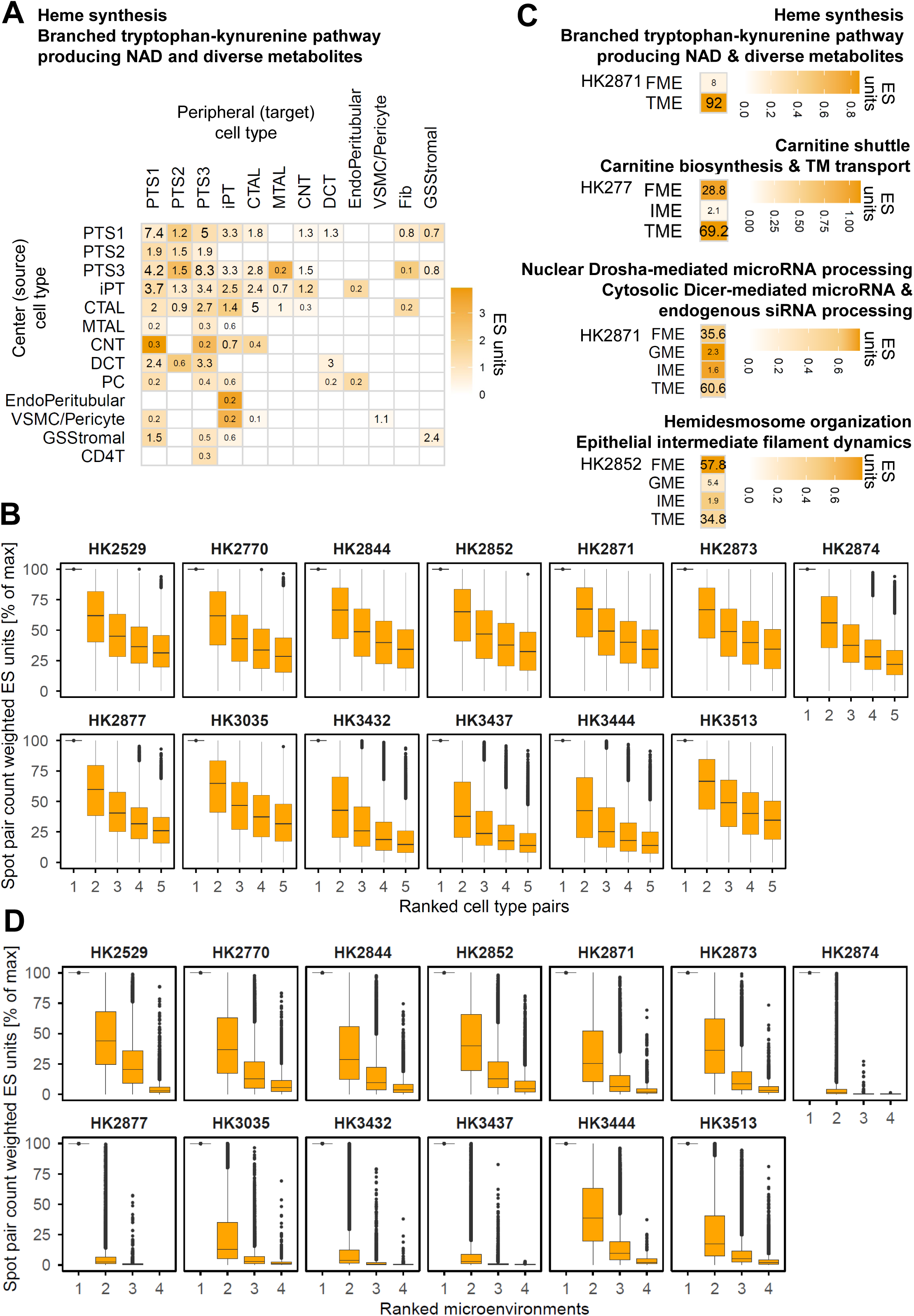
Cell-type selectivity of predicted cross-cellular SCP interactions. **(A)** We repeated our analysis pipeline, but this time averaged the results for each combination of center and peripheral cell type annotations independently. Numbers represent the number of center and peripheral spot pairs (in percent) that are part of that cell type pair, field colors are proportional to mapped (not normalized) ES units between all cell type pairs. **(B)** To systematically analyze potential cell-type selectivity, we multiplied the mapped ES units associated with each SCP and cell type pair with the number of center-peripheral spot pairs participating in that interaction. The resulting estimates of the overall contribution of each cell type pair were normalized by dividing it by the maximum overall contribution for the related SCP pair, followed by ranking of the cell type pairs. For each slice, we then combined the cell type pairs associated with all SCP pairs into groups of same ranks (ignoring which cell types are involved) to show the normalized overall contributions associated with the top 5 ranks. Similarly, we analyzed the distribution of SCP pairs within each microenvironment, shown for the **(C)** same and additional example pairs and **(D)** systematically for all SCP pairs. Note that we ignored mappings between different microenvironments, since these are restricted to the borders between them. TM: transmembrane

How well a given cell type pair represents an SCP pair can be quantified by two measures: the number of center spots participating in that cell type pair (numbers in Fig. 9A) and its (not normalized) ES units (color coded in Fig. 9A). For each SCP pair, we multiplied the number of spots and ES units for each of its cell type pairs, then expressed the results in percent of the highest product. Across all SCP pairs on most slices, these percentages show that the second– and third-strongest cell type pairs retain a large fraction of the strongest pair’s strength (Fig. 9B). Repeating the same comparison using only the number of center spots, or only the ES units, shows how much each value type contributes to the combined results (Suppl. Fig. 7B).

Four different microenvironments, i.e. the tubular, fibrotic, immune and glomerular microenvironments, have been initially identified in the original study.^6^ To investigate whether the identified SCP pairs are selective for a given microenvironment, we repeated the analysis, but this time restricted the averaging to center and peripheral spots assigned to the same microenvironment. Results show that SCP pairs are more selective to one of the four microenvironments than for any single pair of the 16 cell types, though they were not exclusively selective for a single microenvironment (Suppl. Table 8). This is illustrated by example pairs (Fig. 9C, Suppl. Fig. 8A), such as the SCPs involved in carnitine dynamics, microRNA processing or cell adhesion and cytoskeletal dynamics (Fig. 9C), and combined analysis of all SCP pairs on each slice (Fig. 9D and Suppl. Fig. 8B).

In summary, both analyses suggest that biological programs represented by spatially interacting SCPs can involve multiple cell types and can be related to a microenvironment of interest. Nevertheless, our analyses also find that at least some biological programs underlying tissue physiology and pathology are not restricted to a single microenvironment but can be identified across multiple microenvironments.

## Discussion

Extension and optimization of MBCO lead to a significant increase in SCPs and annotated genes. We have kept the general hierarchical organization of SCPs across four levels, with functional redundancies minimized among SCPs annotated to the same level. This organization principle makes downstream analyses easier, as also demonstrated in this study. The additional population of MBCO with metabolites led to an improvement of the predicted interaction networks of functionally related SCPs. This is documented by the prediction of well-known interaction partners for multiple example SCPs that span a broad area of cell biology and by the quantitative confirmation of functional integrity for most liver, cardiac and kidney cell types.

### Locally enriched and tissue-wide functional coordination between SCPs

We used MBCO for the identification of functionally related SCP pairs in human kidney tissue ^7,6^ that are upregulated in neighboring spots, if compared to all other spots on the same tissue slice. Initial analysis suggested that for many SCPs expression patterns are not cell-type-selective. Defining each neighborhood by a circle with a 220 μm radius, we could document that most center SCPs are either supported by expression of the same SCP, or –and to a much larger extent-by the expression of functionally related SCPs in peripheral spots. Spatial enrichment could be documented for more than half of the self-mappings and for a fraction of the mappings between functionally related SCPs. Nevertheless, if directly compared, spatially significant mappings are observed to a similar extent for self-mappings and mappings between functionally related SCPs across all slices. Spot shuffling asks whether an SCP pair is concentrated in a particular region of the tissue section. A pair that is coordinated uniformly across the tissue can therefore be functionally significant without being spatially significant. Loss of spatial significance can also arise when the pair is concentrated within a microenvironment large enough that shuffling does not disrupt it. The two types of significance thus separate local-scale coupling from tissue-scale coordination, and our analyses indicate that the latter is the more common mode. We show that the interaction strength between center and peripheral SCPs largely exceeds the potential interaction of the same SCPs within the center spot. This is not surprising, since our algorithm will only map a peripheral SCP to a different center SCP, if the peripheral SCP is not also upregulated in the center. Additionally, we find that mapped peripheral SCPs are upregulated in only a few neighboring spots, indicating more selective functional coordination than assumed by our circular neighborhood definition.

SCP pairs were identified while ignoring cell type and microenvironment annotations. Post hoc analyses documented that coordinated upregulation of functionally interacting SCPs extended across multiple cell types. Additionally, we documented a tendency of SCP pairs to be linked to a single microenvironment, though not exclusively and with large variation.

### Coordinated expression of SCPs interacting across the extracellular space

Investigation of predicted top SCP pairs identifies SCPs that facilitate structural tissue integrity (e.g. basement membrane and cell-matrix adhesion SCPs), cellular communication (e.g. WNT signaling and modulation), cellular cooperation (e.g. ECM synthesis and processing SCPs) and cellular adaptation (e.g. complement synthesis and inhibition). These examples suggest that functionally related SCPs are asymmetrically active in neighboring spots to generate a tissue-selective function. However, other SCP pairs, e.g. adherens and tight junction organization that –forming parts of the junctional complex-^19^ are normally expressed in both interacting cells and require direct cellular contact, propose phase shifts in translational renewal activities of otherwise symmetric SCP activities. In agreement with asymmetric activities of functionally related SCPs, we have already suggested cooperative activities during kidney fibrosis between neighboring failed-repair proximal tubule cells and cortical myofibroblasts.^15^ Such division of labor could establish a safety mechanism to ensure scar formation only around renal tubules with unrepairable damage. Here, our systematic analysis suggests that the expression of SCPs affecting the extracellular tissue environment and integrity is generally coordinated between neighboring spots, either in locally enriched spots or across the whole slice.

### Coordinated expression of intracellularly interacting SCPs

In addition to SCP pairs that interact within or across the extracellular space, we predict SCP pairs that standard cell biology labels as intracellular interactions. Among those are SCPs pairs involved in mitochondrial dynamics and energy generation, one-carbon, amino acid and urate metabolism, gene expression, proteasomal, vesicle traffic and ciliary dynamics. These findings could represent cooperation between cells in neighboring spots for intracellular tasks. They could potentially cooperate by exchanging cellular mRNA or proteins via extracellular vesicles,^20^ mitochondria via microvesicles ^21,22^ or metabolites for metabolically coupled neighboring spots. Metabolite exchange is well described for neurons and astrocytes, e.g. for cross-cellular coupling of glycolysis and oxidative phosphorylation,^23–25^ as well as for tumor cells and their environment.^26^ As an alternative interpretation, the cross-cellular SCP interactions could indicate synchronized symmetric activities between neighboring spots with shifted translational renewal activities, as also suggested by the SCP pair of adherens and tight junction organization. Another possibility is that the SCP pairs represent a spatial continuum or trajectory in cellular state activities,^15^ where the spatial progression of SCP activities mirrors their functional dependencies. We have observed such a coordination pattern for the temporal organization of SCP activities during neurite outgrowth.^27^ However, SCP expression patterns on selected slices do not suggest organization along spatial gradients in the human kidney tissue studied here.

### Comparison of our approach and results to related concepts in the spatial transcriptomics literature

Our results complement well-studied receptor-ligand interactions between cell types.^7–9,28–30^ Cross-cellular signaling indicates upstream coordination of cellular activities within a region, while our results indicate that additional cooperation and coordination at the level of downstream extra– and intracellular effector functions can also occur.

Two recent studies used machine learning to identify neighborhoods by the predictability of a center spot’s features, such as signaling pathways, from features expressed within the center itself, its immediate neighborhood and neighborhoods defined by more distant spots.^10,11^ Unsurprisingly, the strongest predictor of most features within the center proved to be the features of the center itself. Our study differs from those studies, since our study aims at the identification of functional relationships between spatially adjacent effector SCP sets that can give rise to whole-tissue functions.

Cooperation, adaptation and synchronization between neighboring spots, either locally enriched or distributed over the whole slice, aligns with the recently introduced concept of “functional communities”.^31^ This concept prioritizes intercellular dynamics that enable the “execution of coordinated biologic programs” over the simple cellular composition of a spatial niche as the driver of organ pathology. Functional similarities between adjacent spots, as identified in our study, could help define functional regions of an organ using sets of interacting SCPs rather than by cell type composition alone.

### Limitations

For the interpretation of our results some limitations should be considered. SCPs pairs in neighboring spots that were identified with functional and with or without spatial significance could still be part of different intracellular programs that might have been activated by the same trigger but are otherwise unrelated to each other in the context of a specific whole cell function. For evaluation of such possibilities, a deeper multiscale analysis of intra– and intercellular SCP network relationships needs to be developed. A first step could be to search for SCP pairs within each center spot. Such an analysis depends on penalization of self-mapping, which we explicitly allowed here. Existing or missing functional similarities between identified functionally significant intra– and cross-cellular SCP partners could help separate independent from coordinated activities.

Our analysis was restricted to the 30 SCPs with the highest enrichment scores (ES) in each spot. To assess how strongly each source-target mapping depends on neighboring spots, we calculated the number of ES units of the source SCP that could be mapped to the target SCP within the center spot alone, considering all expressed SCPs. We then applied an additional filter: a mapping was retained only if this within-spot potential did not exceed 80% of the mapping observed across spots. More stringent cutoffs would lead to an enrichment of SCP pairs that indicate stronger cross-cellular dependencies. At the same time, the top 30 SCP restriction also applies to neighboring spots and may reduce the observed mapping of source-target SCP pairs across spots, an effect that is more difficult to quantify.

Our definition of the neighborhood of a center is based on a circle with a radius of 220 μm. While this arbitrary definition is a reasonable size for a spatially interacting region,^32,33^ it most likely represents a simplification of real biology, sometimes by ignoring natural boundaries, such as the Bowman’s capsule, and directional interstitial and fluid flow.^34^ Additionally, because each spot captures multiple cells, contact-dependent interactions such as Notch ligand-receptor signaling may occur within a single spot and therefore may not be reflected in coordinated expression of related SCP pairs across adjacent spots. We did not exclude interactions between tissue cells and potentially bypassing cells within blood vessels, though this should only be a minor fraction of all interactions. More sophisticated definitions of a neighborhood might change the spatial significance for a given SCP pair. Additionally, the spatial significance for an SCP pair of interest is calculated by spot shuffling and should depend on the number of spots expressing either SCP. If, for example, SCPs forming a pair are predominantly expressed within one of the four microenvironments identified in the original study,^6^ the size of the microenvironment could influence the spatial significance observed after spot shuffling. One also should consider that the accuracy of cell-type-selective SCP expression profiles and interactions could be decreased, since the final cell type annotation could represent the most abundant cell type in a spot rather than being representative of all cell types.

One additional limitation concerns the structure of the MBCO network itself. Some SCPs display a higher connectivity than others. This could be due to degree-distribution of certain nodes. Shuffling SCP labels preserves the connectivity distribution of the network but assigns each label to a randomly chosen node, so the null distribution for any given pair is generated from nodes of average connectivity. Central SCPs may therefore reach significance in part because of their position in the network rather than their behavior in the tissue. Nevertheless, biological networks are often scale-free,^35–38^ so that differences in the connectivity across SCPs can –at least partly-reflect real biology. Identification of biologically reasonable SCP pairs supports this interpretation.

### Conclusions

In summary, we use MBCO SCP networks to identify cooperative, adaptive or synchronized effector SCP activities that are either asymmetrically expressed across neighboring tissue spots or exhibit shifted translational renewal or biogenesis patterns. Building on our approach, the integration of inter– and intracellular functional SCP networks can be a starting point for the definition of functional multicellular neighborhoods that form the basis for whole-tissue physiology and pathology.

## Methods

### Population of MBCO with genes and metabolites

To annotate genes and metabolites to MBCO SCPs, we used a slightly updated version of our population algorithm that combines PubMed abstract text mining with statistical enrichment analysis.^1^ Briefly, we defined a PubMed search query for every SCP. Obtained abstracts were screened for genes (including protein complexes and gene families), metabolites (including metabolite groups), diseases, confounding terms as well as protein domains. Though we focused only on genes and metabolites in the following, we screened for the other biological entities as well, since their consideration allows our algorithm to reduce misinterpretation of those entities as genes or metabolites. In contrast to our previous algorithm, we restricted our screening efforts for metabolites to only those text passages that were labeled as metabolites by the NIH text mining system PubTator.^39^ For each set of SCP-selective abstracts, our previous and current algorithm counted how many abstracts mention a biological entity at least once. Abstract counts were subjected to two rounds of enrichment analysis using Fisher’s exact test, with the combined abstract counts obtained for either those SCPs within the same level or with the same parent as a background set. In contrast to our previous algorithm, we did not apply any cutoffs but subjected all identified SCP gene or metabolite annotations to our computer-assisted manual validation pipeline. All SCP gene annotations of the initial version of MBCO were added to either the same or related SCPs in the enhanced version of MBCO, followed by manual reevaluation. Each SCP gene or metabolite annotation is supported by at least one reference. Additionally, genes were added to SCPs of levels 3 and 4 after expert curation from selected reviews and UniProt ^40^ or NCBI gene summaries,^41^ that are also given on the GeneCards website.^42^ Candidate gene-SCP annotations proposed by GPT-6 (OpenAI, ChatGPT) were evaluated and accepted only when literature references supporting the annotation could be added.

All annotated genes and all genes identified in at least one PubMed abstract were defined as the MBCO background genes.

### Identification of functional SCP interactions

Our previous and current population algorithm calculates the selectivity of an SCP gene or metabolite annotation using Fisher’s exact test and all genes or metabolites annotated to all SCPs of the same level (or with the same parents) as the background set. The selectivity score is given by the –log_10_(p-value). The nature of the algorithm makes SCPs of the same level (or with the same parents) compete for genes and metabolites. The more abstracts mention a gene for one SCP, the less significant is the association of that gene with another SCP. This allows us to predict functional relationships between SCPs of the same level by leave-one out analysis. Our published algorithm iteratively removes each level 2 and level 3 SCP from the ontology and quantifies the significance changes for gene and metabolite annotations calculated for each remaining SCP. Our previous algorithm focused on significance rank changes, while our current algorithm focused on the significance scores (i.e., –log_10_(p-values)) themselves. In addition, we here only considered significance scores calculated with all SCPs of the same level as a background set and ignored scores for all SCPs with the same parent. To normalize those changes, we here used the sum of all significance scores in the left-out SCP as a reference. As previously applied as well,^1^ only genes manually validated as true positives for the removed SCP were considered. In case of metabolites, we only considered those metabolites that were annotated as true positives to both, the removed and remaining SCPs. The calculated interaction scores for each pair of removed and remaining SCPs based on genes or metabolites were weighted by 0.7 and 0.3, respectively, and then summed up. Results obtained for the same SCP pair where each SCP can either be the removed or the remaining SCP were combined by adding up their interaction scores.

To ensure that well-known interactions, i.e. between the SCP pairs “Beta-oxidation” and “Citric acid cycle” as well as “Mitochondrial import and oxidative decarboxylation of pyruvate” and “Glycolysis and gluconeogenesis” were identified by our algorithm, we increased the abstract counts for acetyl-CoA and pyruvate within the related SCPs, so that they constitute 10% of the total abstract counts of those SCPs.

### Quantification of cellular functional integrities

Marker genes of cardiac,^13^ liver ^12^ and kidney ^14^ cell types were calculated using the Seurat functionality FindMarkers ^43^ with all cells of the other cell types of the same study as a reference. Top 500 marker genes were subjected to enrichment analysis using MBCO level 3 SCPs and Fisher’s exact test. We defined the experimental background genes as those genes that were expressed in at least one cell of the same study, MBCO background genes were extracted from the ontology. Only SCP genes and marker genes that were also part of the experimental and ontology background gene sets, respectively, were kept for enrichment analysis. Significant SCPs (p-value ≤ 0.05) of each cell type were mapped onto MBCO SCP networks, followed by calculation of all shortest pathways between any (source and target) pair of SCPs using Dijkstra’s algorithm. Shortest pathways starting at each source SCP were averaged, after weighting them by the –log_10_(p-values) obtained for the target SCPs. Average shortest pathways were averaged again across all source SCPs, weighting them by the –log_10_(p-values) of the source SCPs. Resulting double-average shortest pathway lengths were defined as 100%. For each dataset, we repeated the analysis 100 times after SCP label shuffling in the MBCO network. Only labels of SCPs predicted for at least one cell type of the same dataset were shuffled. Double-average shortest pathway lengths obtained for each shuffled network were normalized towards the double-average shortest pathways obtained for the real network. Empirical p-values were calculated as the fraction of the 101 results, comprising the real network and the 100 shuffled networks, whose double-average shortest pathway length was equal to or smaller than that of the real network (minimum possible p-value = 1/101).

### Mapping of SCP activities between neighboring spots

We used the MBCO SCP network to analyze the functional relationships between SCPs expressed in neighboring spots on reference and diseased kidney tissues. For quality control, we kept only spots with at least 100 total UMIs and 200 detected genes as well as three median absolute deviations (MADs) below the median of the log_10_-transformed UMIs or gene counts. Additionally, we removed spots, if one gene accounted for more than 20% of their UMIs. We also implemented a cutoff that removed spots exceeding three median absolute deviations above the median mitochondrial gene counts, although mitochondrial genes were not detected. After spot filtering, genes were removed if they were detected in fewer than 10 spots or if more than 50% of their normalized expression came from a single spot. Described spot quality control measures were suggested and implemented by Claude Opus 5.5, except that we lowered the minimum UMI counts from 500 to 100.

For each gene, we compared its gene expression within each spot against the average of all other spots on the same slide to calculate a log_2_ fold change. For each slice, we defined the experimental background genes as those genes that were expressed in at least one spot. MBCO background genes were extracted from the ontology. Only SCP genes and DEGs that were also part of the experimental and ontology background gene sets, respectively, were kept for the following analysis. SCPs with less than 5 remaining genes were removed. Genes of each spot that were upregulated with a log_2_ fold change of at least 0.5 were subjected to pathway enrichment analysis using the UCell algorithm^44^ and MBCO level 3 SCPs. Spots with non-zero counts for fewer than 5% of all genes were excluded. UCell’s ‘maxRank’ parameter was set to the median number of detected genes across the remaining spots, and ‘ties.method’ was set to “average”. We arbitrarily kept the top 30 SCPs expressed in each spot. Our following algorithm for the mapping of functionally related SCPs between any two spots quantifies the functional relationship between co-expressed SCPs in two adjacent spots. To simplify the implementation, we normalized the enrichment scores of all retained SCPs in each spot so that they summed to 200 enrichment score (ES) units, rather than introducing a conversion factor when comparing SCP expression between two spots. The enrichment scores of SCPs outside the top 30 were normalized using the same denominator, so that their potential influence, which would otherwise be missed, could be quantified at a later stage as well.

We defined the neighborhood or site of each (center) spot as all peripheral spots that lie within a range of 220 μm. SCPs predicted for each pair of center and peripheral spots were mapped onto the MBCO SCP network, followed by calculation of all shortest pathway lengths between any pair of center (source) and peripheral (target) SCPs, using Dijkstra’s algorithm.^45^ We used the Hungarian algorithm ^46^ (modified from ^47^) to map ES units between the center spot and each peripheral spot such that each center and each peripheral ES unit is mapped only once and the total length of the shortest pathways between related SCPs is minimized. For each center SCP, we then averaged the mapped ES units with each peripheral SCP across all peripheral spots. To account for spatial separation between the center and peripheral spots, the mapped ES units were weighted using a Gaussian radial basis (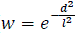, d: distance between center and peripheral spot; l = site; radius = 220 μm). To obtain a single value for each SCP pair per slice, these averages were then combined across all center spots on the same slice as a weighted mean, with weights reflecting the number and distances of peripheral spots associated with each center spot.

For all analyses, we only considered those center SCPs that were identified in at least 20 spots on the same slice or on the same slice within the same cell type or microenvironment (see below).

### Identification of functionally and spatially significant SCP mapping pairs

To identify mapped center and peripheral SCP pairs with functional or spatial significance, we repeated the analysis after shuffling the SCP labels in the MBCO network or the spots on each slice 300 times. Only labels of SCPs that were predicted for at least one spot on the same slice were shuffled. For each SCP pair we counted how often the double averaged mapped ES units obtained with or without SCP label or spot shuffling were equal or larger than the ES units obtained with the real network or on the real slice. SCP mappings with empirical p-values (minimum possible p-value: 1/301) equal or smaller than 0.1 were defined as functionally or spatially significant. For all SCP pairs in which the potential interaction between the source and target SCPs within the center spot exceeded 80% of their observed interaction with the peripheral spots expressing the target SCP (see below), we classified the functional and spatial results as non-significant. This classification is a simplification: had the target SCP expressed in the center spot been considered, the peripheral SCP would have been mapped to itself in the center.

### Investigation of SCP pairs

To compare peripheral SCP mappings across all center SCPs, we normalized all double-averaged ES units for each center SCP across all peripheral SCP mappings to add up to 100. We use the term normalized ES units in the following. Center and peripheral SCP mappings obtained for each slice were subjected to pairwise correlation analysis of mapped normalized ES units under real conditions, followed by hierarchical clustering.

For each center SCP, we split the peripheral SCP mappings into three groups: mappings to the same peripheral SCP (self-mappings), mappings to peripheral SCPs with and mappings to peripheral SCPs without functional significance. Normalized ES units for all center SCPs on the same slice annotated to each group were visualized in the same boxplots. We further split self-mappings and mappings with functional significance into those with and without spatial significance, prior to re-visualization.

SCP pairs on each slice were ranked by mapped normalized ES units and example pairs connecting different SCPs were selected for visualization of their expression values on example slices.

### Quantification of the potential interactions of source and target SCPs in the center

For each pair of different SCPs, we calculated how many ES units of the center SCP could be mapped to the peripheral SCP within the center spot itself. Since each ES unit of the source SCP is mapped to one ES unit of the target SCP, this value represents the minimum number of ES units predicted for the source or target SCP in the center. As described above, our main analysis was restricted to the top 30 SCPs with the highest expression values. To avoid missing potential mappings between a source SCP and a target SCP in the center spot, we also included all SCPs below this threshold. To quantify if mappings between center and peripheral SCPs are representative for the whole circular neighborhood or only involve single peripheral spots, we additionally calculated the doubly averaged mapped ES units between each center and peripheral SCP pair, considering only those peripheral spots that express the peripheral SCP, while ignoring all others. For simplicity, and unlike in the main analysis, we here averaged the ES units of each center-peripheral SCP pair across all center spots on the slice without weighting by the number of peripheral spots and their distances. As outlined above, for SCP pairs whose potential interaction within the center spot exceeded 80% of the observed interaction with the peripheral spots expressing the target SCP, we set both the spatial and functional permutation results to non-significant. As outlined above, this classification is a simplification: had the target SCP expressed in the center spot been considered, the peripheral SCP would have been mapped to itself in the center.

### Characterization of cell-type– and microenvironment-selective SCP pairs

To assess the selectivity of identified SCP pairs for cell type pairs, we repeated the analysis separately for each combination of center and peripheral spot cell types. For each center spot, we averaged the ES units of each SCP pair separately for each peripheral cell type, using only the peripheral spots annotated with that cell type. We then averaged these values for each SCP pair across all center spots of the same cell type, yielding one double-averaged ES unit value per center-peripheral cell type pair. In addition, we counted the number of center-peripheral spot pairs contributing to each SCP pair within each cell type pair and expressed this count as a percentage of all center-peripheral spot pairs on the slice. To assess the overall representation of a cell-type pair for a given SCP pair in terms of both intensity and frequency, we multiplied these two values to obtain spot-pair-count weighted ES units.

To examine the cell type pair selectivity of SCP interactions more generally, we identified, for each SCP pair on a slice, the cell type pair with the highest double-averaged ES units, the highest number of center-peripheral spot pairs, and the highest spot-pair-count-weighted ES units. We then expressed each value as a percentage of the corresponding maximum. For each SCP pair and measure, cell type pairs were ranked in descending order of the related value. The top-ranked values for all SCP pairs on the same slice were then visualized together.

The authors of the original study ^6^ identified four different microenvironments on their slices: the tubular, glomerular, fibrotic and immune microenvironment (TME, GME, FME and IME, respectively). To assess the selectivity of identified SCP pairs for microenvironments, we repeated the analysis as described above but considered only combinations between center and peripheral spots that were annotated to the same microenvironment.

### Supplementary Tables

**Supplementary Table 1**. **Cell-type-selective SCPs.** Enrichment results for **(A)** liver, **(B)** heart and **(C)** kidney cell types. Cell-type-selective marker genes were calculated using Seurat FindMarkers and subjected to enrichment analysis using MBCO level 3 SCPs and Fisher’s exact test.

**Supplementary Table 2. Cell-type-selective functional integrities**. For each (A) liver, **(B)** heart and **(C)** kidney cell type, pairwise distances between significantly expressed SCPs were calculated within the functional SCP networks. Distances were averaged after weighting them by the –log10(p-values) of the connected SCPs. A null distribution was generated by shuffling the network labels of those SCPs that were predicted in any cell type of the same dataset, followed by calculation of empirical p-values.

**Supplementary Table 3. Spot-selective enrichment results.** Upregulated genes on each spot, if compared to all other spots, were subjected to pathway enrichment analysis using the Ulm algorithm and level 3 SCPs. Top 30 pathways were kept for the slices **(A)** HK2529, **(B)** HK2770, **(C)** HK2844, **(D)** HK2852, **(E)** HK2871, **(F)** HK2873, **(G)** HK2874, **(H)** HK2877, **(I)** HK3035, **(J)** HK3432, **(K)** HK3437, **(L)** HK3444 and **(M)** HK3513.

**Supplementary Table 4. Cell-type-annotation-selective enrichment results.** Enrichment results were averaged for all spots annotated to the same cell type for the slices **(A)** HK2529, **(B)** HK2770, **(C)** HK2844, **(D)** HK2852, **(E)** HK2871, **(F)** HK2873, **(G)** HK2874, **(H)** HK2877, **(I)** HK3035, **(J)** HK3432, **(K)** HK3437, **(L)** HK3444 and **(M)** HK3513.

**Supplementary Table 5. Functional significance for SCP mappings.** SCP pair were identified and tested for functional significance as described in the methods section on slices **(A)** HK2529, **(B)** HK2770, **(C)** HK2844, **(D)** HK2852, **(E)** HK2871, **(F)** HK2873, **(G)** HK2874, **(H)** HK2877, **(I)** HK3035, **(J)** HK3432, **(K)** HK3437, **(L)** HK3444 and **(M)** HK3513.

**Supplementary Table 6. Spatial significance for SCP mappings.** Identified SCP pairs were tested for spatial significance as described in the methods section on slices **(A)** HK2529, **(B)** HK2770, **(C)** HK2844, **(D)** HK2852, **(E)** HK2871, **(F)** HK2873, **(G)** HK2874, **(H)** HK2877, **(I)** HK3035, **(J)** HK3432, **(K)** HK3437, **(L)** HK3444 and **(M)** HK3513.

**Supplementary Table 7. Selected SCP pairs between pairs of cell type annotations.** We repeated our analysis pipeline, but this time averaged the results for each combination of center and peripheral cell type annotations independently. The table contains the mapping statistics for selected SCP pairs for slices **(A)** HK2529, **(B)** HK2770, **(C)** HK2844, **(D)** HK2852, **(E)** HK2871, **(F)** HK2873, **(G)** HK2874, **(H)** HK2877, **(I)** HK3035, **(J)** HK3432, **(K)** HK3437, **(L)** HK3444 and **(M)** HK3513.

**Supplementary Table 8. Selected SCP pairs within the same microenvironment.** We repeated our analysis pipeline, but this time averaged the results only across center and peripheral spots that belonged to the same microenvironment. The table contains the mapping statistics for selected SCP pairs for slices **(A)** HK2529, **(B)** HK2770, **(C)** HK2844, **(D)** HK2852, **(E)** HK2871, **(F)** HK2873, **(G)** HK2874, **(H)** HK2877, **(I)** HK3035, **(J)** HK3432, **(K)** HK3437, **(L)** HK3444 and **(M)** HK3513.

## Acknowledgements

We thank Drs. Amin Abedini and Katalin Susztak for providing the final Seurat object of their spatial transcriptomics study ^6^ that we used to analyze our approach.

## Use of AI

For multiple SCPs, we used GPT-6 (OpenAI, ChatGPT) to obtain additional candidate gene annotations that were then manually evaluated and only used if supporting evidence could be found in the literature. References were searched with and without help of GPT-6. Some SCP names were discussed with GPT-6.

Quality control steps for spatial transcriptomics data were suggested and implemented by Claude Opus 5.5 (after minor modification).

Claude Opus 5.5 was used to write R code for some data analysis steps and visualization figures.

During the preparation of this manuscript, we used Claude Opus 5.5 to correct typos, grammar and style. After using this tool, we rechecked and rewrote the content as needed.

We take full responsibility for all content including that generated with AI-assistance.

## Code and data availability

Analysis code has been deposited at github.com/SBCNY/MBCO_SCP_interaction_adjacent_tissue_spots.

MBC Ontologies are deposited at mbc-ontology.org / github.com/SBCNY/Molecular-Biology-of-the-Cell.

