## Supplementary Figures 1-5 for "Functional cooperation, adaptation and synchronization of subcellular processes in cellular neighborhoods as inferred from spatial transcriptomics"

### Abstract

Tissue-level functions emerge from pathway activities distributed across neighboring cells. Yet analyses of spatial transcriptomics mostly focus on cell type composition or signaling through ligand-receptor pairs, leaving spatial relationships between effector processes undescribed. We inferred networks of functionally interacting subcellular processes (SCPs) that are independent of shared gene products and lie beyond sibling relationships defined in parent-child hierarchies. Using those networks for the analysis of published spatial transcriptomics datasets from human kidney biopsies, we identified functionally related SCP activities between a tissue spot and its neighbors. Identified pairs of adjacent SCPs indicate functional cooperation, e.g. basement membrane assembly and cell-matrix adhesion, functional adaption, e.g. complement synthesis and inhibition, and functional synchronization, e.g. folate cycle and cobalamin metabolism or pre-mRNA splicing and nuclear mRNA export. Two independent null models, shuffling of SCP labels within the network and shuffling of spots on the tissue slice, distinguish pairs that are functionally non-random and pairs that are additionally enriched within the immediate neighborhood, separating local from tissue-wide coordination. Our results suggest that tissue physiology reflects not only cell type composition but also functional coordination between neighboring regions.

**Supplementary Figures 1 – 5**

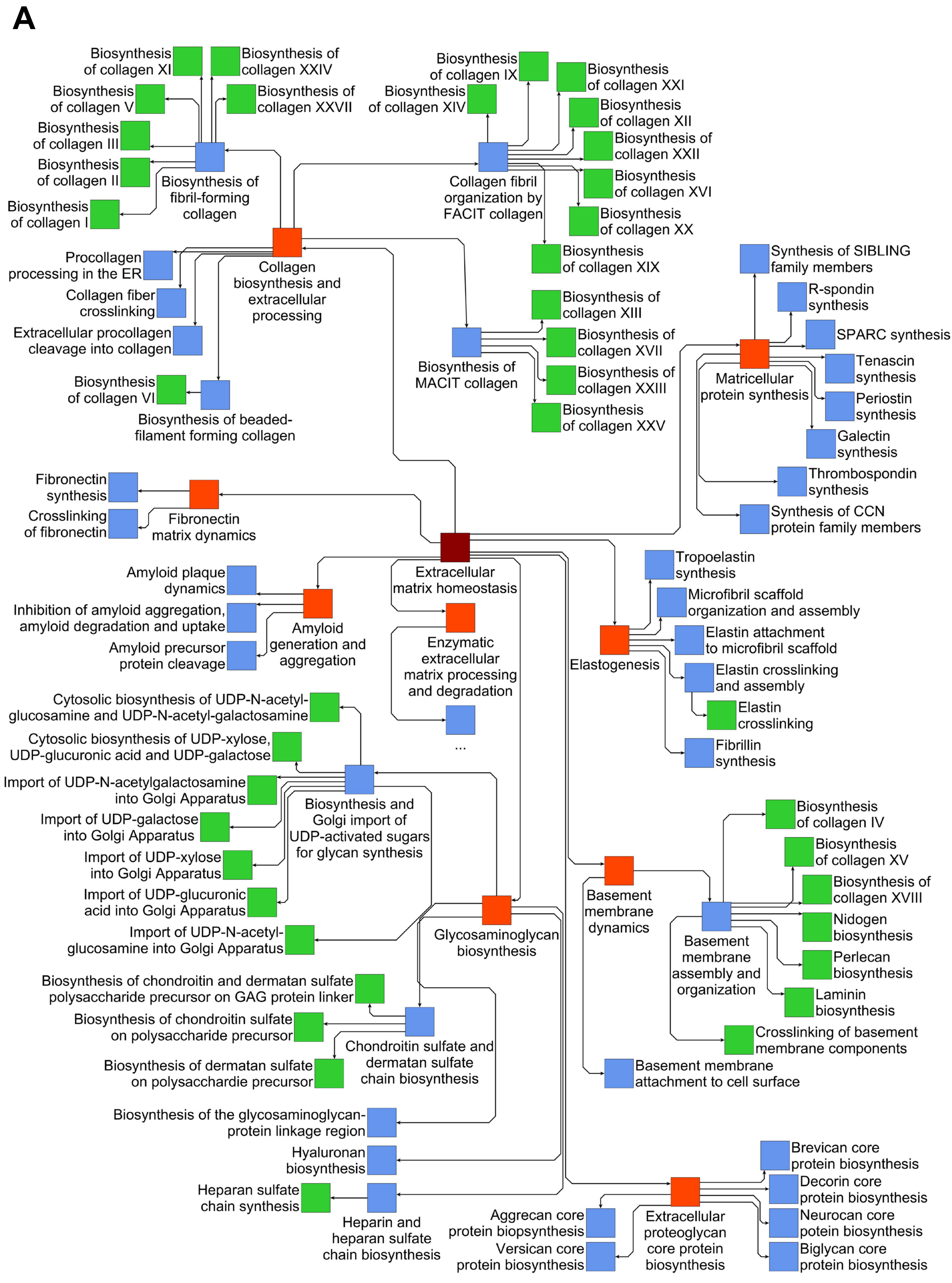


**Supplementary figure 1**

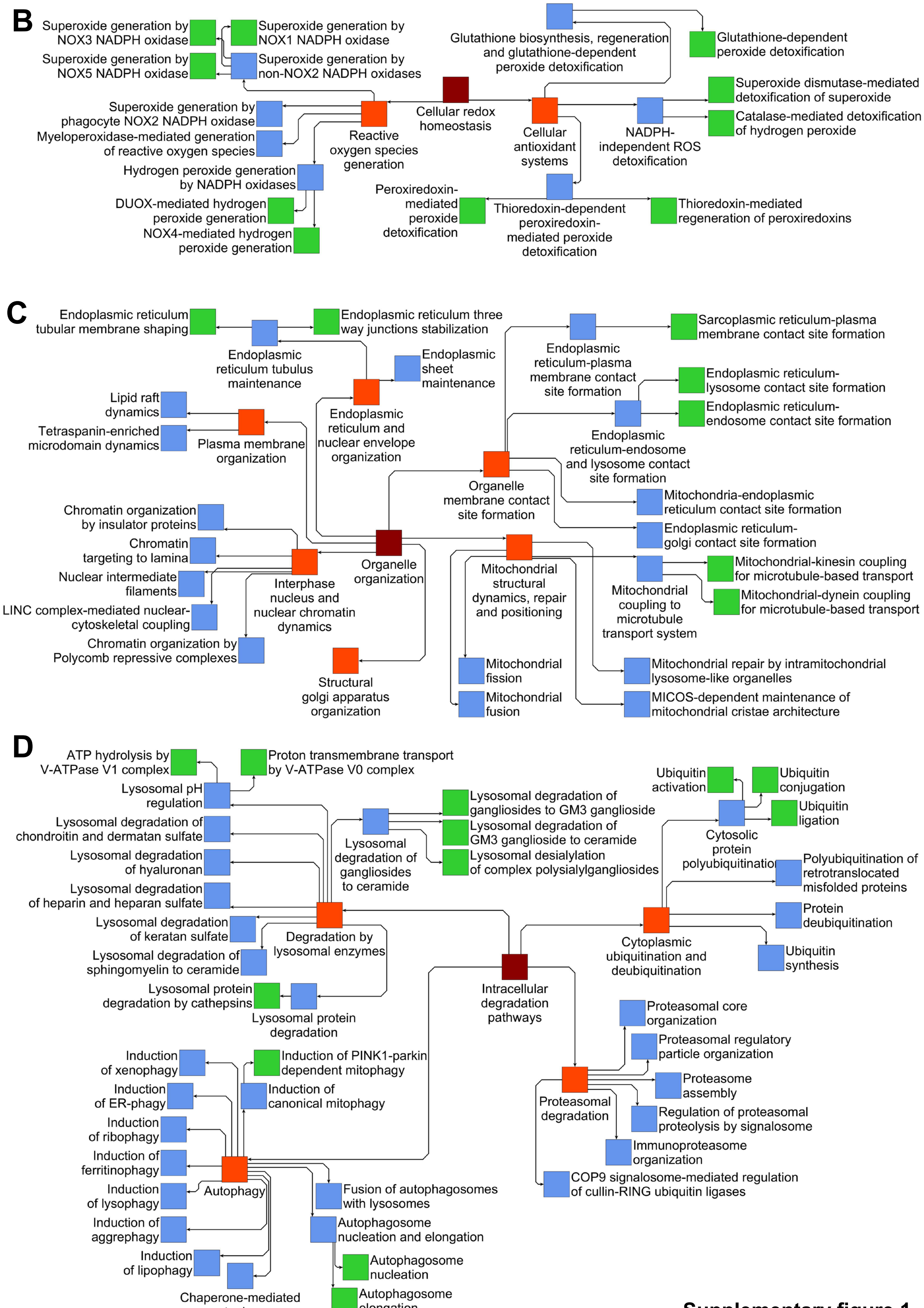


**Supplementary figure 1**

E

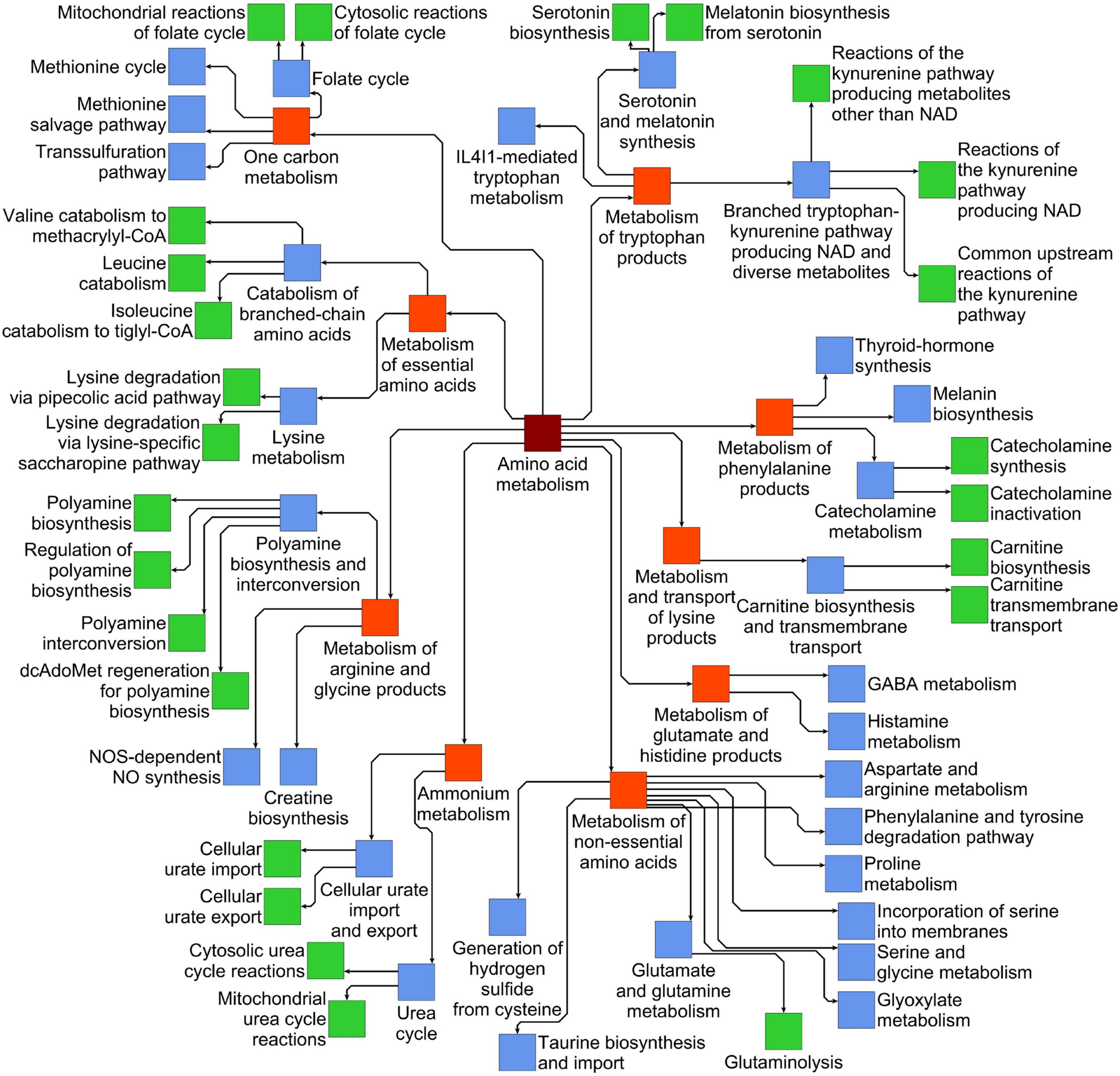


F

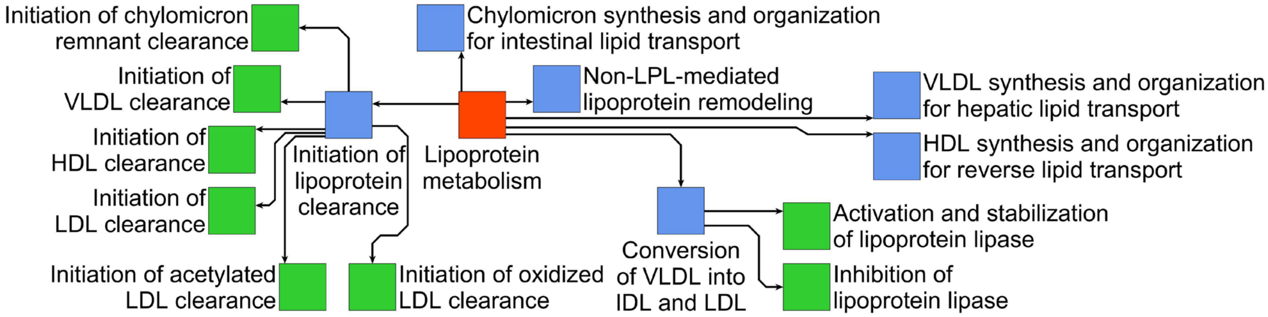


**Supplementary figure 1**

G

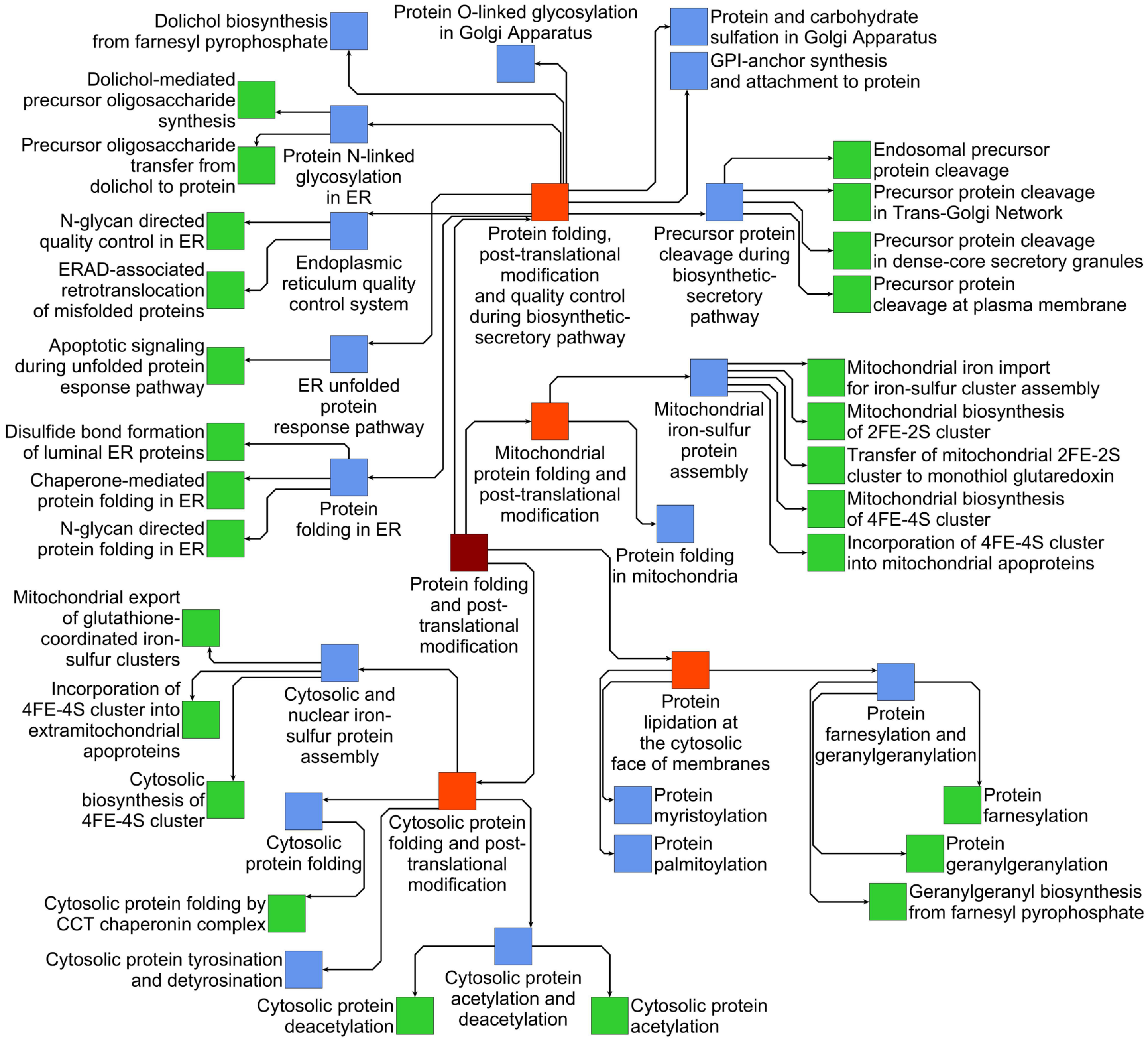


**Supplementary figure 1**

H

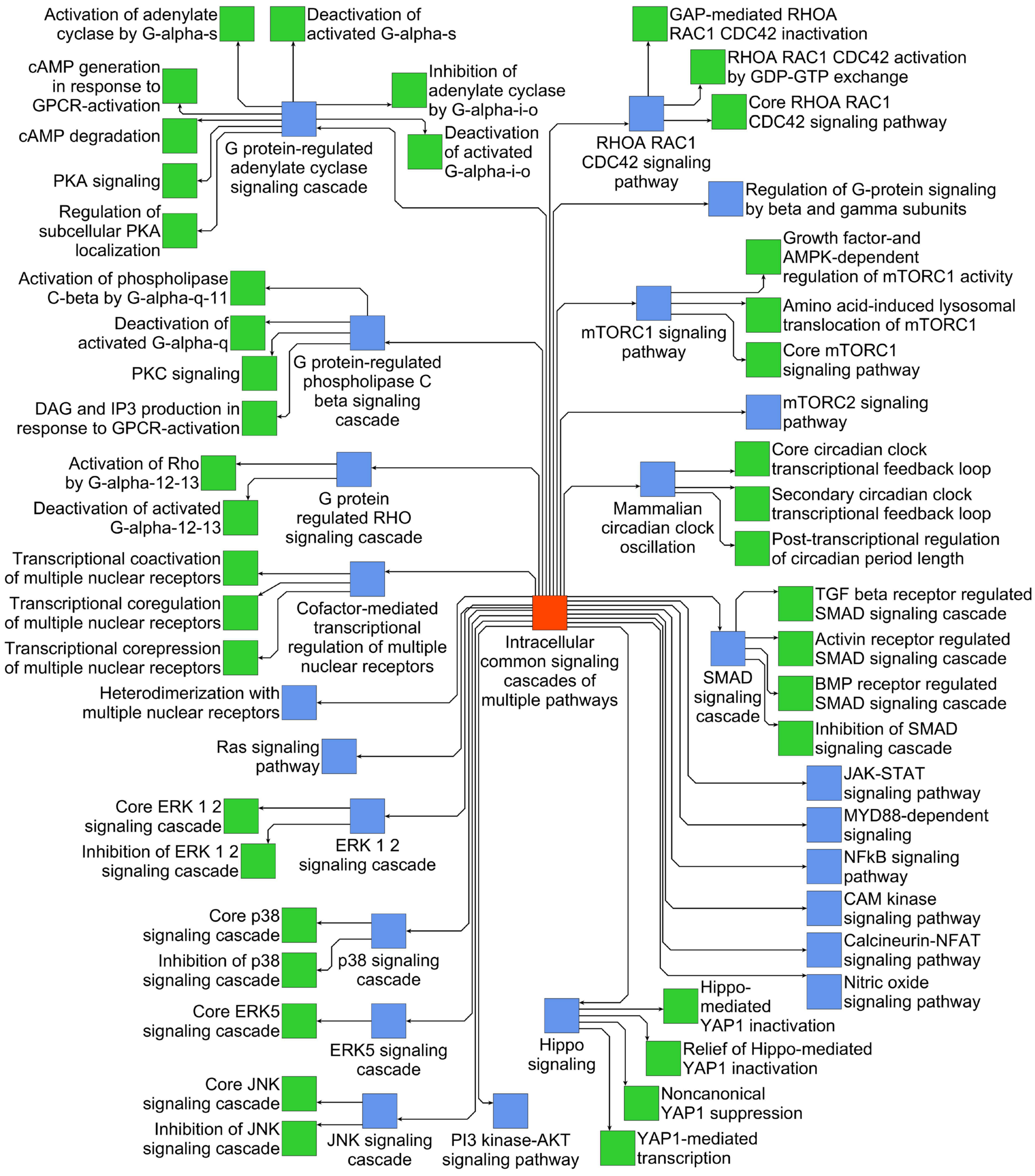


**Supplementary figure 1**

I

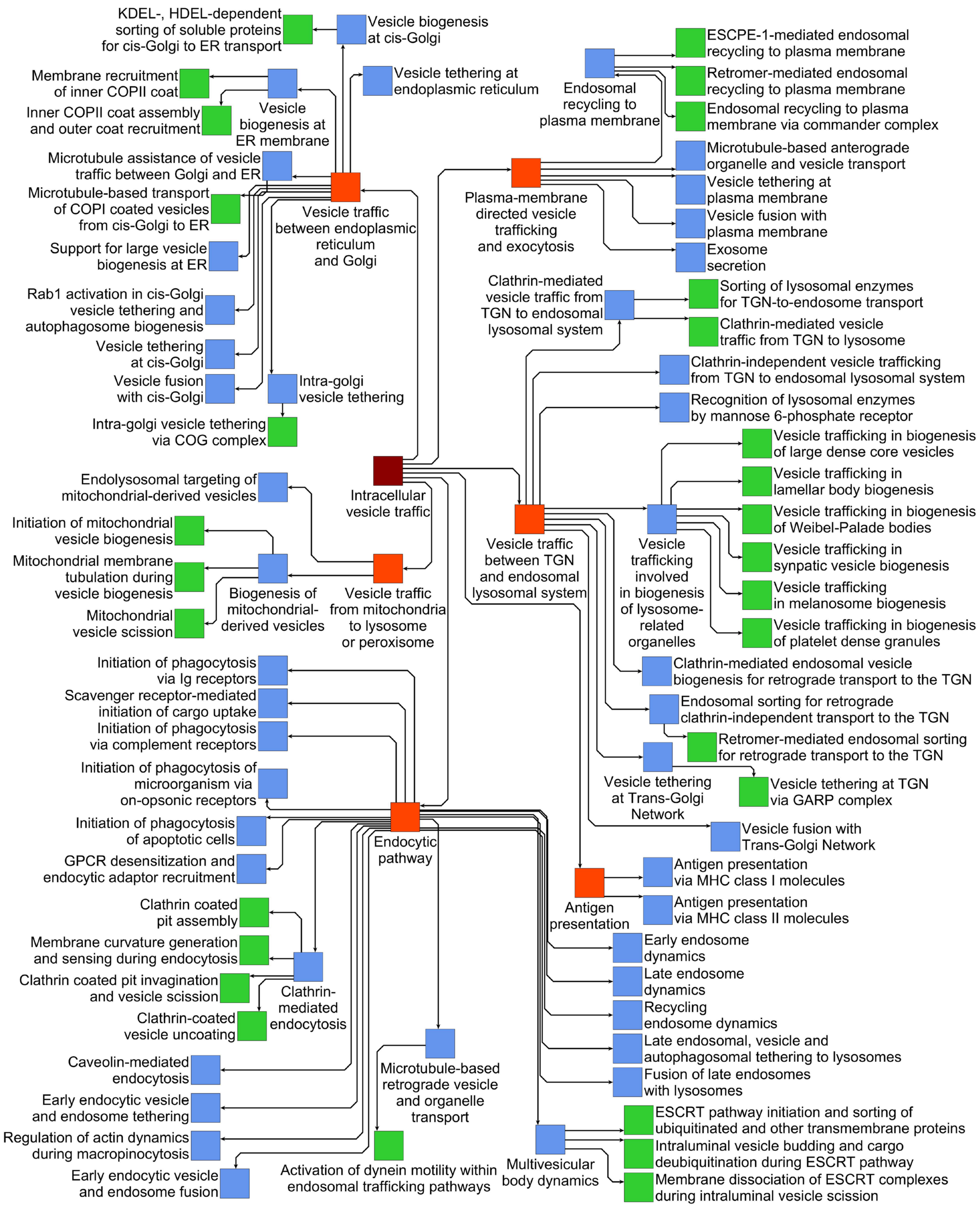


Supplementary figure 1

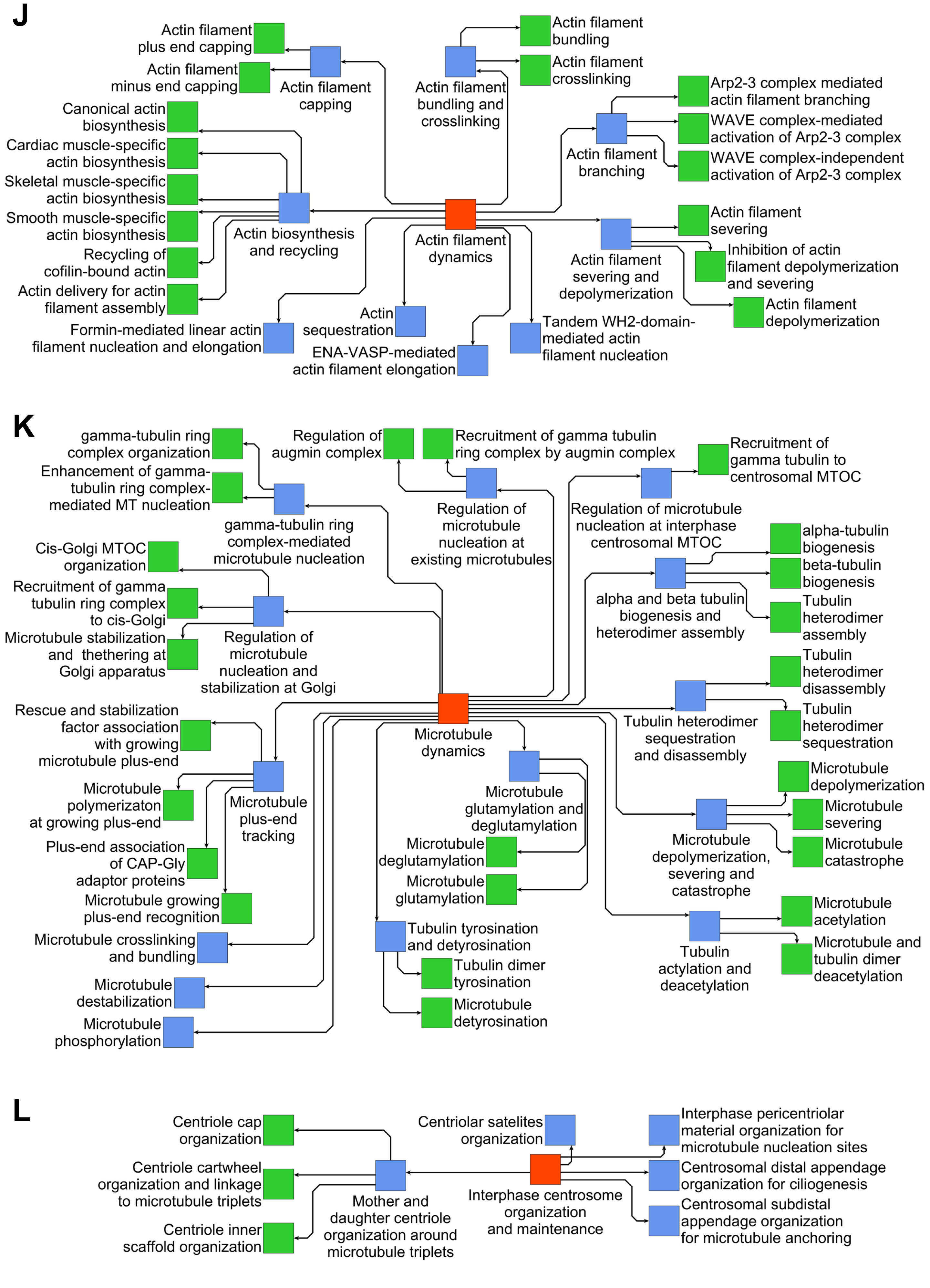


**Supplementary figure 1**

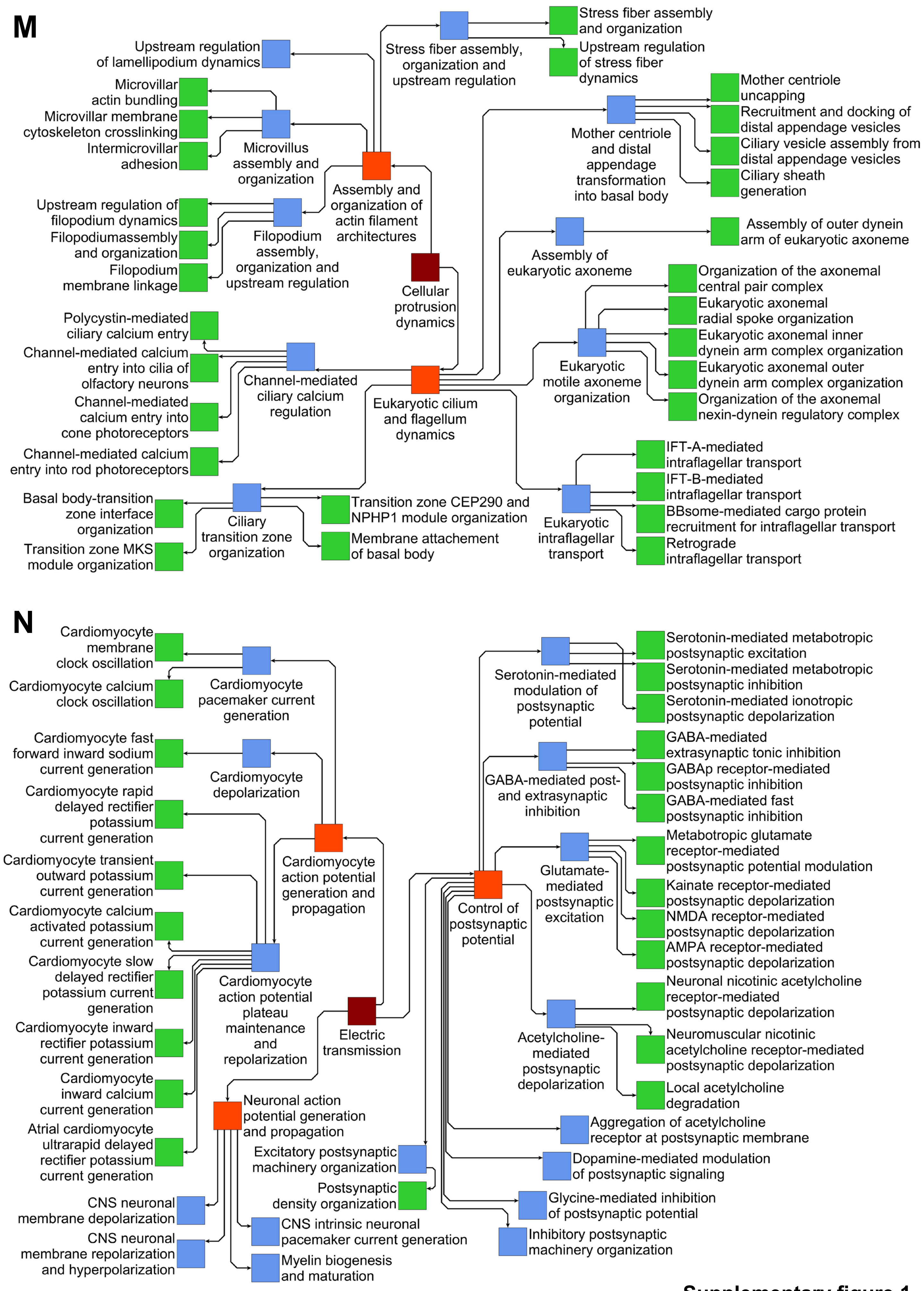


**Supplementary figure 1**

O

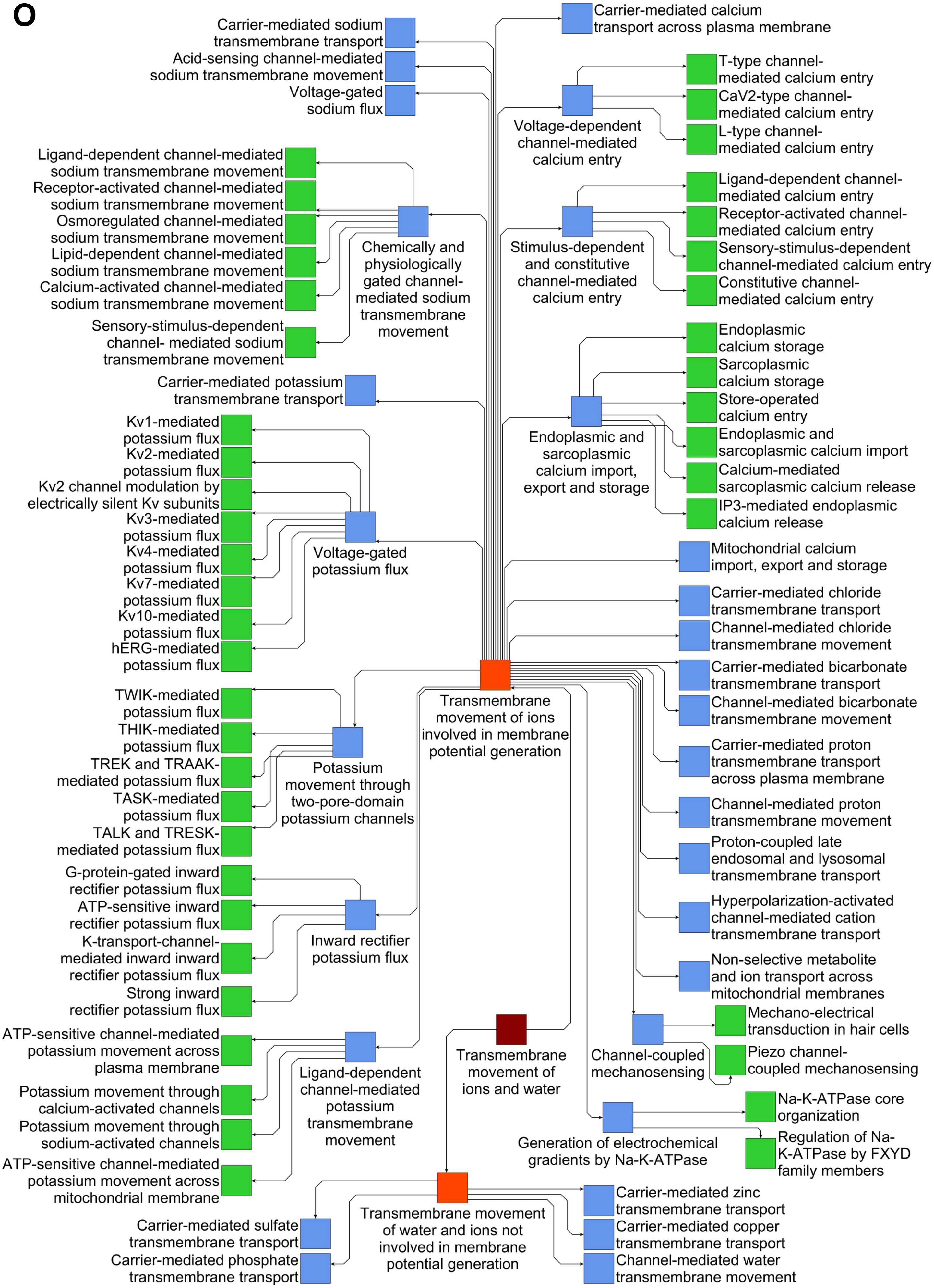


**Supplementary figure 1**

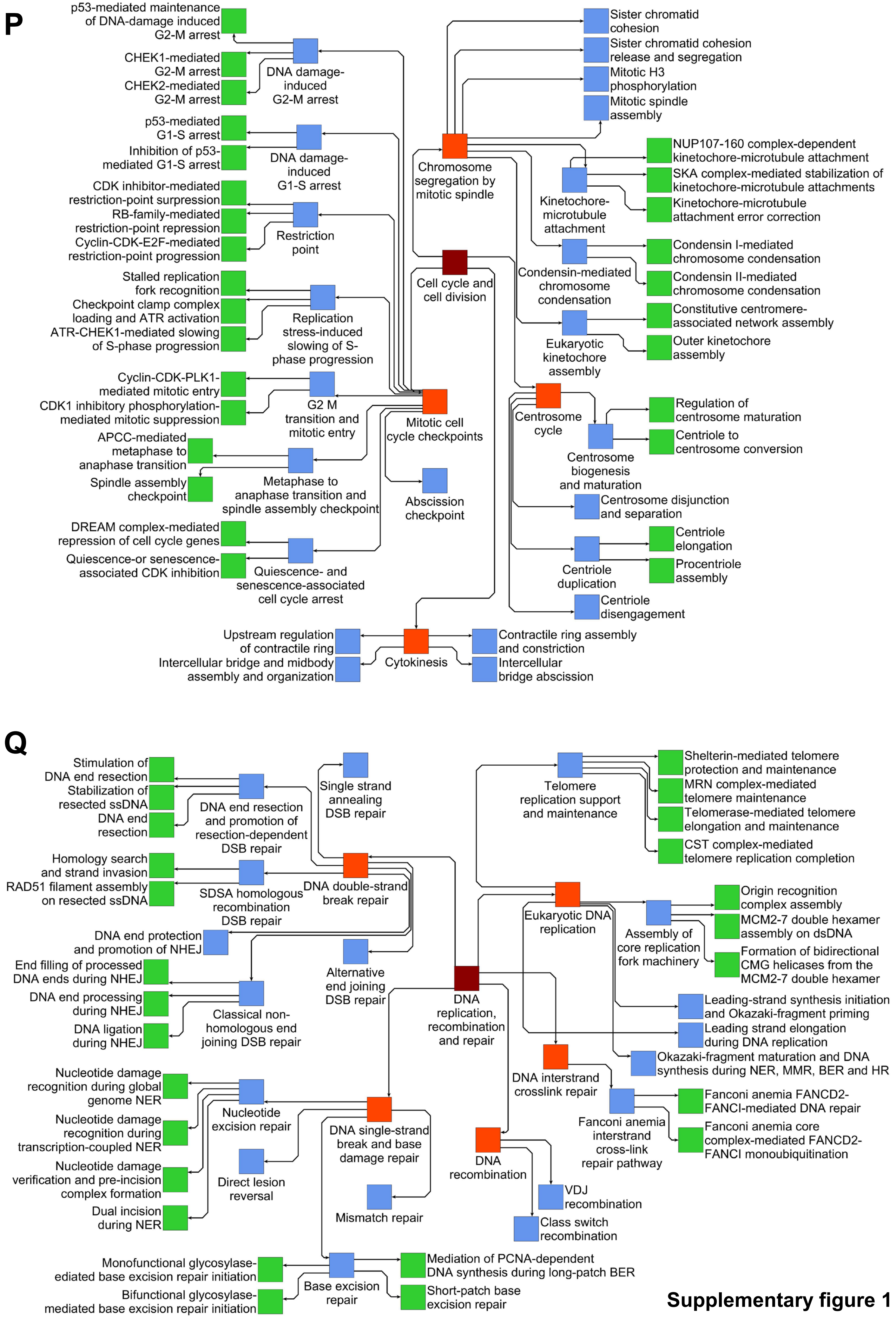


**Supplementary figure 1**

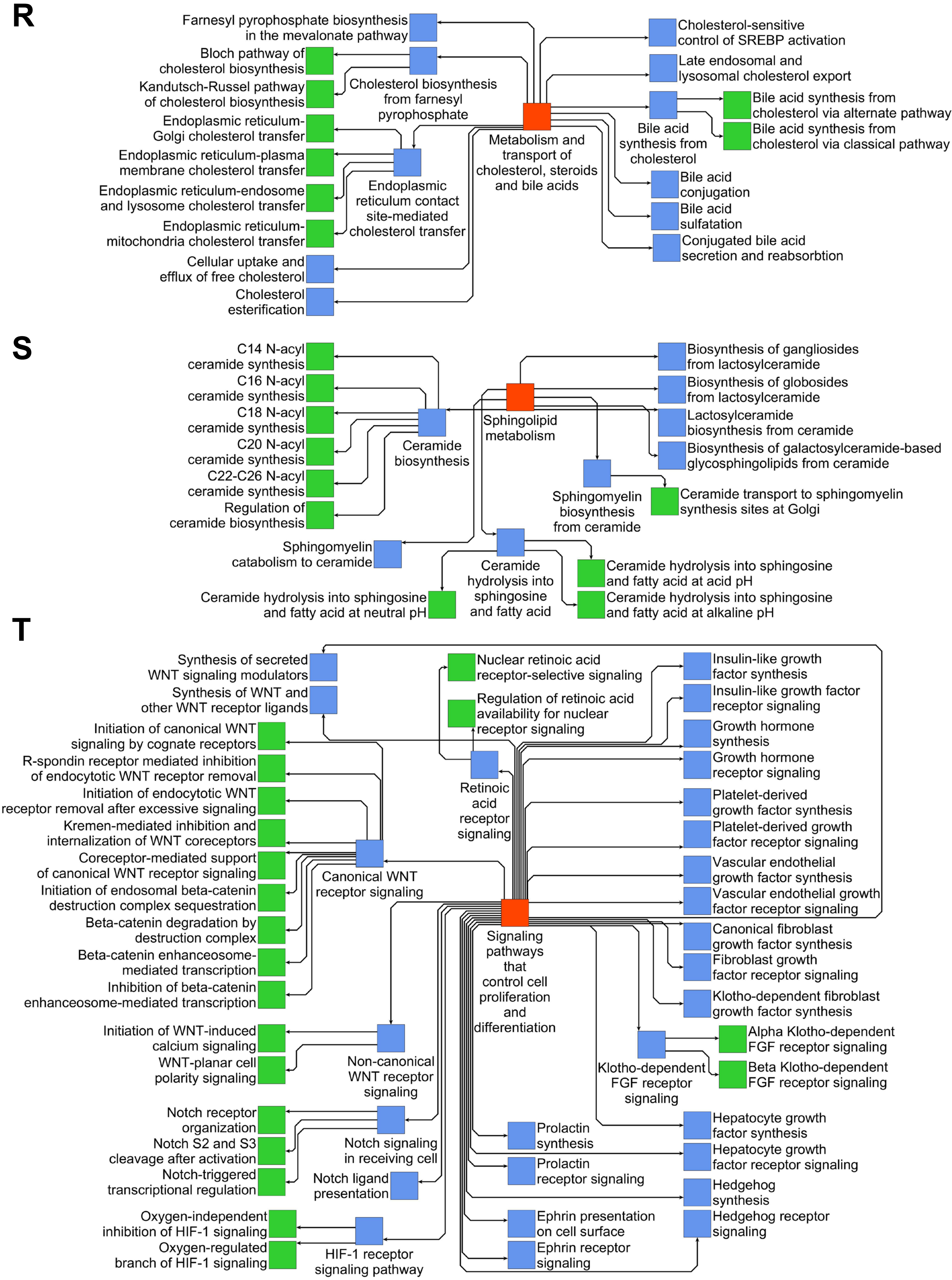


**Supplementary figure 1**

**Supplementary Figure 1. Selected hierarchical relationships.** Selected children of the level 1 or level 2 SCPs **(A)** ‘Extracellular matrix homeostasis’, **(B)** ‘Cellular redox homeostasis’, **(C)** ‘Organelle organization’, **(D)** ‘Intracellular degradation pathways’ and **(E)** ‘Amino acid metabolism’, **(F)** ‘Lipoprotein metabolism’, **(G)** ‘Protein folding and post-translational modification’, **(H)** ‘Intracellular common signaling cascades of multiple pathways’, **(I)** ‘Intracellular vesicle traffic’, **(J)** ‘Actin filament dynamics’, **(K)** ‘Microtubule dynamics’, **(L)** ‘Interphase centrosome organization and maintenance’, **(M)** ‘Cellular protrusion dynamics’, **(N)** ‘Electric transmission’, **(O)** ‘Transmembrane movement of ions involved in membrane potential generation’, **(P)** ‘Cell cycle and cell division’, **(Q)** ‘DNA replication, recombination and repair’, **(R)** ‘Metabolism and transport of cholesterol, steroids and bile acids, **(S)** ‘Sphingolipid metabolism’ and **(T)** ‘Signaling pathways that control cell proliferation and differentiation’ are shown. Level 1, 2, 3 and 4 SCPs are colored in dark red, red, blue and green, respectively.

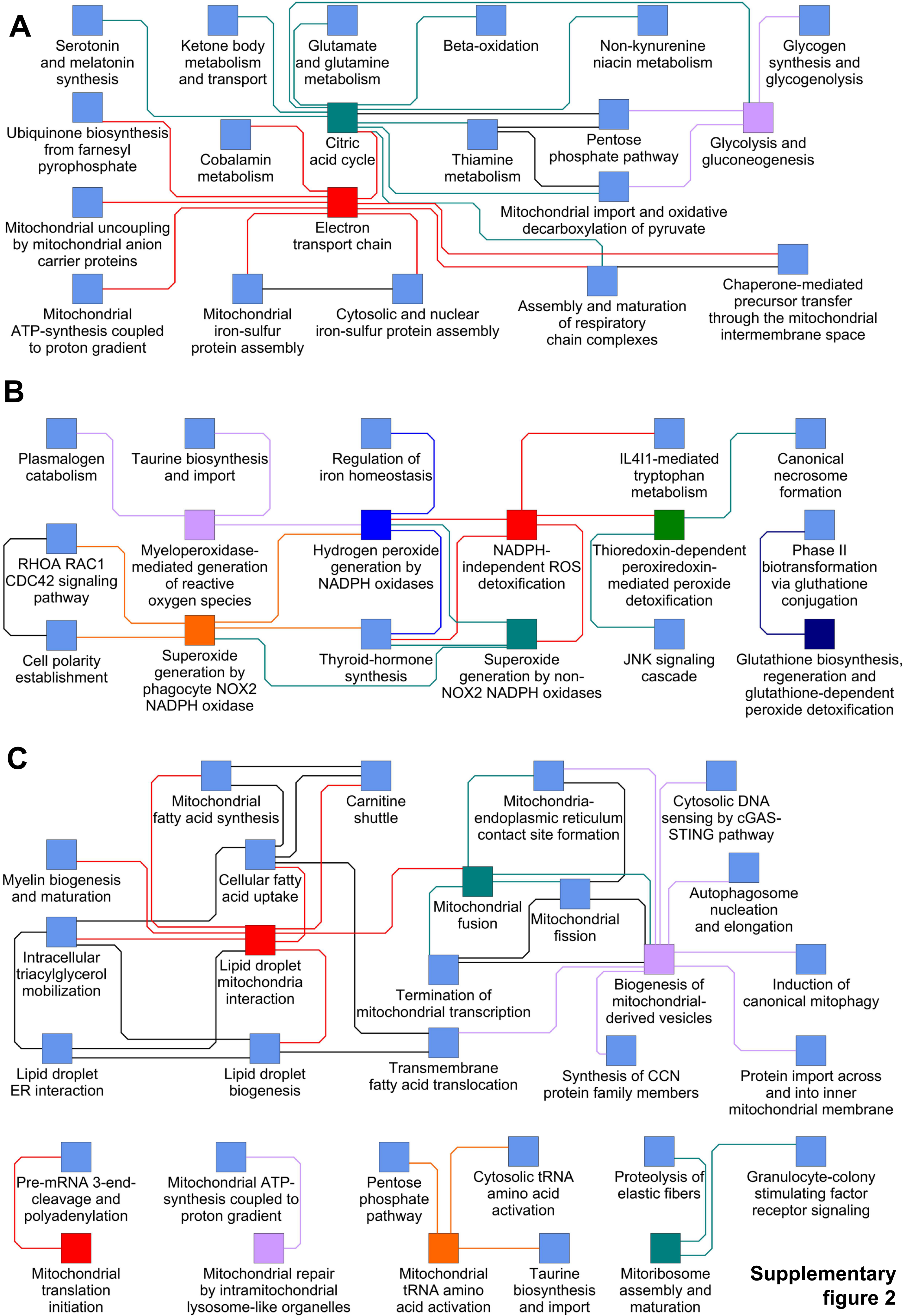


**Supplementary figure 2**

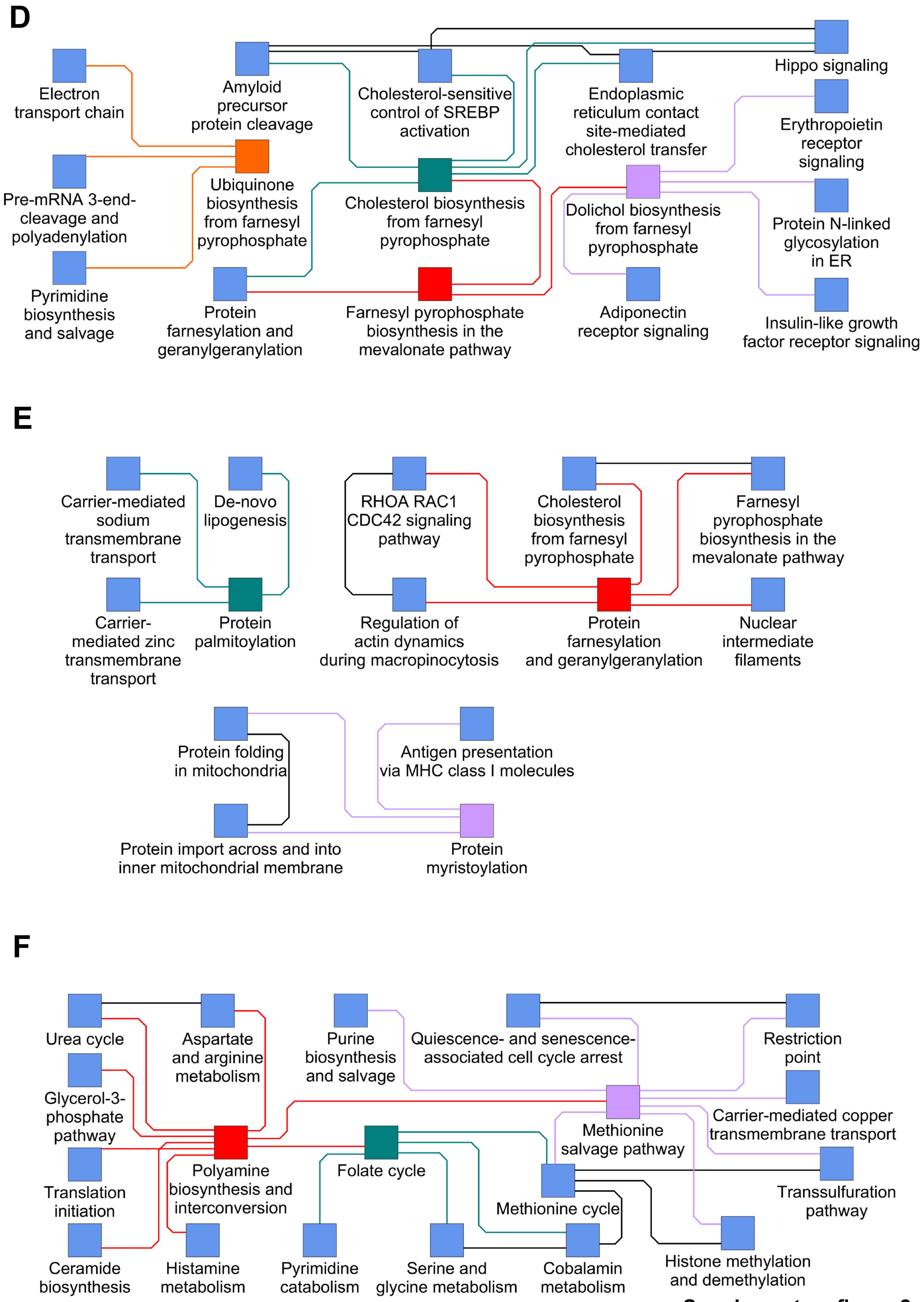


**Supplementary figure 2**

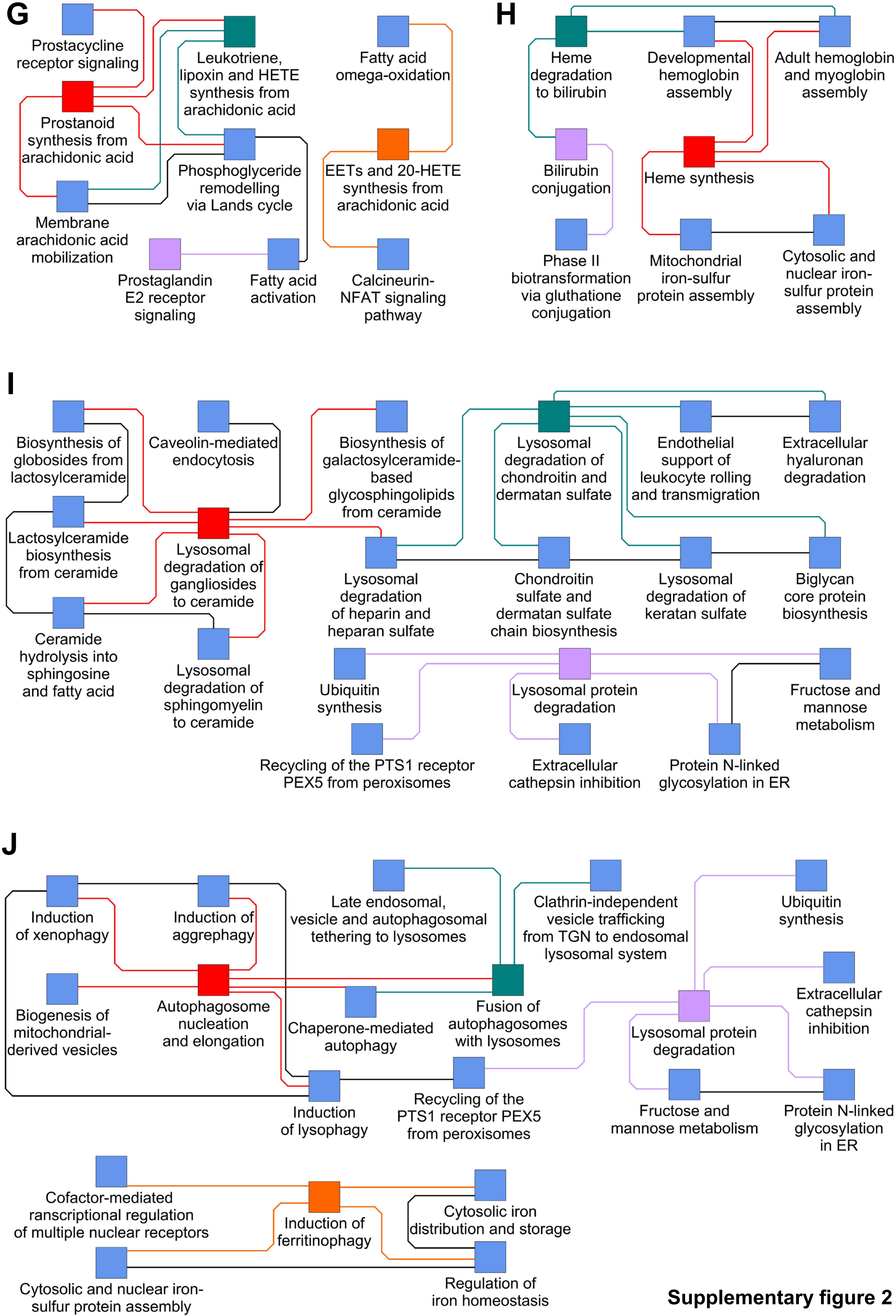


**Supplementary figure 2**

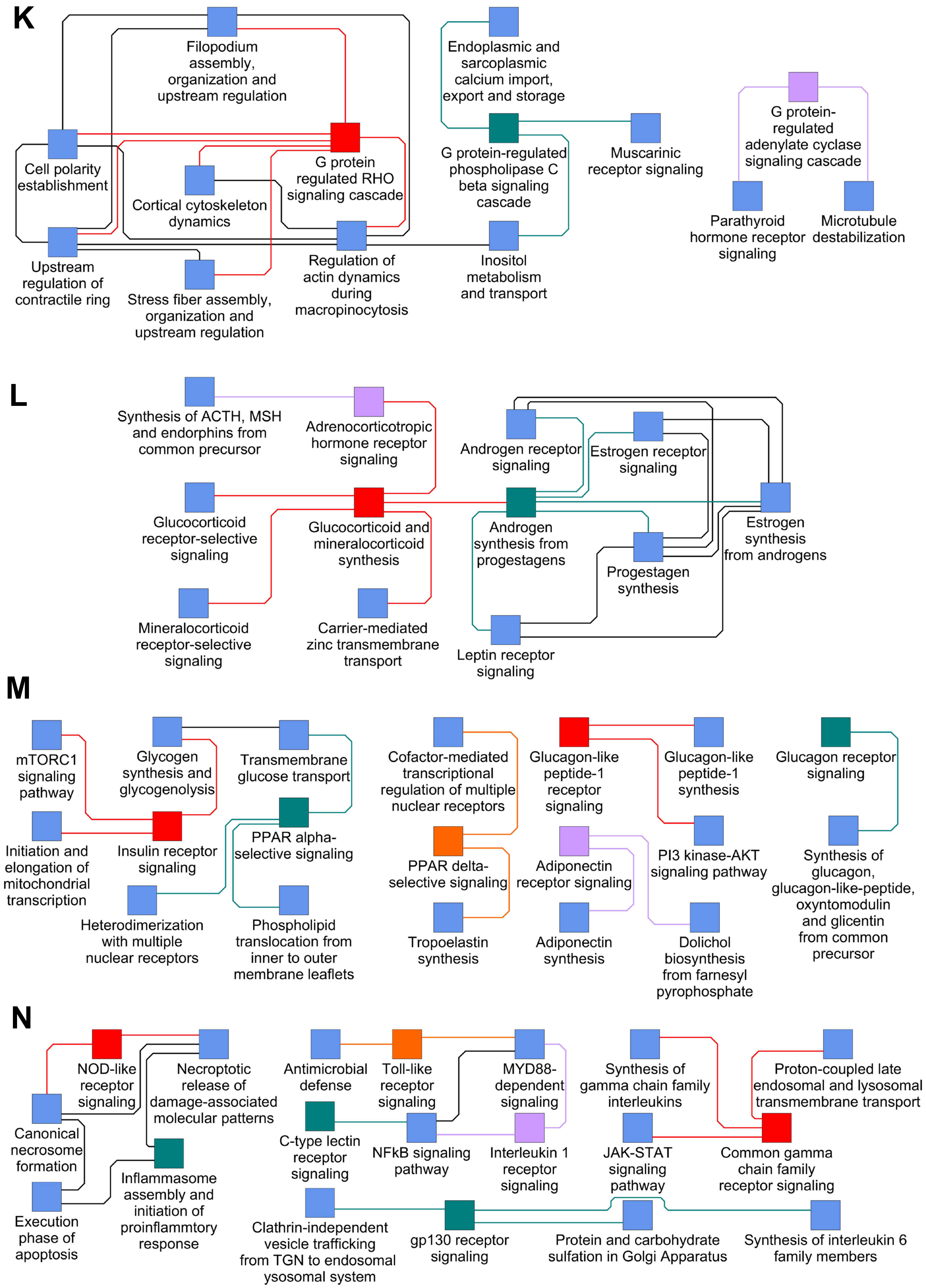


**Supplementary figure 2**

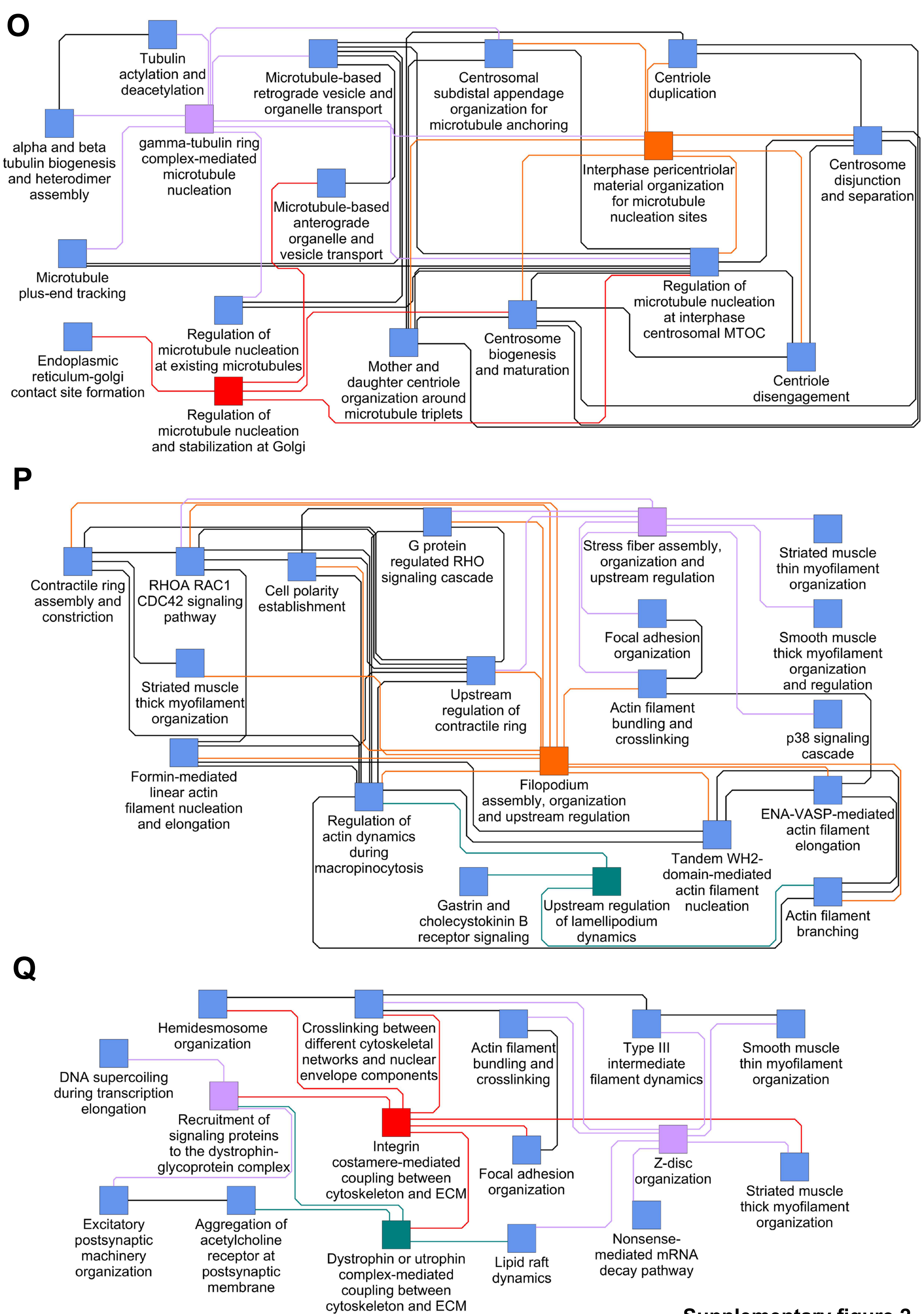


**Supplementary figure 2**

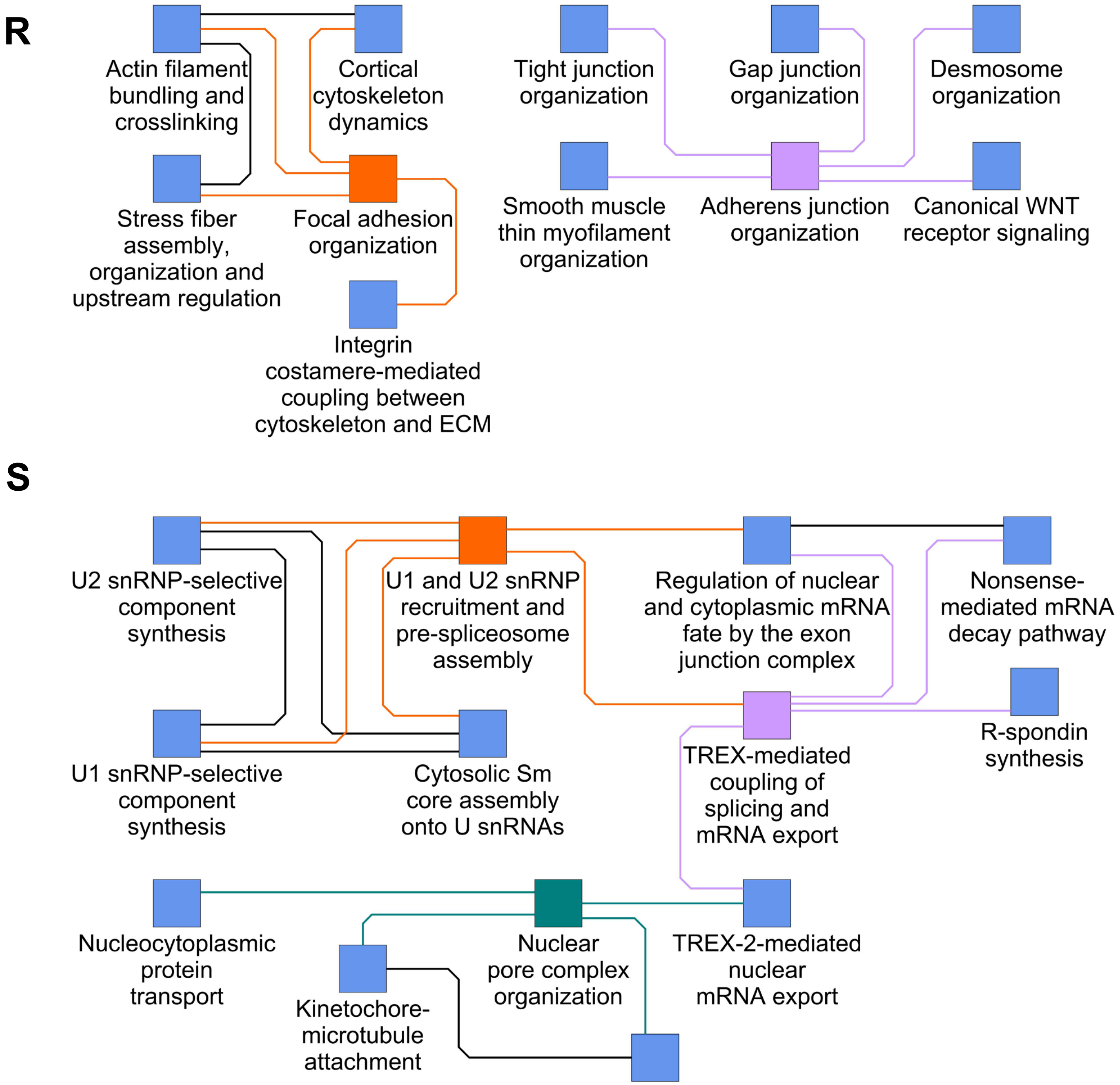


**Supplementary figure 2**

**Supplementary Figure 2. Selected functional relationships between level 3 SCPs.** All direct neighbors (blue) for selected SCPs (dark blue) involved in **(A)** mitochondrial energy generation, **(B)** antioxidative defense, **(C)** mitochondrial dynamics, **(D)** mevalonate and connected synthesis pathways, **(E)** protein lipidation, **(F)** polyamine, methionine and folate metabolism, **(G)** eicosanoid metabolism, **(H)** heme metabolism, **(I)** lysosomal metabolic pathways, **(J)** autophagy , **(K)** G protein-mediated signaling, **(L)** steroid hormone synthesis and signaling **(M)** metabolic signaling pathways, **(N)** inflammatory signaling pathways, **(O)** microtubule dynamics, **(P)** actin based cytoskeletal structures, **(Q)** costamere-mediated adhesion, **(R)** canonical cell adhesion and **(S)** nuclear mRNA dynamics. The top 25% of all functional level 3 SCP interactions were considered. Seed SCPs and edges that connect them to their directed neighbors are colored red, orange, dark cyan, purple or dark blue. Direct neighbor SCPs are colored in light blue, edges connecting direct neighbor SCPs with each other are colored black.

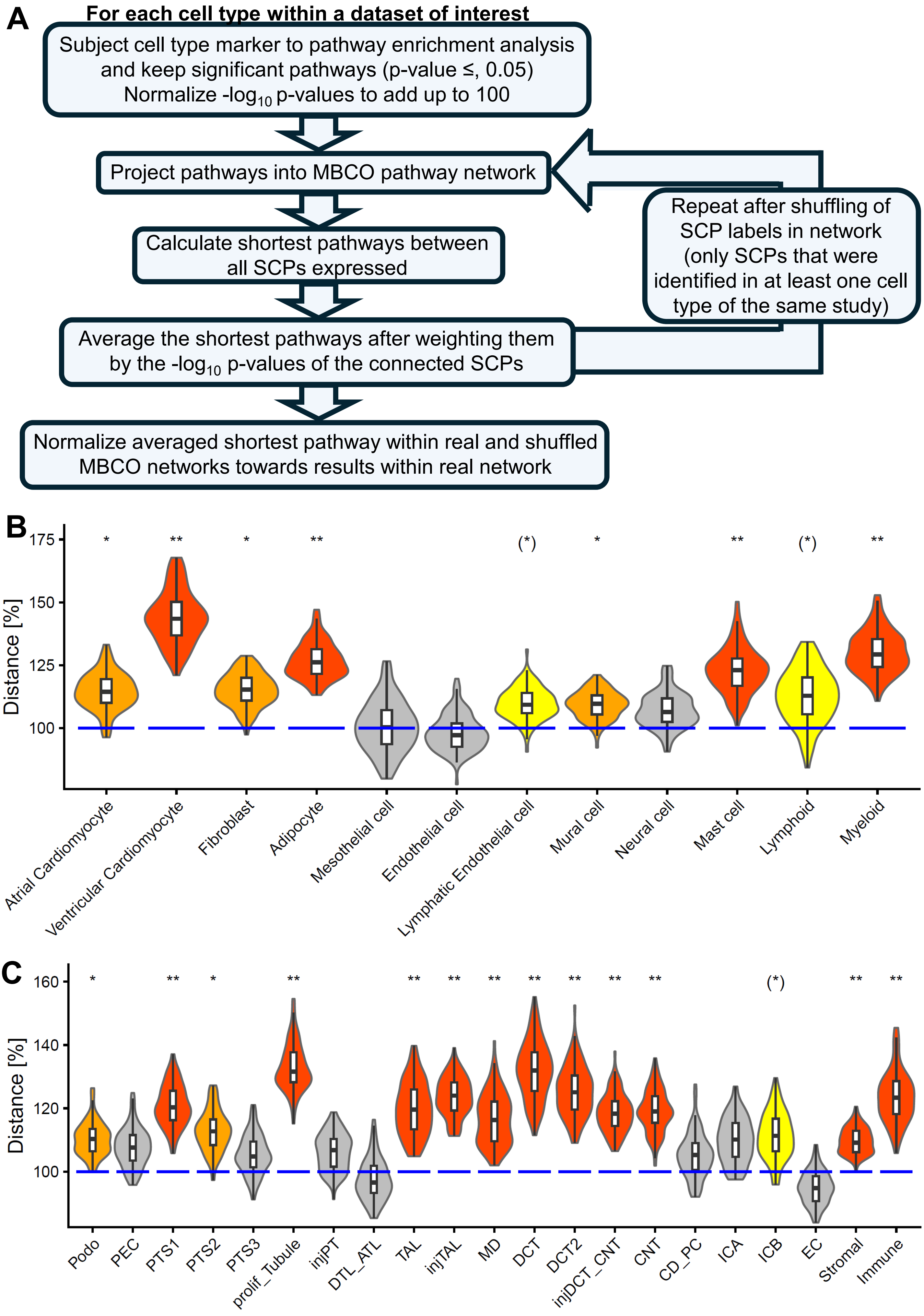


**Supplementary figure 3**

**Supplementary Figure 3. Functional integrity of SCPs expressed in the same cell type. (A)** The flow chart summarizes the main steps for the calculation of the functional integrity of SCPs expressed in the same cell type. Marker genes of **(B)** cardiac and **(C)** kidney cell types were subjected to pathway enrichment analysis. For each cell type we calculated the pairwise distances between all significant SCPs within the inferred network of functional relationships. All distances were averaged after weighting them by $-\log_{10}$ p-values of the connected SCPs and defined as 100% (blue lines). Null distributions (violins) were generated with the same algorithm after network label shuffling between SCPs predicted in at least one cell type for 100 times. Empirical p-values were calculated. Red, orange, gray: (*), *, **: $0.1 \geq p > 0.05$, $0.05 \geq p > 0.01$, $p \leq 0.01$.

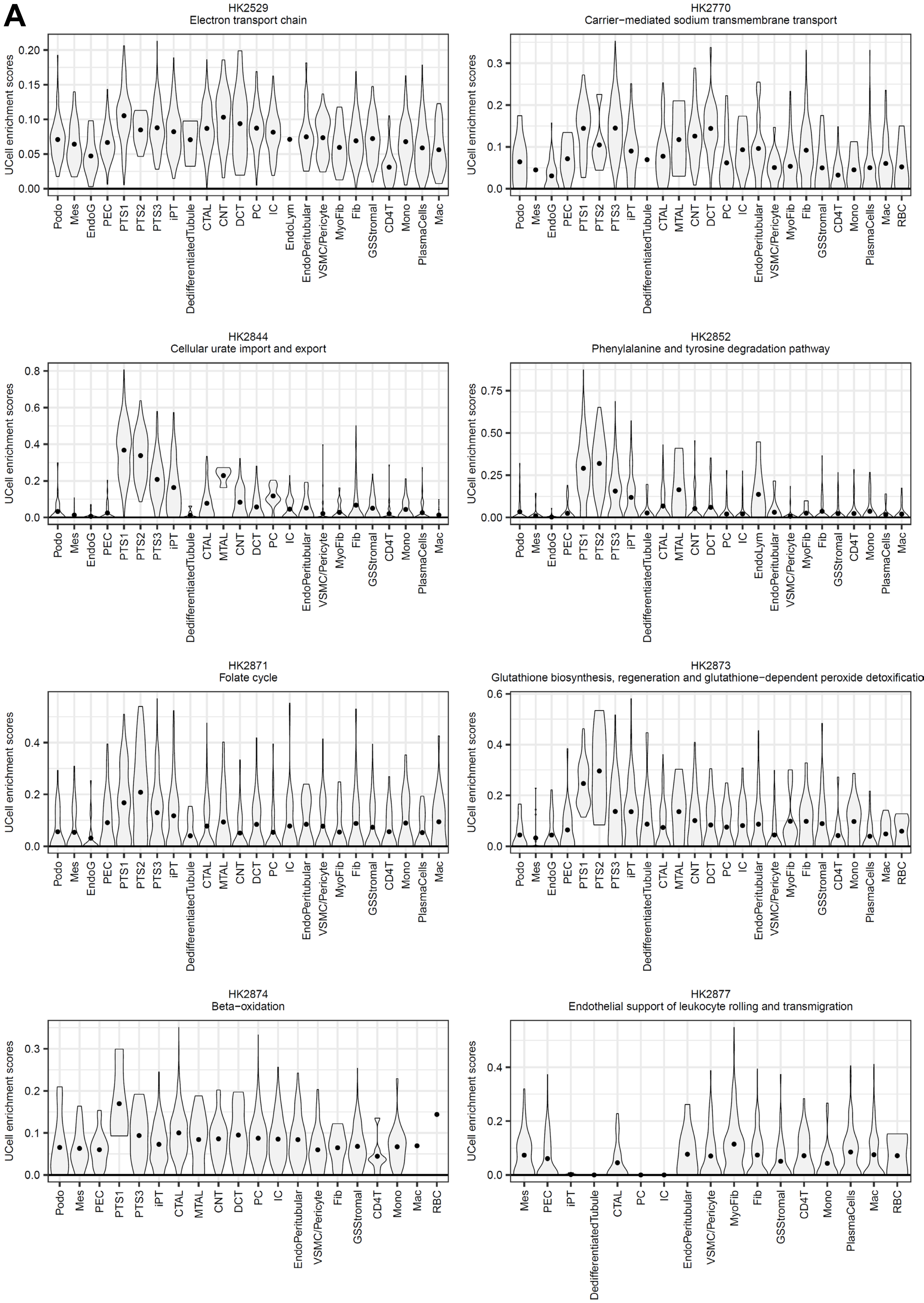


**Supplementary figure 4**

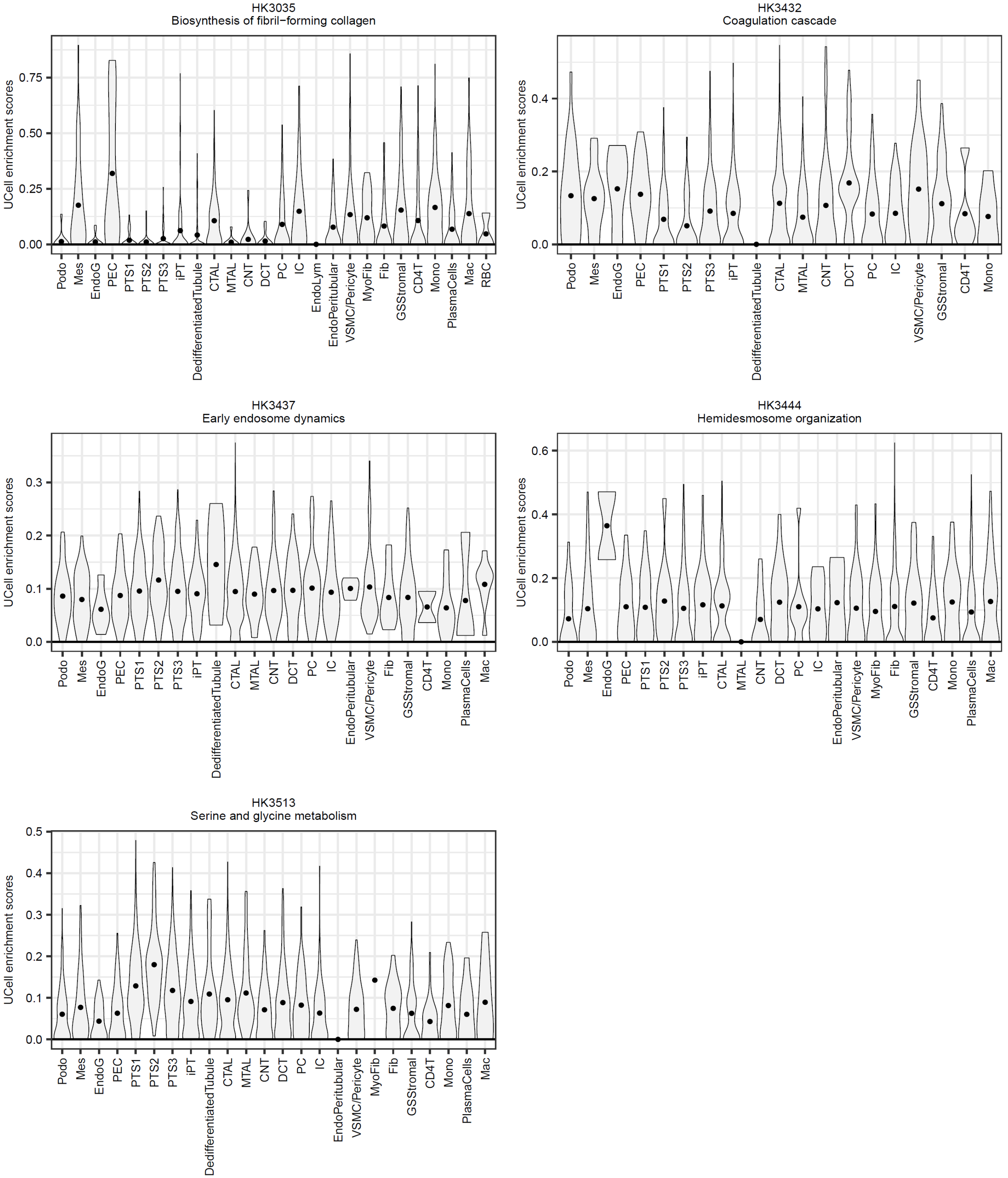


**Supplementary figure 4**

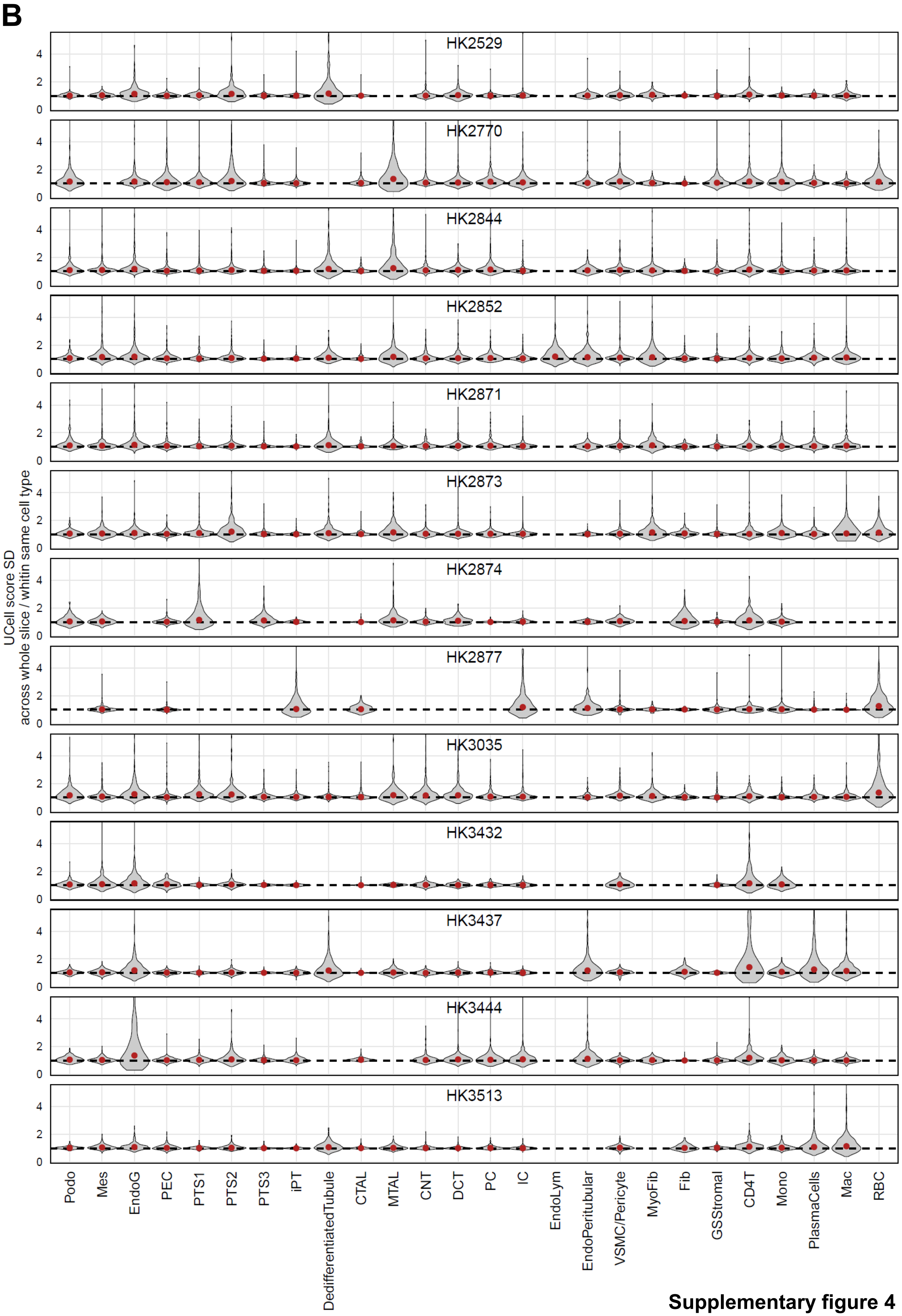


**Supplementary figure 4**

**Supplementary Figure 4. Predicted SCP activities vary across all spots annotated to the same cell type.** For each spot, we identified genes upregulated relative to the mean expression across all other spots and subjected them to enrichment analysis with MBCO level 3 SCPs, using UCell. **(A)** Violin plots show enrichment scores for selected SCPs of all spots annotated to the same cell types on the given slice. **(B)** For each SCP on a given slice, we calculated the ratio of the standard deviations across all spots and across all spots annotated to the same cell type. Red dots indicate medians. For this summary figure, only cell types annotated to at least 100 spots were considered.

**A**

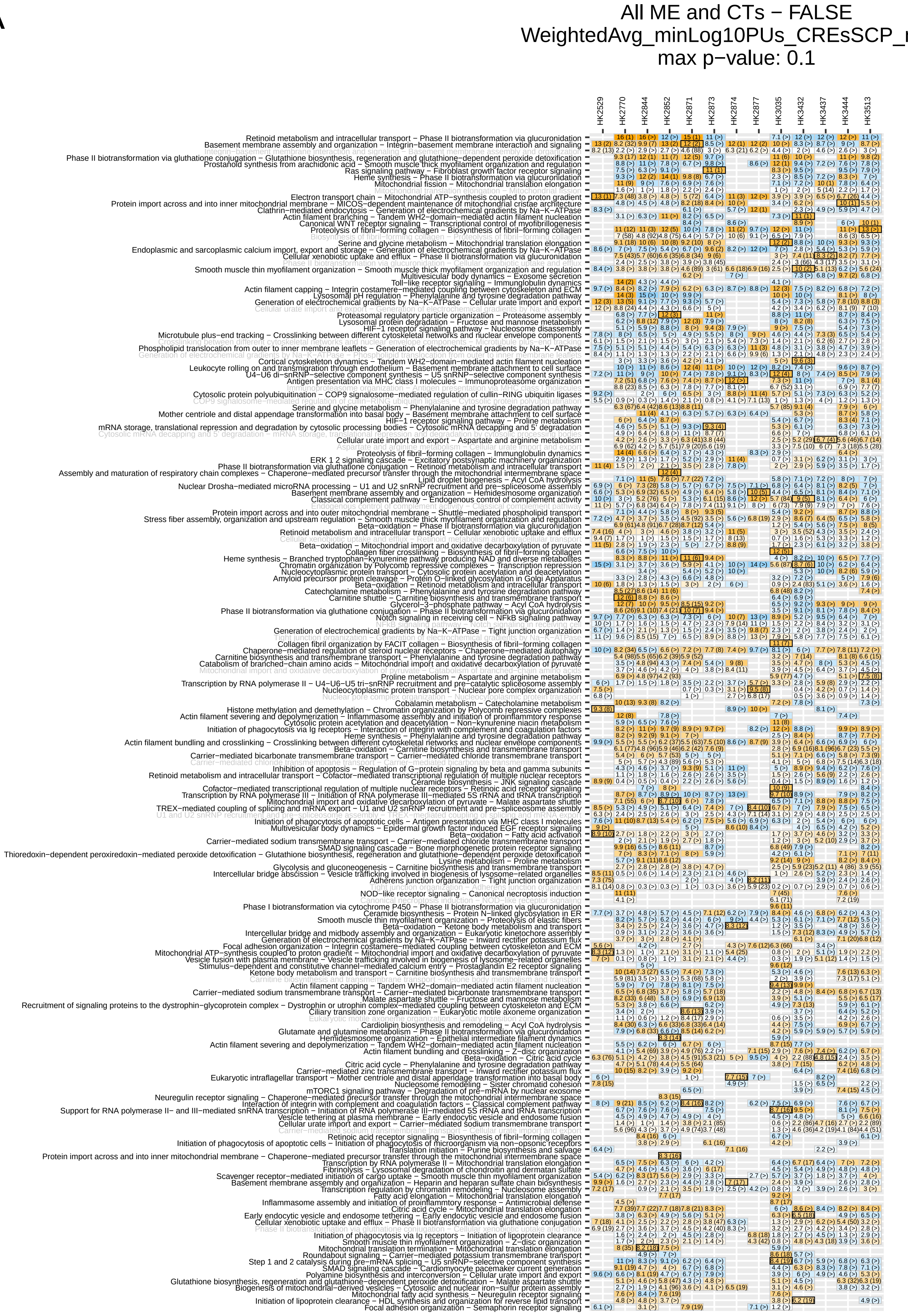
All ME and CTs − FALSE
WeightedAvg_minLog10PUs_CREsSCP_r
max p−value: 0.1
HK2529
HK2770
HK2844
HK2852
HK2871
HK2873
HK2874
HK2877
HK3035
HK3432
HK3437
HK3444
HK3513
Retinoid metabolism and intracellular transport − Phase II biotransformation via glucuronidation
Basement membrane assembly and organization − Integrin−basement membrane interaction and signaling
Integrin−basement membrane interaction and signaling − Basement membrane assembly and organization
Phase II biotransformation via glutathione conjugation − Glutathione biosynthesis, regeneration and glutathione−dependent peroxide detoxification
Prostanoid synthesis from arachidonic acid − Smooth muscle thick myofilament organization and regulation
Ras signaling pathway − Fibroblast growth factor receptor signaling
Heme synthesis − Phase II biotransformation via glucuronidation
Mitochondrial fission − Mitochondrial translation elongation
Mitochondrial translation elongation − Mitochondrial fission
Electron transport chain − Mitochondrial ATP−synthesis coupled to proton gradient
Protein import across and into inner mitochondrial membrane − MICOS−dependent maintenance of mitochondrial cristae architecture
Clathrin−mediated endocytosis − Generation of electrochemical gradients by Na−K−ATPase
Actin filament branching − Tandem WH2−domain−mediated actin filament nucleation
Canonical WNT receptor signaling − Transcriptional control of myofibrillogenesis
Proteolysis of fibril−forming collagen − Biosynthesis of fibril−forming collagen
Biosynthesis of fibril−forming collagen − Proteolysis of fibril−forming collagen
Serine and glycine metabolism − Mitochondrial translation elongation
Endoplasmic and sarcoplasmic calcium import, export and storage − Generation of electrochemical gradients by Na−K−ATPase
Cellular xenobiotic uptake and efflux − Phase II biotransformation via glucuronidation
Phase II biotransformation via glucuronidation − Cellular xenobiotic uptake and efflux
Smooth muscle thin myofilament organization − Smooth muscle thick myofilament organization and regulation
Multivesicular body dynamics − Exosome secretion
Toll−like receptor signaling − Immunglobulin dynamics
Actin filament capping − Integrin costamere−mediated coupling between cytoskeleton and ECM
Lysosomal pH regulation − Phenylalanine and tyrosine degradation pathway
Generation of electrochemical gradients by Na−K−ATPase − Cellular urate import and export
Cellular urate import and export − Generation of electrochemical gradients by Na−K−ATPase
Proteasomal regulatory particle organization − Proteasome assembly
Lysosomal protein degradation − Fructose and mannose metabolism
HIF−1 receptor signaling pathway − Nucleosome disassembly
Microtubule plus−end tracking − Crosslinking between different cytoskeletal networks and nuclear envelope components
Crosslinking between different cytoskeletal networks and nuclear envelope components − Microtubule plus−end tracking
Phospholipid translocation from outer to inner membrane leaflets − Generation of electrochemical gradients by Na−K−ATPase
Generation of electrochemical gradients by Na−K−ATPase − Phospholipid translocation from outer to inner membrane leaflets
Cortical cytoskeleton dynamics − Tandem WH2−domain−mediated actin filament nucleation
Leukocyte rolling on and transmigration through endothelium − Basement membrane attachment to cell surface
U4−U6 di−snRNP−selective component synthesis − U5 snRNP−selective component synthesis
Antigen presentation via MHC class I molecules − Immunoproteasome organization
Immunoproteasome organization − Antigen presentation via MHC class I molecules
Cytosolic protein polyubiquitination − COP9 signalosome−mediated regulation of cullin−RING ubiquitin ligases
COP9 signalosome−mediated regulation of cullin−RING ubiquitin ligases − Cytosolic protein polyubiquitination
Serine and glycine metabolism − Phenylalanine and tyrosine degradation pathway
Mother centriole and distal appendage transformation into basal body − Basement membrane attachment to cell surface
HIF−1 receptor signaling pathway − Proline metabolism
mRNA storage, translational repression and degradation by cytosolic processing bodies − Cytosolic mRNA decapping and 5' degradation
Cytosolic mRNA decapping and 5' degradation − mRNA storage, translational repression and degradation by cytosolic processing bodies
Cellular urate import and export − Aspartate and arginine metabolism
Aspartate and arginine metabolism − Cellular urate import and export
Proteolysis of fibril−forming collagen − Immunglobulin dynamics
ERK 1 2 signaling cascade − Excitatory postsynaptic machinery organization
Phase II biotransformation via glutathione conjugation − Retinoid metabolism and intracellular transport
Assembly and maturation of respiratory chain complexes − Chaperone−mediated precursor transfer through the mitochondrial intermembrane space
Lipid droplet biogenesis − Acyl CoA hydrolysis
Nuclear Drosha−mediated microRNA processing − U1 and U2 snRNP recruitment and pre−spliceosome assembly
Basement membrane assembly and organization − Hemidesmosome organization
Classical complement pathway − Endogenous control of complement activity
Endogenous control of complement activity − Classical complement pathway
Protein import across and into outer mitochondrial membrane − Shuttle−mediated phospholipid transport
Stress fiber assembly, organization and upstream regulation − Smooth muscle thick myofilament organization and regulation
Beta−oxidation − Phase II biotransformation via glucuronidation
Retinoid metabolism and intracellular transport − Cellular xenobiotic uptake and efflux
Cellular xenobiotic uptake and efflux − Retinoid metabolism and intracellular transport
Beta−oxidation − Mitochondrial import and oxidative decarboxylation of pyruvate
Collagen fiber crosslinking − Biosynthesis of fibril−forming collagen
Heme synthesis − Branched tryptophan−kynurenine pathway producing NAD and diverse metabolites
Chromatin organization by Polycomb repressive complexes − Transcription repression
Nucleocytoplasmic protein transport − Cytosolic protein acetylation and deacetylation
Amyloid precursor protein cleavage − Protein O−linked glycosylation in Golgi Apparatus
Beta−oxidation − Retinoid metabolism and intracellular transport
Catecholamine metabolism − Phenylalanine and tyrosine degradation pathway
Carnitine shuttle − Carnitine biosynthesis and transmembrane transport
Glycerol−3−phosphate pathway − Acyl CoA hydrolysis
Phase II biotransformation via glutathione conjugation − Phase II biotransformation via glucuronidation
Notch signaling in receiving cell − NFkB signaling pathway
NFkB signaling pathway − Notch signaling in receiving cell
Generation of electrochemical gradients by Na−K−ATPase − Tight junction organization
Tight junction organization − Generation of electrochemical gradients by Na−K−ATPase
Collagen fibril organization by FACIT collagen − Biosynthesis of fibril−forming collagen
Chaperone−mediated regulation of steroid nuclear receptors − Chaperone−mediated autophagy
Carnitine biosynthesis and transmembrane transport − Phenylalanine and tyrosine degradation pathway
Catabolism of branched−chain amino acids − Mitochondrial import and oxidative decarboxylation of pyruvate
Mitochondrial import and oxidative decarboxylation of pyruvate − Catabolism of branched−chain amino acids
Proline metabolism − Aspartate and arginine metabolism
Transcription by RNA polymerase II − U4−U6−U5 tri−snRNP recruitment and pre−catalytic spliceosome assembly
Nucleocytoplasmic protein transport − Nuclear pore complex organization
Nuclear pore complex organization − Nucleocytoplasmic protein transport
Cobalamin metabolism − Catecholamine metabolism
Histone methylation and demethylation − Chromatin organization by Polycomb repressive complexes
Actin filament severing and depolymerization − Inflammasome assembly and initiation of proinflammtory response
Cytosolic protein acetylation and deacetylation − Non−kynurenine niacin metabolism
Initiation of phagocytosis via Ig receptors − Interaction of integrin with complement and coagulation factors
Heme synthesis − Phenylalanine and tyrosine degradation pathway
Actin filament bundling and crosslinking − Crosslinking between different cytoskeletal networks and nuclear envelope components
Beta−oxidation − Carnitine biosynthesis and transmembrane transport
Carrier−mediated bicarbonate transmembrane transport − Carrier−mediated chloride transmembrane transport
Carrier−mediated chloride transmembrane transport − Carrier−mediated bicarbonate transmembrane transport
Inhibition of apoptosis − Regulation of G−protein signaling by beta and gamma subunits
Retinoid metabolism and intracellular transport − Cofactor−mediated transcriptional regulation of multiple nuclear receptors
Ceramide biosynthesis − JNK signaling cascade
Cofactor−mediated transcriptional regulation of multiple nuclear receptors − Retinoic acid receptor signaling
Transcription by RNA polymerase III − Initiation of RNA polymerase III−mediated 5S rRNA and tRNA transcription
Mitochondrial import and oxidative decarboxylation of pyruvate − Malate aspartate shuttle
TREX−mediated coupling of splicing and mRNA export − U1 and U2 snRNP recruitment and pre−spliceosome assembly
U1 and U2 snRNP recruitment and pre−spliceosome assembly − TREX−mediated coupling of splicing and mRNA export
Initiation of phagocytosis of apoptotic cells − Antigen presentation via MHC class I molecules
Multivesicular body dynamics − Epidermal growth factor induced EGF receptor signaling
Beta−oxidation − Fatty acid activation
Carrier−mediated sodium transmembrane transport − Carrier−mediated chloride transmembrane transport
SMAD signaling cascade − Bone morphogenetic protein receptor signaling
Thioredoxin−dependent peroxiredoxin−mediated peroxide detoxification − Glutathione biosynthesis, regeneration and glutathione−dependent peroxide detoxification
Lysine metabolism − Proline metabolism
Glycolysis and gluconeogenesis − Carnitine biosynthesis and transmembrane transport
Intercellular bridge abscission − Vesicle trafficking involved in biogenesis of lysosome−related organelles
Adherens junction organization − Tight junction organization
Tight junction organization − Adherens junction organization
NOD−like receptor signaling − Canonical necroptosis induction
Canonical necroptosis induction − NOD−like receptor signaling
Phase I biotransformation via cytochrome P450 − Phase II biotransformation via glucuronidation
Ceramide biosynthesis − Protein N−linked glycosylation in ER
Smooth muscle thin myofilament organization − Proteolysis of elastic fibers
Beta−oxidation − Ketone body metabolism and transport
Intercellular bridge and midbody assembly and organization − Eukaryotic kinetochore assembly
Generation of electrochemical gradients by Na−K−ATPase − Inward rectifier potassium flux
Focal adhesion organization − Integrin costamere−mediated coupling between cytoskeleton and ECM
Mitochondrial ATP−synthesis coupled to proton gradient − Mitochondrial import and oxidative decarboxylation of pyruvate
Vesicle fusion with plasma membrane − Vesicle trafficking involved in biogenesis of lysosome−related organelles
Stimulus−dependent and constitutive channel−mediated calcium entry − Prostaglandin E2 receptor signaling
Ketone body metabolism and transport − Carnitine biosynthesis and transmembrane transport
Carnitine biosynthesis and transmembrane transport − Ketone body metabolism and transport
Actin filament capping − Tandem WH2−domain−mediated actin filament nucleation
Carrier−mediated sodium transmembrane transport − Carrier−mediated bicarbonate transmembrane transport
Malate aspartate shuttle − Fructose and mannose metabolism
Recruitment of signaling proteins to the dystrophin−glycoprotein complex − Dystrophin or utrophin complex−mediated coupling between cytoskeleton and ECM
Ciliary transition zone organization − Eukaryotic motile axoneme organization
Eukaryotic motile axoneme organization − Ciliary transition zone organization
Cardiolipin biosynthesis and remodeling − Acyl CoA hydrolysis
Glutamate and glutamine metabolism − Phase II biotransformation via glucuronidation
Hemidesmosome organization − Epithelial intermediate filament dynamics
Actin filament severing and depolymerization − Tandem WH2−domain−mediated actin filament nucleation
Actin filament bundling and crosslinking − Z−disc organization
Beta−oxidation − Citric acid cycle
Citric acid cycle − Phenylalanine and tyrosine degradation pathway
Carrier−mediated zinc transmembrane transport − Inward rectifier potassium flux
Eukaryotic intraflagellar transport − Mother centriole and distal appendage transformation into basal body
Nucleosome remodeling − Sister chromatid cohesion
mTORC1 signaling pathway − Degradation of pre−mRNA by nuclear exosome
Neuregulin receptor signaling − Chaperone−mediated precursor transfer through the mitochondrial intermembrane space
Interaction of integrin with complement and coagulation factors − Classical complement pathway
Support for RNA polymerase II− and III−mediated snRNA transcription − Initiation of RNA polymerase III−mediated 5S rRNA and tRNA transcription
Vesicle tethering at plasma membrane − Early endocytic vesicle and endosome fusion
Cellular urate import and export − Carrier−mediated sodium transmembrane transport
Carrier−mediated sodium transmembrane transport − Cellular urate import and export
Retinoic acid receptor signaling − Biosynthesis of fibril−forming collagen
Initiation of phagocytosis of apoptotic cells − Initiation of phagocytosis of microorganism via non−opsonic receptors
Translation initiation − Purine biosynthesis and salvage
Protein import across and into inner mitochondrial membrane − Chaperone−mediated precursor transfer through the mitochondrial intermembrane space
Transcription by RNA polymerase II − Mitochondrial translation elongation
Fibrinolysis − Lysosomal degradation of chondroitin and dermatan sulfate
Scavenger receptor−mediated initiation of cargo uptake − Smooth muscle thin myofilament organization
Basement membrane assembly and organization − Heparin and heparan sulfate chain biosynthesis
Transcription regulation by chromatin remodeling − Nucleosome assembly
Fatty acid elongation − Mitochondrial translation elongation
Inflammasome assembly and initiation of proinflammtory response − Antimicrobial defense
Citric acid cycle − Mitochondrial translation elongation
Early endocytic vesicle and endosome tethering − Early endocytic vesicle and endosome fusion
Cellular xenobiotic uptake and efflux − Phase II biotransformation via glutathione conjugation
Phase II biotransformation via glutathione conjugation − Cellular xenobiotic uptake and efflux
Initiation of phagocytosis via Ig receptors − Initiation of lipoprotein clearance
Smooth muscle thin myofilament organization − Z−disc organization
Mitochondrial translation termination − Mitochondrial translation elongation
Roundabout signaling − Carrier−mediated potassium transmembrane transport
Step 1 and 2 catalysis during pre−mRNA splicing − U5 snRNP−selective component synthesis
SMAD signaling cascade − Cardiomyocyte pacemaker current generation
Polyamine biosynthesis and interconversion − Cellular urate import and export
Glutathione biosynthesis, regeneration and glutathione−dependent peroxide detoxification − Malate aspartate shuttle
Biogenesis of mitochondrial−derived vesicles − Cytosolic and nuclear iron−sulfur protein assembly
Mitochondrial fatty acid synthesis − Neuregulin receptor signaling
Initiation of lipoprotein clearance − HDL synthesis and organization for reverse lipid transport
Focal adhesion organization − Semaphorin receptor signaling

B

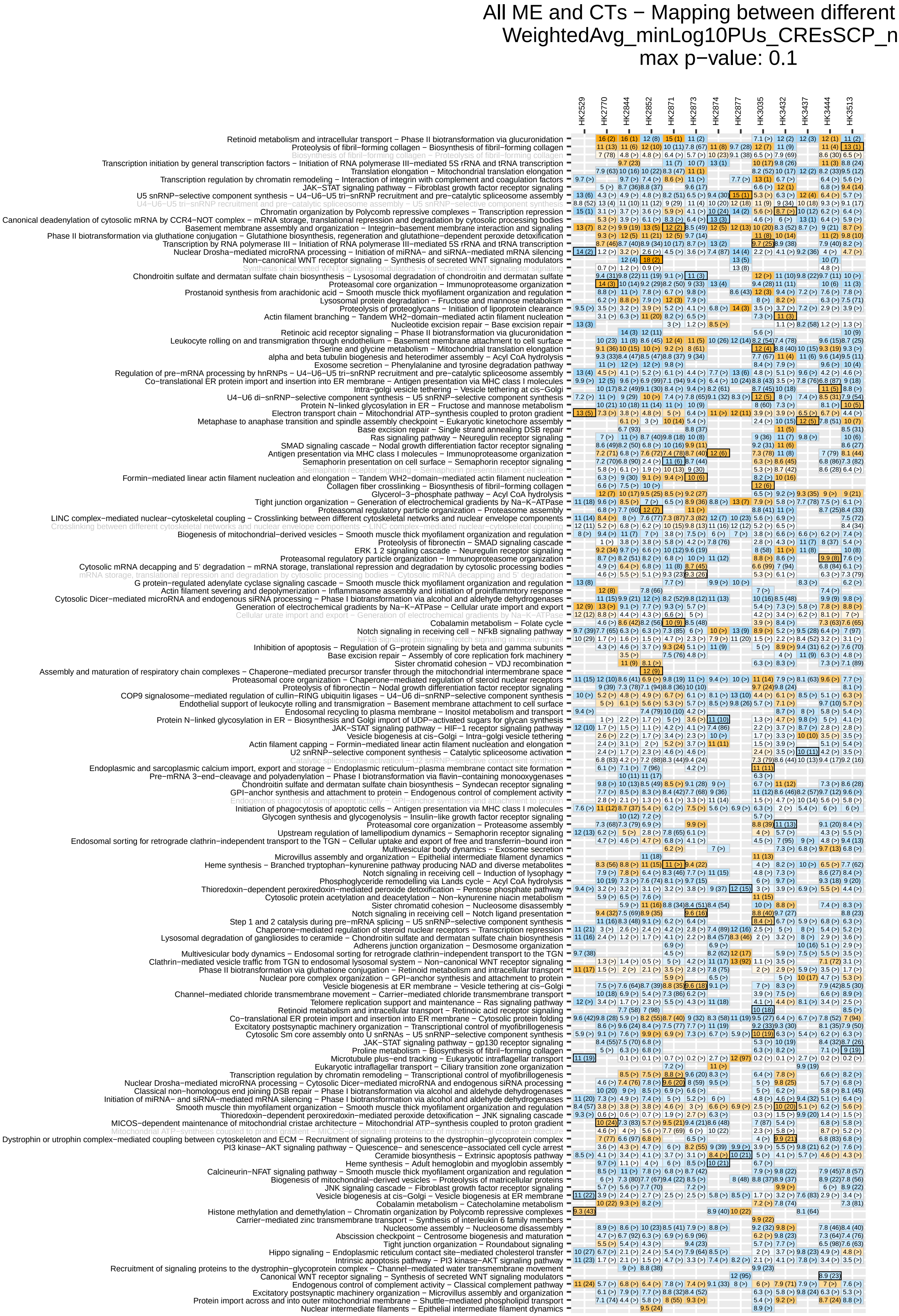
All ME and CTs − Mapping between different
WeightedAvg_minLog10PUs_CREsSCP_n
max p−value: 0.1
HK2529
HK2770
HK2844
HK2852
HK2871
HK2873
HK2874
HK2877
HK3035
HK3432
HK3437
HK3444
HK3513
Retinoid metabolism and intracellular transport − Phase II biotransformation via glucuronidation
Proteolysis of fibril−forming collagen − Biosynthesis of fibril−forming collagen
Biosynthesis of fibril−forming collagen − Proteolysis of fibril−forming collagen
Transcription initiation by general transcription factors − Initiation of RNA polymerase III−mediated 5S rRNA and tRNA transcription
Translation elongation − Mitochondrial translation elongation
Transcription regulation by chromatin remodeling − Interaction of integrin with complement and coagulation factors
JAK−STAT signaling pathway − Fibroblast growth factor receptor signaling
U5 snRNP−selective component synthesis − U4−U6−U5 tri−snRNP recruitment and pre−catalytic spliceosome assembly
U4−U6−U5 tri−snRNP recruitment and pre−catalytic spliceosome assembly − U5 snRNP−selective component synthesis
Chromatin organization by Polycomb repressive complexes − Transcription repression
Canonical deadenylation of cytosolic mRNA by CCR4−NOT complex − mRNA storage, translational repression and degradation by cytosolic processing bodies
Basement membrane assembly and organization − Integrin−basement membrane interaction and signaling
Phase II biotransformation via glutathione conjugation − Glutathione biosynthesis, regeneration and glutathione−dependent peroxide detoxification
Transcription by RNA polymerase III − Initiation of RNA polymerase III−mediated 5S rRNA and tRNA transcription
Nuclear Drosha−mediated microRNA processing − Initiation of miRNA− and siRNA−mediated mRNA silencing
Non−canonical WNT receptor signaling − Synthesis of secreted WNT signaling modulators
Synthesis of secreted WNT signaling modulators − Non−canonical WNT receptor signaling
Chondroitin sulfate and dermatan sulfate chain biosynthesis − Lysosomal degradation of chondroitin and dermatan sulfate
Proteasomal core organization − Immunoproteasome organization
Prostanoid synthesis from arachidonic acid − Smooth muscle thick myofilament organization and regulation
Lysosomal protein degradation − Fructose and mannose metabolism
Proteolysis of proteoglycans − Initiation of lipoprotein clearance
Actin filament branching − Tandem WH2−domain−mediated actin filament nucleation
Nucleotide excision repair − Base excision repair
Retinoic acid receptor signaling − Phase II biotransformation via glucuronidation
Leukocyte rolling on and transmigration through endothelium − Basement membrane attachment to cell surface
Serine and glycine metabolism − Mitochondrial translation elongation
alpha and beta tubulin biogenesis and heterodimer assembly − Acyl CoA hydrolysis
Exosome secretion − Phenylalanine and tyrosine degradation pathway
Regulation of pre−mRNA processing by hnRNPs − U4−U6−U5 tri−snRNP recruitment and pre−catalytic spliceosome assembly
Co−translational ER protein import and insertion into ER membrane − Antigen presentation via MHC class I molecules
Intra−golgi vesicle tethering − Vesicle tethering at cis−Golgi
U4−U6 di−snRNP−selective component synthesis − U5 snRNP−selective component synthesis
Protein N−linked glycosylation in ER − Fructose and mannose metabolism
Electron transport chain − Mitochondrial ATP−synthesis coupled to proton gradient
Metaphase to anaphase transition and spindle assembly checkpoint − Eukaryotic kinetochore assembly
Base excision repair − Single strand annealing DSB repair
Ras signaling pathway − Neuregulin receptor signaling
SMAD signaling cascade − Nodal growth differentiation factor receptor signaling
Antigen presentation via MHC class I molecules − Immunoproteasome organization
Semaphorin presentation on cell surface − Semaphorin receptor signaling
Semaphorin receptor signaling − Semaphorin presentation on cell surface
Formin−mediated linear actin filament nucleation and elongation − Tandem WH2−domain−mediated actin filament nucleation
Collagen fiber crosslinking − Biosynthesis of fibril−forming collagen
Glycerol−3−phosphate pathway − Acyl CoA hydrolysis
Tight junction organization − Generation of electrochemical gradients by Na−K−ATPase
Proteasomal regulatory particle organization − Proteasome assembly
LINC complex−mediated nuclear−cytoskeletal coupling − Crosslinking between different cytoskeletal networks and nuclear envelope components
Crosslinking between different cytoskeletal networks and nuclear envelope components − LINC complex−mediated nuclear−cytoskeletal coupling
Biogenesis of mitochondrial−derived vesicles − Smooth muscle thick myofilament organization and regulation
Proteolysis of fibronectin − SMAD signaling cascade
ERK 1 2 signaling cascade − Neuregulin receptor signaling
Proteasomal regulatory particle organization − Immunoproteasome organization
Cytosolic mRNA decapping and 5' degradation − mRNA storage, translational repression and degradation by cytosolic processing bodies
mRNA storage, translational repression and degradation by cytosolic processing bodies − Cytosolic mRNA decapping and 5' degradation
G protein−regulated adenylate cyclase signaling cascade − Smooth muscle thick myofilament organization and regulation
Actin filament severing and depolymerization − Inflammasome assembly and initiation of proinflammtory response
Cytosolic Dicer−mediated microRNA and endogenous siRNA processing − Phase I biotransformation via alcohol and aldehyde dehydrogenases
Generation of electrochemical gradients by Na−K−ATPase − Cellular urate import and export
Cellular urate import and export − Generation of electrochemical gradients by Na−K−ATPase
Cobalamin metabolism − Folate cycle
Notch signaling in receiving cell − NFkB signaling pathway
NFkB signaling pathway − Notch signaling in receiving cell
Inhibition of apoptosis − Regulation of G−protein signaling by beta and gamma subunits
Base excision repair − Assembly of core replication fork machinery
Sister chromatid cohesion − VDJ recombination
Assembly and maturation of respiratory chain complexes − Chaperone−mediated precursor transfer through the mitochondrial intermembrane space
Proteasomal core organization − Chaperone−mediated regulation of steroid nuclear receptors
Proteolysis of fibronectin − Nodal growth differentiation factor receptor signaling
COP9 signalosome−mediated regulation of cullin−RING ubiquitin ligases − U4−U6 di−snRNP−selective component synthesis
Endothelial support of leukocyte rolling and transmigration − Basement membrane attachment to cell surface
Endosomal recycling to plasma membrane − Inositol metabolism and transport
Protein N−linked glycosylation in ER − Biosynthesis and Golgi import of UDP−activated sugars for glycan synthesis
JAK−STAT signaling pathway − HIF−1 receptor signaling pathway
Vesicle biogenesis at cis−Golgi − Intra−golgi vesicle tethering
Actin filament capping − Formin−mediated linear actin filament nucleation and elongation
U2 snRNP−selective component synthesis − Catalytic spliceosome activation
Catalytic spliceosome activation − U2 snRNP−selective component synthesis
Endoplasmic and sarcoplasmic calcium import, export and storage − Endoplasmic reticulum−plasma membrane contact site formation
Pre−mRNA 3'−end−cleavage and polyadenylation − Phase I biotransformation via flavin−containing monooxygenases
Chondroitin sulfate and dermatan sulfate chain biosynthesis − Syndecan receptor signaling
GPI−anchor synthesis and attachment to protein − Endogenous control of complement activity
Endogenous control of complement activity − GPI−anchor synthesis and attachment to protein
Initiation of phagocytosis of apoptotic cells − Antigen presentation via MHC class I molecules
Glycogen synthesis and glycogenolysis − Insulin−like growth factor receptor signaling
Proteasomal core organization − Proteasome assembly
Upstream regulation of lamellipodium dynamics − Semaphorin receptor signaling
Endosomal sorting for retrograde clathrin−independent transport to the TGN − Cellular uptake and export of free and transferrin−bound iron
Multivesicular body dynamics − Exosome secretion
Microvillus assembly and organization − Epithelial intermediate filament dynamics
Heme synthesis − Branched tryptophan−kynurenine pathway producing NAD and diverse metabolites
Notch signaling in receiving cell − Induction of lysophagy
Phosphoglyceride remodelling via Lands cycle − Acyl CoA hydrolysis
Thioredoxin−dependent peroxiredoxin−mediated peroxide detoxification − Pentose phosphate pathway
Cytosolic protein acetylation and deacetylation − Non−kynurenine niacin metabolism
Sister chromatid cohesion − Nucleosome disassembly
Notch signaling in receiving cell − Notch ligand presentation
Step 1 and 2 catalysis during pre−mRNA splicing − U5 snRNP−selective component synthesis
Chaperone−mediated regulation of steroid nuclear receptors − Transcription repression
Lysosomal degradation of gangliosides to ceramide − Chondroitin sulfate and dermatan sulfate chain biosynthesis
Adherens junction organization − Desmosome organization
Multivesicular body dynamics − Endosomal sorting for retrograde clathrin−independent transport to the TGN
Clathrin−mediated vesicle traffic from TGN to endosomal lysosomal system − Non−canonical WNT receptor signaling
Phase II biotransformation via glutathione conjugation − Retinoid metabolism and intracellular transport
Nuclear pore complex organization − GPI−anchor synthesis and attachment to protein
Vesicle biogenesis at ER membrane − Vesicle tethering at cis−Golgi
Channel−mediated chloride transmembrane movement − Carrier−mediated chloride transmembrane transport
Telomere replication support and maintenance − Ras signaling pathway
Retinoid metabolism and intracellular transport − Retinoic acid receptor signaling
Co−translational ER protein import and insertion into ER membrane − Cytosolic protein folding
Excitatory postsynaptic machinery organization − Transcriptional control of myofibrillogenesis
Cytosolic Sm core assembly onto U snRNAs − U5 snRNP−selective component synthesis
JAK−STAT signaling pathway − gp130 receptor signaling
Proline metabolism − Biosynthesis of fibril−forming collagen
Microtubule plus−end tracking − Eukaryotic intraflagellar transport
Eukaryotic intraflagellar transport − Ciliary transition zone organization
Transcription regulation by chromatin remodeling − Transcriptional control of myofibrillogenesis
Nuclear Drosha−mediated microRNA processing − Cytosolic Dicer−mediated microRNA and endogenous siRNA processing
Classical non−homologous end joining DSB repair − Phase I biotransformation via alcohol and aldehyde dehydrogenases
Initiation of miRNA− and siRNA−mediated mRNA silencing − Phase I biotransformation via alcohol and aldehyde dehydrogenases
Smooth muscle thin myofilament organization − Smooth muscle thick myofilament organization and regulation
Thioredoxin−dependent peroxiredoxin−mediated peroxide detoxification − JNK signaling cascade
MICOS−dependent maintenance of mitochondrial cristae architecture − Mitochondrial ATP−synthesis coupled to proton gradient
Mitochondrial ATP−synthesis coupled to proton gradient − MICOS−dependent maintenance of mitochondrial cristae architecture
Dystrophin or utrophin complex−mediated coupling between cytoskeleton and ECM − Recruitment of signaling proteins to the dystrophin−glycoprotein complex
PI3 kinase−AKT signaling pathway − Quiescence− and senescence−associated cell cycle arrest
Ceramide biosynthesis − Extrinsic apoptosis pathway
Heme synthesis − Adult hemoglobin and myoglobin assembly
Calcineurin−NFAT signaling pathway − Smooth muscle thick myofilament organization and regulation
Biogenesis of mitochondrial−derived vesicles − Proteolysis of matricellular proteins
JNK signaling cascade − Fibroblast growth factor receptor signaling
Vesicle biogenesis at cis−Golgi − Vesicle biogenesis at ER membrane
Cobalamin metabolism − Catecholamine metabolism
Histone methylation and demethylation − Chromatin organization by Polycomb repressive complexes
Carrier−mediated zinc transmembrane transport − Synthesis of interleukin 6 family members
Nucleosome assembly − Nucleosome disassembly
Abscission checkpoint − Centrosome biogenesis and maturation
Tight junction organization − Roundabout signaling
Hippo signaling − Endoplasmic reticulum contact site−mediated cholesterol transfer
Intrinsic apoptosis pathway − PI3 kinase−AKT signaling pathway
Recruitment of signaling proteins to the dystrophin−glycoprotein complex − Channel−mediated water transmembrane movement
Canonical WNT receptor signaling − Synthesis of secreted WNT signaling modulators
Endogenous control of complement activity − Classical complement pathway
Excitatory postsynaptic machinery organization − Microvillus assembly and organization
Protein import across and into outer mitochondrial membrane − Shuttle−mediated phospholipid transport
Nuclear intermediate filaments − Epithelial intermediate filament dynamics

**Supplementary Figure 5. Top Interactions between SCPs with functional and potential spatial significance. (A)** Identified interactions between center and peripheral SCPs with functional and spatial significance were ranked within each dataset by spatial significance. The top 20 interactions of each dataset were curated and results are shown for those across all datasets, if functionally significant. Numbers show ES units, numbers in brackets spatial p-value ranks. Orange/blue fields show interactions between SCPs with functional and spatial or only functional significance, respectively. Framed fields indicate selections for follow-up visualizations, e.g., SCP pair expression on the given slice. Reverse SCP pairs (pairs already shown in the opposite direction) are shown in light gray with faded field colors. **(B)** Identified interactions between center and peripheral SCPs with functional significance (independently of spatial significance) are shown as in A. In contrast to A, numbers in brackets show ES score ranks. Note that some of the SCP pairs overlap with the SCP pairs curated in A.
