## Supplementary Figures 6-8 for "Functional cooperation, adaptation and synchronization of subcellular processes in cellular neighborhoods as inferred from spatial transcriptomics"

**Abstract**

Tissue-level functions emerge from pathway activities distributed across neighboring cells. Yet analyses of spatial transcriptomics mostly focus on cell type composition or signaling through ligand-receptor pairs, leaving spatial relationships between effector processes undescribed. We inferred networks of functionally interacting subcellular processes (SCPs) that are independent of shared gene products and lie beyond sibling relationships defined in parent-child hierarchies. Using those networks for the analysis of published spatial transcriptomics datasets from human kidney biopsies, we identified functionally related SCP activities between a tissue spot and its neighbors. Identified pairs of adjacent SCPs indicate functional cooperation, e.g. basement membrane assembly and cell-matrix adhesion, functional adaption, e.g. complement synthesis and inhibition, and functional synchronization, e.g. folate cycle and cobalamin metabolism or pre-mRNA splicing and nuclear mRNA export. Two independent null models, shuffling of SCP labels within the network and shuffling of spots on the tissue slice, distinguish pairs that are functionally non-random and pairs that are additionally enriched within the immediate neighborhood, separating local from tissue-wide coordination. Our results suggest that tissue physiology reflects not only cell type composition but also functional coordination between neighboring regions.

**Supplementary Figures 6 - 8**

**A**

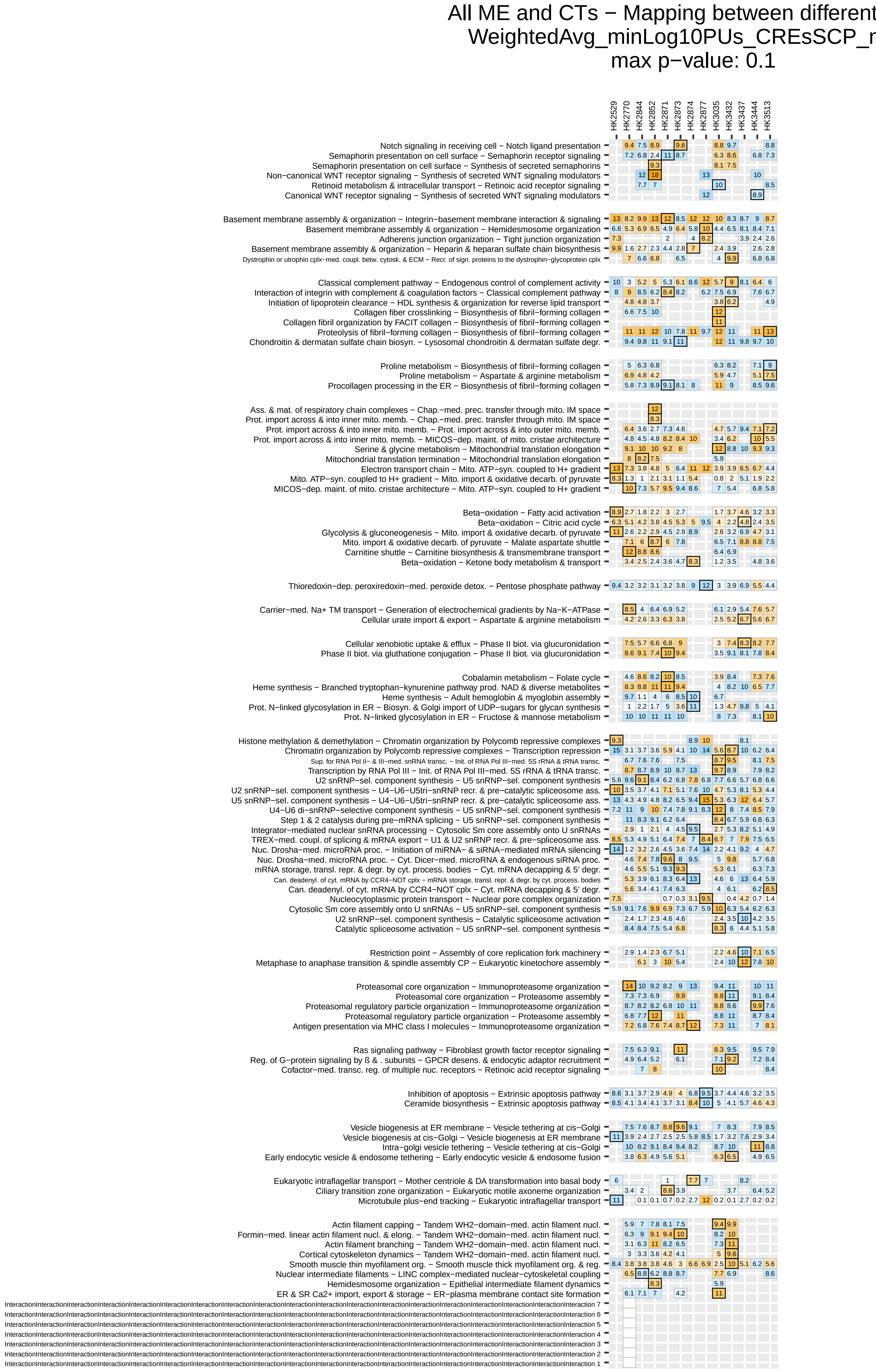
All ME and CTs − Mapping between different
WeightedAvg_minLog10PUs_CREsSCP_r
max p−value: 0.1
HK2529
HK2770
HK2844
HK2852
HK2871
HK2873
HK2874
HK2877
HK3035
HK3432
HK3437
HK3444
HK3513
Notch signaling in receiving cell − Notch ligand presentation
Semaphorin presentation on cell surface − Semaphorin receptor signaling
Semaphorin presentation on cell surface − Synthesis of secreted semaphorins
Non−canonical WNT receptor signaling − Synthesis of secreted WNT signaling modulators
Retinoid metabolism & intracellular transport − Retinoic acid receptor signaling
Canonical WNT receptor signaling − Synthesis of secreted WNT signaling modulators
Basement membrane assembly & organization − Integrin−basement membrane interaction & signaling
Basement membrane assembly & organization − Hemidesmosome organization
Adherens junction organization − Tight junction organization
Basement membrane assembly & organization − Heparin & heparan sulfate chain biosynthesis
Dystrophin or utrophin cplx−med. coupl. betw. cytosk. & ECM − Recr. of sign. proteins to the dystrophin−glycoprotein cplx
Classical complement pathway − Endogenous control of complement activity
Interaction of integrin with complement & coagulation factors − Classical complement pathway
Initiation of lipoprotein clearance − HDL synthesis & organization for reverse lipid transport
Collagen fiber crosslinking − Biosynthesis of fibril−forming collagen
Collagen fibril organization by FACIT collagen − Biosynthesis of fibril−forming collagen
Proteolysis of fibril−forming collagen − Biosynthesis of fibril−forming collagen
Chondroitin & dermatan sulfate chain biosyn. − Lysosomal chondroitin & dermatan sulfate degr.
Proline metabolism − Biosynthesis of fibril−forming collagen
Proline metabolism − Aspartate & arginine metabolism
Procollagen processing in the ER − Biosynthesis of fibril−forming collagen
Ass. & mat. of respiratory chain complexes − Chap.−med. prec. transfer through mito. IM space
Prot. import across & into inner mito. memb. − Chap.−med. prec. transfer through mito. IM space
Prot. import across & into inner mito. memb. − Prot. import across & into outer mito. memb.
Prot. import across & into inner mito. memb. − MICOS−dep. maint. of mito. cristae architecture
Serine & glycine metabolism − Mitochondrial translation elongation
Mitochondrial translation termination − Mitochondrial translation elongation
Electron transport chain − Mito. ATP−syn. coupled to H+ gradient
Mito. ATP−syn. coupled to H+ gradient − Mito. import & oxidative decarb. of pyruvate
MICOS−dep. maint. of mito. cristae architecture − Mito. ATP−syn. coupled to H+ gradient
Beta−oxidation − Fatty acid activation
Beta−oxidation − Citric acid cycle
Glycolysis & gluconeogenesis − Mito. import & oxidative decarb. of pyruvate
Mito. import & oxidative decarb. of pyruvate − Malate aspartate shuttle
Carnitine shuttle − Carnitine biosynthesis & transmembrane transport
Beta−oxidation − Ketone body metabolism & transport
Thioredoxin−dep. peroxiredoxin−med. peroxide detox. − Pentose phosphate pathway
Carrier−med. Na+ TM transport − Generation of electrochemical gradients by Na−K−ATPase
Cellular urate import & export − Aspartate & arginine metabolism
Cellular xenobiotic uptake & efflux − Phase II biot. via glucuronidation
Phase II biot. via gluthatione conjugation − Phase II biot. via glucuronidation
Cobalamin metabolism − Folate cycle
Heme synthesis − Branched tryptophan−kynurenine pathway prod. NAD & diverse metabolites
Heme synthesis − Adult hemoglobin & myoglobin assembly
Prot. N−linked glycosylation in ER − Biosyn. & Golgi import of UDP−sugars for glycan synthesis
Prot. N−linked glycosylation in ER − Fructose & mannose metabolism
Histone methylation & demethylation − Chromatin organization by Polycomb repressive complexes
Chromatin organization by Polycomb repressive complexes − Transcription repression
Sup. for RNA Pol II− & III−med. snRNA transc. − Init. of RNA Pol III−med. 5S rRNA & tRNA transc.
Transcription by RNA Pol III − Init. of RNA Pol III−med. 5S rRNA & tRNA transc.
U2 snRNP−sel. component synthesis − U5 snRNP−sel. component synthesis
U2 snRNP−sel. component synthesis − U4−U6−U5tri−snRNP recr. & pre−catalytic spliceosome ass.
U5 snRNP−sel. component synthesis − U4−U6−U5tri−snRNP recr. & pre−catalytic spliceosome ass.
U4−U6 di−snRNP−selective component synthesis − U5 snRNP−sel. component synthesis
Step 1 & 2 catalysis during pre−mRNA splicing − U5 snRNP−sel. component synthesis
Integrator−mediated nuclear snRNA processing − Cytosolic Sm core assembly onto U snRNAs
TREX−med. coupl. of splicing & mRNA export − U1 & U2 snRNP recr. & pre−spliceosome ass.
Nuc. Drosha−med. microRNA proc. − Initiation of miRNA− & siRNA−mediated mRNA silencing
Nuc. Drosha−med. microRNA proc. − Cyt. Dicer−med. microRNA & endogenous siRNA proc.
mRNA storage, transl. repr. & degr. by cyt. process. bodies − Cyt. mRNA decapping & 5' degr.
Can. deadenyl. of cyt. mRNA by CCR4−NOT cplx − mRNA storage, transl. repr. & degr. by cyt. process. bodies
Can. deadenyl. of cyt. mRNA by CCR4−NOT cplx − Cyt. mRNA decapping & 5' degr.
Nucleocytoplasmic protein transport − Nuclear pore complex organization
Cytosolic Sm core assembly onto U snRNAs − U5 snRNP−sel. component synthesis
U2 snRNP−sel. component synthesis − Catalytic spliceosome activation
Catalytic spliceosome activation − U5 snRNP−sel. component synthesis
Restriction point − Assembly of core replication fork machinery
Metaphase to anaphase transition & spindle assembly CP − Eukaryotic kinetochore assembly
Proteasomal core organization − Immunoproteasome organization
Proteasomal core organization − Proteasome assembly
Proteasomal regulatory particle organization − Immunoproteasome organization
Proteasomal regulatory particle organization − Proteasome assembly
Antigen presentation via MHC class I molecules − Immunoproteasome organization
Ras signaling pathway − Fibroblast growth factor receptor signaling
Reg. of G−protein signaling by ß & . subunits − GPCR desens. & endocytic adaptor recruitment
Cofactor−med. transc. reg. of multiple nuc. receptors − Retinoic acid receptor signaling
Inhibition of apoptosis − Extrinsic apoptosis pathway
Ceramide biosynthesis − Extrinsic apoptosis pathway
Vesicle biogenesis at ER membrane − Vesicle tethering at cis−Golgi
Vesicle biogenesis at cis−Golgi − Vesicle biogenesis at ER membrane
Intra−golgi vesicle tethering − Vesicle tethering at cis−Golgi
Early endocytic vesicle & endosome tethering − Early endocytic vesicle & endosome fusion
Eukaryotic intraflagellar transport − Mother centriole & DA transformation into basal body
Ciliary transition zone organization − Eukaryotic motile axoneme organization
Microtubule plus−end tracking − Eukaryotic intraflagellar transport
Actin filament capping − Tandem WH2−domain−med. actin filament nucl.
Formin−med. linear actin filament nucl. & elong. − Tandem WH2−domain−med. actin filament nucl.
Actin filament branching − Tandem WH2−domain−med. actin filament nucl.
Cortical cytoskeleton dynamics − Tandem WH2−domain−med. actin filament nucl.
Smooth muscle thin myofilament org. − Smooth muscle thick myofilament org. & reg.
Nuclear intermediate filaments − LINC complex−mediated nuclear−cytoskeletal coupling
Hemidesmosome organization − Epithelial intermediate filament dynamics
ER & SR Ca2+ import, export & storage − ER−plasma membrane contact site formation
InteractionInteractionInteractionInteractionInteractionInteractionInteractionInteractionInteractionInteractionInteractionInteractionInteractionInteractionInteractionInteractionInteractionInteractionInteractionInteraction 7
InteractionInteractionInteractionInteractionInteractionInteractionInteractionInteractionInteractionInteractionInteractionInteractionInteractionInteractionInteractionInteractionInteractionInteractionInteractionInteraction 6
InteractionInteractionInteractionInteractionInteractionInteractionInteractionInteractionInteractionInteractionInteractionInteractionInteractionInteractionInteractionInteractionInteractionInteractionInteractionInteraction 5
InteractionInteractionInteractionInteractionInteractionInteractionInteractionInteractionInteractionInteractionInteractionInteractionInteractionInteractionInteractionInteractionInteractionInteractionInteractionInteraction 4
InteractionInteractionInteractionInteractionInteractionInteractionInteractionInteractionInteractionInteractionInteractionInteractionInteractionInteractionInteractionInteractionInteractionInteractionInteractionInteraction 3
InteractionInteractionInteractionInteractionInteractionInteractionInteractionInteractionInteractionInteractionInteractionInteractionInteractionInteractionInteractionInteractionInteractionInteractionInteractionInteraction 2
InteractionInteractionInteractionInteractionInteractionInteractionInteractionInteractionInteractionInteractionInteractionInteractionInteractionInteractionInteractionInteractionInteractionInteractionInteractionInteraction 1

B

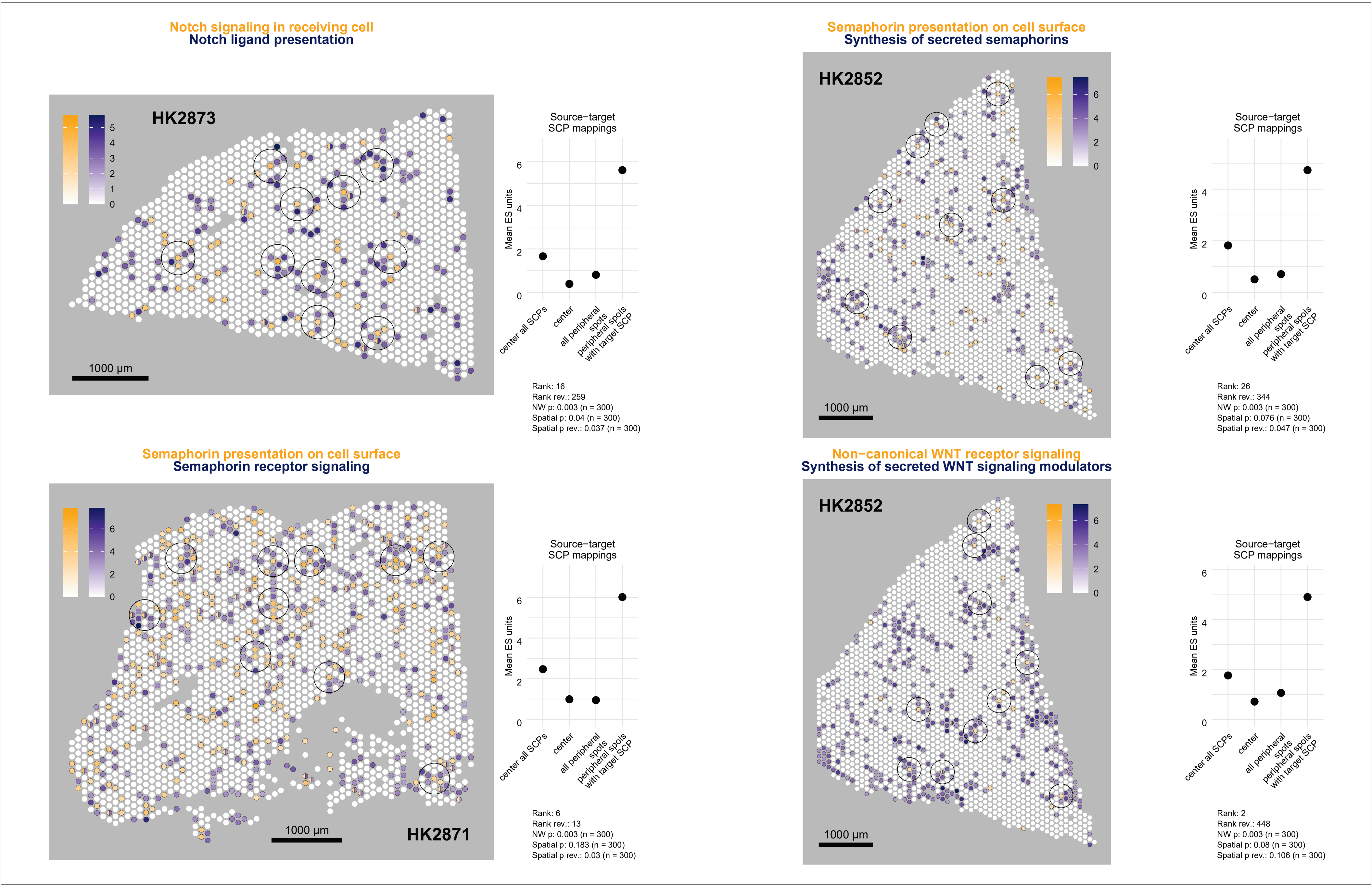
Notch signaling in receiving cell
Notch ligand presentation
HK2873
1000 µm
Source−target
SCP mappings
Mean ES units
center all SCPs
center
all peripheral spots
peripheral spots with target SCP
Rank: 16
Rank rev.: 259
NW p: 0.003 (n = 300)
Spatial p: 0.04 (n = 300)
Spatial p rev.: 0.037 (n = 300)
Semaphorin presentation on cell surface
Synthesis of secreted semaphorins
HK2852
1000 µm
Source−target
SCP mappings
Mean ES units
center all SCPs
center
all peripheral spots
peripheral spots with target SCP
Rank: 26
Rank rev.: 344
NW p: 0.003 (n = 300)
Spatial p: 0.076 (n = 300)
Spatial p rev.: 0.047 (n = 300)
Semaphorin presentation on cell surface
Semaphorin receptor signaling
HK2871
1000 µm
Source−target
SCP mappings
Mean ES units
center all SCPs
center
all peripheral spots
peripheral spots with target SCP
Rank: 6
Rank rev.: 13
NW p: 0.003 (n = 300)
Spatial p: 0.183 (n = 300)
Spatial p rev.: 0.03 (n = 300)
Non−canonical WNT receptor signaling
Synthesis of secreted WNT signaling modulators
HK2852
1000 µm
Source−target
SCP mappings
Mean ES units
center all SCPs
center
all peripheral spots
peripheral spots with target SCP
Rank: 2
Rank rev.: 448
NW p: 0.003 (n = 300)
Spatial p: 0.08 (n = 300)
Spatial p rev.: 0.106 (n = 300)

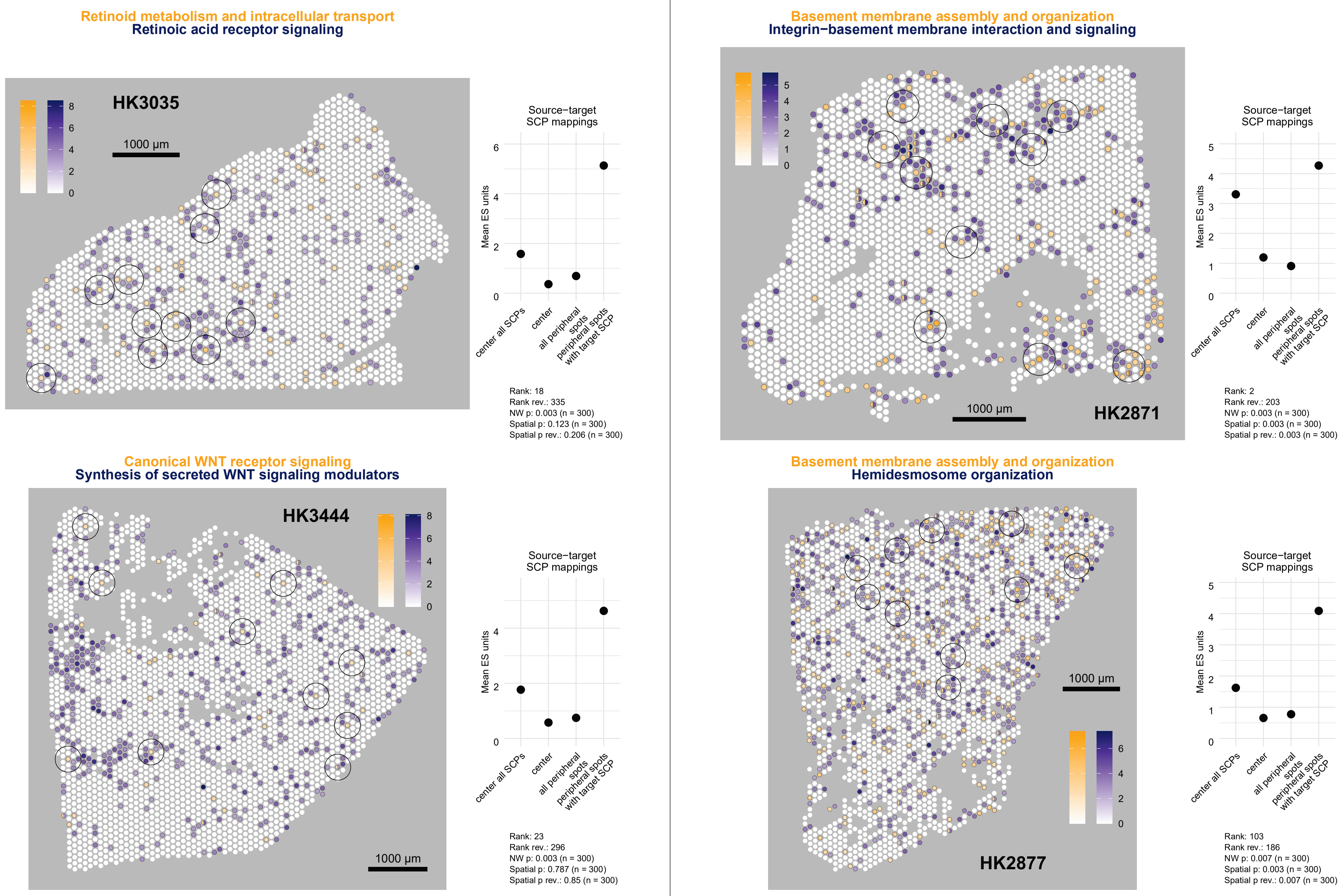

Retinoid metabolism and intracellular transport
Retinoic acid receptor signaling
HK3035
1000 µm
0
2
4
6
8
Source−target
SCP mappings
Mean ES units
center all SCPs
center
all peripheral spots
peripheral spots with target SCP
Rank: 18
Rank rev.: 335
NW p: 0.003 (n = 300)
Spatial p: 0.123 (n = 300)
Spatial p rev.: 0.206 (n = 300)
Basement membrane assembly and organization
Integrin−basement membrane interaction and signaling
HK2871
1000 µm
0
1
2
3
4
5
Source−target
SCP mappings
Mean ES units
center all SCPs
center
all peripheral spots
peripheral spots with target SCP
Rank: 2
Rank rev.: 203
NW p: 0.003 (n = 300)
Spatial p: 0.003 (n = 300)
Spatial p rev.: 0.003 (n = 300)
Canonical WNT receptor signaling
Synthesis of secreted WNT signaling modulators
HK3444
1000 µm
0
2
4
6
8
Source−target
SCP mappings
Mean ES units
center all SCPs
center
all peripheral spots
peripheral spots with target SCP
Rank: 23
Rank rev.: 296
NW p: 0.003 (n = 300)
Spatial p: 0.787 (n = 300)
Spatial p rev.: 0.85 (n = 300)
Basement membrane assembly and organization
Hemidesmosome organization
HK2877
1000 µm
0
2
4
6
Source−target
SCP mappings
Mean ES units
center all SCPs
center
all peripheral spots
peripheral spots with target SCP
Rank: 103
Rank rev.: 186
NW p: 0.007 (n = 300)
Spatial p: 0.003 (n = 300)
Spatial p rev.: 0.007 (n = 300)

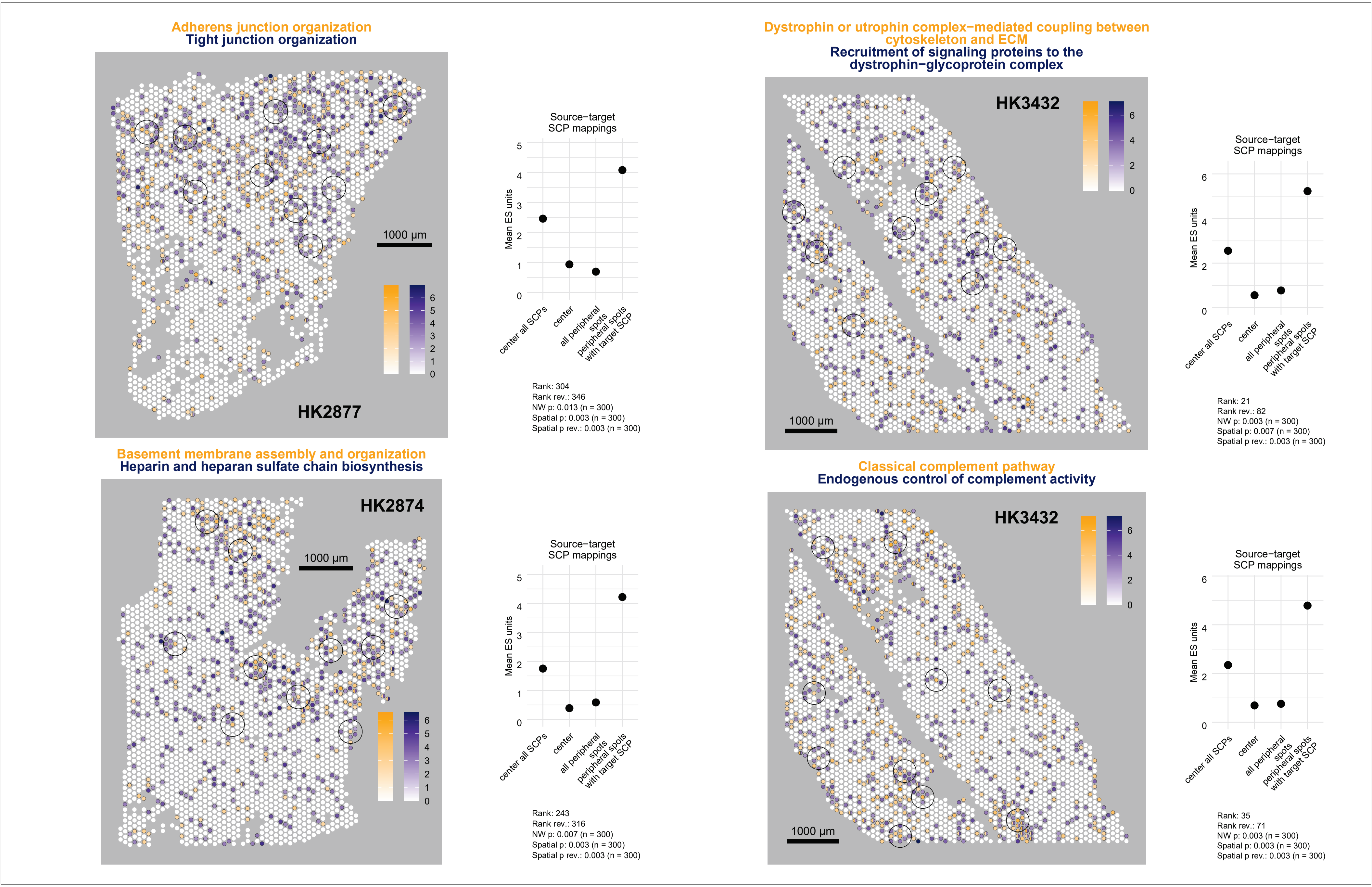

Adherens junction organization
Tight junction organization
HK2877
1000 µm
0
1
2
3
4
5
6
Source−target
SCP mappings
Mean ES units
center all SCPs
center
all peripheral spots
peripheral spots with target SCP
Rank: 304
Rank rev.: 346
NW p: 0.013 (n = 300)
Spatial p: 0.003 (n = 300)
Spatial p rev.: 0.003 (n = 300)
Basement membrane assembly and organization
Heparin and heparan sulfate chain biosynthesis
HK2874
1000 µm
Source−target
SCP mappings
Mean ES units
center all SCPs
center
all peripheral spots
peripheral spots with target SCP
Rank: 243
Rank rev.: 316
NW p: 0.007 (n = 300)
Spatial p: 0.003 (n = 300)
Spatial p rev.: 0.003 (n = 300)
Dystrophin or utrophin complex−mediated coupling between cytoskeleton and ECM
Recruitment of signaling proteins to the dystrophin−glycoprotein complex
HK3432
1000 µm
Source−target
SCP mappings
Mean ES units
center all SCPs
center
all peripheral spots
peripheral spots with target SCP
Rank: 21
Rank rev.: 82
NW p: 0.003 (n = 300)
Spatial p: 0.007 (n = 300)
Spatial p rev.: 0.003 (n = 300)
Classical complement pathway
Endogenous control of complement activity
HK3432
1000 µm
Source−target
SCP mappings
Mean ES units
center all SCPs
center
all peripheral spots
peripheral spots with target SCP
Rank: 35
Rank rev.: 71
NW p: 0.003 (n = 300)
Spatial p: 0.003 (n = 300)
Spatial p rev.: 0.003 (n = 300)

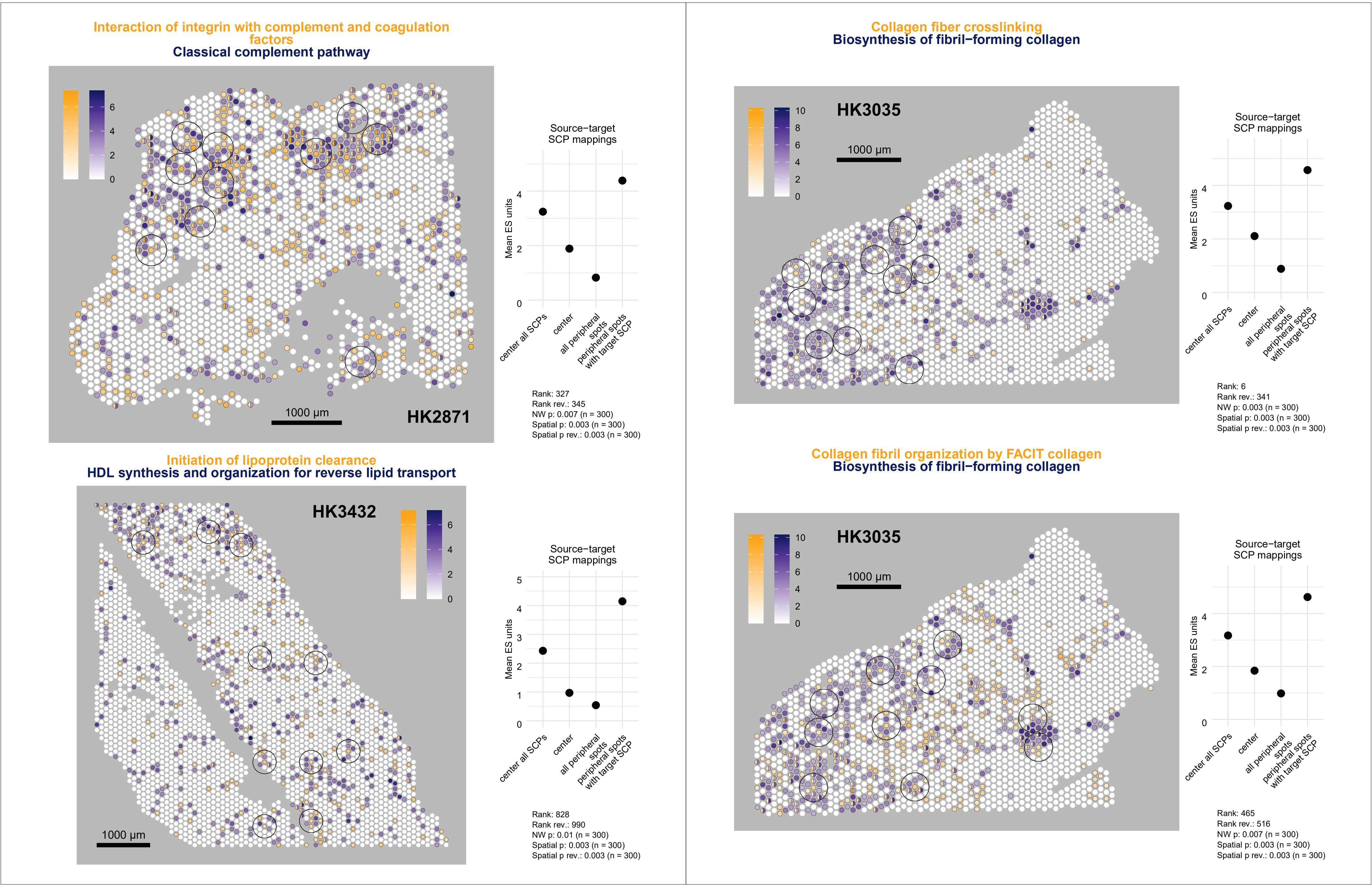
Interaction of integrin with complement and coagulation factors
Classical complement pathway
HK2871
1000 µm
Source−target SCP mappings
Mean ES units
center all SCPs
center
all peripheral spots
peripheral spots with target SCP
Rank: 327
Rank rev.: 345
NW p: 0.007 (n = 300)
Spatial p: 0.003 (n = 300)
Spatial p rev.: 0.003 (n = 300)
Collagen fiber crosslinking
Biosynthesis of fibril−forming collagen
HK3035
1000 µm
Source−target SCP mappings
Mean ES units
center all SCPs
center
all peripheral spots
peripheral spots with target SCP
Rank: 6
Rank rev.: 341
NW p: 0.003 (n = 300)
Spatial p: 0.003 (n = 300)
Spatial p rev.: 0.003 (n = 300)
Initiation of lipoprotein clearance
HDL synthesis and organization for reverse lipid transport
HK3432
1000 µm
Source−target SCP mappings
Mean ES units
center all SCPs
center
all peripheral spots
peripheral spots with target SCP
Rank: 828
Rank rev.: 990
NW p: 0.01 (n = 300)
Spatial p: 0.003 (n = 300)
Spatial p rev.: 0.003 (n = 300)
Collagen fibril organization by FACIT collagen
Biosynthesis of fibril−forming collagen
HK3035
1000 µm
Source−target SCP mappings
Mean ES units
center all SCPs
center
all peripheral spots
peripheral spots with target SCP
Rank: 465
Rank rev.: 516
NW p: 0.007 (n = 300)
Spatial p: 0.003 (n = 300)
Spatial p rev.: 0.003 (n = 300)

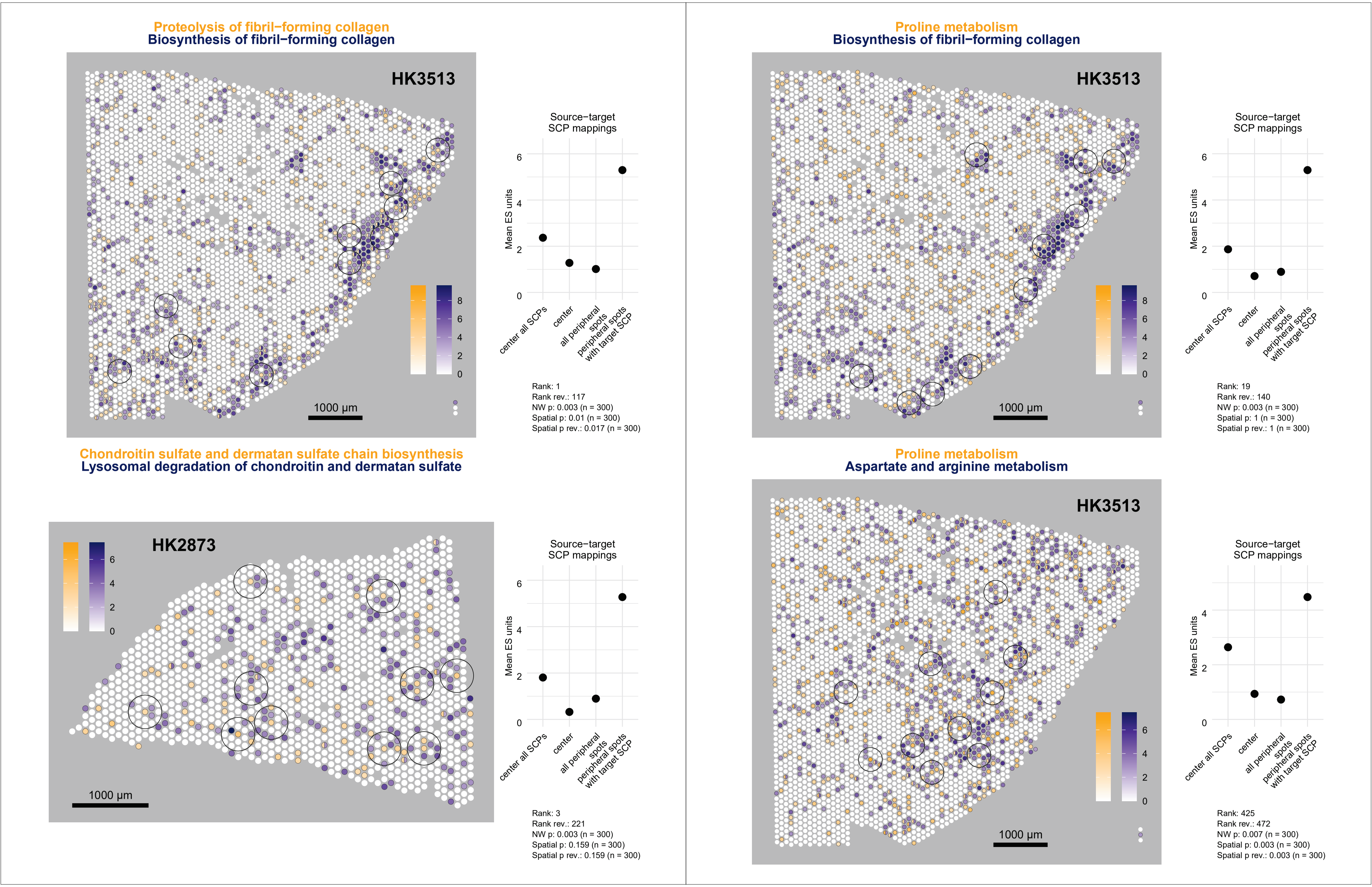

Proteolysis of fibril−forming collagen
Biosynthesis of fibril−forming collagen
HK3513
1000 µm
Source−target SCP mappings
Mean ES units
center all SCPs
center
all peripheral spots
peripheral spots with target SCP
Rank: 1
Rank rev.: 117
NW p: 0.003 (n = 300)
Spatial p: 0.01 (n = 300)
Spatial p rev.: 0.017 (n = 300)
Proline metabolism
Biosynthesis of fibril−forming collagen
HK3513
1000 µm
Source−target SCP mappings
Mean ES units
center all SCPs
center
all peripheral spots
peripheral spots with target SCP
Rank: 19
Rank rev.: 140
NW p: 0.003 (n = 300)
Spatial p: 1 (n = 300)
Spatial p rev.: 1 (n = 300)
Chondroitin sulfate and dermatan sulfate chain biosynthesis
Lysosomal degradation of chondroitin and dermatan sulfate
HK2873
1000 µm
Source−target SCP mappings
Mean ES units
center all SCPs
center
all peripheral spots
peripheral spots with target SCP
Rank: 3
Rank rev.: 221
NW p: 0.003 (n = 300)
Spatial p: 0.159 (n = 300)
Spatial p rev.: 0.159 (n = 300)
Proline metabolism
Aspartate and arginine metabolism
HK3513
1000 µm
Source−target SCP mappings
Mean ES units
center all SCPs
center
all peripheral spots
peripheral spots with target SCP
Rank: 425
Rank rev.: 472
NW p: 0.007 (n = 300)
Spatial p: 0.003 (n = 300)
Spatial p rev.: 0.003 (n = 300)

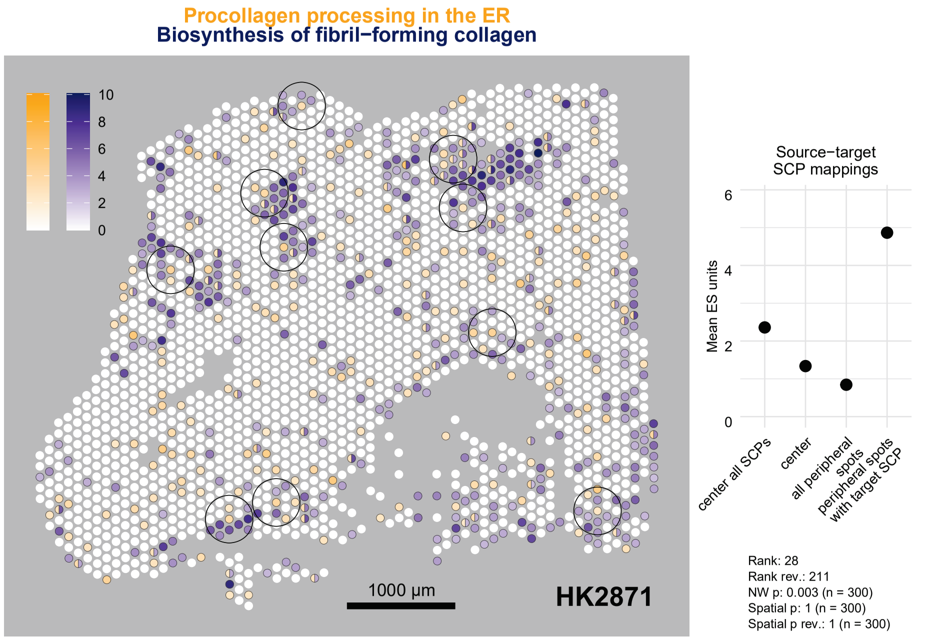
Procollagen processing in the ER
Biosynthesis of fibril−forming collagen
HK2871
1000 µm
0
2
4
6
8
10
Source−target
SCP mappings
Mean ES units
center all SCPs
center
all peripheral spots
peripheral spots with target SCP
Rank: 28
Rank rev.: 211
NW p: 0.003 (n = 300)
Spatial p: 1 (n = 300)
Spatial p rev.: 1 (n = 300)

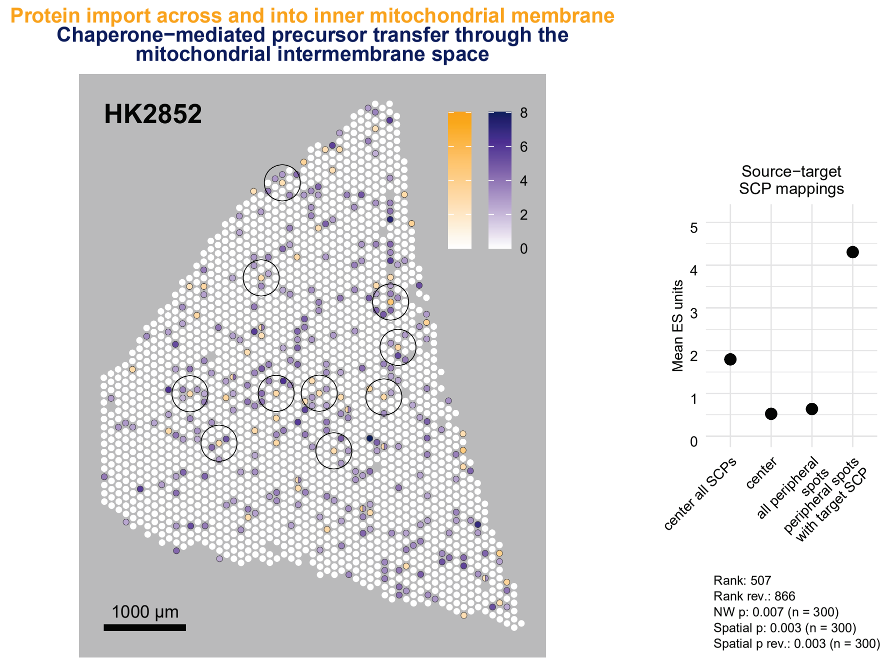
Protein import across and into inner mitochondrial membrane
Chaperone−mediated precursor transfer through the mitochondrial intermembrane space
HK2852
1000 µm
0
2
4
6
8
Source−target
SCP mappings
Mean ES units
center all SCPs
center
all peripheral spots
peripheral spots with target SCP
Rank: 507
Rank rev.: 866
NW p: 0.007 (n = 300)
Spatial p: 0.003 (n = 300)
Spatial p rev.: 0.003 (n = 300)

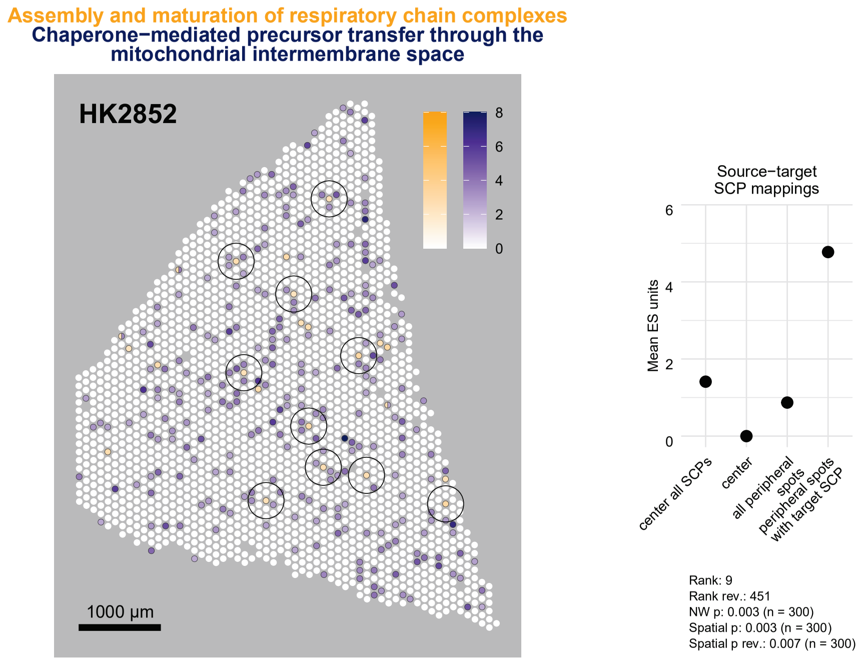
Assembly and maturation of respiratory chain complexes
Chaperone−mediated precursor transfer through the mitochondrial intermembrane space
HK2852
1000 µm
0
2
4
6
8
Source−target
SCP mappings
Mean ES units
center all SCPs
center
all peripheral spots
peripheral spots with target SCP
Rank: 9
Rank rev.: 451
NW p: 0.003 (n = 300)
Spatial p: 0.003 (n = 300)
Spatial p rev.: 0.007 (n = 300)

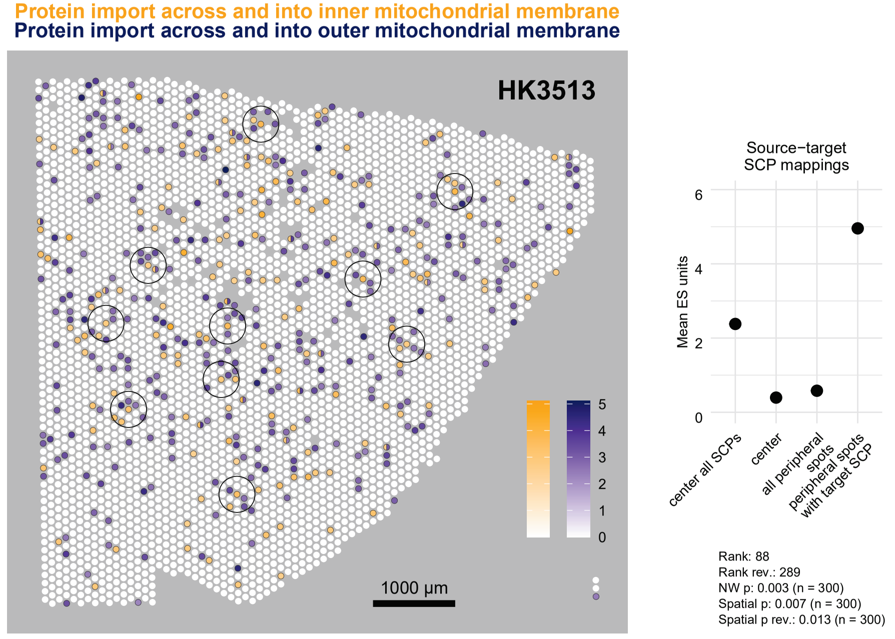
Protein import across and into inner mitochondrial membrane
Protein import across and into outer mitochondrial membrane
HK3513
1000 µm
0
1
2
3
4
5
Source−target
SCP mappings
Mean ES units
center all SCPs
center
all peripheral spots
peripheral spots with target SCP
Rank: 88
Rank rev.: 289
NW p: 0.003 (n = 300)
Spatial p: 0.007 (n = 300)
Spatial p rev.: 0.013 (n = 300)

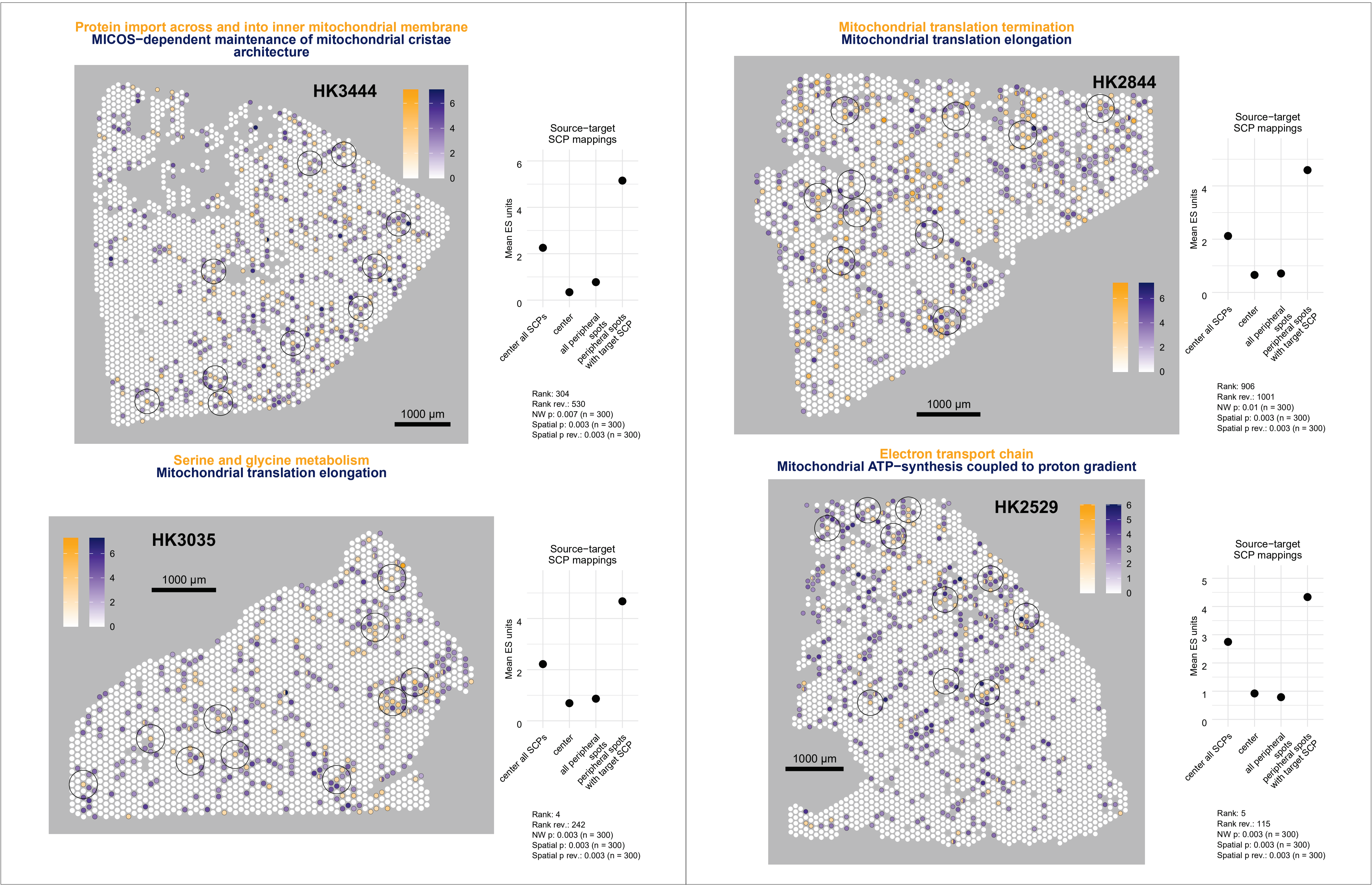

Protein import across and into inner mitochondrial membrane
MICOS−dependent maintenance of mitochondrial cristae architecture
HK3444
1000 µm
Source−target SCP mappings
Mean ES units
center all SCPs
center
all peripheral spots
peripheral spots with target SCP
Rank: 304
Rank rev.: 530
NW p: 0.007 (n = 300)
Spatial p: 0.003 (n = 300)
Spatial p rev.: 0.003 (n = 300)
Mitochondrial translation termination
Mitochondrial translation elongation
HK2844
1000 µm
Source−target SCP mappings
Mean ES units
center all SCPs
center
all peripheral spots
peripheral spots with target SCP
Rank: 906
Rank rev.: 1001
NW p: 0.01 (n = 300)
Spatial p: 0.003 (n = 300)
Spatial p rev.: 0.003 (n = 300)
Serine and glycine metabolism
Mitochondrial translation elongation
HK3035
1000 µm
Source−target SCP mappings
Mean ES units
center all SCPs
center
all peripheral spots
peripheral spots with target SCP
Rank: 4
Rank rev.: 242
NW p: 0.003 (n = 300)
Spatial p: 0.003 (n = 300)
Spatial p rev.: 0.003 (n = 300)
Electron transport chain
Mitochondrial ATP−synthesis coupled to proton gradient
HK2529
1000 µm
Source−target SCP mappings
Mean ES units
center all SCPs
center
all peripheral spots
peripheral spots with target SCP
Rank: 5
Rank rev.: 115
NW p: 0.003 (n = 300)
Spatial p: 0.003 (n = 300)
Spatial p rev.: 0.003 (n = 300)

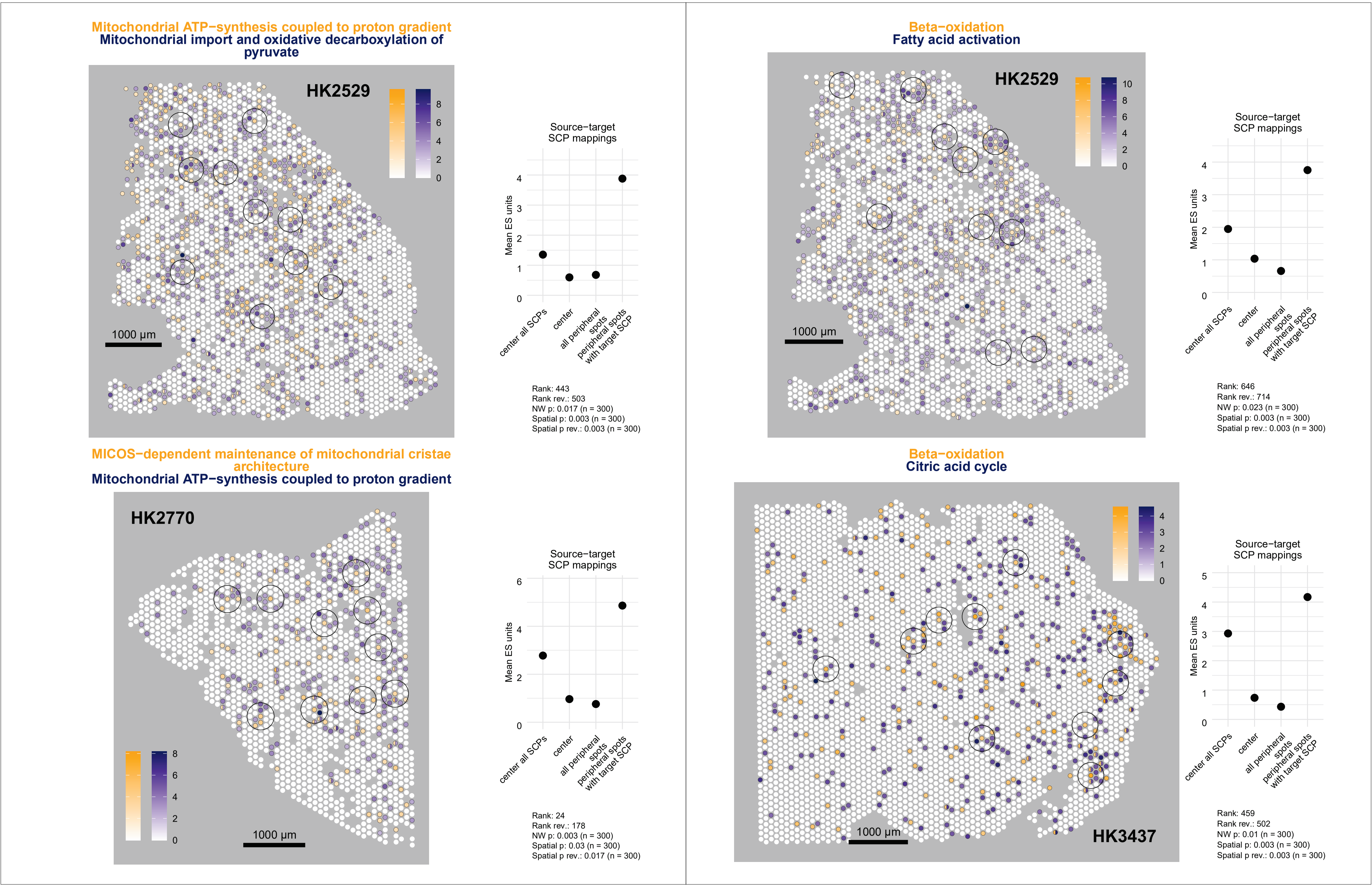

Mitochondrial ATP−synthesis coupled to proton gradient
Mitochondrial import and oxidative decarboxylation of pyruvate
HK2529
1000 µm
Source−target SCP mappings
Mean ES units
center all SCPs
center
all peripheral spots
peripheral spots with target SCP
Rank: 443
Rank rev.: 503
NW p: 0.017 (n = 300)
Spatial p: 0.003 (n = 300)
Spatial p rev.: 0.003 (n = 300)
Beta−oxidation
Fatty acid activation
HK2529
1000 µm
Source−target SCP mappings
Mean ES units
center all SCPs
center
all peripheral spots
peripheral spots with target SCP
Rank: 646
Rank rev.: 714
NW p: 0.023 (n = 300)
Spatial p: 0.003 (n = 300)
Spatial p rev.: 0.003 (n = 300)
MICOS−dependent maintenance of mitochondrial cristae architecture
Mitochondrial ATP−synthesis coupled to proton gradient
HK2770
1000 µm
Source−target SCP mappings
Mean ES units
center all SCPs
center
all peripheral spots
peripheral spots with target SCP
Rank: 24
Rank rev.: 178
NW p: 0.003 (n = 300)
Spatial p: 0.03 (n = 300)
Spatial p rev.: 0.017 (n = 300)
Beta−oxidation
Citric acid cycle
HK3437
1000 µm
Source−target SCP mappings
Mean ES units
center all SCPs
center
all peripheral spots
peripheral spots with target SCP
Rank: 459
Rank rev.: 502
NW p: 0.01 (n = 300)
Spatial p: 0.003 (n = 300)
Spatial p rev.: 0.003 (n = 300)

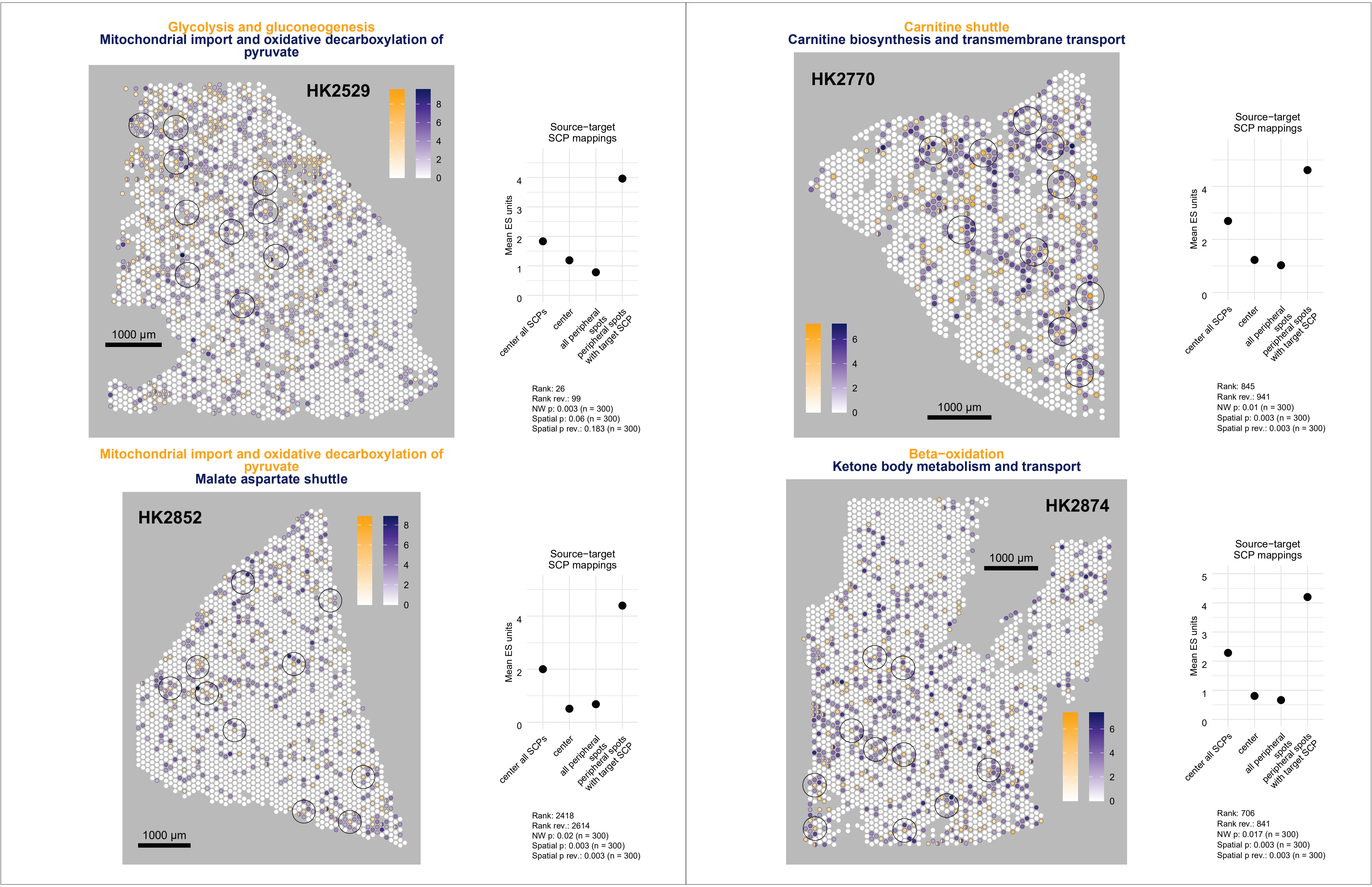

Glycolysis and gluconeogenesis
Mitochondrial import and oxidative decarboxylation of pyruvate
HK2529
1000 µm
Source−target SCP mappings
Mean ES units
center all SCPs
center
all peripheral spots
peripheral spots with target SCP
Rank: 26
Rank rev.: 99
NW p: 0.003 (n = 300)
Spatial p: 0.06 (n = 300)
Spatial p rev.: 0.183 (n = 300)
Carnitine shuttle
Carnitine biosynthesis and transmembrane transport
HK2770
1000 µm
Source−target SCP mappings
Mean ES units
center all SCPs
center
all peripheral spots
peripheral spots with target SCP
Rank: 845
Rank rev.: 941
NW p: 0.01 (n = 300)
Spatial p: 0.003 (n = 300)
Spatial p rev.: 0.003 (n = 300)
Mitochondrial import and oxidative decarboxylation of pyruvate
Malate aspartate shuttle
HK2852
1000 µm
Source−target SCP mappings
Mean ES units
center all SCPs
center
all peripheral spots
peripheral spots with target SCP
Rank: 2418
Rank rev.: 2614
NW p: 0.02 (n = 300)
Spatial p: 0.003 (n = 300)
Spatial p rev.: 0.003 (n = 300)
Beta−oxidation
Ketone body metabolism and transport
HK2874
1000 µm
Source−target SCP mappings
Mean ES units
center all SCPs
center
all peripheral spots
peripheral spots with target SCP
Rank: 706
Rank rev.: 841
NW p: 0.017 (n = 300)
Spatial p: 0.003 (n = 300)
Spatial p rev.: 0.003 (n = 300)

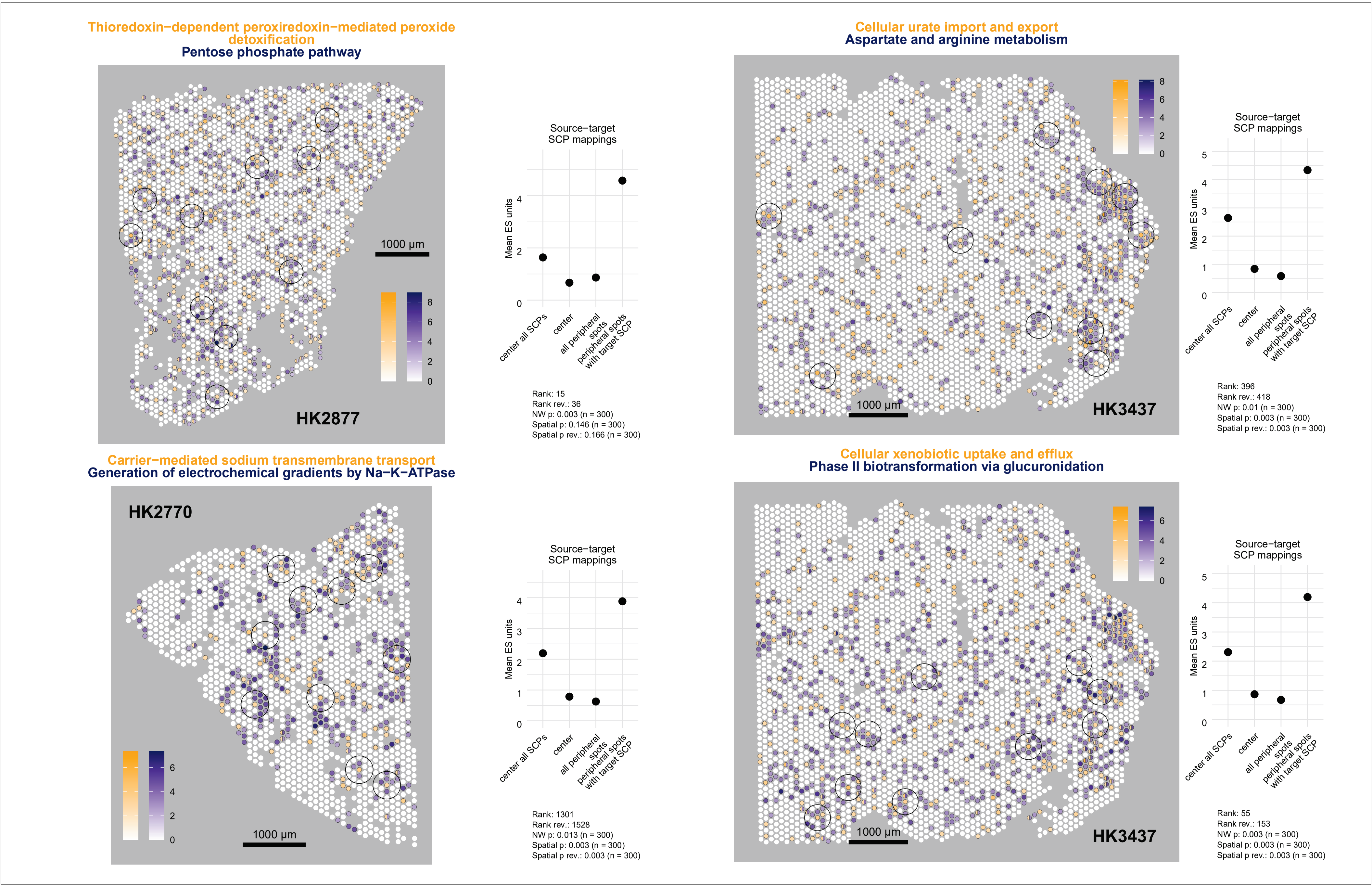

Thioredoxin−dependent peroxiredoxin−mediated peroxide detoxification
Pentose phosphate pathway
HK2877
1000 µm
0 2 4 6 8
Source−target SCP mappings
Mean ES units
0 2 4
center all SCPs
center
all peripheral spots
peripheral spots with target SCP
Rank: 15
Rank rev.: 36
NW p: 0.003 (n = 300)
Spatial p: 0.146 (n = 300)
Spatial p rev.: 0.166 (n = 300)
Cellular urate import and export
Aspartate and arginine metabolism
HK3437
1000 µm
0 2 4 6 8
Source−target SCP mappings
Mean ES units
0 1 2 3 4 5
center all SCPs
center
all peripheral spots
peripheral spots with target SCP
Rank: 396
Rank rev.: 418
NW p: 0.01 (n = 300)
Spatial p: 0.003 (n = 300)
Spatial p rev.: 0.003 (n = 300)
Carrier−mediated sodium transmembrane transport
Generation of electrochemical gradients by Na−K−ATPase
HK2770
1000 µm
0 2 4 6
Source−target SCP mappings
Mean ES units
0 1 2 3 4
center all SCPs
center
all peripheral spots
peripheral spots with target SCP
Rank: 1301
Rank rev.: 1528
NW p: 0.013 (n = 300)
Spatial p: 0.003 (n = 300)
Spatial p rev.: 0.003 (n = 300)
Cellular xenobiotic uptake and efflux
Phase II biotransformation via glucuronidation
HK3437
1000 µm
0 2 4 6
Source−target SCP mappings
Mean ES units
0 1 2 3 4 5
center all SCPs
center
all peripheral spots
peripheral spots with target SCP
Rank: 55
Rank rev.: 153
NW p: 0.003 (n = 300)
Spatial p: 0.003 (n = 300)
Spatial p rev.: 0.003 (n = 300)

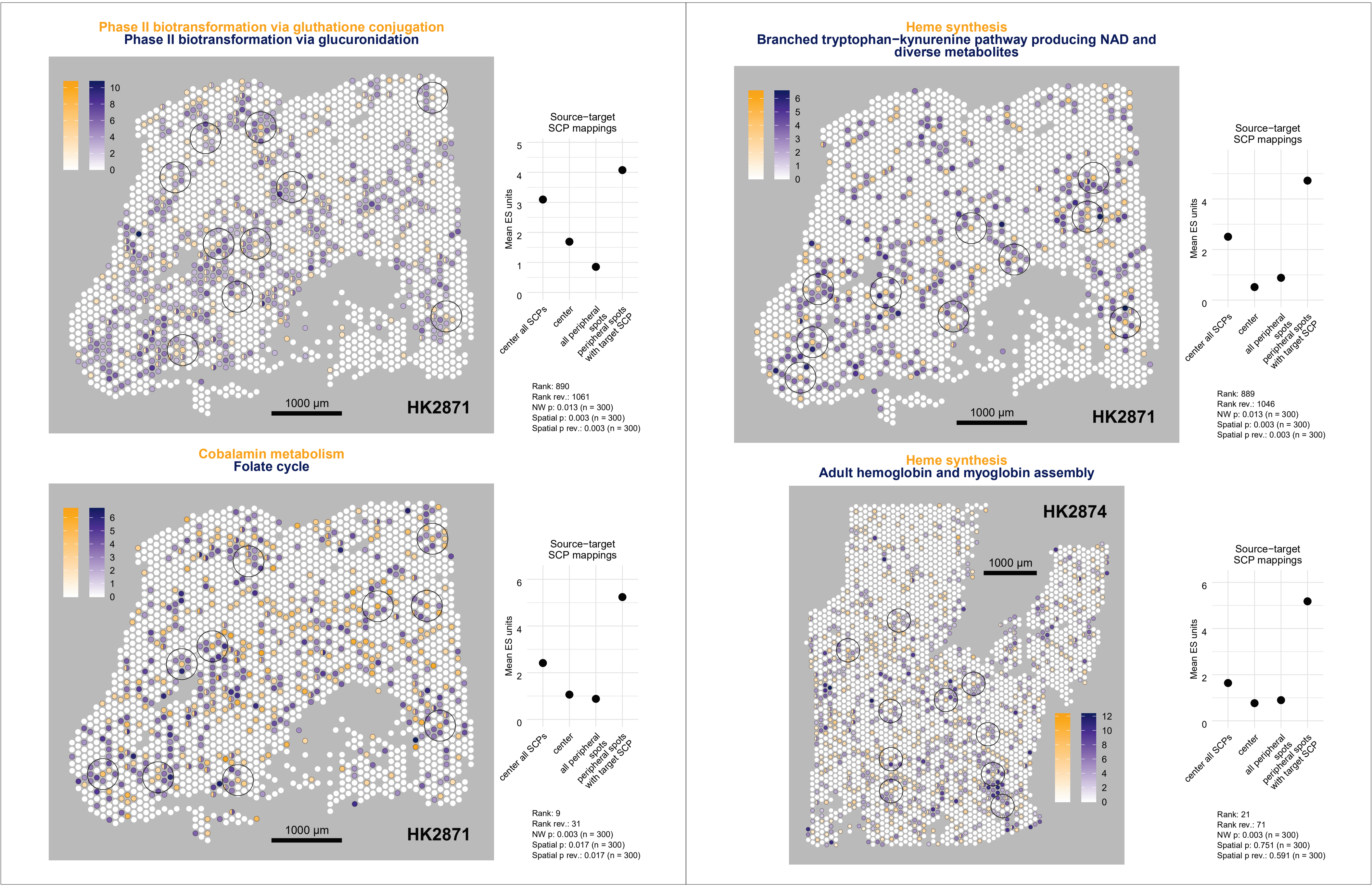

Phase II biotransformation via gluthatione conjugation
Phase II biotransformation via glucuronidation
HK2871
1000 µm
Source−target
SCP mappings
Mean ES units
center all SCPs
center
all peripheral spots
peripheral spots with target SCP
Rank: 890
Rank rev.: 1061
NW p: 0.013 (n = 300)
Spatial p: 0.003 (n = 300)
Spatial p rev.: 0.003 (n = 300)
Heme synthesis
Branched tryptophan−kynurenine pathway producing NAD and diverse metabolites
HK2871
1000 µm
Source−target
SCP mappings
Mean ES units
center all SCPs
center
all peripheral spots
peripheral spots with target SCP
Rank: 889
Rank rev.: 1046
NW p: 0.013 (n = 300)
Spatial p: 0.003 (n = 300)
Spatial p rev.: 0.003 (n = 300)
Cobalamin metabolism
Folate cycle
HK2871
1000 µm
Source−target
SCP mappings
Mean ES units
center all SCPs
center
all peripheral spots
peripheral spots with target SCP
Rank: 9
Rank rev.: 31
NW p: 0.003 (n = 300)
Spatial p: 0.017 (n = 300)
Spatial p rev.: 0.017 (n = 300)
Heme synthesis
Adult hemoglobin and myoglobin assembly
HK2874
1000 µm
Source−target
SCP mappings
Mean ES units
center all SCPs
center
all peripheral spots
peripheral spots with target SCP
Rank: 21
Rank rev.: 71
NW p: 0.003 (n = 300)
Spatial p: 0.751 (n = 300)
Spatial p rev.: 0.591 (n = 300)

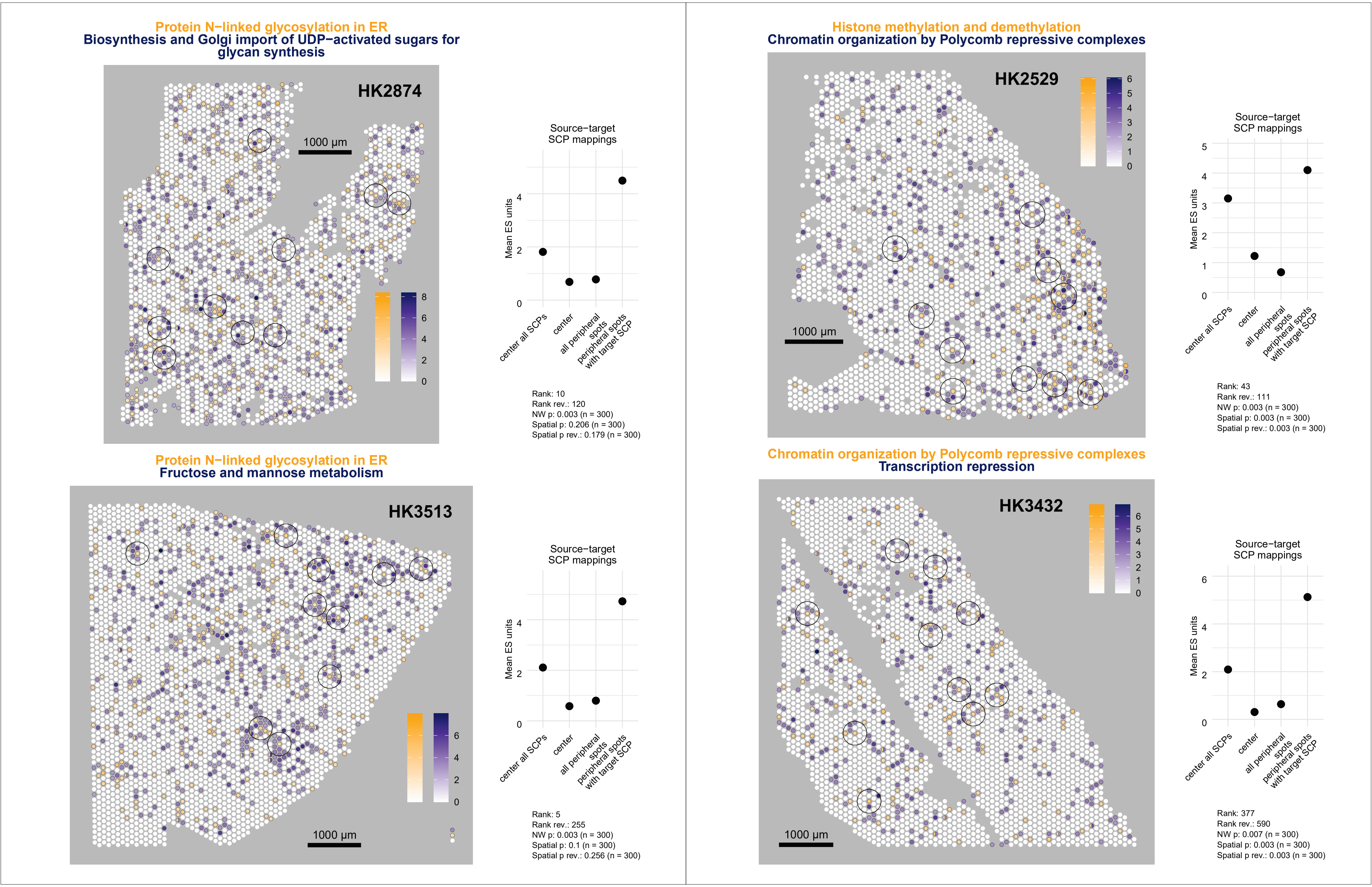
Protein N−linked glycosylation in ER
Biosynthesis and Golgi import of UDP−activated sugars for glycan synthesis
HK2874
1000 µm
0
2
4
6
8
Source−target SCP mappings
Mean ES units
0
2
4
center all SCPs
center
all peripheral spots
peripheral spots with target SCP
Rank: 10
Rank rev.: 120
NW p: 0.003 (n = 300)
Spatial p: 0.206 (n = 300)
Spatial p rev.: 0.179 (n = 300)
Histone methylation and demethylation
Chromatin organization by Polycomb repressive complexes
HK2529
1000 µm
0
1
2
3
4
5
6
Source−target SCP mappings
Mean ES units
0
1
2
3
4
5
center all SCPs
center
all peripheral spots
peripheral spots with target SCP
Rank: 43
Rank rev.: 111
NW p: 0.003 (n = 300)
Spatial p: 0.003 (n = 300)
Spatial p rev.: 0.003 (n = 300)
Protein N−linked glycosylation in ER
Fructose and mannose metabolism
HK3513
1000 µm
0
2
4
6
Source−target SCP mappings
Mean ES units
0
2
4
center all SCPs
center
all peripheral spots
peripheral spots with target SCP
Rank: 5
Rank rev.: 255
NW p: 0.003 (n = 300)
Spatial p: 0.1 (n = 300)
Spatial p rev.: 0.256 (n = 300)
Chromatin organization by Polycomb repressive complexes
Transcription repression
HK3432
1000 µm
0
1
2
3
4
5
6
Source−target SCP mappings
Mean ES units
0
2
4
6
center all SCPs
center
all peripheral spots
peripheral spots with target SCP
Rank: 377
Rank rev.: 590
NW p: 0.007 (n = 300)
Spatial p: 0.003 (n = 300)
Spatial p rev.: 0.003 (n = 300)

**Support for RNA polymerase II− and III−mediated snRNA transcription**
**Initiation of RNA polymerase III−mediated 5S rRNA and tRNA transcription**

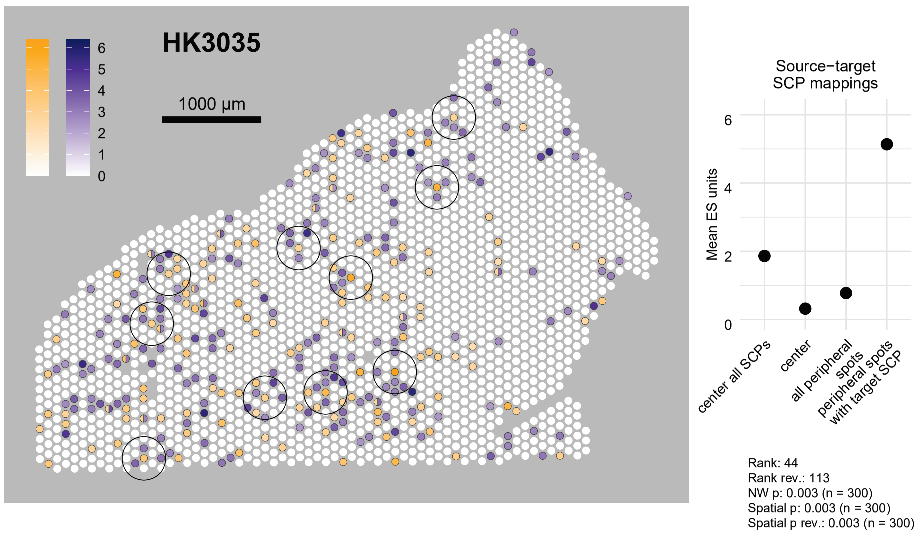


**U2 snRNP−selective component synthesis**
**U5 snRNP−selective component synthesis**

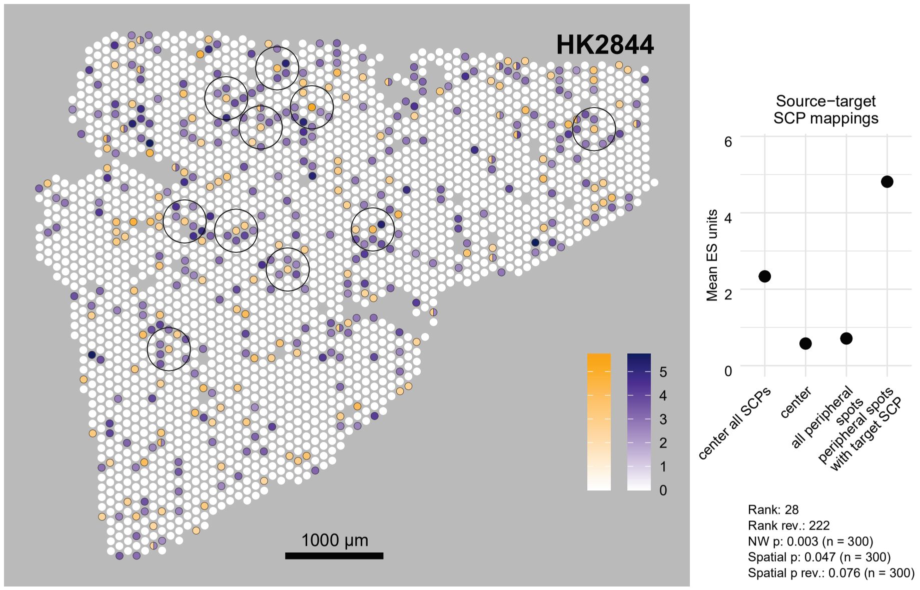


**Transcription by RNA polymerase III**
**Initiation of RNA polymerase III−mediated 5S rRNA and tRNA transcription**

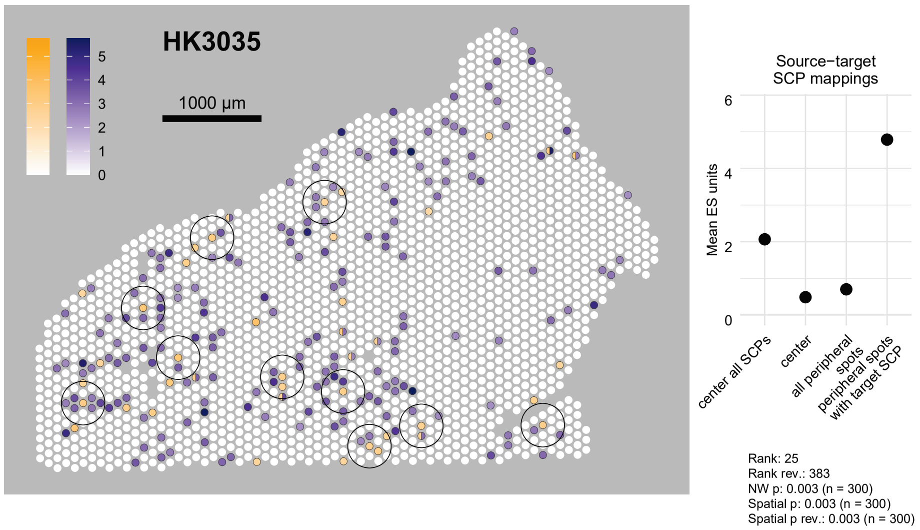


**U2 snRNP−selective component synthesis**
**U4−U6−U5 tri−snRNP recruitment and pre−catalytic spliceosome assembly**

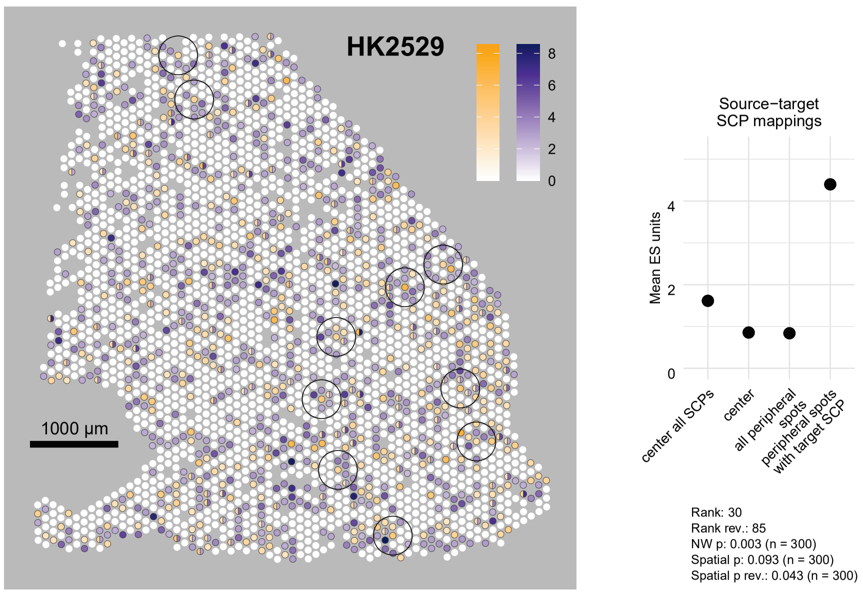

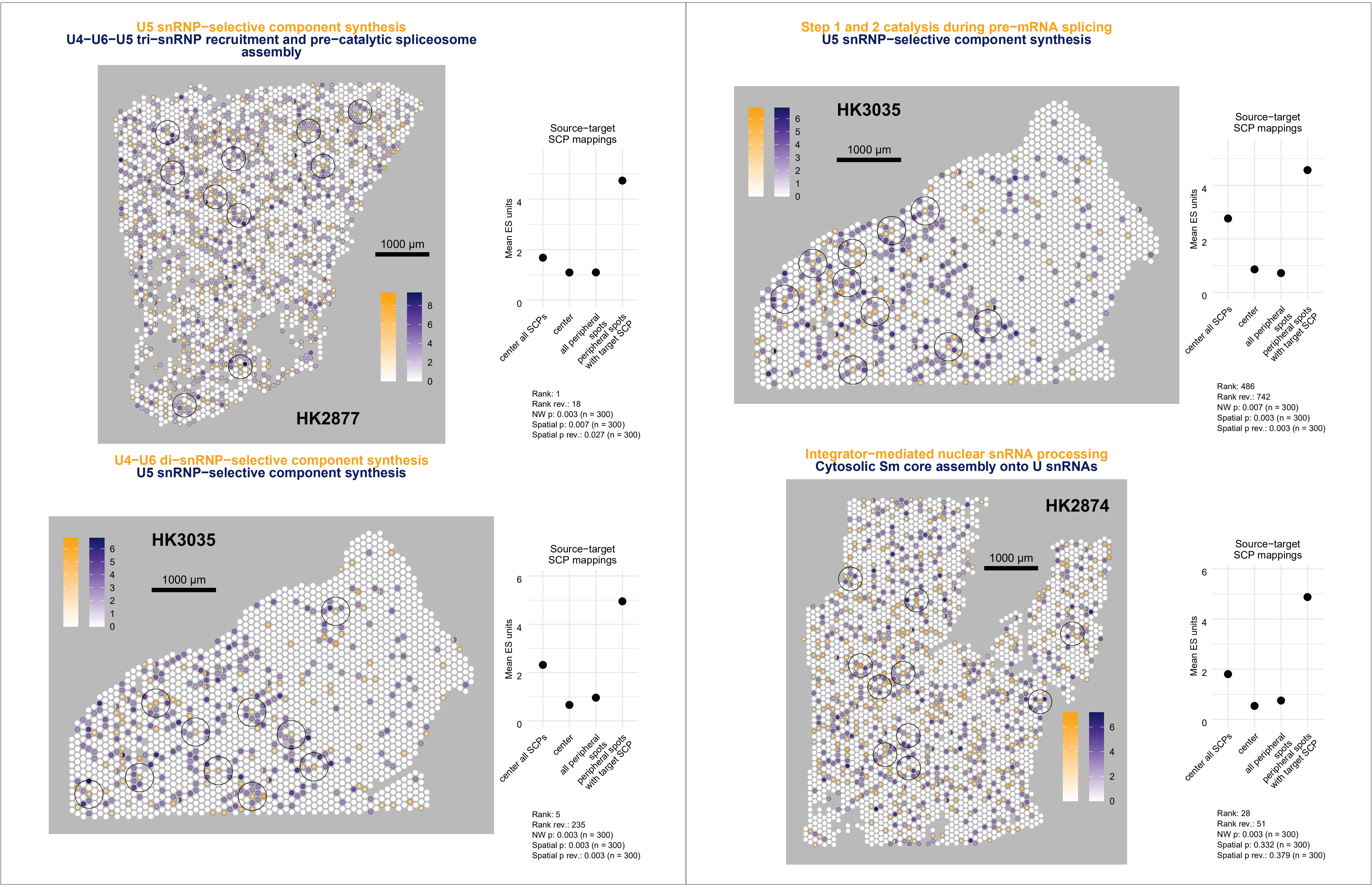

U5 snRNP−selective component synthesis
U4−U6−U5 tri−snRNP recruitment and pre−catalytic spliceosome assembly
HK2877
1000 µm
0 2 4 6 8
Source−target SCP mappings
Mean ES units
0 2 4
center all SCPs
center
all peripheral spots
peripheral spots with target SCP
Rank: 1
Rank rev.: 18
NW p: 0.003 (n = 300)
Spatial p: 0.007 (n = 300)
Spatial p rev.: 0.027 (n = 300)
Step 1 and 2 catalysis during pre−mRNA splicing
U5 snRNP−selective component synthesis
HK3035
1000 µm
0 1 2 3 4 5 6
Source−target SCP mappings
Mean ES units
0 2 4
center all SCPs
center
all peripheral spots
peripheral spots with target SCP
Rank: 486
Rank rev.: 742
NW p: 0.007 (n = 300)
Spatial p: 0.003 (n = 300)
Spatial p rev.: 0.003 (n = 300)
U4−U6 di−snRNP−selective component synthesis
U5 snRNP−selective component synthesis
HK3035
1000 µm
0 1 2 3 4 5 6
Source−target SCP mappings
Mean ES units
0 2 4 6
center all SCPs
center
all peripheral spots
peripheral spots with target SCP
Rank: 5
Rank rev.: 235
NW p: 0.003 (n = 300)
Spatial p: 0.003 (n = 300)
Spatial p rev.: 0.003 (n = 300)
Integrator−mediated nuclear snRNA processing
Cytosolic Sm core assembly onto U snRNAs
HK2874
1000 µm
0 2 4 6
Source−target SCP mappings
Mean ES units
0 2 4 6
center all SCPs
center
all peripheral spots
peripheral spots with target SCP
Rank: 28
Rank rev.: 51
NW p: 0.003 (n = 300)
Spatial p: 0.332 (n = 300)
Spatial p rev.: 0.379 (n = 300)

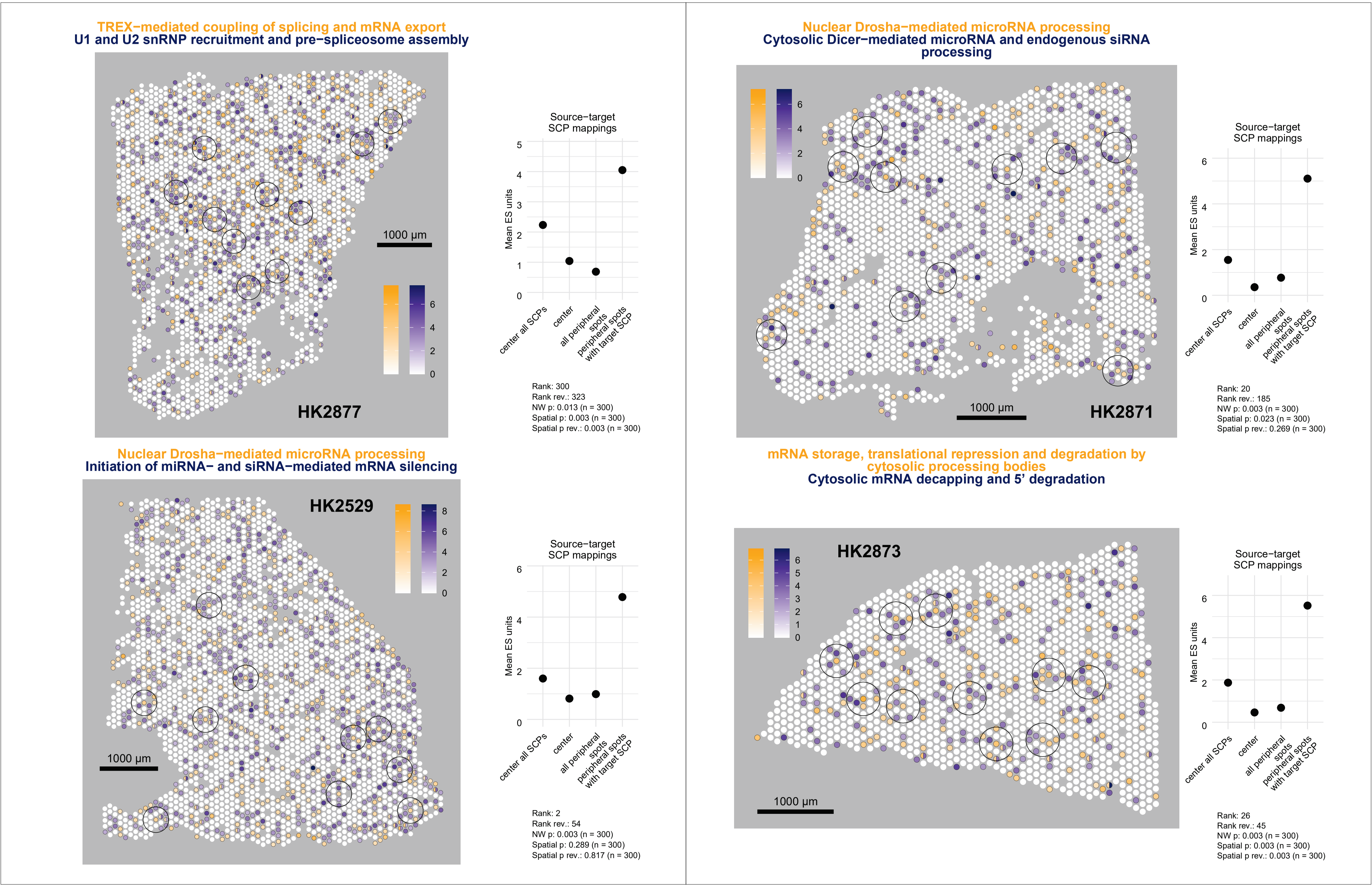
TREX−mediated coupling of splicing and mRNA export
U1 and U2 snRNP recruitment and pre−spliceosome assembly
1000 µm
0
2
4
6
HK2877
Source−target
SCP mappings
Mean ES units
0
1
2
3
4
5
center all SCPs
center
all peripheral spots
peripheral spots with target SCP
Rank: 300
Rank rev.: 323
NW p: 0.013 (n = 300)
Spatial p: 0.003 (n = 300)
Spatial p rev.: 0.003 (n = 300)
Nuclear Drosha−mediated microRNA processing
Cytosolic Dicer−mediated microRNA and endogenous siRNA processing
0
2
4
6
1000 µm
HK2871
Source−target
SCP mappings
Mean ES units
0
2
4
6
center all SCPs
center
all peripheral spots
peripheral spots with target SCP
Rank: 20
Rank rev.: 185
NW p: 0.003 (n = 300)
Spatial p: 0.023 (n = 300)
Spatial p rev.: 0.269 (n = 300)
Nuclear Drosha−mediated microRNA processing
Initiation of miRNA− and siRNA−mediated mRNA silencing
HK2529
0
2
4
6
8
1000 µm
Source−target
SCP mappings
Mean ES units
0
2
4
6
center all SCPs
center
all peripheral spots
peripheral spots with target SCP
Rank: 2
Rank rev.: 54
NW p: 0.003 (n = 300)
Spatial p: 0.289 (n = 300)
Spatial p rev.: 0.817 (n = 300)
mRNA storage, translational repression and degradation by cytosolic processing bodies
Cytosolic mRNA decapping and 5’ degradation
HK2873
0
1
2
3
4
5
6
1000 µm
Source−target
SCP mappings
Mean ES units
0
2
4
6
center all SCPs
center
all peripheral spots
peripheral spots with target SCP
Rank: 26
Rank rev.: 45
NW p: 0.003 (n = 300)
Spatial p: 0.003 (n = 300)
Spatial p rev.: 0.003 (n = 300)

Canonical deadenylation of cytosolic mRNA by CCR4−NOT complex
mRNA storage, translational repression and degradation by cytosolic processing bodies

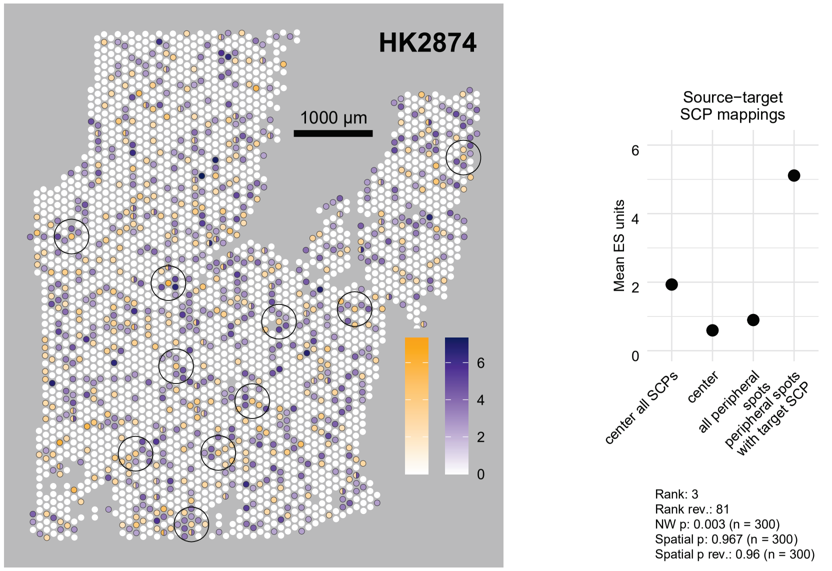


Canonical deadenylation of cytosolic mRNA by CCR4−NOT complex
Cytosolic mRNA decapping and 5’ degradation

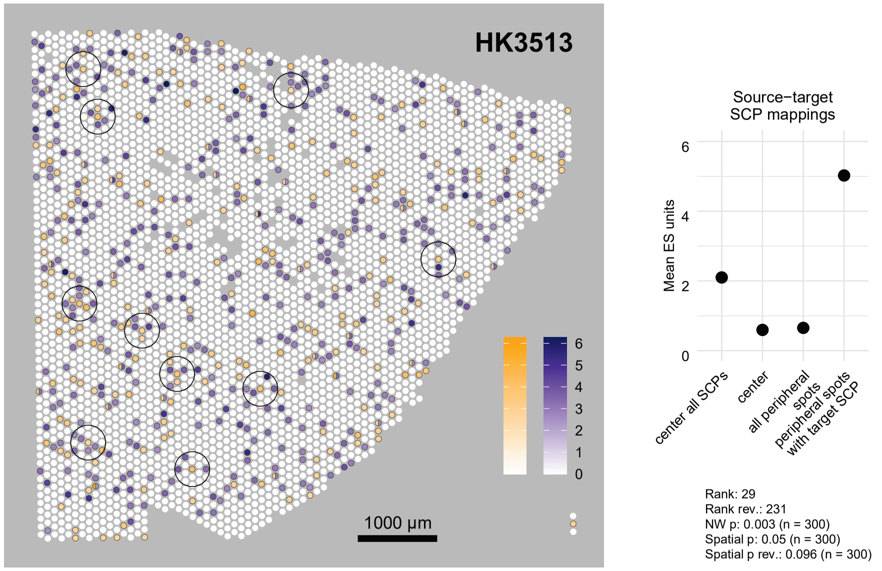


Nucleocytoplasmic protein transport
Nuclear pore complex organization

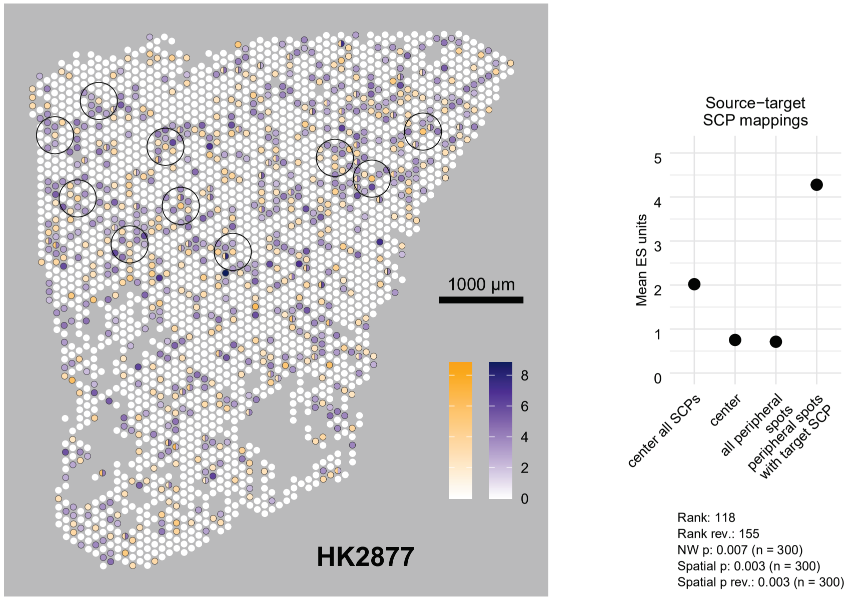


Cytosolic Sm core assembly onto U snRNAs
U5 snRNP−selective component synthesis

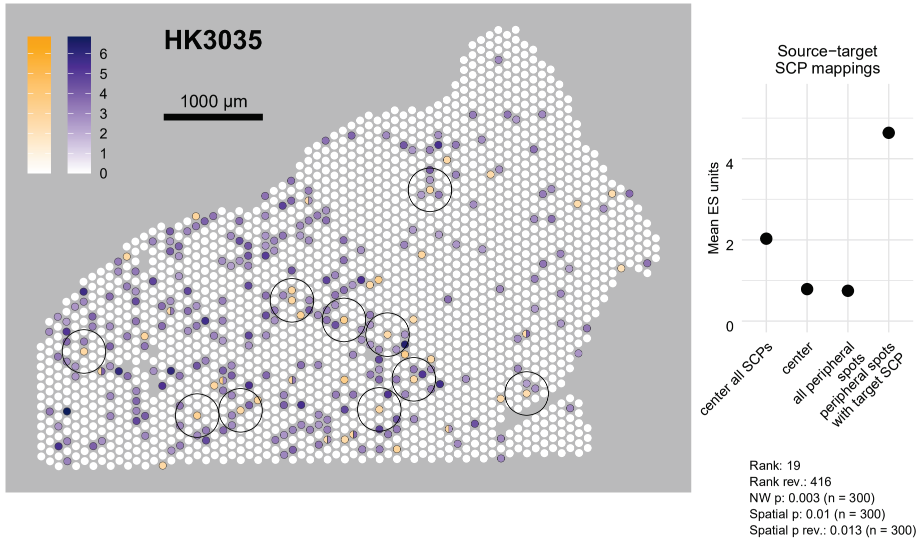

U2 snRNP−selective component synthesis
Catalytic spliceosome activation

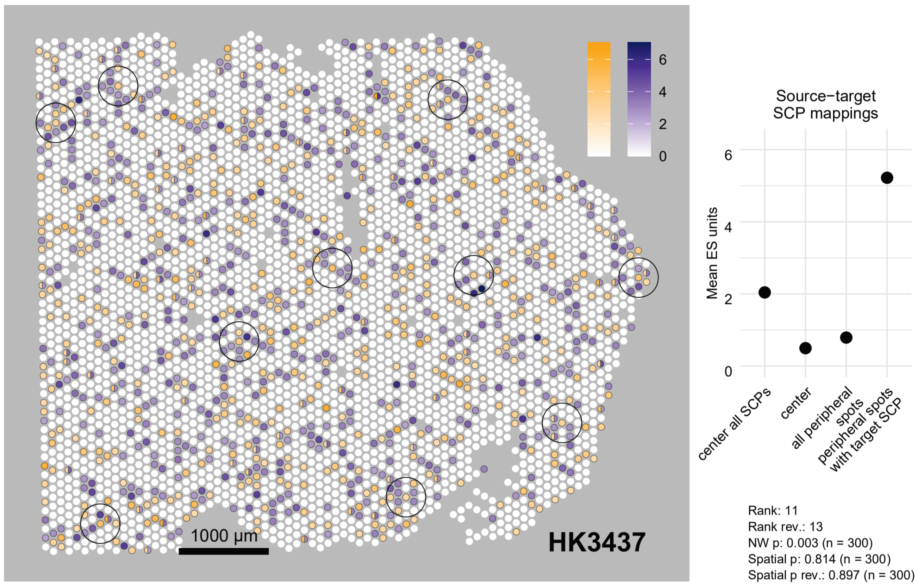


Restriction point
Assembly of core replication fork machinery

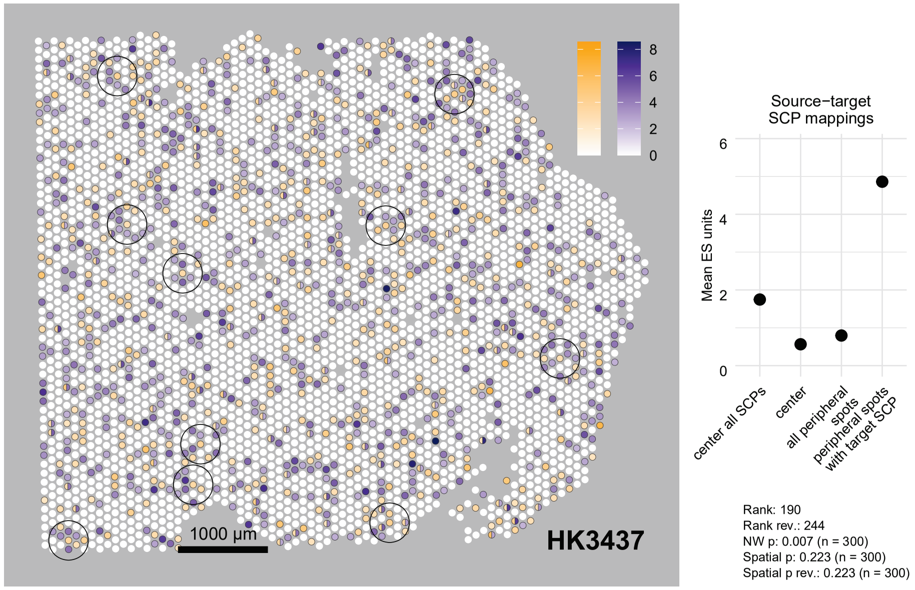


Catalytic spliceosome activation
U5 snRNP−selective component synthesis

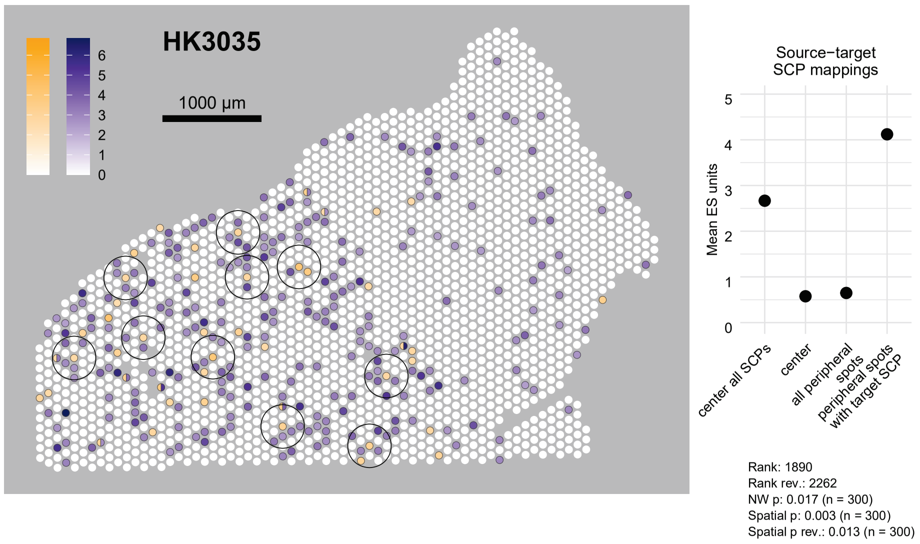


Metaphase to anaphase transition and spindle assembly checkpoint
Eukaryotic kinetochore assembly

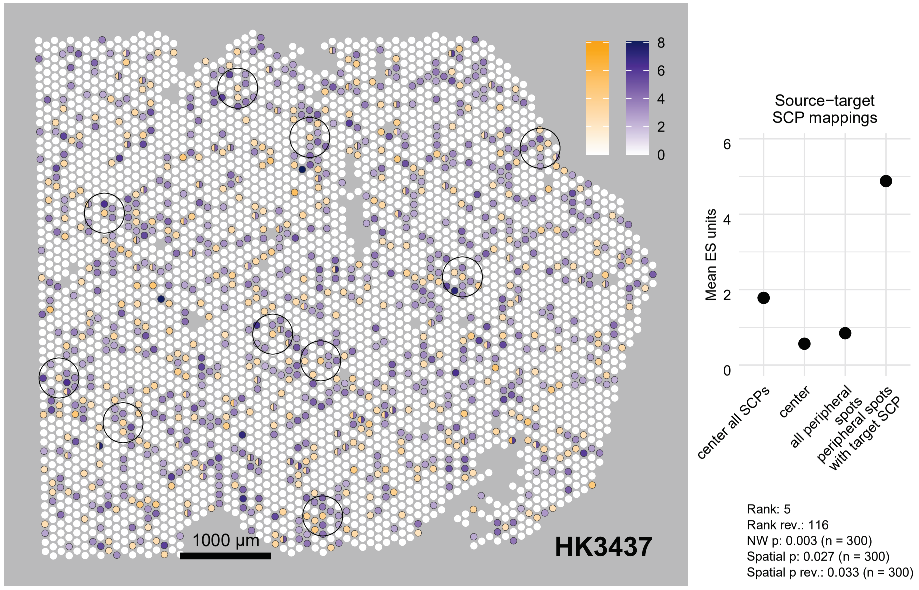

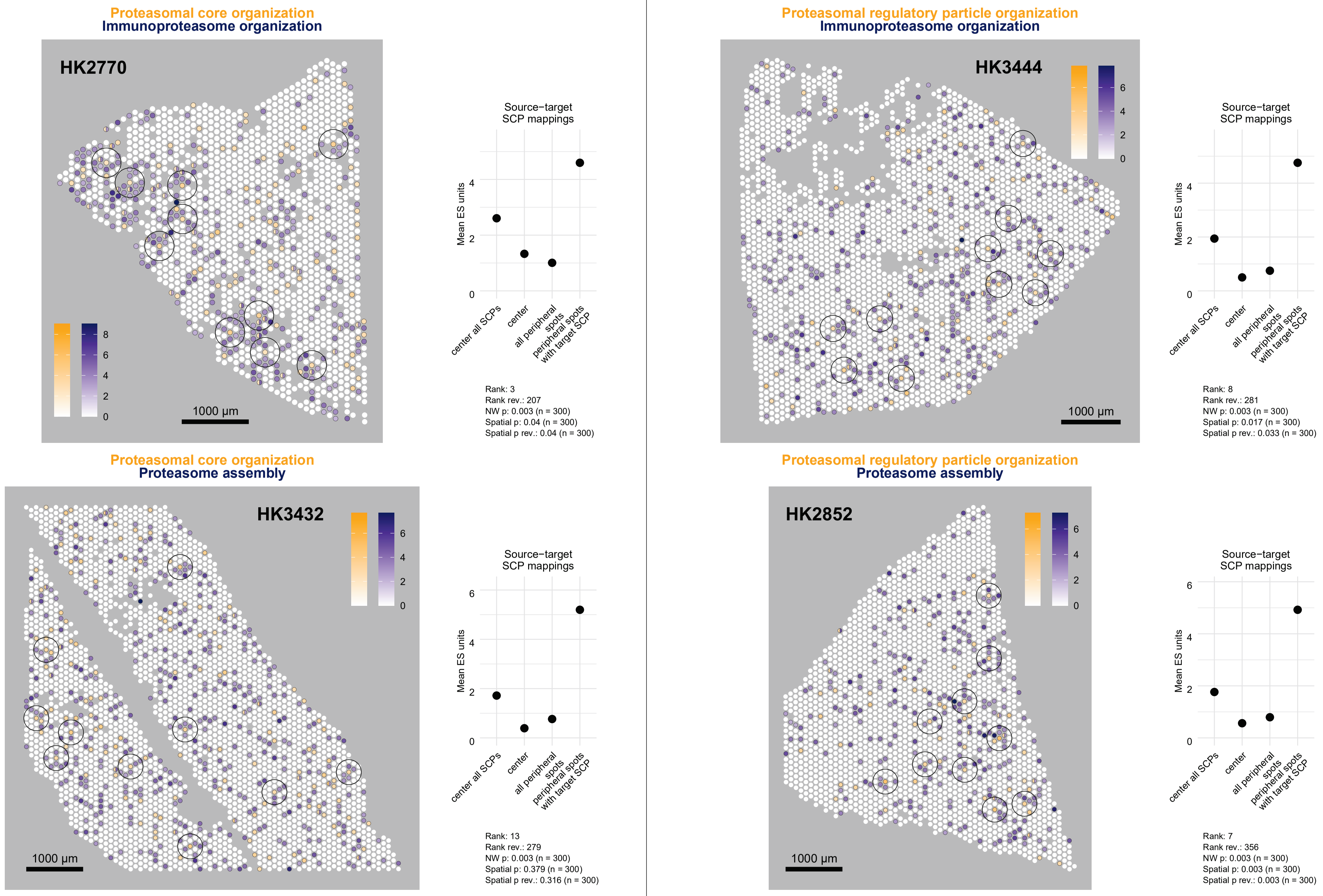
Proteasomal core organization
Immunoproteasome organization
HK2770
1000 µm
Source−target
SCP mappings
Mean ES units
center all SCPs
center
all peripheral spots
peripheral spots with target SCP
Rank: 3
Rank rev.: 207
NW p: 0.003 (n = 300)
Spatial p: 0.04 (n = 300)
Spatial p rev.: 0.04 (n = 300)
Proteasomal regulatory particle organization
Immunoproteasome organization
HK3444
1000 µm
Source−target
SCP mappings
Mean ES units
center all SCPs
center
all peripheral spots
peripheral spots with target SCP
Rank: 8
Rank rev.: 281
NW p: 0.003 (n = 300)
Spatial p: 0.017 (n = 300)
Spatial p rev.: 0.033 (n = 300)
Proteasomal core organization
Proteasome assembly
HK3432
1000 µm
Source−target
SCP mappings
Mean ES units
center all SCPs
center
all peripheral spots
peripheral spots with target SCP
Rank: 13
Rank rev.: 279
NW p: 0.003 (n = 300)
Spatial p: 0.379 (n = 300)
Spatial p rev.: 0.316 (n = 300)
Proteasomal regulatory particle organization
Proteasome assembly
HK2852
1000 µm
Source−target
SCP mappings
Mean ES units
center all SCPs
center
all peripheral spots
peripheral spots with target SCP
Rank: 7
Rank rev.: 356
NW p: 0.003 (n = 300)
Spatial p: 0.003 (n = 300)
Spatial p rev.: 0.003 (n = 300)

**Antigen presentation via MHC class I molecules**
**Immunoproteasome organization**

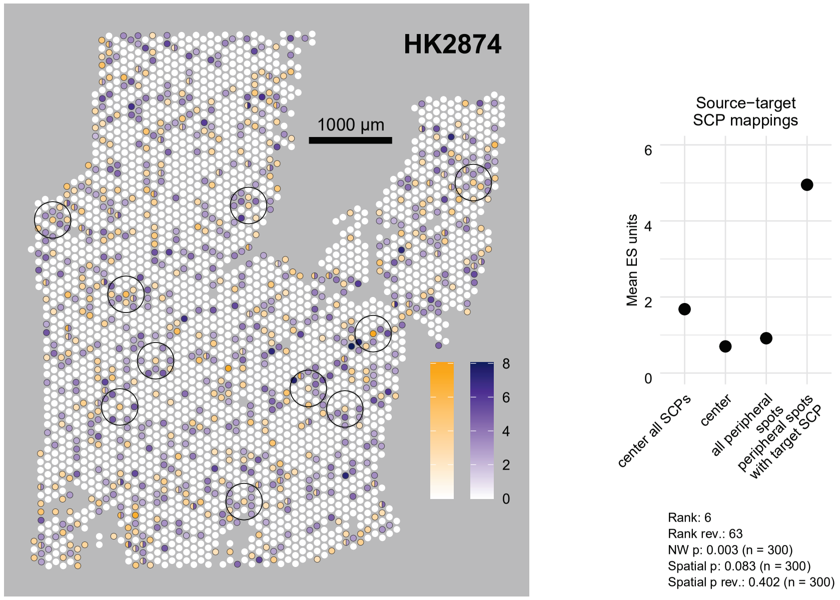


**Regulation of G−protein signaling by beta and gamma subunits**
**GPCR desensitization and endocytic adaptor recruitment**

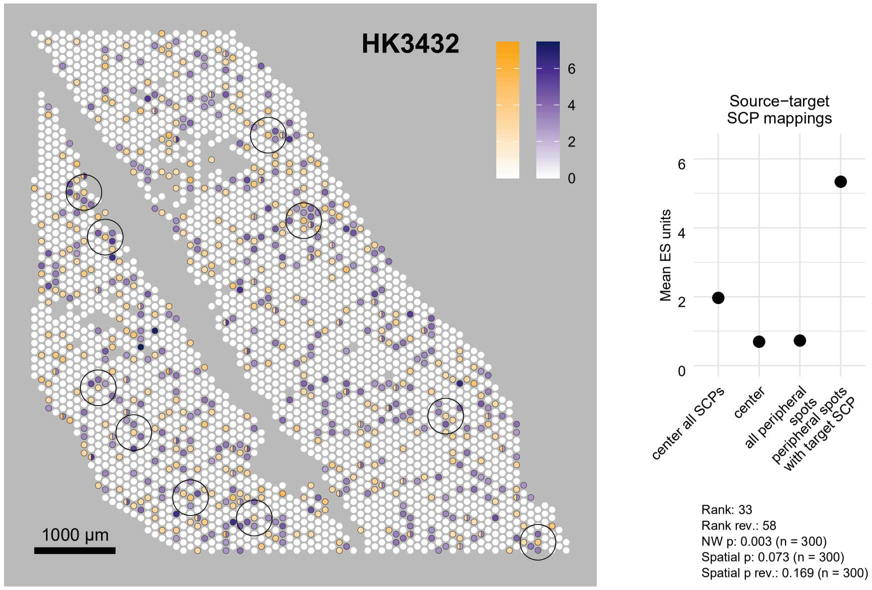


**Ras signaling pathway**
**Fibroblast growth factor receptor signaling**

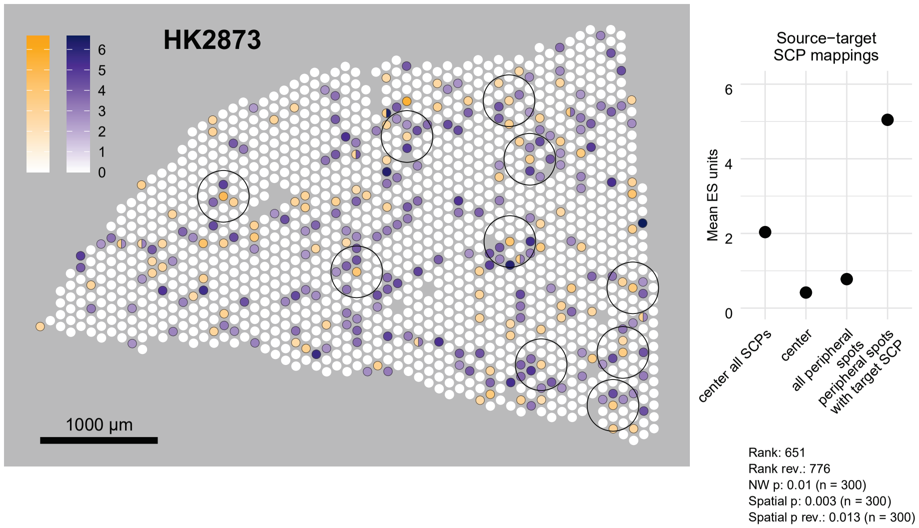


**Cofactor−mediated transcriptional regulation of multiple nuclear receptors**
**Retinoic acid receptor signaling**

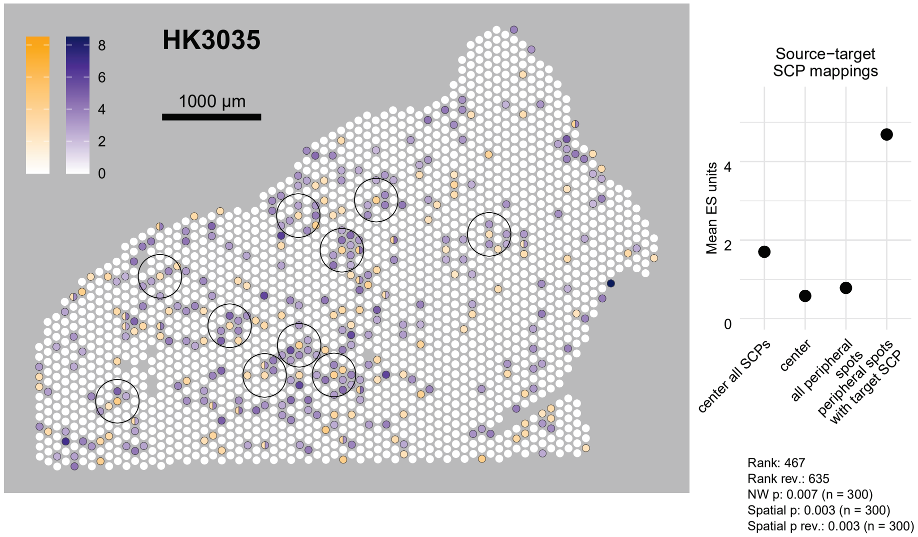

Inhibition of apoptosis
Extrinsic apoptosis pathway

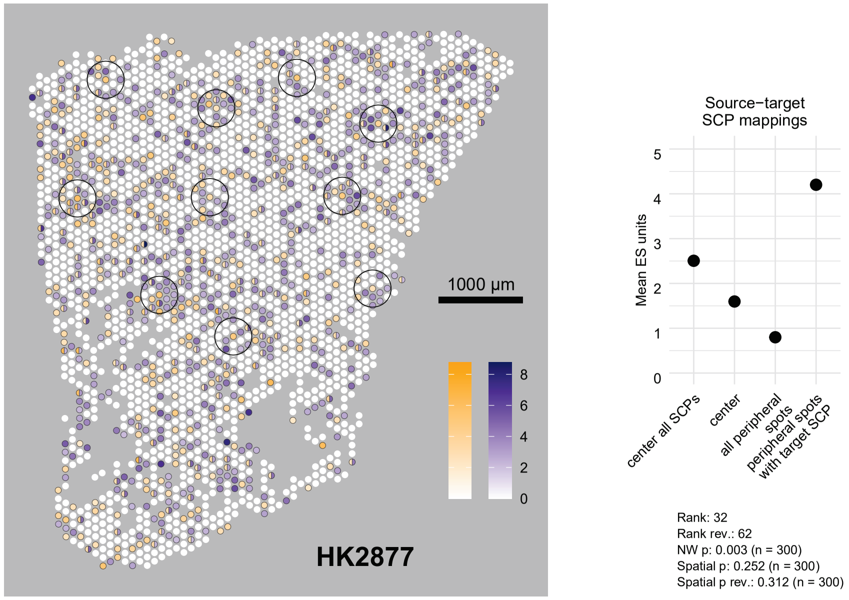


Vesicle biogenesis at ER membrane
Vesicle tethering at cis−Golgi

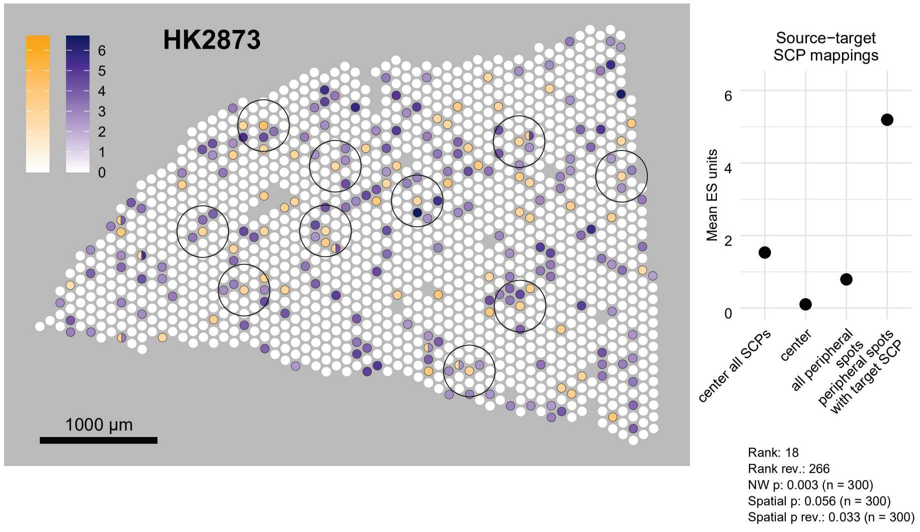


Ceramide biosynthesis
Extrinsic apoptosis pathway

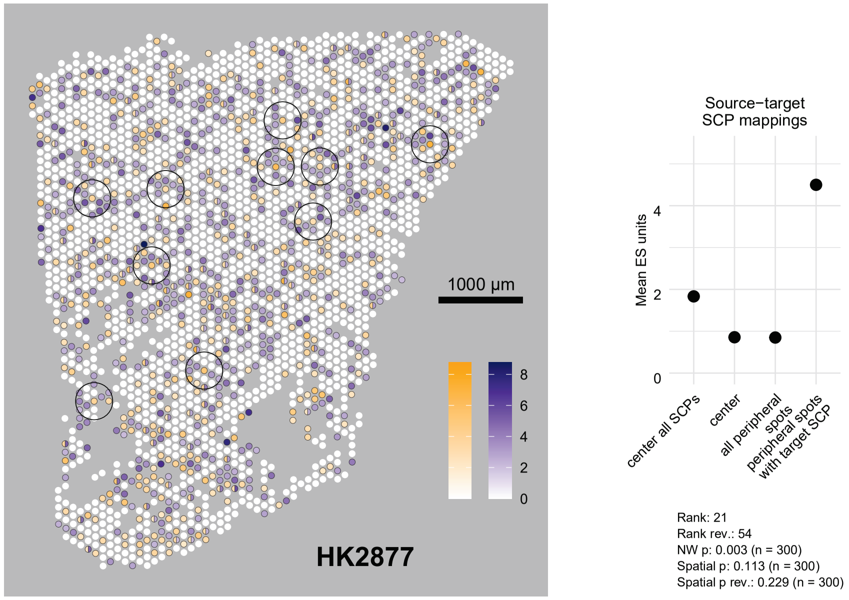


Vesicle biogenesis at cis−Golgi
Vesicle biogenesis at ER membrane

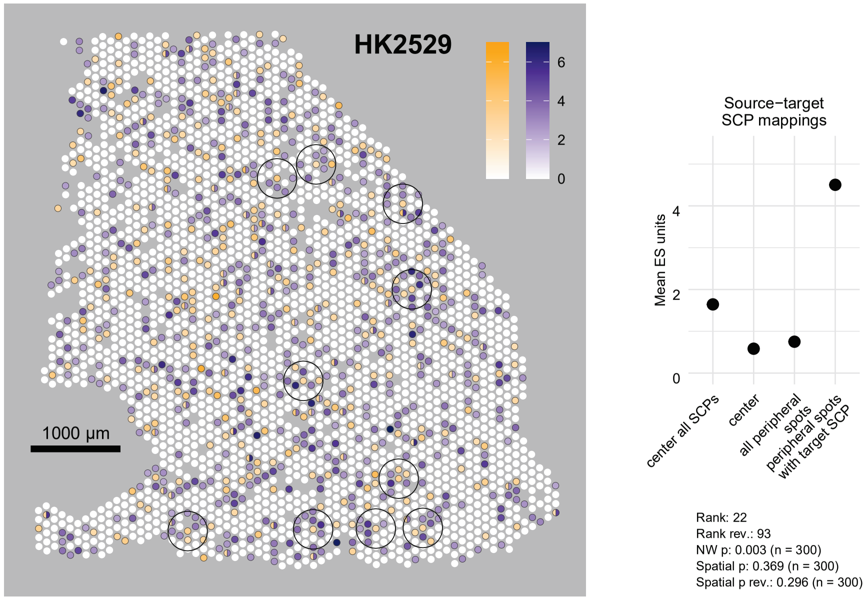

Intra−golgi vesicle tethering
Vesicle tethering at cis−Golgi

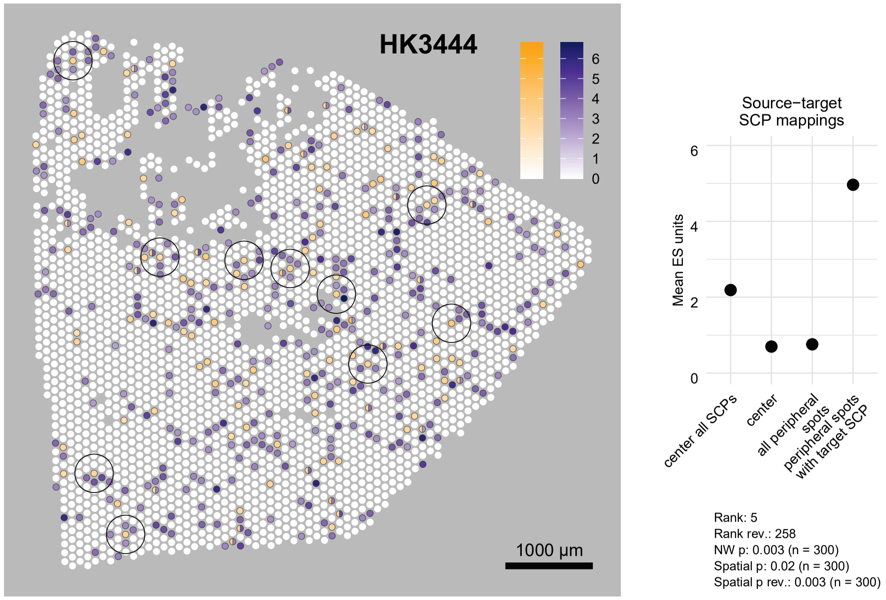


Eukaryotic intraflagellar transport
Mother centriole and distal appendage transformation into basal body

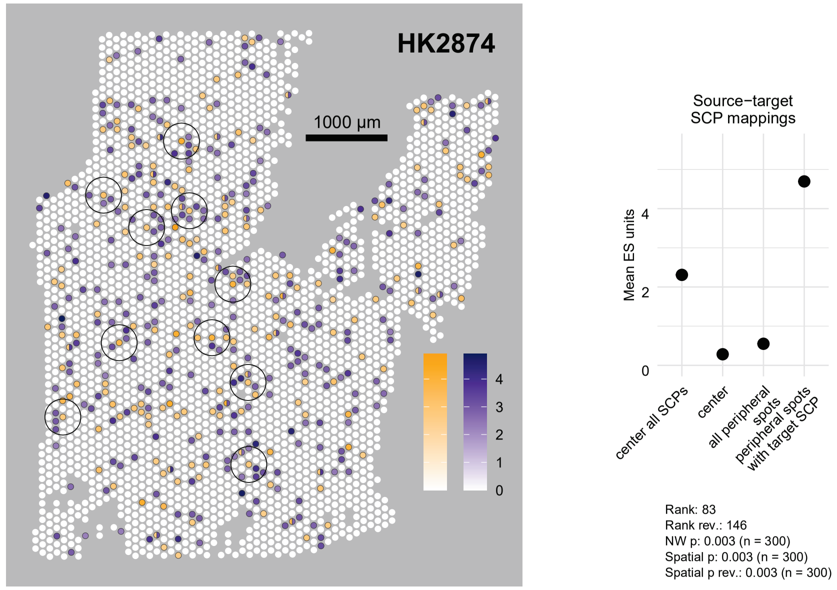


Early endocytic vesicle and endosome tethering
Early endocytic vesicle and endosome fusion

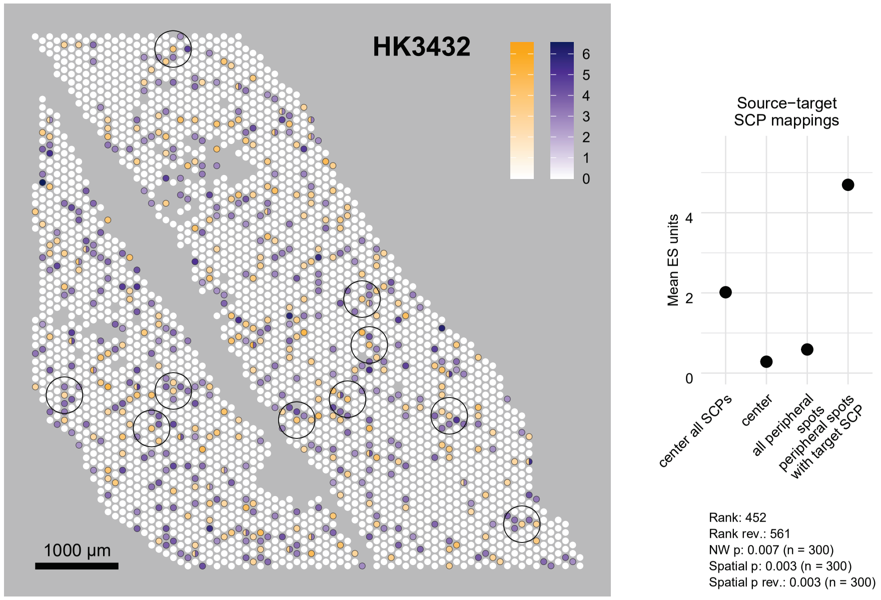


Ciliary transition zone organization
Eukaryotic motile axoneme organization

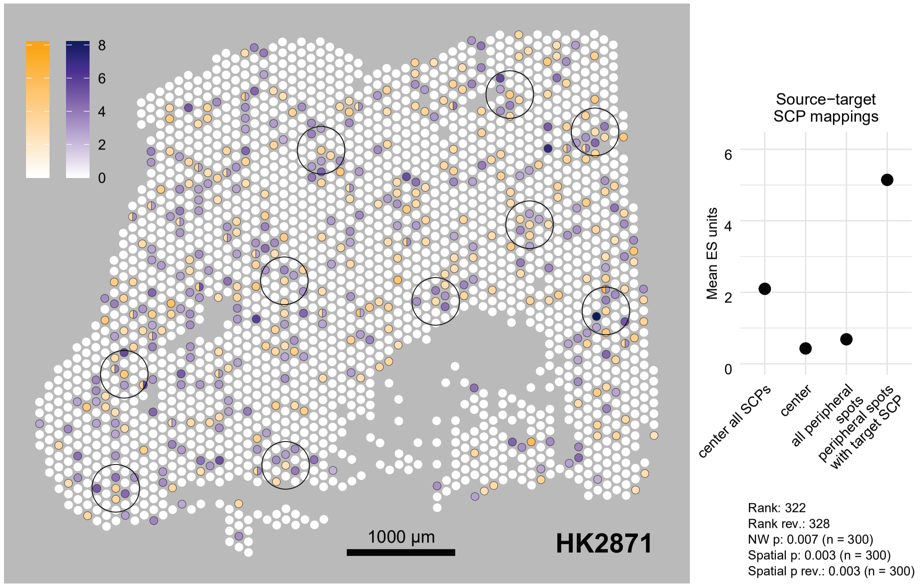

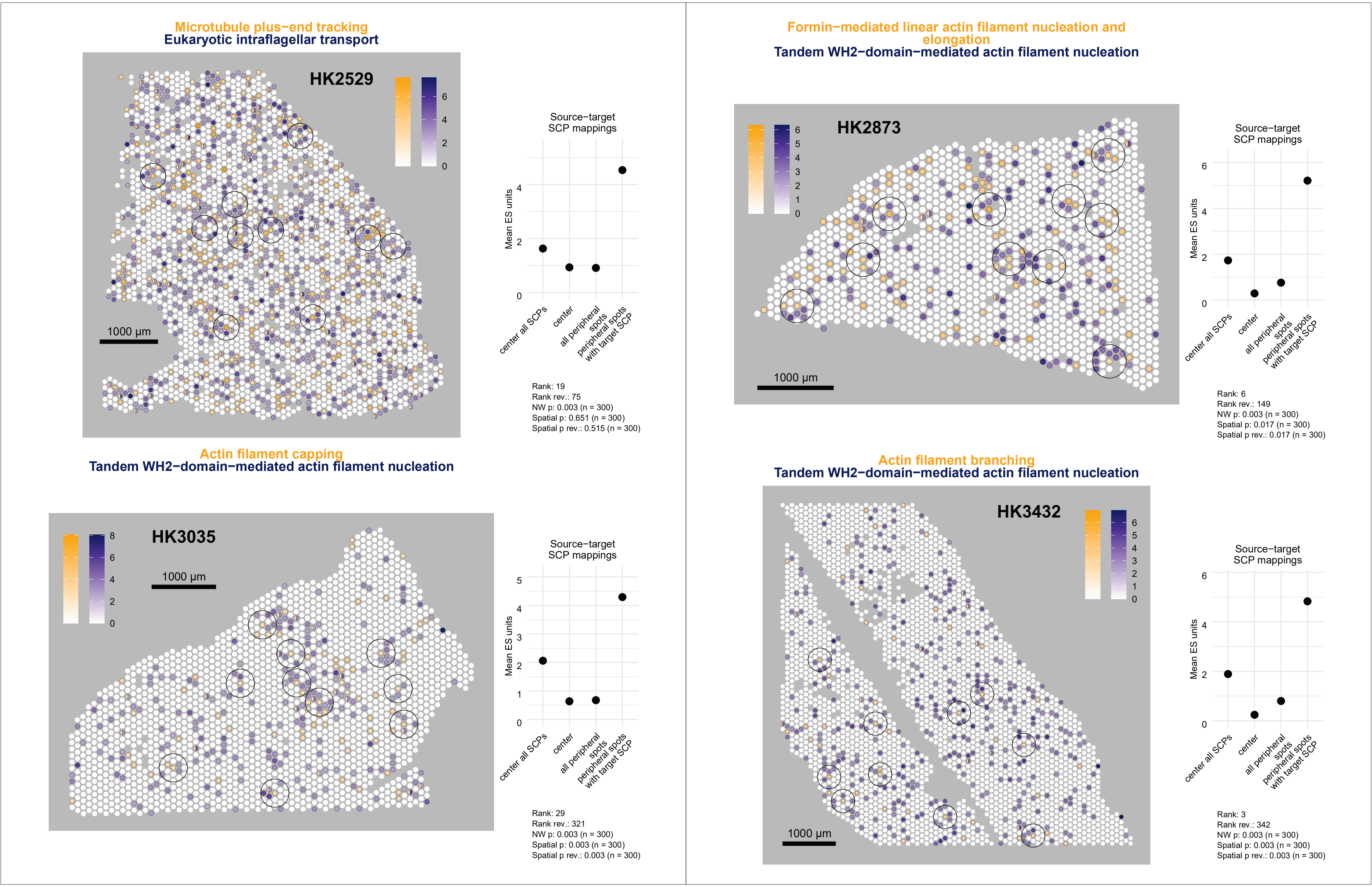

Microtubule plus−end tracking
Eukaryotic intraflagellar transport
HK2529
1000 µm
0 2 4 6
Source−target
SCP mappings
Mean ES units
0 2 4
center all SCPs
center
all peripheral spots
peripheral spots with target SCP
Rank: 19
Rank rev.: 75
NW p: 0.003 (n = 300)
Spatial p: 0.651 (n = 300)
Spatial p rev.: 0.515 (n = 300)
Formin−mediated linear actin filament nucleation and elongation
Tandem WH2−domain−mediated actin filament nucleation
HK2873
1000 µm
0 1 2 3 4 5 6
Source−target
SCP mappings
Mean ES units
0 2 4 6
center all SCPs
center
all peripheral spots
peripheral spots with target SCP
Rank: 6
Rank rev.: 149
NW p: 0.003 (n = 300)
Spatial p: 0.017 (n = 300)
Spatial p rev.: 0.017 (n = 300)
Actin filament capping
Tandem WH2−domain−mediated actin filament nucleation
HK3035
1000 µm
0 2 4 6 8
Source−target
SCP mappings
Mean ES units
0 1 2 3 4 5
center all SCPs
center
all peripheral spots
peripheral spots with target SCP
Rank: 29
Rank rev.: 321
NW p: 0.003 (n = 300)
Spatial p: 0.003 (n = 300)
Spatial p rev.: 0.003 (n = 300)
Actin filament branching
Tandem WH2−domain−mediated actin filament nucleation
HK3432
1000 µm
0 1 2 3 4 5 6
Source−target
SCP mappings
Mean ES units
0 2 4 6
center all SCPs
center
all peripheral spots
peripheral spots with target SCP
Rank: 3
Rank rev.: 342
NW p: 0.003 (n = 300)
Spatial p: 0.003 (n = 300)
Spatial p rev.: 0.003 (n = 300)

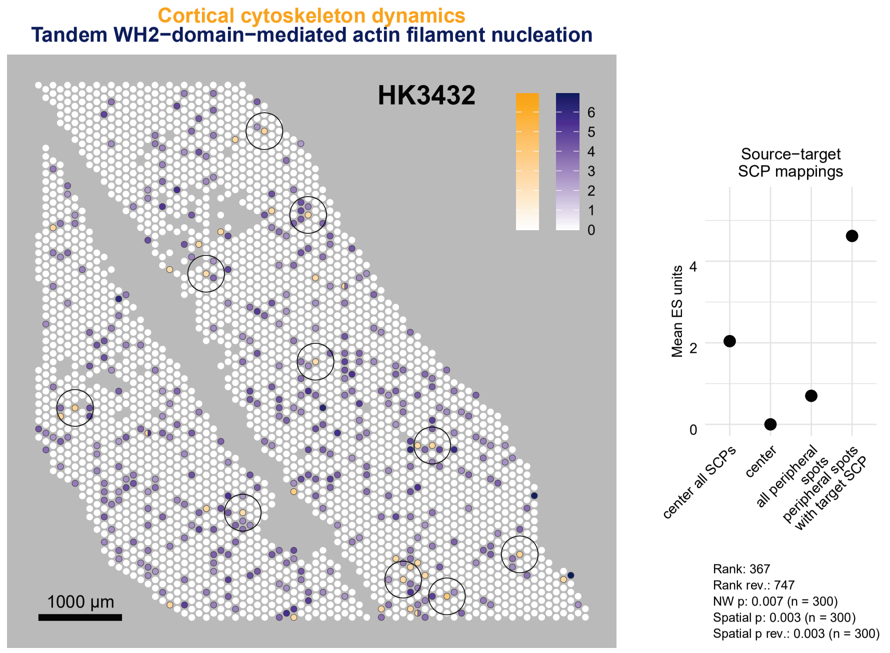

Cortical cytoskeleton dynamics
Tandem WH2−domain−mediated actin filament nucleation
HK3432
1000 µm
Source−target SCP mappings
Mean ES units
center all SCPs
center
all peripheral spots
peripheral spots with target SCP
Rank: 367
Rank rev.: 747
NW p: 0.007 (n = 300)
Spatial p: 0.003 (n = 300)
Spatial p rev.: 0.003 (n = 300)


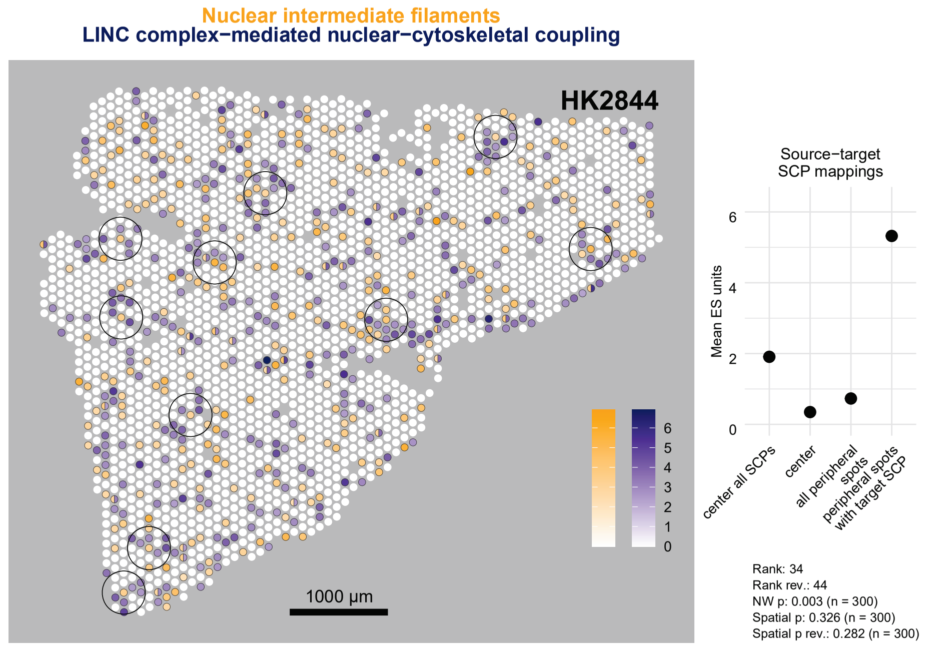

Nuclear intermediate filaments
LINC complex−mediated nuclear−cytoskeletal coupling
HK2844
1000 µm
Source−target SCP mappings
Mean ES units
center all SCPs
center
all peripheral spots
peripheral spots with target SCP
Rank: 34
Rank rev.: 44
NW p: 0.003 (n = 300)
Spatial p: 0.326 (n = 300)
Spatial p rev.: 0.282 (n = 300)


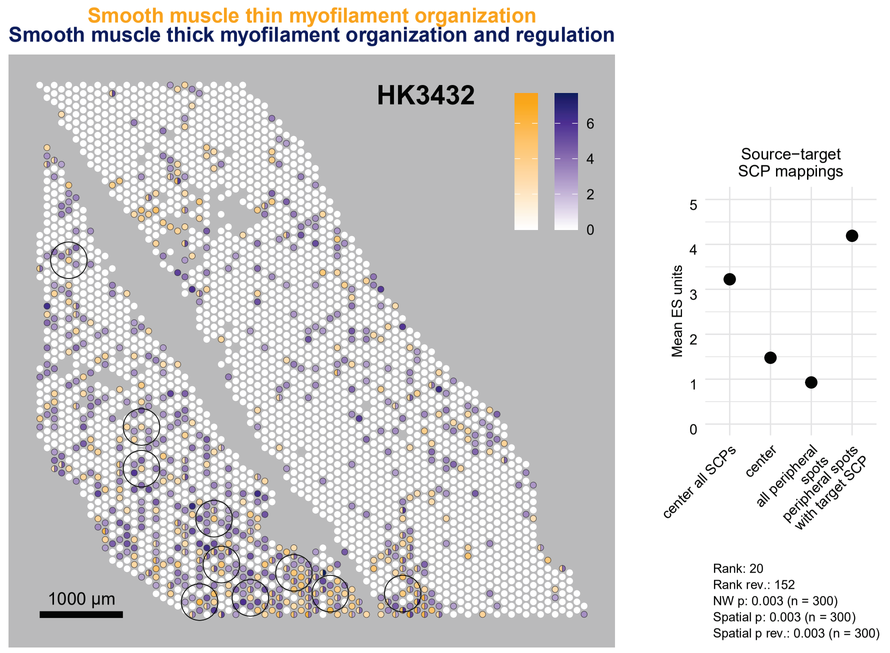

Smooth muscle thin myofilament organization
Smooth muscle thick myofilament organization and regulation
HK3432
1000 µm
Source−target SCP mappings
Mean ES units
center all SCPs
center
all peripheral spots
peripheral spots with target SCP
Rank: 20
Rank rev.: 152
NW p: 0.003 (n = 300)
Spatial p: 0.003 (n = 300)
Spatial p rev.: 0.003 (n = 300)


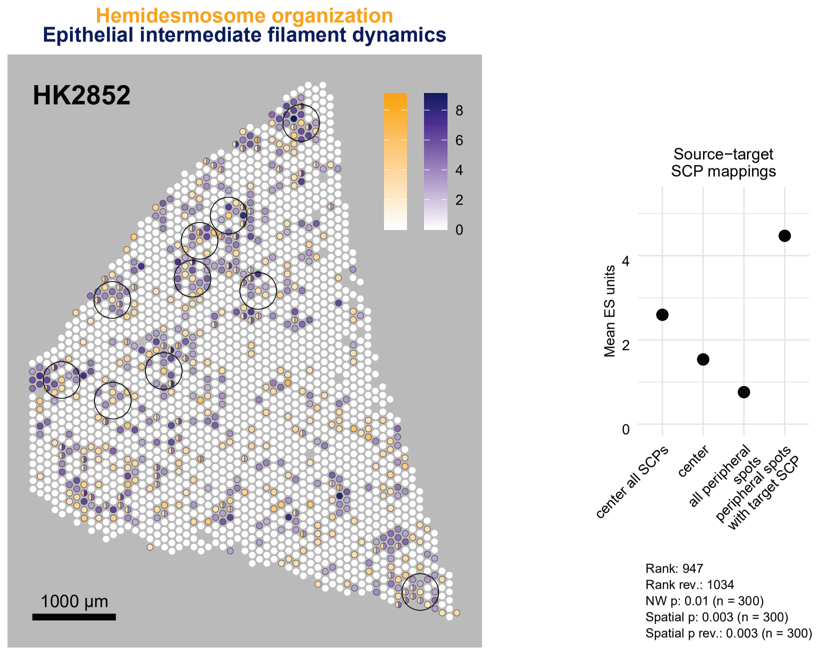

Hemidesmosome organization
Epithelial intermediate filament dynamics
HK2852
1000 µm
Source−target SCP mappings
Mean ES units
center all SCPs
center
all peripheral spots
peripheral spots with target SCP
Rank: 947
Rank rev.: 1034
NW p: 0.01 (n = 300)
Spatial p: 0.003 (n = 300)
Spatial p rev.: 0.003 (n = 300)

**Endoplasmic and sarcoplasmic calcium import, export and storage**
**Endoplasmic reticulum−plasma membrane contact site formation**

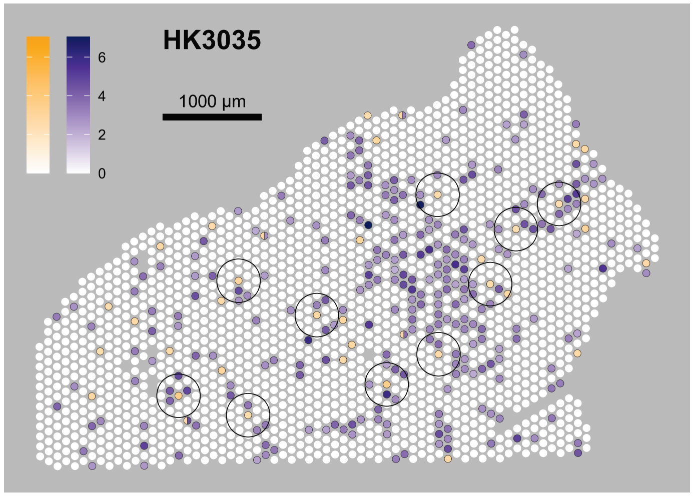


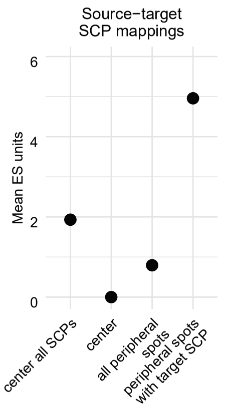


Rank: 11
Rank rev.: 419
NW p: 0.003 (n = 300)
Spatial p: 0.007 (n = 300)
Spatial p rev.: 0.017 (n = 300)

**Supplementary Figure 6. Spatial expression of selected SCP interactions on selected slices. (A)** Results obtained for selected SCP interactions, curated from supplementary figures 5A and B. Numbers show ES scores, numbers in brackets ES score ranks, frames selections for follow-up visualizations, e.g., SCP pair expression on the given slice. Some of the SCP pairs are also shown in figure 6. **(B)** Spatial expression maps of the same SCP pairs in the same order. Orange and blue indicate ES units of the source SCP (center spot) and the target SCP (peripheral spots), respectively. Spots split into orange and blue halves have both SCPs among their top 30 SCPs. Circles frame sites with the highest mapped ES score units. Diagrams show mapped ES units between center and peripheral spots, averaged either across all peripheral spots or only across peripheral spots expressing the target SCP among their top 30 SCPs. These are compared with the potential mapping of the source to the target SCP within the center spot, which indicates whether a source-target pair depends on neighboring spots or is already present in the center spot. Potential mappings were calculated as the smaller of the two SCPs' ES units in the center spot, i.e., the number of ES units that would be mapped if all ES units of both SCPs were mapped to each other. Since only the top 30 SCPs per spot are considered, we additionally calculated potential mapping with all SCPs expressed in the center. Expression maps and diagrams of five selected SCPs are also shown in figure 7.

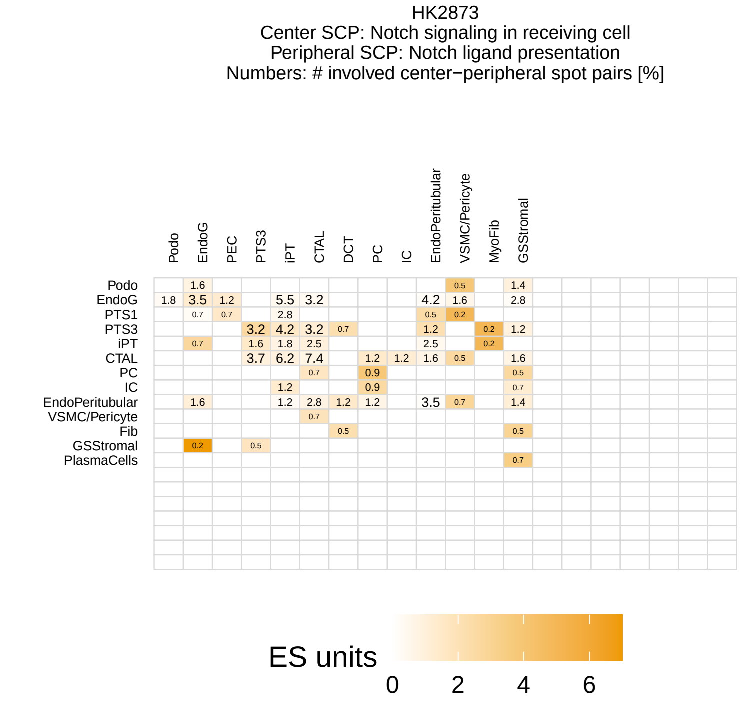
HK2873
Center SCP: Notch signaling in receiving cell
Peripheral SCP: Notch ligand presentation
Numbers: # involved center−peripheral spot pairs [%]
Podo
EndoG
PEC
PTS3
iPT
CTAL
DCT
PC
IC
EndoPeritubular
VSMC/Pericyte
MyoFib
GSStromal
PTS1
Fib
PlasmaCells
ES units
0
2
4
6

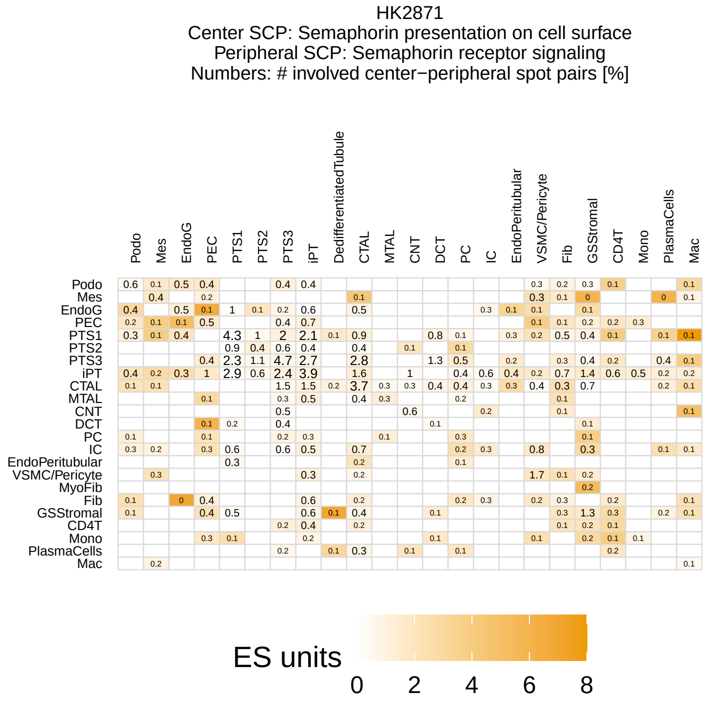
HK2871
Center SCP: Semaphorin presentation on cell surface
Peripheral SCP: Semaphorin receptor signaling
Numbers: # involved center−peripheral spot pairs [%]
Podo
Mes
EndoG
PEC
PTS1
PTS2
PTS3
iPT
DedifferentiatedTubule
CTAL
MTAL
CNT
DCT
PC
IC
EndoPeritubular
VSMC/Pericyte
MyoFib
Fib
GSStromal
CD4T
Mono
PlasmaCells
Mac
ES units
0
2
4
6
8

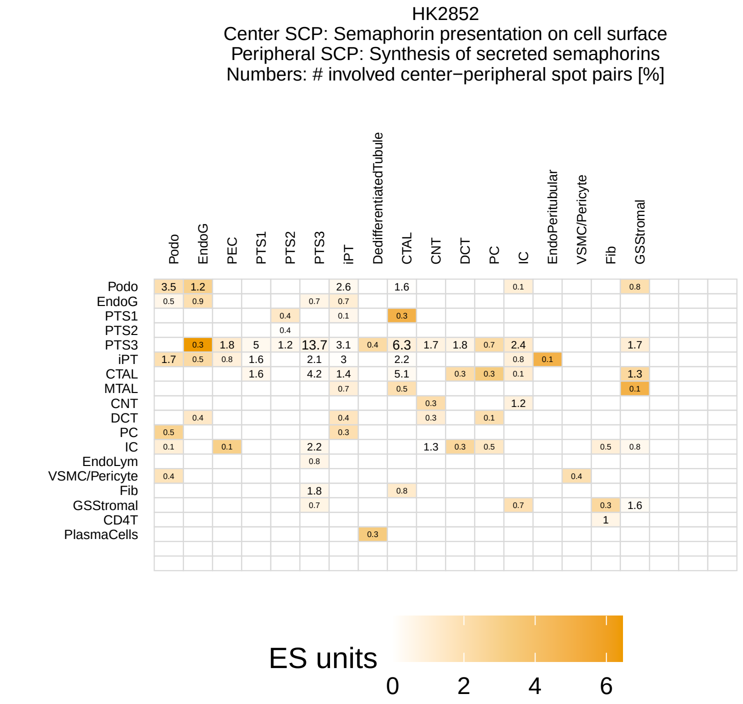
HK2852
Center SCP: Semaphorin presentation on cell surface
Peripheral SCP: Synthesis of secreted semaphorins
Numbers: # involved center−peripheral spot pairs [%]
Podo
EndoG
PEC
PTS1
PTS2
PTS3
iPT
DedifferentiatedTubule
CTAL
CNT
DCT
PC
IC
EndoPeritubular
VSMC/Pericyte
Fib
GSStromal
MTAL
EndoLym
CD4T
PlasmaCells
ES units
0
2
4
6

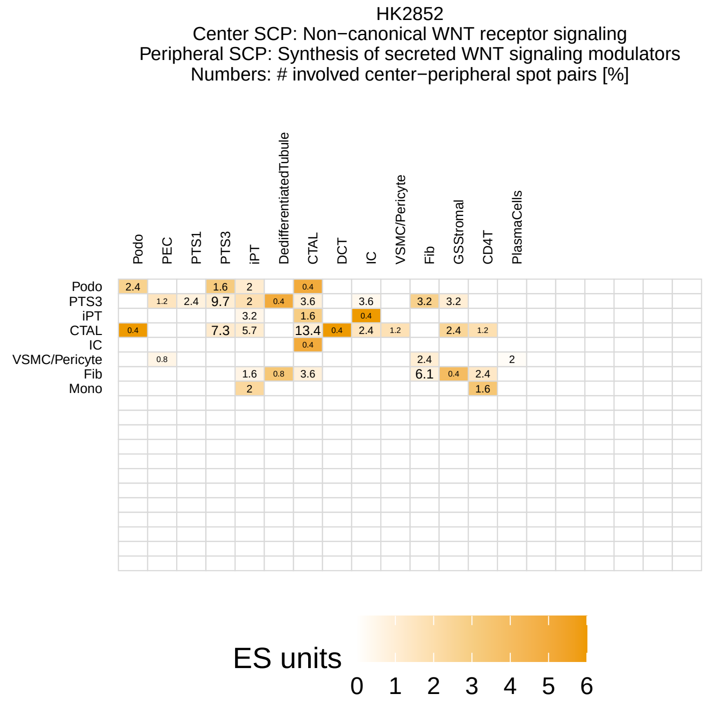
HK2852
Center SCP: Non−canonical WNT receptor signaling
Peripheral SCP: Synthesis of secreted WNT signaling modulators
Numbers: # involved center−peripheral spot pairs [%]
Podo
PEC
PTS1
PTS3
iPT
DedifferentiatedTubule
CTAL
DCT
IC
VSMC/Pericyte
Fib
GSStromal
CD4T
PlasmaCells
Mono
ES units
0
1
2
3
4
5
6

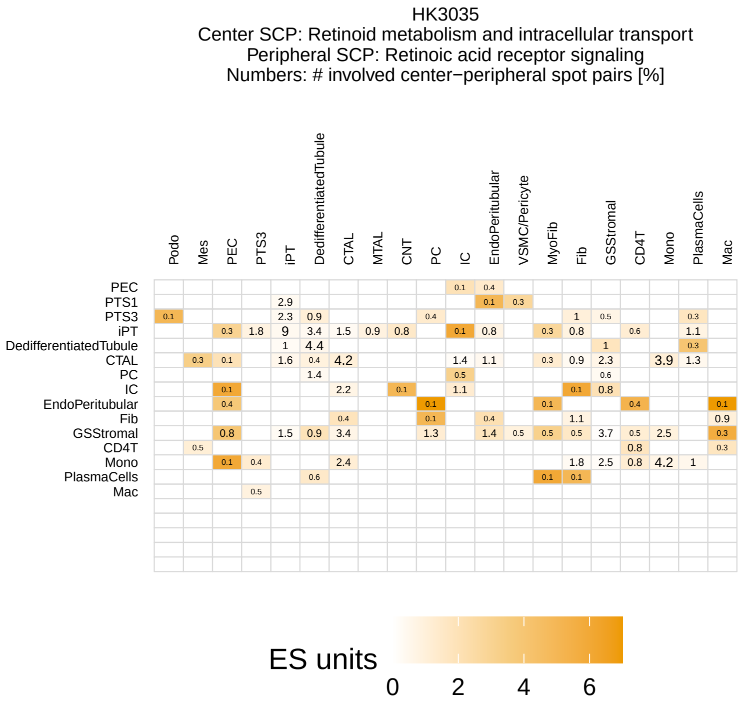
HK3035
Center SCP: Retinoid metabolism and intracellular transport
Peripheral SCP: Retinoic acid receptor signaling
Numbers: # involved center−peripheral spot pairs [%]
Podo
Mes
PEC
PTS3
iPT
DedifferentiatedTubule
CTAL
MTAL
CNT
PC
IC
EndoPeritubular
VSMC/Pericyte
MyoFib
Fib
GSStromal
CD4T
Mono
PlasmaCells
Mac
PTS1
ES units
0
2
4
6

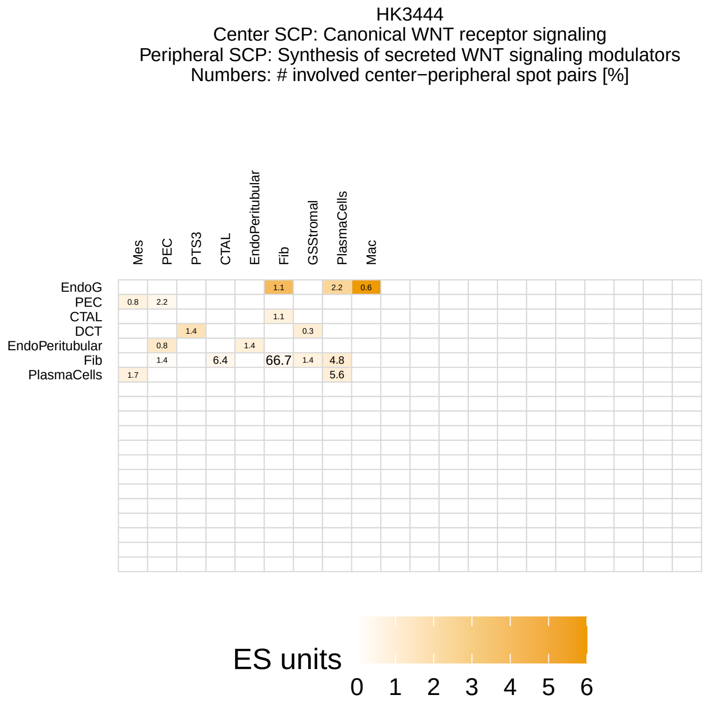
HK3444
Center SCP: Canonical WNT receptor signaling
Peripheral SCP: Synthesis of secreted WNT signaling modulators
Numbers: # involved center−peripheral spot pairs [%]
Mes
PEC
PTS3
CTAL
EndoPeritubular
Fib
GSStromal
PlasmaCells
Mac
EndoG
DCT
ES units
0
1
2
3
4
5
6

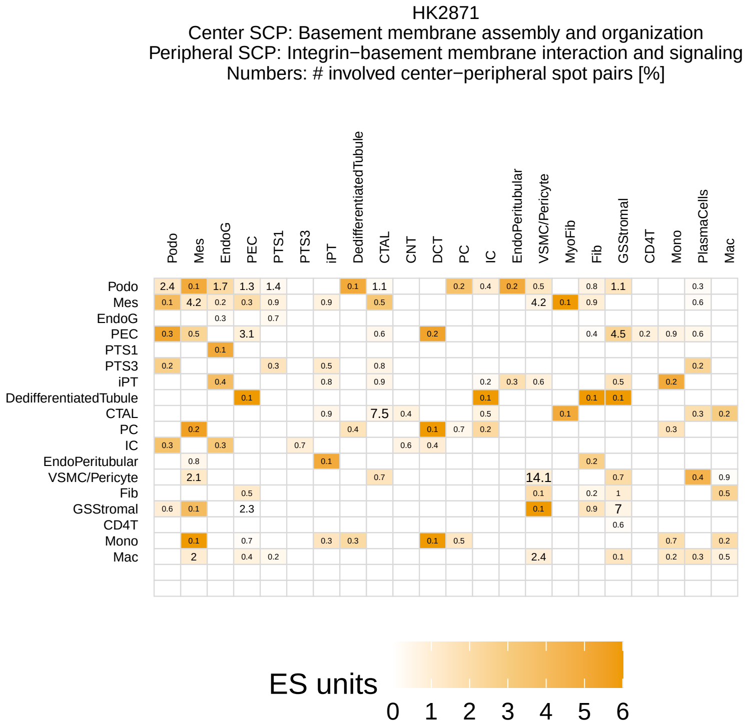
HK2871
Center SCP: Basement membrane assembly and organization
Peripheral SCP: Integrin−basement membrane interaction and signaling
Numbers: # involved center−peripheral spot pairs [%]
ES units
0 1 2 3 4 5 6

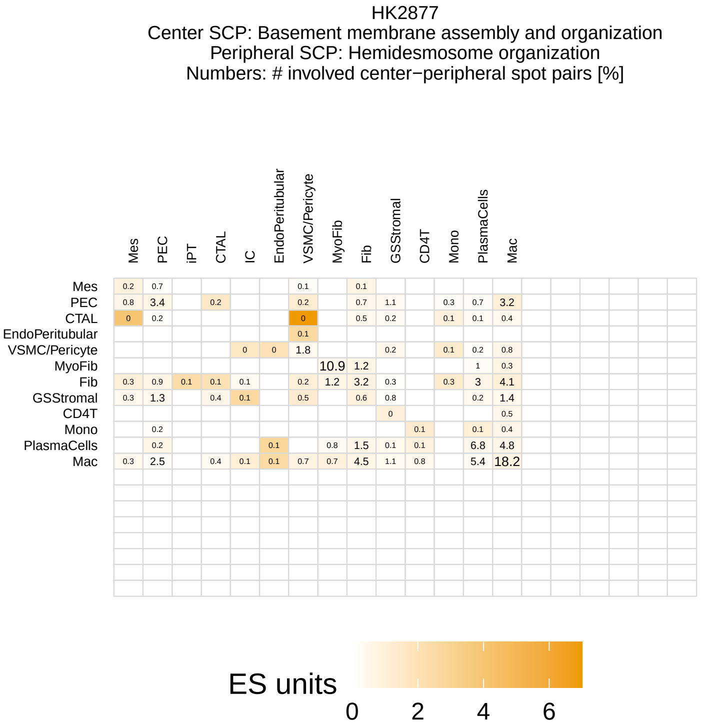
HK2877
Center SCP: Basement membrane assembly and organization
Peripheral SCP: Hemidesmosome organization
Numbers: # involved center−peripheral spot pairs [%]
ES units
0 2 4 6

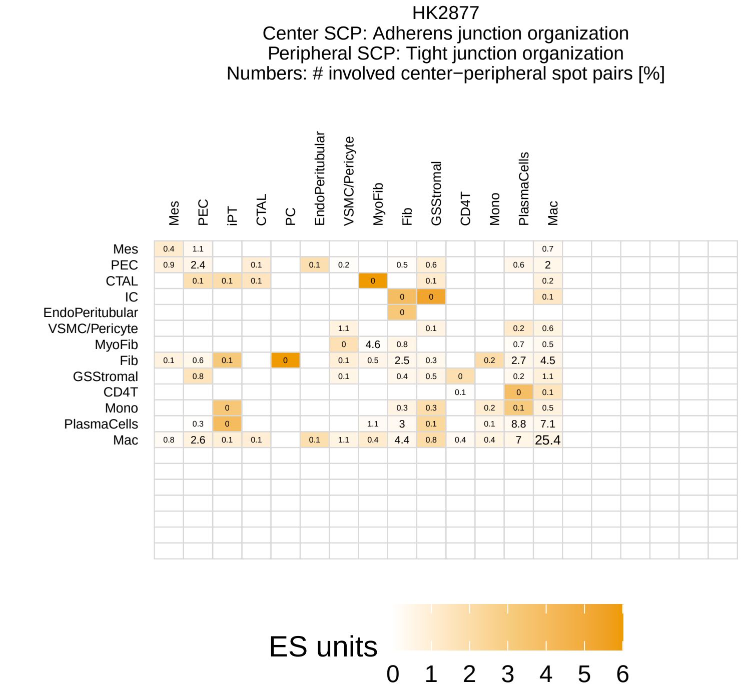
HK2877
Center SCP: Adherens junction organization
Peripheral SCP: Tight junction organization
Numbers: # involved center−peripheral spot pairs [%]
ES units
0 1 2 3 4 5 6

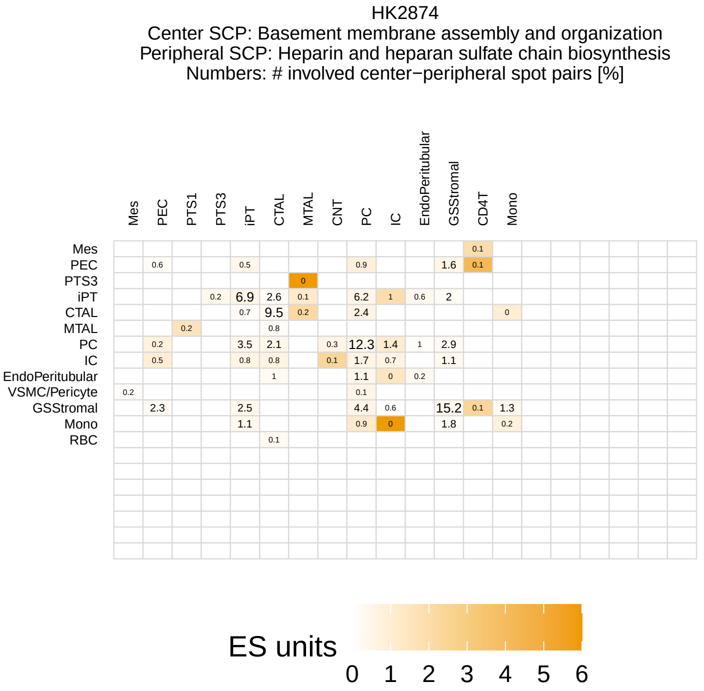
HK2874
Center SCP: Basement membrane assembly and organization
Peripheral SCP: Heparin and heparan sulfate chain biosynthesis
Numbers: # involved center−peripheral spot pairs [%]
ES units
0 1 2 3 4 5 6

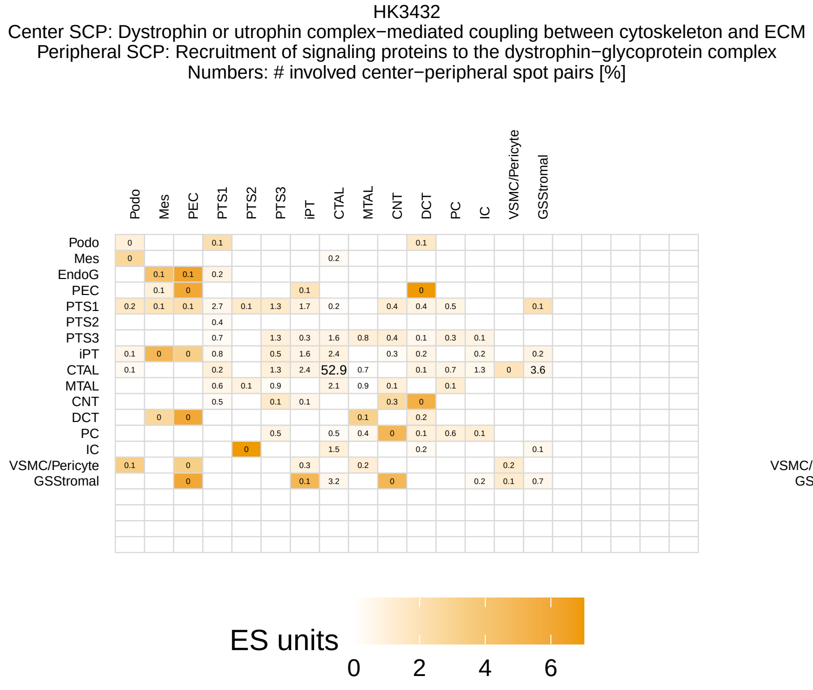
HK3432
Center SCP: Dystrophin or utrophin complex−mediated coupling between cytoskeleton and ECM
Peripheral SCP: Recruitment of signaling proteins to the dystrophin−glycoprotein complex
Numbers: # involved center−peripheral spot pairs [%]
ES units
0 2 4 6

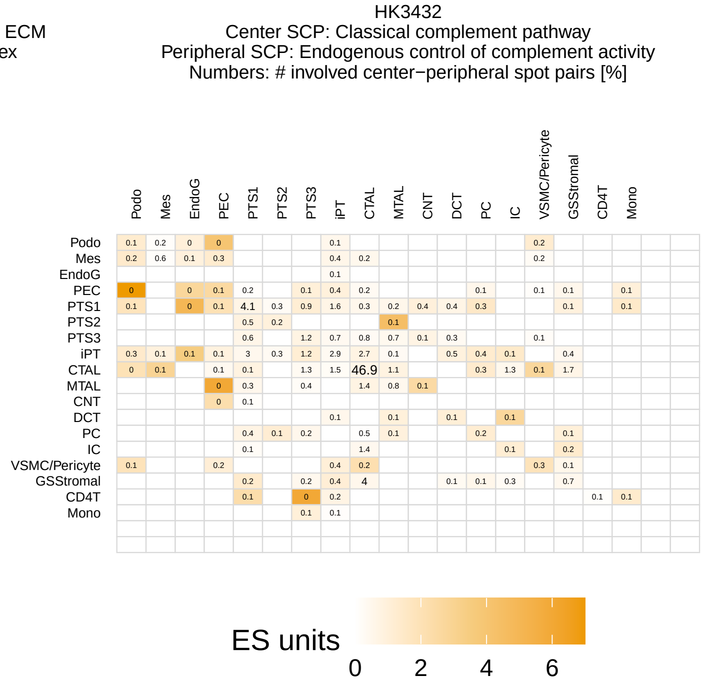
HK3432
Center SCP: Classical complement pathway
Peripheral SCP: Endogenous control of complement activity
Numbers: # involved center−peripheral spot pairs [%]
ES units
0 2 4 6

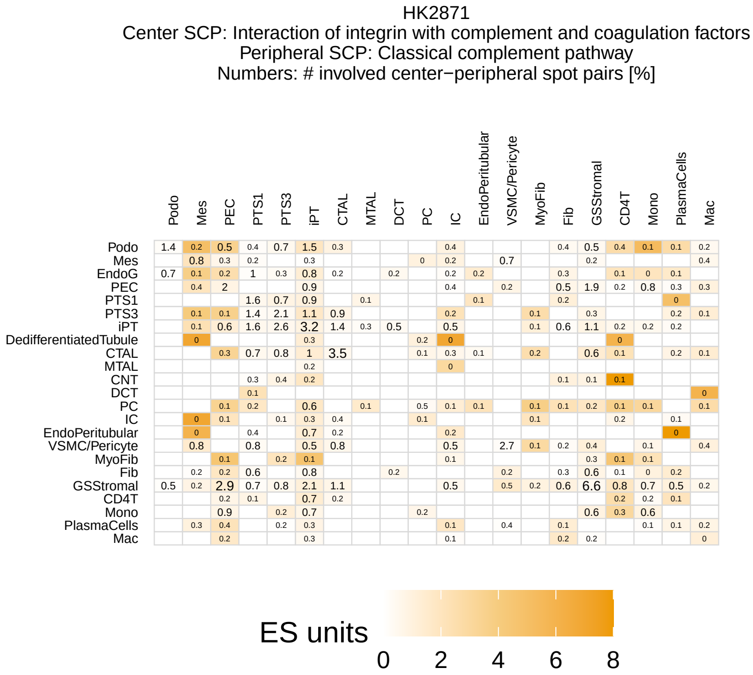
HK2871
Center SCP: Interaction of integrin with complement and coagulation factors
Peripheral SCP: Classical complement pathway
Numbers: # involved center−peripheral spot pairs [%]
ES units
0 2 4 6 8

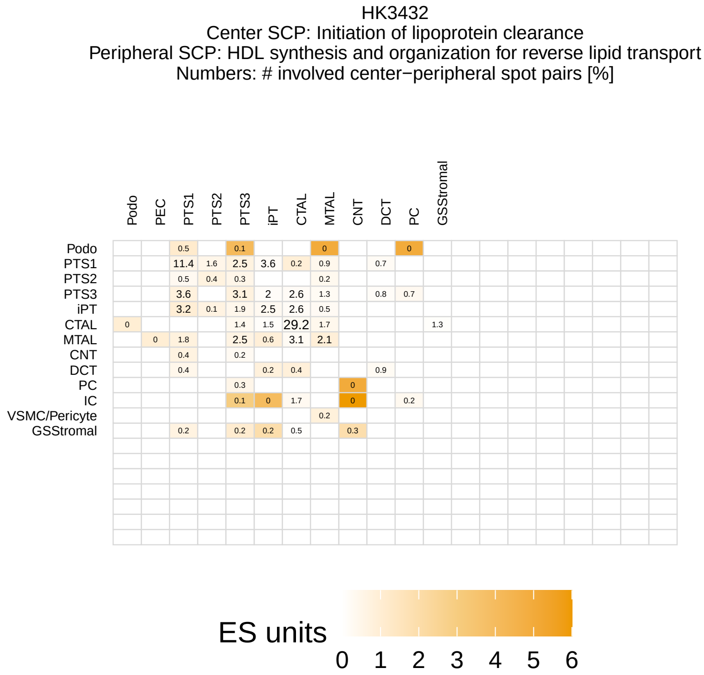
HK3432
Center SCP: Initiation of lipoprotein clearance
Peripheral SCP: HDL synthesis and organization for reverse lipid transport
Numbers: # involved center−peripheral spot pairs [%]
ES units
0 1 2 3 4 5 6

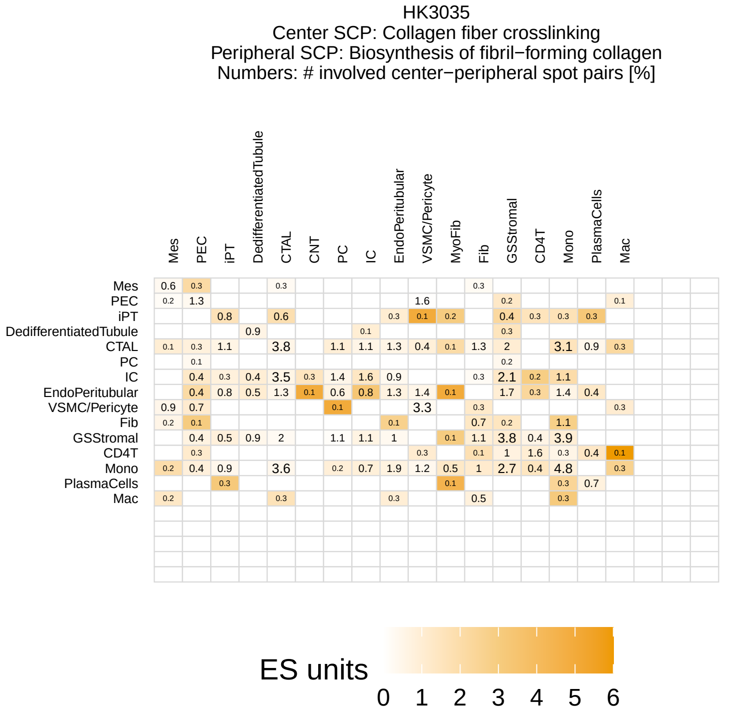
HK3035
Center SCP: Collagen fiber crosslinking
Peripheral SCP: Biosynthesis of fibril−forming collagen
Numbers: # involved center−peripheral spot pairs [%]
ES units
0 1 2 3 4 5 6

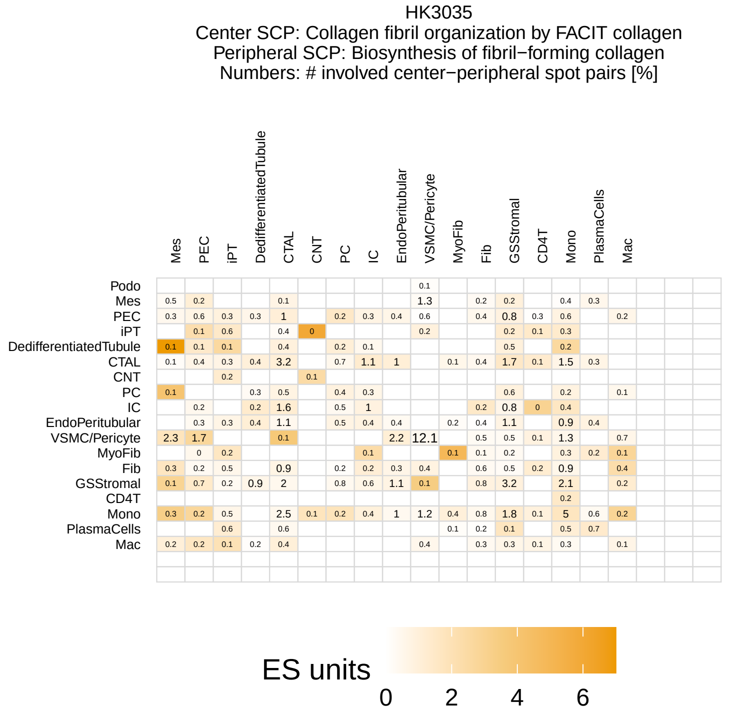
HK3035
Center SCP: Collagen fibril organization by FACIT collagen
Peripheral SCP: Biosynthesis of fibril−forming collagen
Numbers: # involved center−peripheral spot pairs [%]
ES units
0 2 4 6

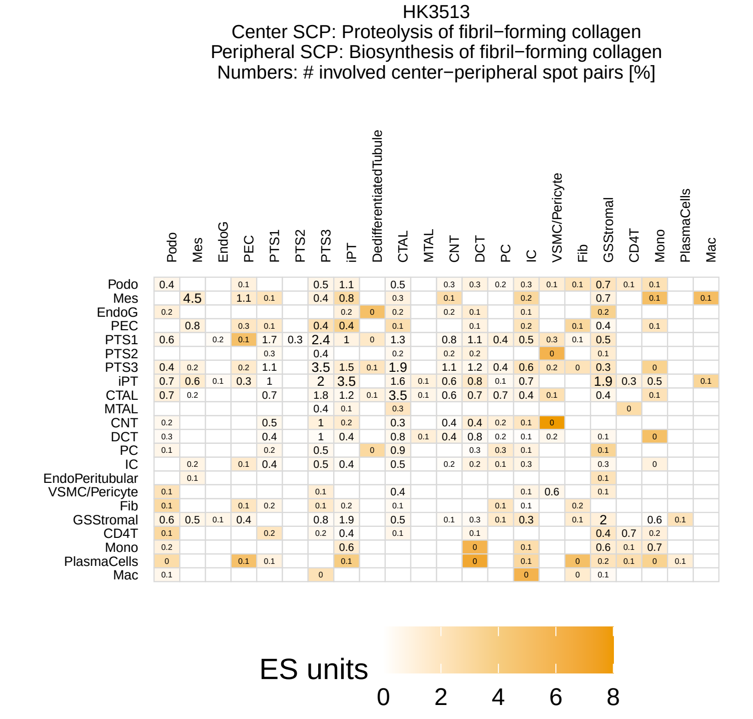
HK3513
Center SCP: Proteolysis of fibril−forming collagen
Peripheral SCP: Biosynthesis of fibril−forming collagen
Numbers: # involved center−peripheral spot pairs [%]
ES units
0 2 4 6 8

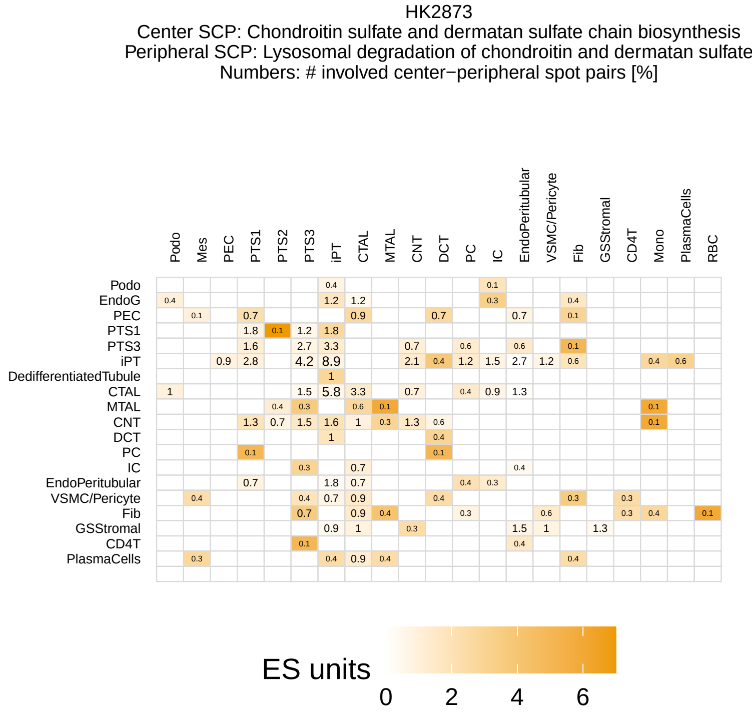
HK2873
Center SCP: Chondroitin sulfate and dermatan sulfate chain biosynthesis
Peripheral SCP: Lysosomal degradation of chondroitin and dermatan sulfate
Numbers: # involved center−peripheral spot pairs [%]
ES units
0 2 4 6

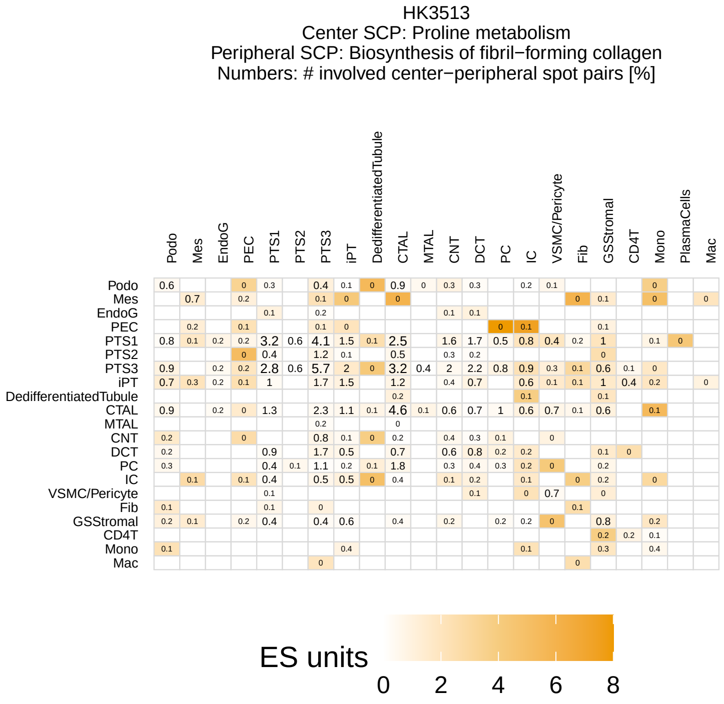
HK3513
Center SCP: Proline metabolism
Peripheral SCP: Biosynthesis of fibril−forming collagen
Numbers: # involved center−peripheral spot pairs [%]
ES units
0 2 4 6 8

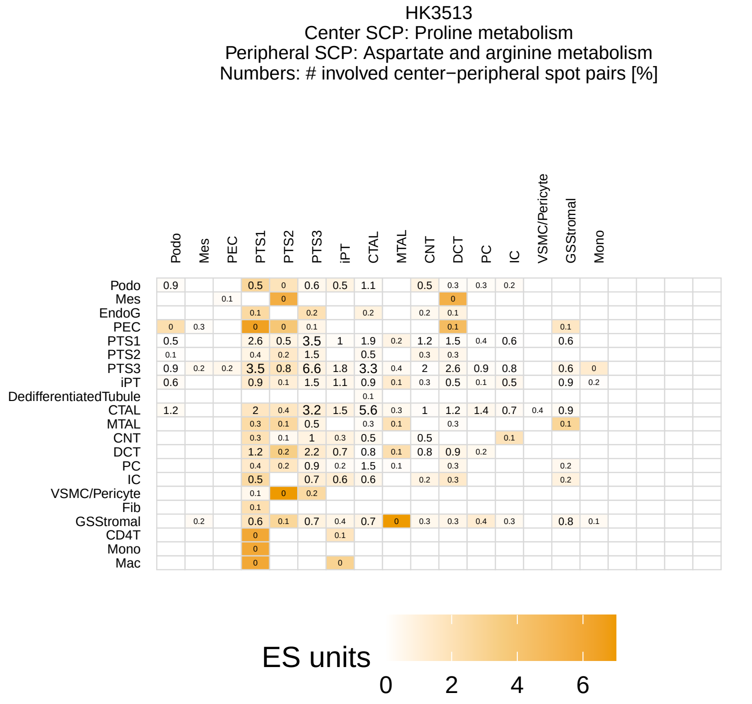
HK3513
Center SCP: Proline metabolism
Peripheral SCP: Aspartate and arginine metabolism
Numbers: # involved center−peripheral spot pairs [%]
ES units
0 2 4 6

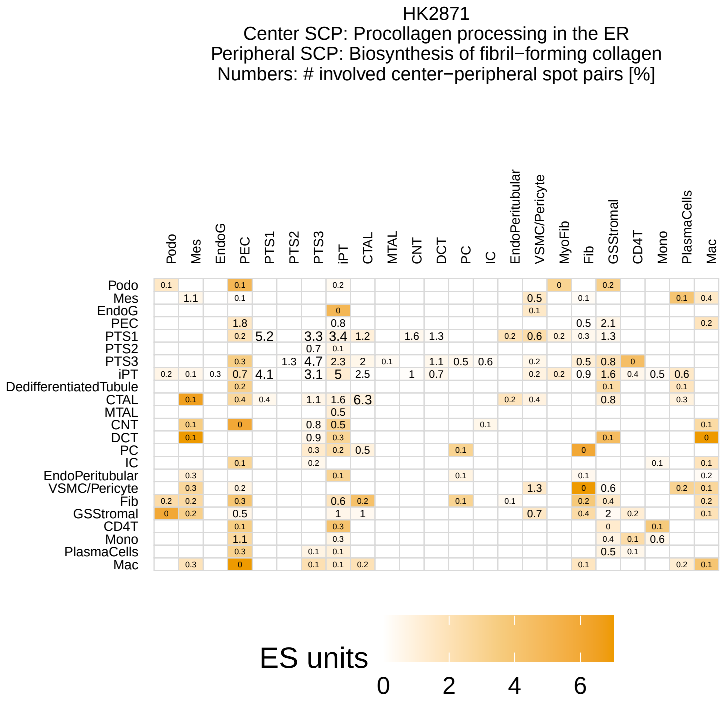
HK2871
Center SCP: Procollagen processing in the ER
Peripheral SCP: Biosynthesis of fibril−forming collagen
Numbers: # involved center−peripheral spot pairs [%]
ES units
0 2 4 6

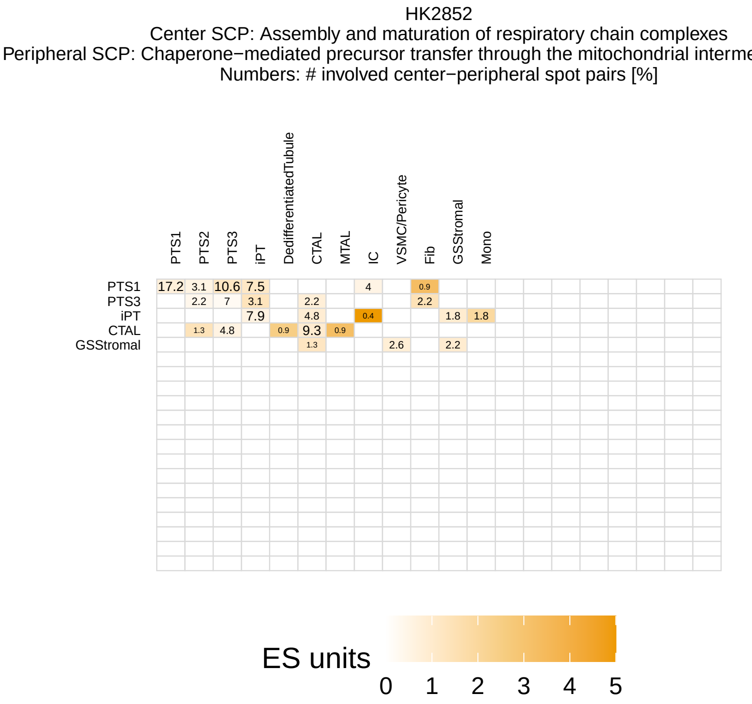
HK2852
Center SCP: Assembly and maturation of respiratory chain complexes
Peripheral SCP: Chaperone−mediated precursor transfer through the mitochondrial interme
Numbers: # involved center−peripheral spot pairs [%]
ES units
0 1 2 3 4 5

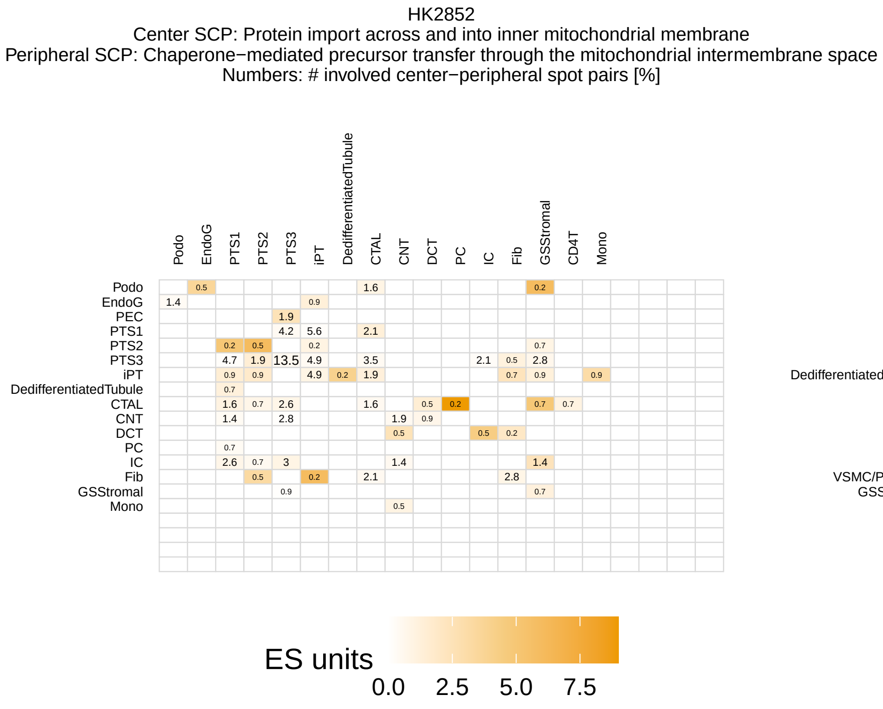
HK2852
Center SCP: Protein import across and into inner mitochondrial membrane
Peripheral SCP: Chaperone−mediated precursor transfer through the mitochondrial intermembrane space
Numbers: # involved center−peripheral spot pairs [%]
ES units
0.0 2.5 5.0 7.5

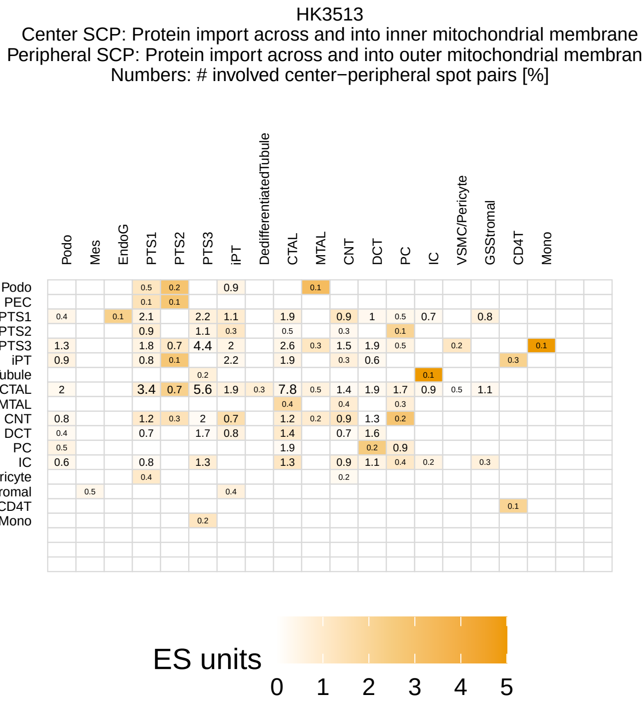
HK3513
Center SCP: Protein import across and into inner mitochondrial membrane
Peripheral SCP: Protein import across and into outer mitochondrial membran
Numbers: # involved center−peripheral spot pairs [%]
ES units
0 1 2 3 4 5

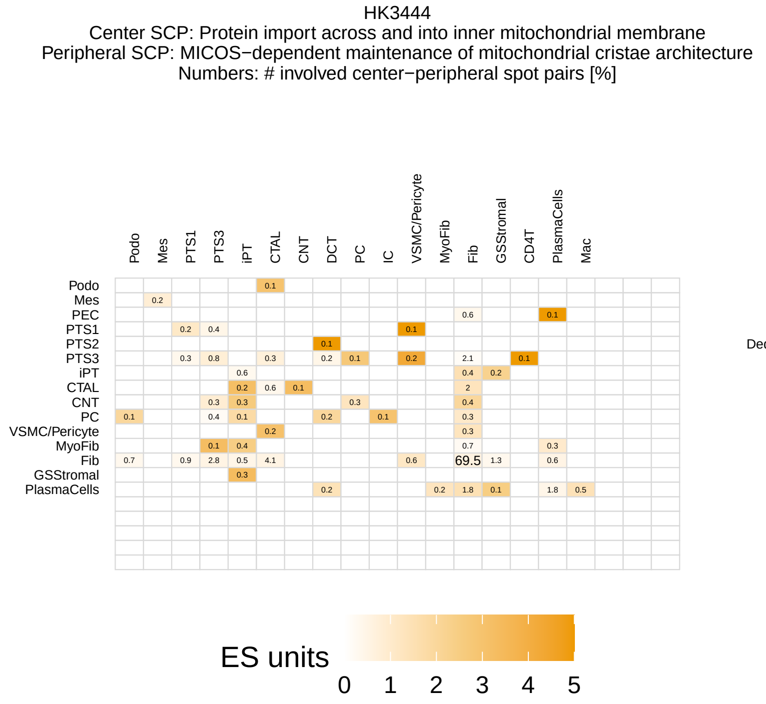
HK3444
Center SCP: Protein import across and into inner mitochondrial membrane
Peripheral SCP: MICOS−dependent maintenance of mitochondrial cristae architecture
Numbers: # involved center−peripheral spot pairs [%]
Podo Mes PTS1 PTS3 iPT CTAL CNT DCT PC IC VSMC/Pericyte MyoFib Fib GSStromal CD4T PlasmaCells Mac
Podo Mes PEC PTS1 PTS2 PTS3 iPT CTAL CNT PC VSMC/Pericyte MyoFib Fib GSStromal PlasmaCells
ES units
0 1 2 3 4 5

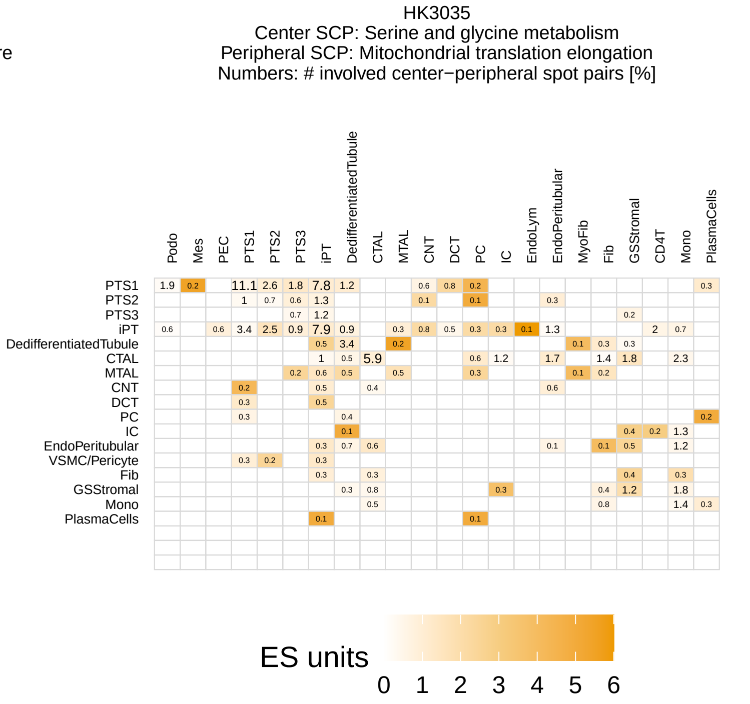
HK3035
Center SCP: Serine and glycine metabolism
Peripheral SCP: Mitochondrial translation elongation
Numbers: # involved center−peripheral spot pairs [%]
Podo Mes PEC PTS1 PTS2 PTS3 iPT DedifferentiatedTubule CTAL MTAL CNT DCT PC IC EndoLym EndoPeritubular MyoFib Fib GSStromal CD4T Mono PlasmaCells
PTS1 PTS2 PTS3 iPT DedifferentiatedTubule CTAL MTAL CNT DCT PC IC EndoPeritubular VSMC/Pericyte Fib GSStromal Mono PlasmaCells
ES units
0 1 2 3 4 5 6

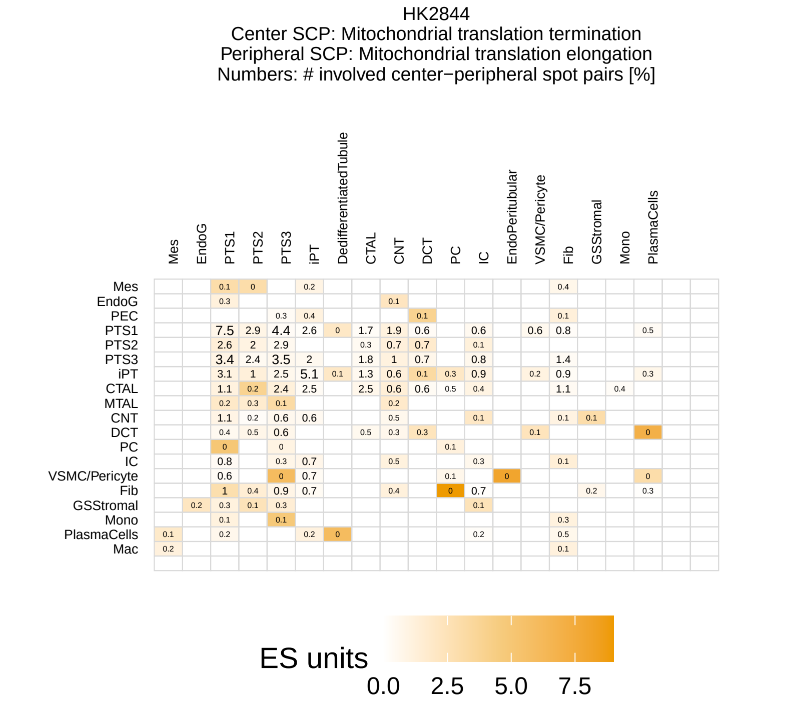
HK2844
Center SCP: Mitochondrial translation termination
Peripheral SCP: Mitochondrial translation elongation
Numbers: # involved center−peripheral spot pairs [%]
Mes EndoG PTS1 PTS2 PTS3 iPT DedifferentiatedTubule CTAL CNT DCT PC IC EndoPeritubular VSMC/Pericyte Fib GSStromal Mono PlasmaCells
Mes EndoG PEC PTS1 PTS2 PTS3 iPT CTAL MTAL CNT DCT PC IC VSMC/Pericyte Fib GSStromal Mono PlasmaCells Mac
ES units
0.0 2.5 5.0 7.5

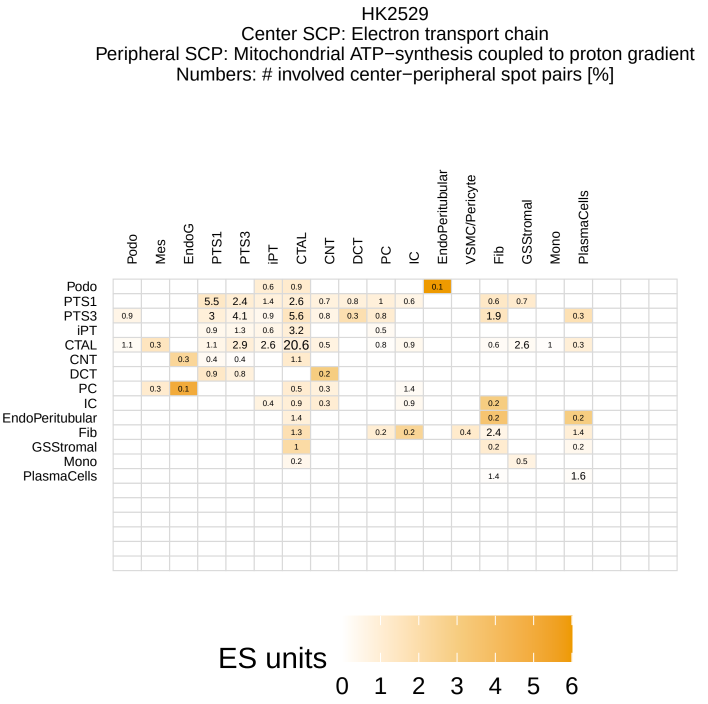
HK2529
Center SCP: Electron transport chain
Peripheral SCP: Mitochondrial ATP−synthesis coupled to proton gradient
Numbers: # involved center−peripheral spot pairs [%]
Podo Mes EndoG PTS1 PTS3 iPT CTAL CNT DCT PC IC EndoPeritubular VSMC/Pericyte Fib GSStromal Mono PlasmaCells
Podo PTS1 PTS3 iPT CTAL CNT DCT PC IC EndoPeritubular Fib GSStromal Mono PlasmaCells
ES units
0 1 2 3 4 5 6

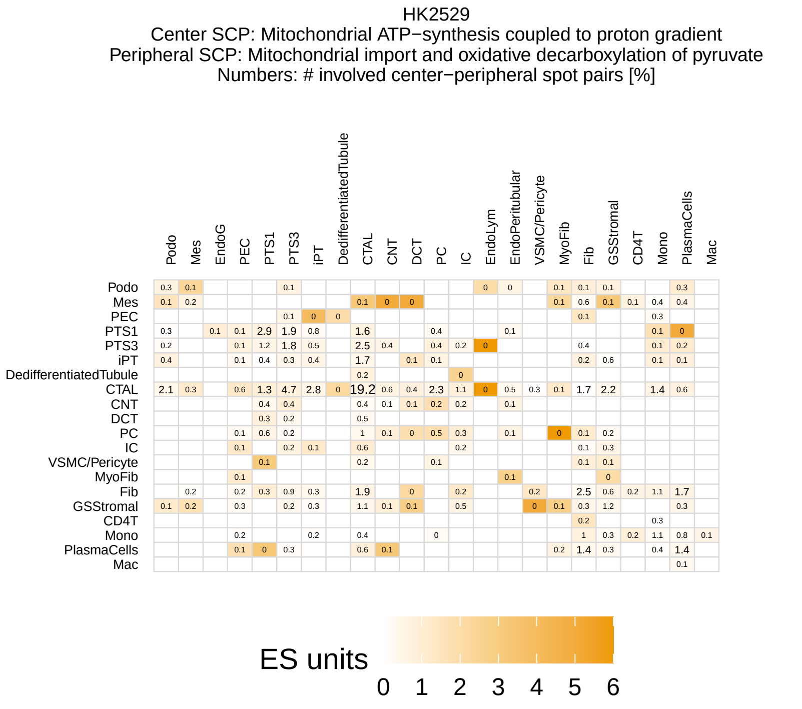
HK2529
Center SCP: Mitochondrial ATP−synthesis coupled to proton gradient
Peripheral SCP: Mitochondrial import and oxidative decarboxylation of pyruvate
Numbers: # involved center−peripheral spot pairs [%]
Podo Mes EndoG PEC PTS1 PTS3 iPT DedifferentiatedTubule CTAL CNT DCT PC IC EndoLym EndoPeritubular VSMC/Pericyte MyoFib Fib GSStromal CD4T Mono PlasmaCells Mac
Podo Mes PEC PTS1 PTS3 iPT DedifferentiatedTubule CTAL CNT DCT PC IC VSMC/Pericyte MyoFib Fib GSStromal CD4T Mono PlasmaCells Mac
ES units
0 1 2 3 4 5 6

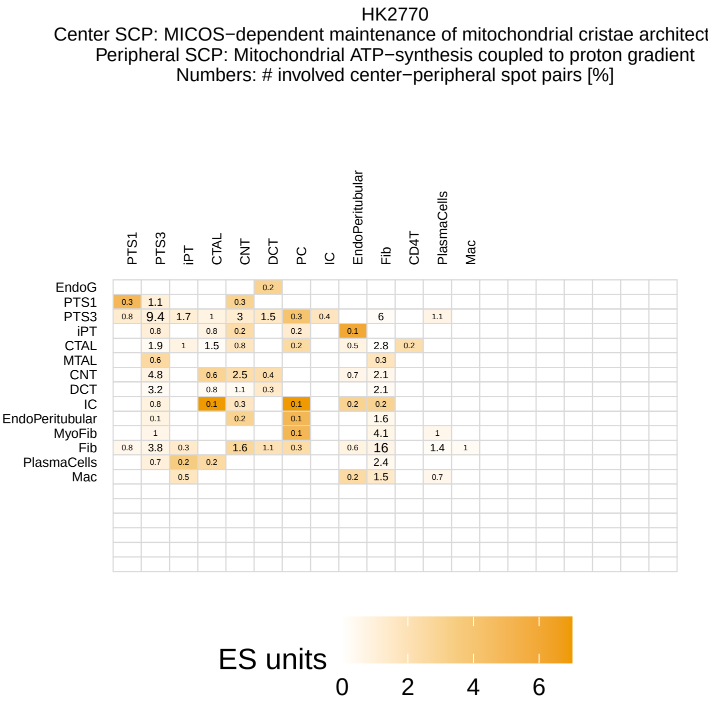
HK2770
Center SCP: MICOS−dependent maintenance of mitochondrial cristae architect
Peripheral SCP: Mitochondrial ATP−synthesis coupled to proton gradient
Numbers: # involved center−peripheral spot pairs [%]
PTS1 PTS3 iPT CTAL CNT DCT PC IC EndoPeritubular Fib CD4T PlasmaCells Mac
EndoG PTS1 PTS3 iPT CTAL MTAL CNT DCT IC EndoPeritubular MyoFib Fib PlasmaCells Mac
ES units
0 2 4 6

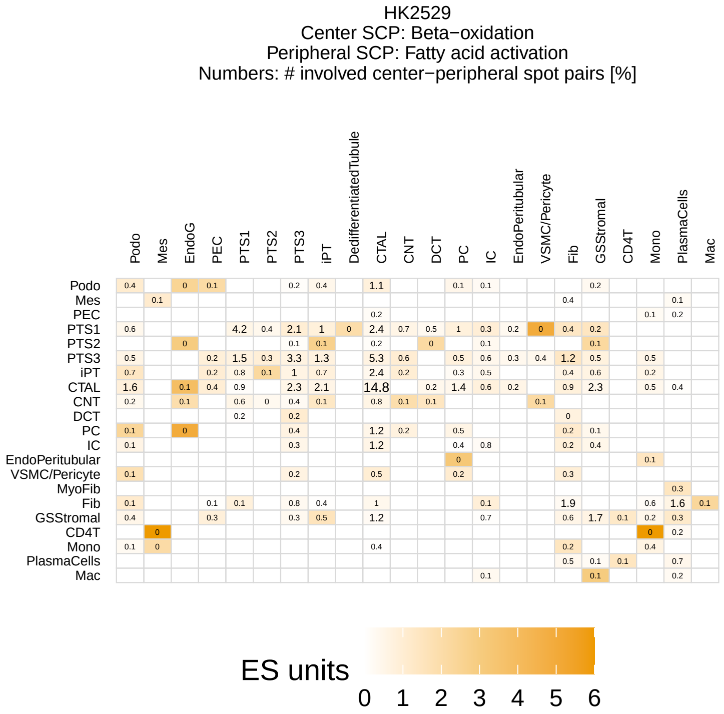
HK2529
Center SCP: Beta−oxidation
Peripheral SCP: Fatty acid activation
Numbers: # involved center−peripheral spot pairs [%]
ES units
0 1 2 3 4 5 6

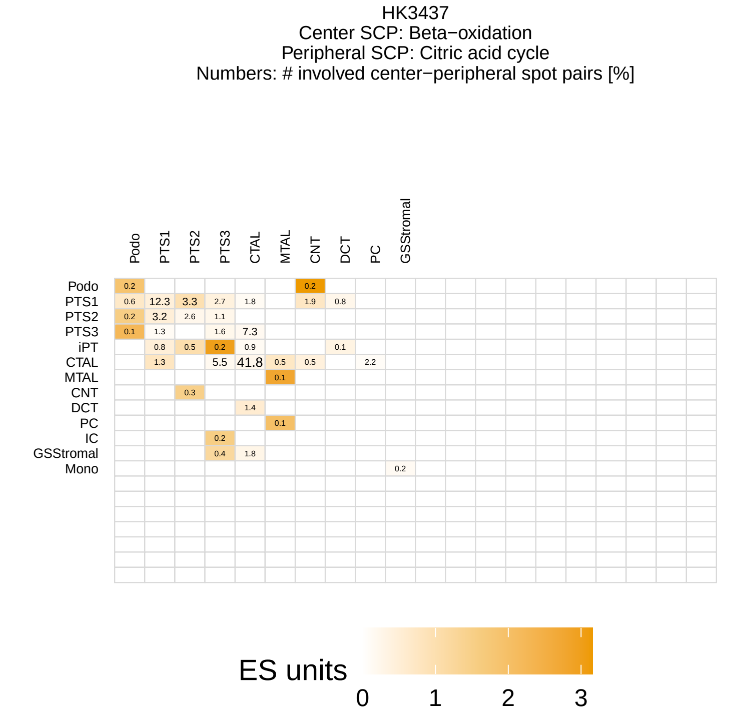
HK3437
Center SCP: Beta−oxidation
Peripheral SCP: Citric acid cycle
Numbers: # involved center−peripheral spot pairs [%]
ES units
0 1 2 3

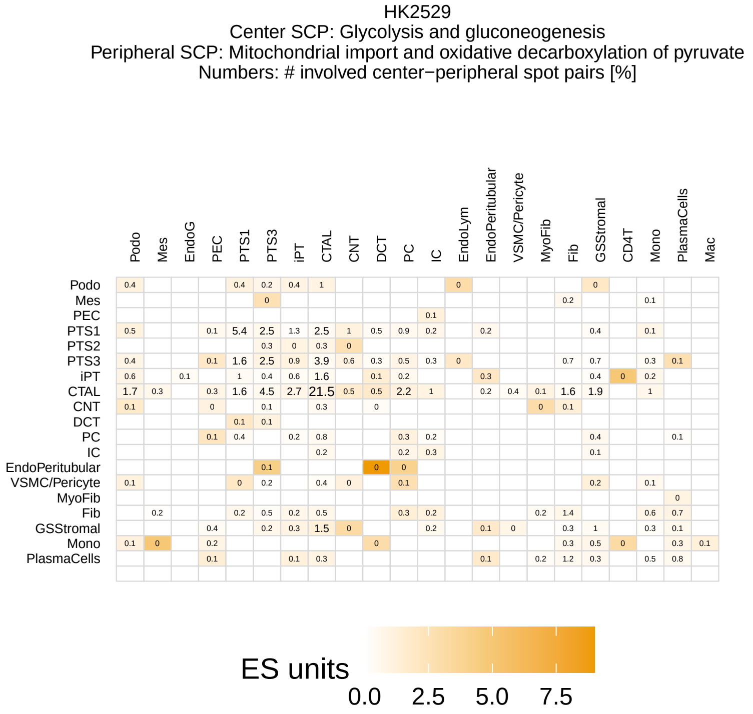
HK2529
Center SCP: Glycolysis and gluconeogenesis
Peripheral SCP: Mitochondrial import and oxidative decarboxylation of pyruvate
Numbers: # involved center−peripheral spot pairs [%]
ES units
0.0 2.5 5.0 7.5

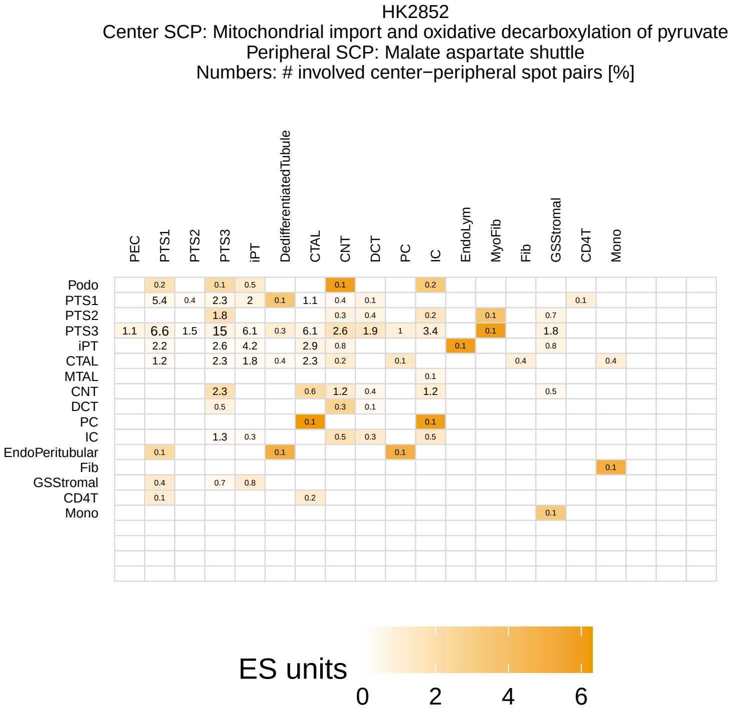
HK2852
Center SCP: Mitochondrial import and oxidative decarboxylation of pyruvate
Peripheral SCP: Malate aspartate shuttle
Numbers: # involved center−peripheral spot pairs [%]
ES units
0 2 4 6

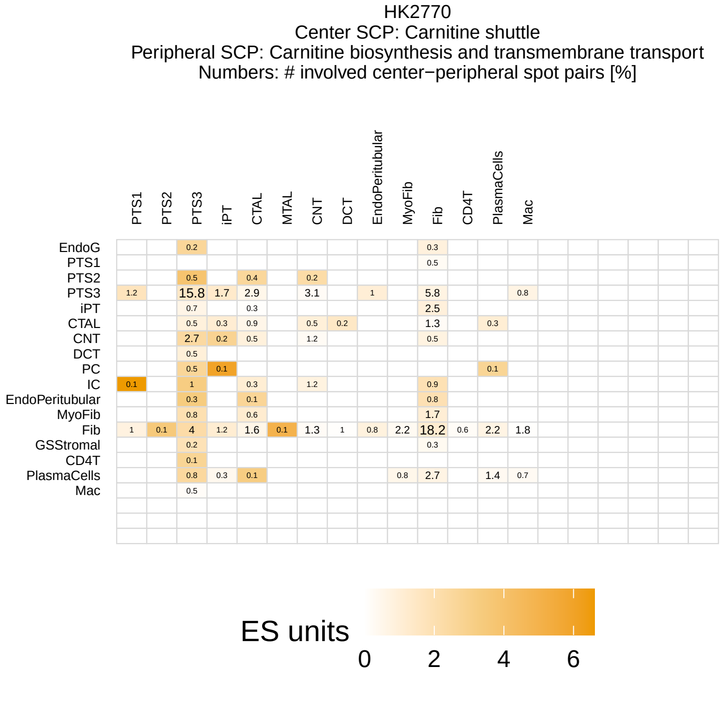
HK2770
Center SCP: Carnitine shuttle
Peripheral SCP: Carnitine biosynthesis and transmembrane transport
Numbers: # involved center−peripheral spot pairs [%]
ES units
0 2 4 6

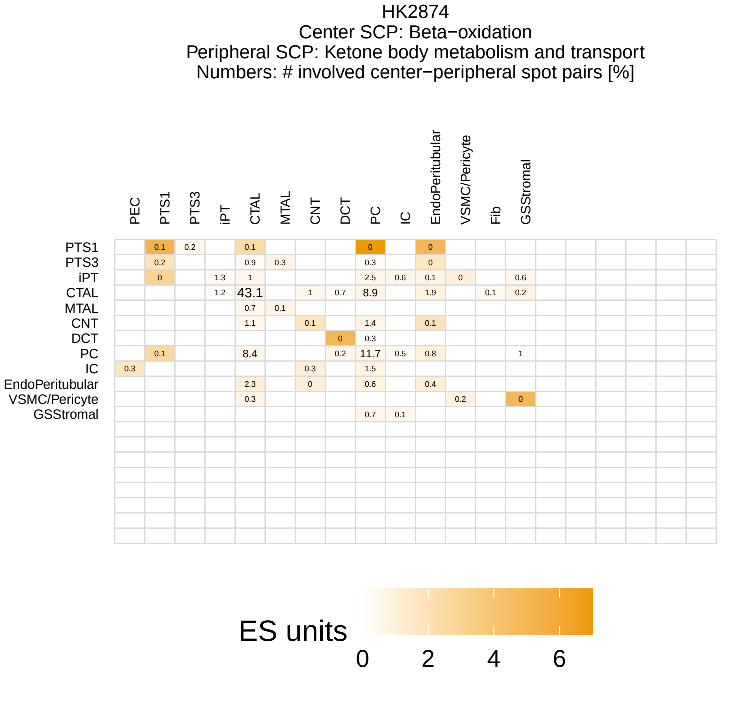
HK2874
Center SCP: Beta−oxidation
Peripheral SCP: Ketone body metabolism and transport
Numbers: # involved center−peripheral spot pairs [%]
ES units
0 2 4 6

HK2877
Center SCP: Thioredoxin−dependent peroxiredoxin−mediated peroxide detoxification
Peripheral SCP: Pentose phosphate pathway
Numbers: # involved center−peripheral spot pairs [%]

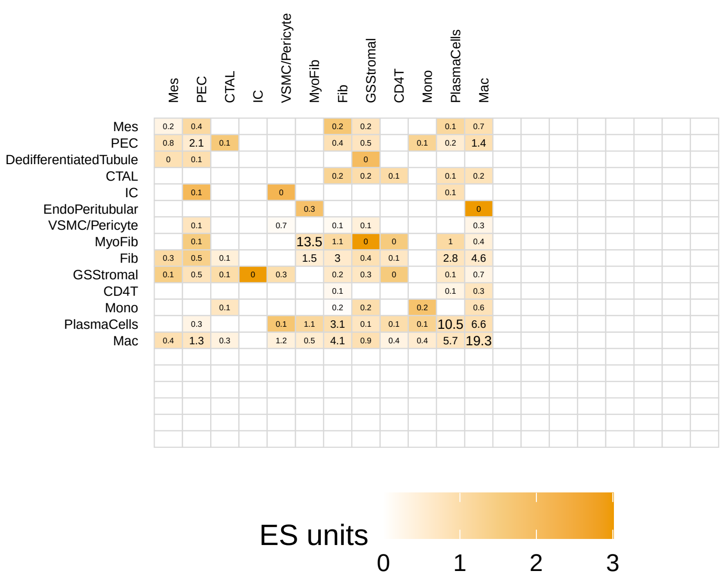


HK2770
Center SCP: Carrier−mediated sodium transmembrane transport
Peripheral SCP: Generation of electrochemical gradients by Na−K−ATPase
Numbers: # involved center−peripheral spot pairs [%]

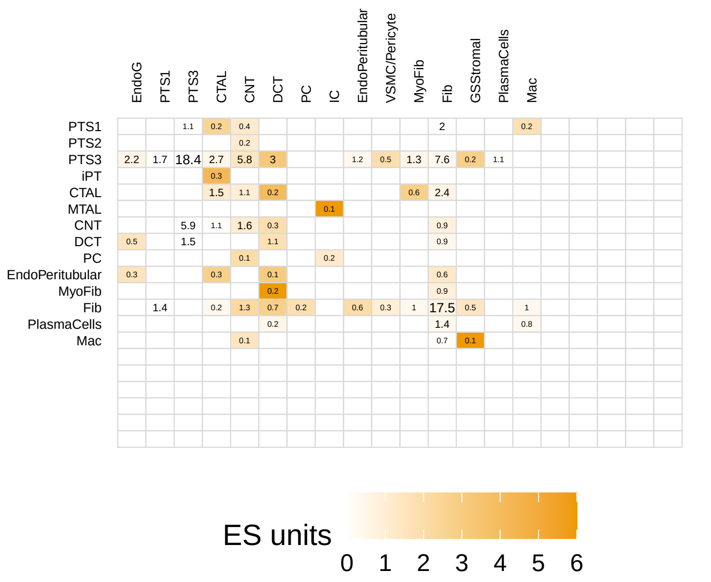


HK3437
Center SCP: Cellular urate import and export
Peripheral SCP: Aspartate and arginine metabolism
Numbers: # involved center−peripheral spot pairs [%]

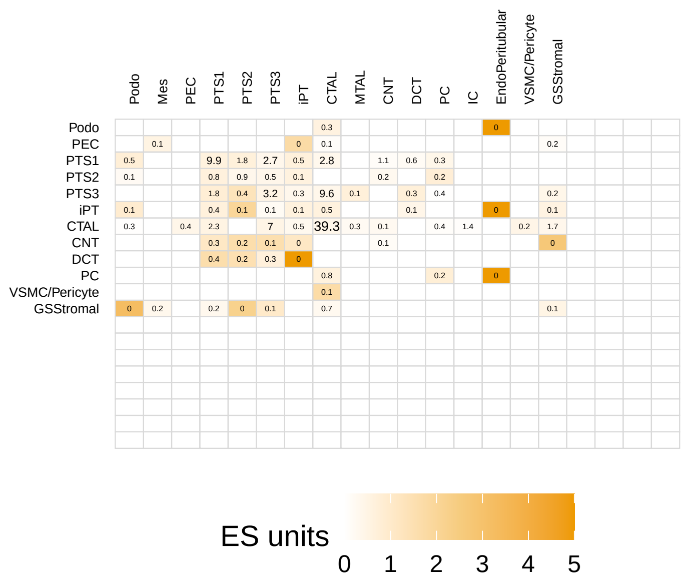


HK3437
Center SCP: Cellular xenobiotic uptake and efflux
Peripheral SCP: Phase II biotransformation via glucuronidation
Numbers: # involved center−peripheral spot pairs [%]

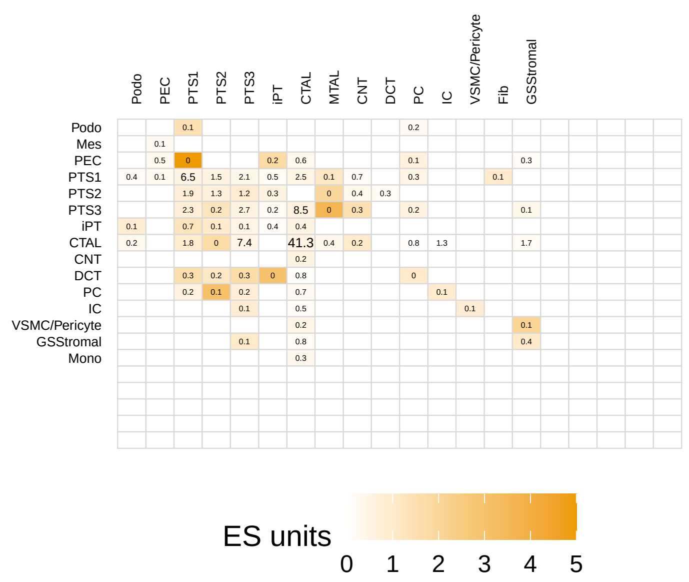


HK2871
Center SCP: Phase II biotransformation via gluthatione conjugation
Peripheral SCP: Phase II biotransformation via glucuronidation
Numbers: # involved center−peripheral spot pairs [%]

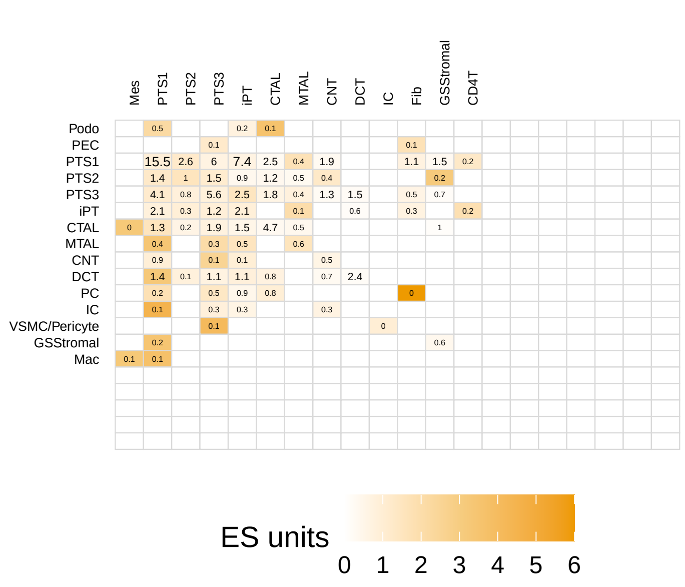


HK2871
Center SCP: Cobalamin metabolism
Peripheral SCP: Folate cycle
Numbers: # involved center−peripheral spot pairs [%]

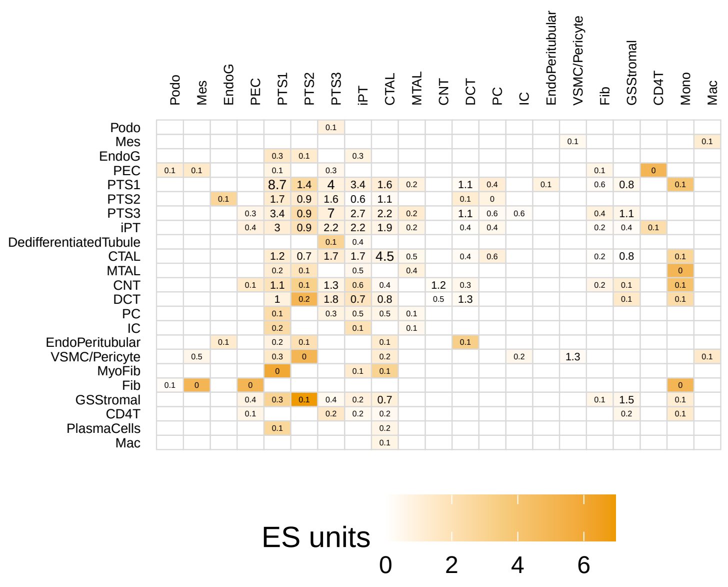

HK2871
Center SCP: Heme synthesis
Peripheral SCP: Branched tryptophan−kynurenine pathway producing NAD and diverse metabolites
Numbers: # involved center−peripheral spot pairs [%]

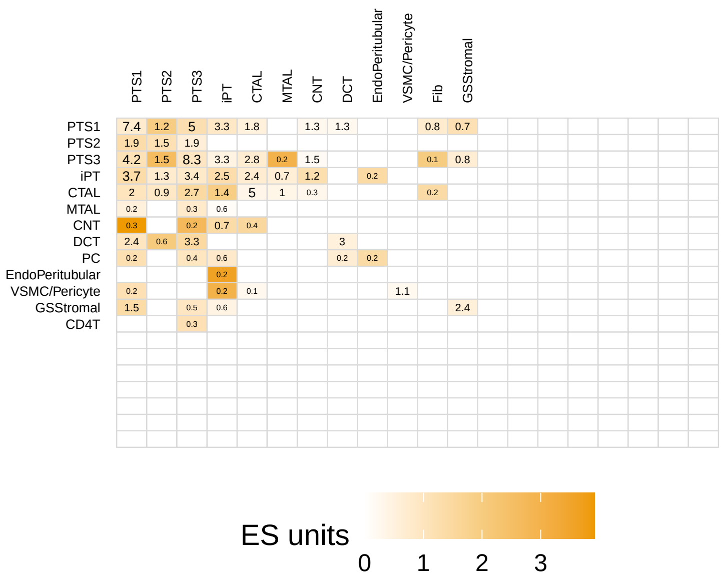


HK2874
Center SCP: Heme synthesis
Peripheral SCP: Adult hemoglobin and myoglobin assembly
Numbers: # involved center−peripheral spot pairs [%]

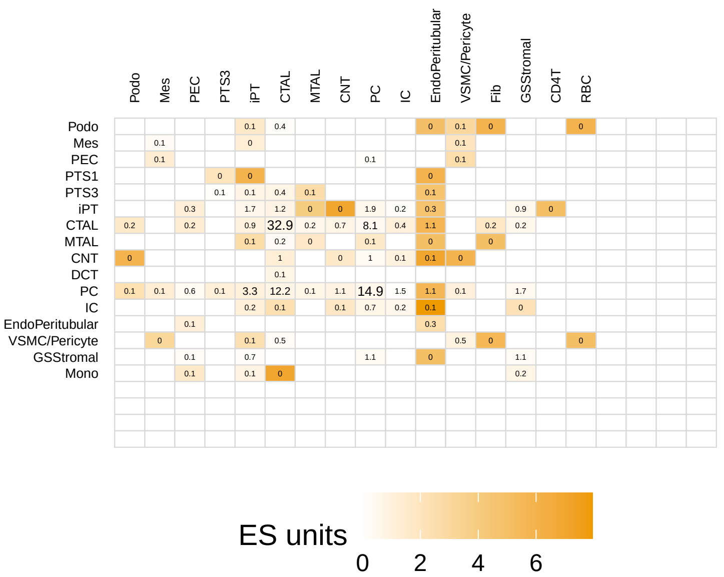


HK2874
Center SCP: Protein N−linked glycosylation in ER
Peripheral SCP: Biosynthesis and Golgi import of UDP−activated sugars for glycan synthesis
Numbers: # involved center−peripheral spot pairs [%]

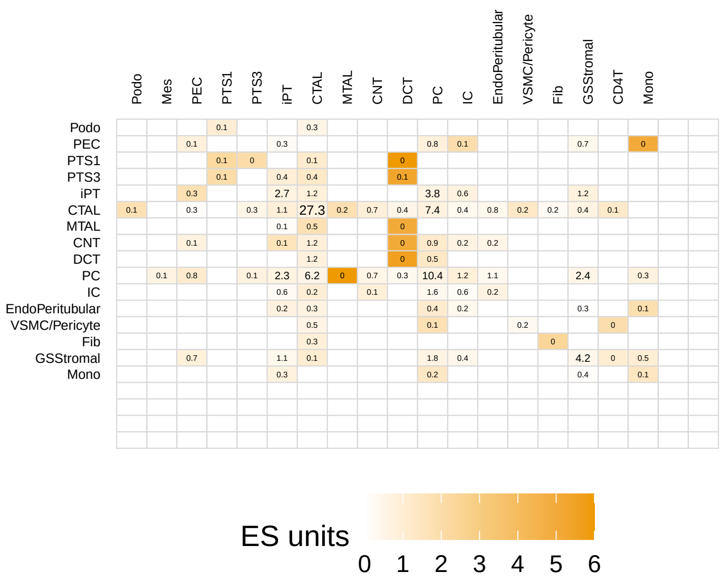


HK3513
Center SCP: Protein N−linked glycosylation in ER
Peripheral SCP: Fructose and mannose metabolism
Numbers: # involved center−peripheral spot pairs [%]

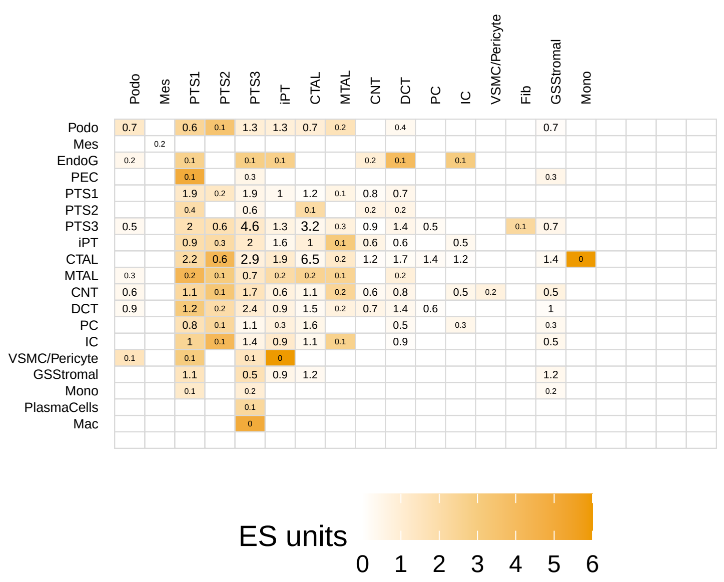


HK2529
Center SCP: Histone methylation and demethylation
Peripheral SCP: Chromatin organization by Polycomb repressive complexes
Numbers: # involved center−peripheral spot pairs [%]

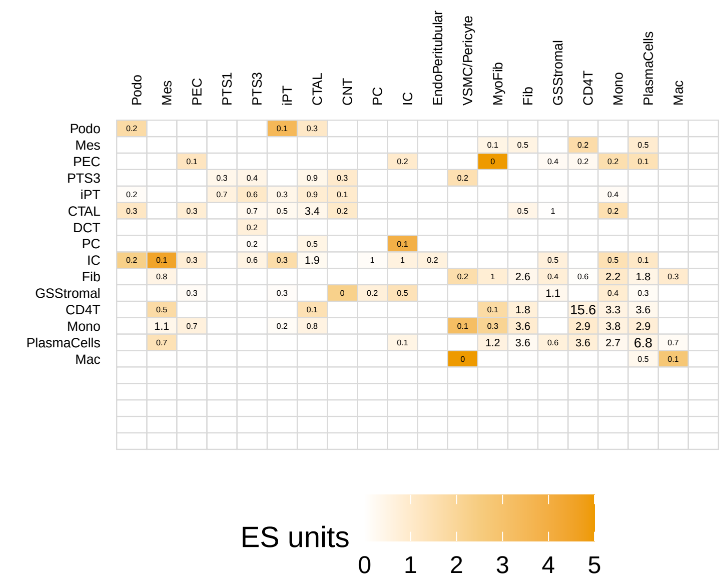


HK3432
Center SCP: Chromatin organization by Polycomb repressive complexes
Peripheral SCP: Transcription repression
Numbers: # involved center−peripheral spot pairs [%]

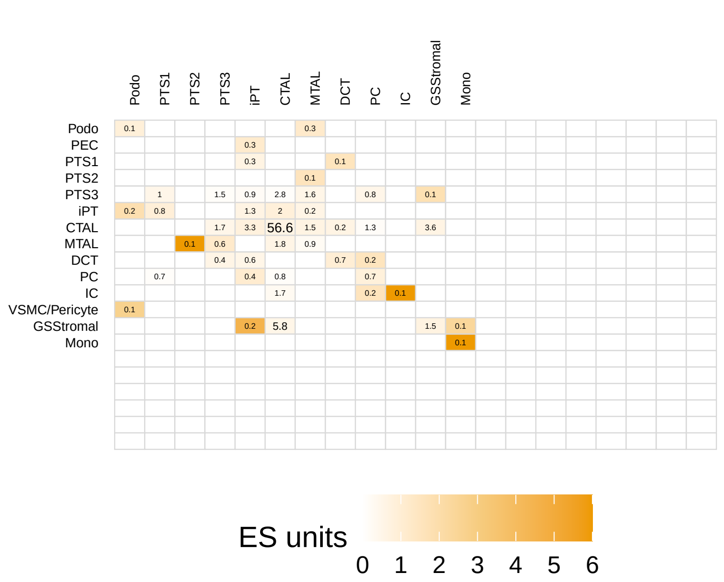

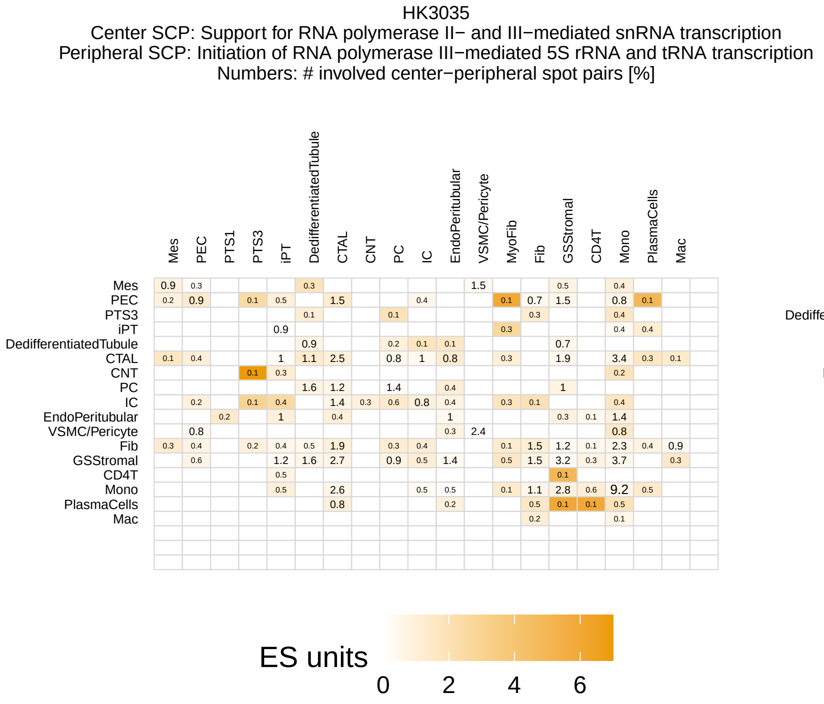
HK3035
Center SCP: Support for RNA polymerase II− and III−mediated snRNA transcription
Peripheral SCP: Initiation of RNA polymerase III−mediated 5S rRNA and tRNA transcription
Numbers: # involved center−peripheral spot pairs [%]
ES units
0 2 4 6

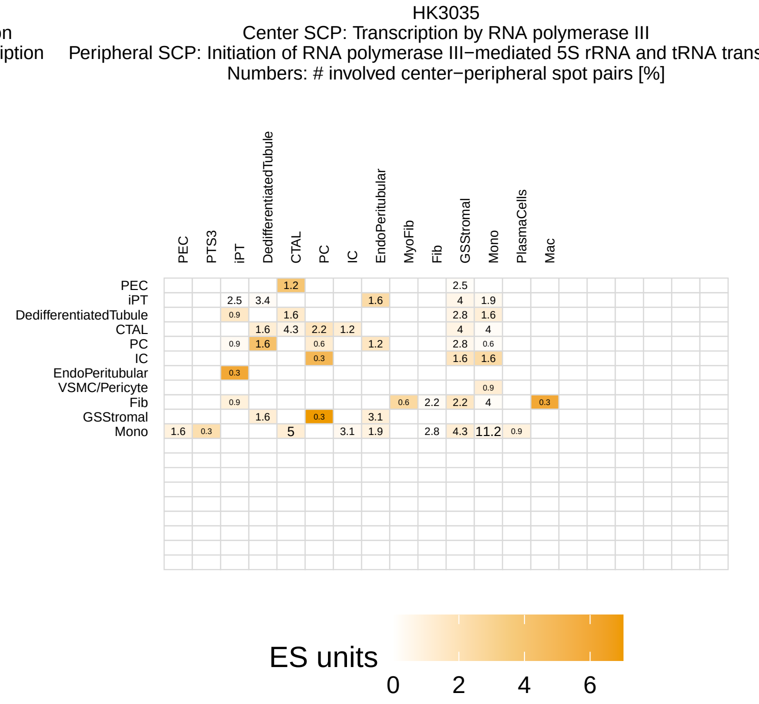
HK3035
Center SCP: Transcription by RNA polymerase III
Peripheral SCP: Initiation of RNA polymerase III−mediated 5S rRNA and tRNA trans
Numbers: # involved center−peripheral spot pairs [%]
ES units
0 2 4 6

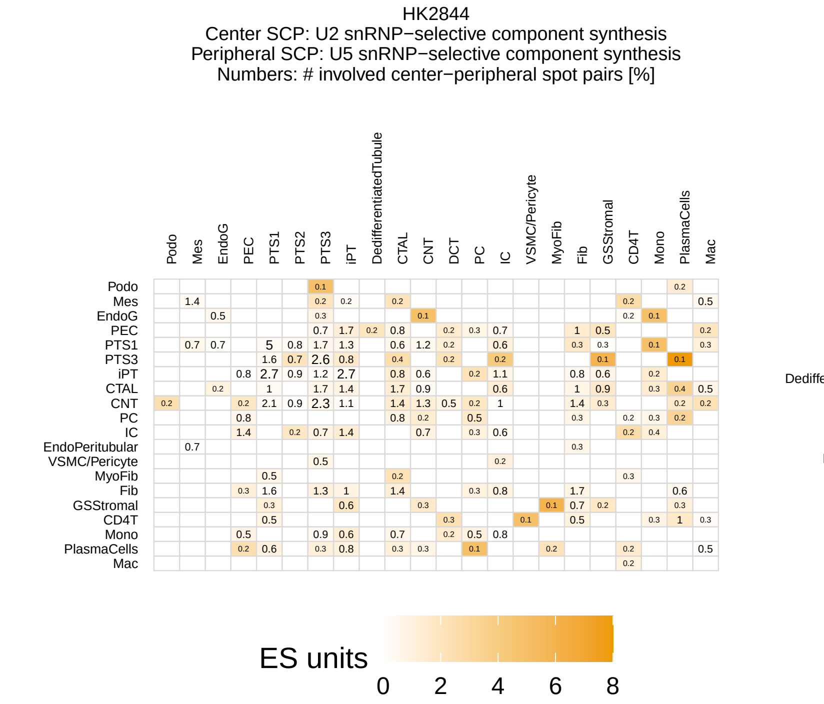
HK2844
Center SCP: U2 snRNP−selective component synthesis
Peripheral SCP: U5 snRNP−selective component synthesis
Numbers: # involved center−peripheral spot pairs [%]
ES units
0 2 4 6 8

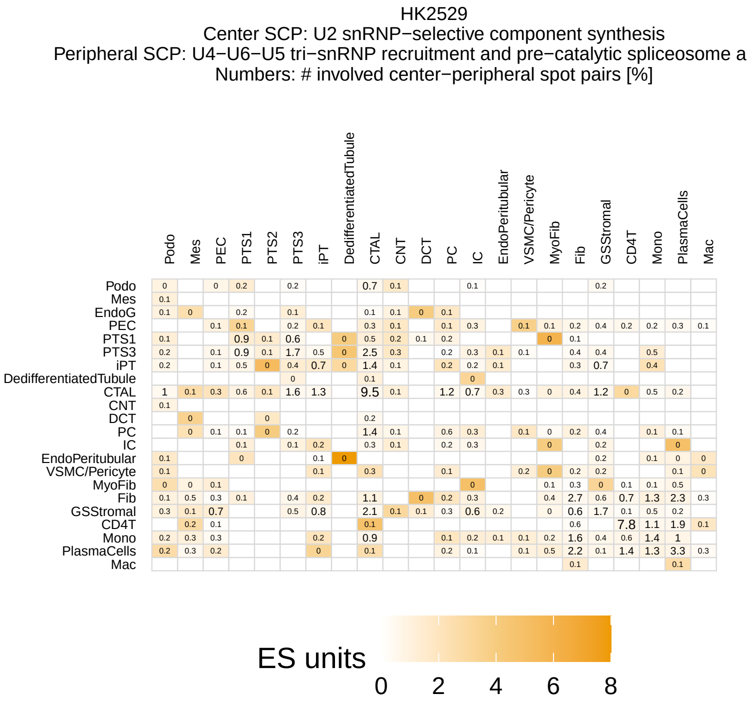
HK2529
Center SCP: U2 snRNP−selective component synthesis
Peripheral SCP: U4−U6−U5 tri−snRNP recruitment and pre−catalytic spliceosome a
Numbers: # involved center−peripheral spot pairs [%]
ES units
0 2 4 6 8

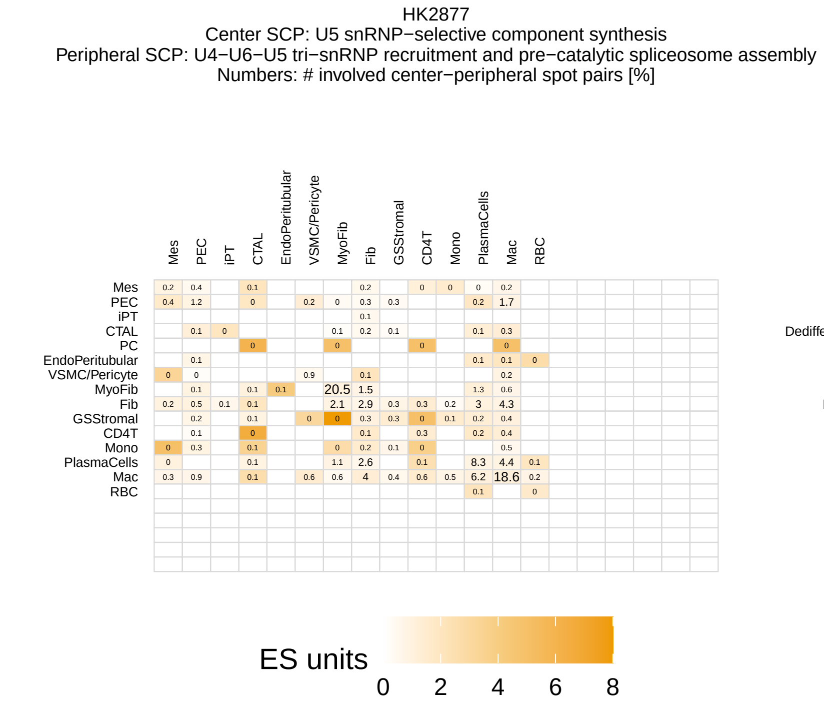
HK2877
Center SCP: U5 snRNP−selective component synthesis
Peripheral SCP: U4−U6−U5 tri−snRNP recruitment and pre−catalytic spliceosome assembly
Numbers: # involved center−peripheral spot pairs [%]
ES units
0 2 4 6 8

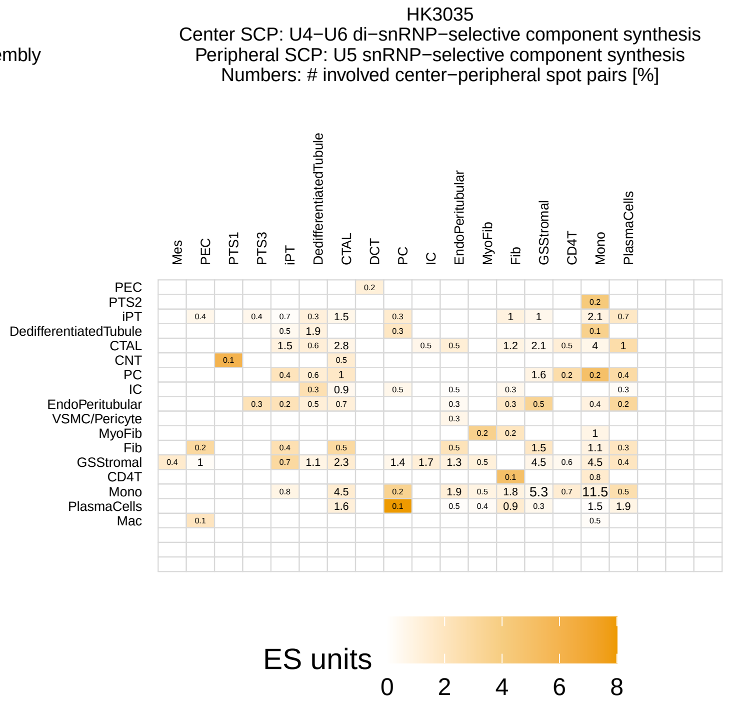
HK3035
Center SCP: U4−U6 di−snRNP−selective component synthesis
Peripheral SCP: U5 snRNP−selective component synthesis
Numbers: # involved center−peripheral spot pairs [%]
ES units
0 2 4 6 8

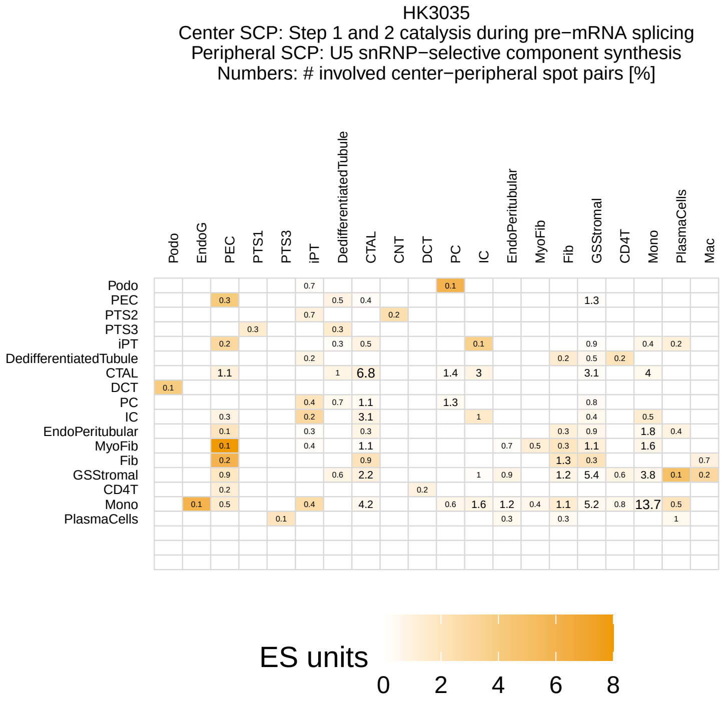
HK3035
Center SCP: Step 1 and 2 catalysis during pre−mRNA splicing
Peripheral SCP: U5 snRNP−selective component synthesis
Numbers: # involved center−peripheral spot pairs [%]
ES units
0 2 4 6 8

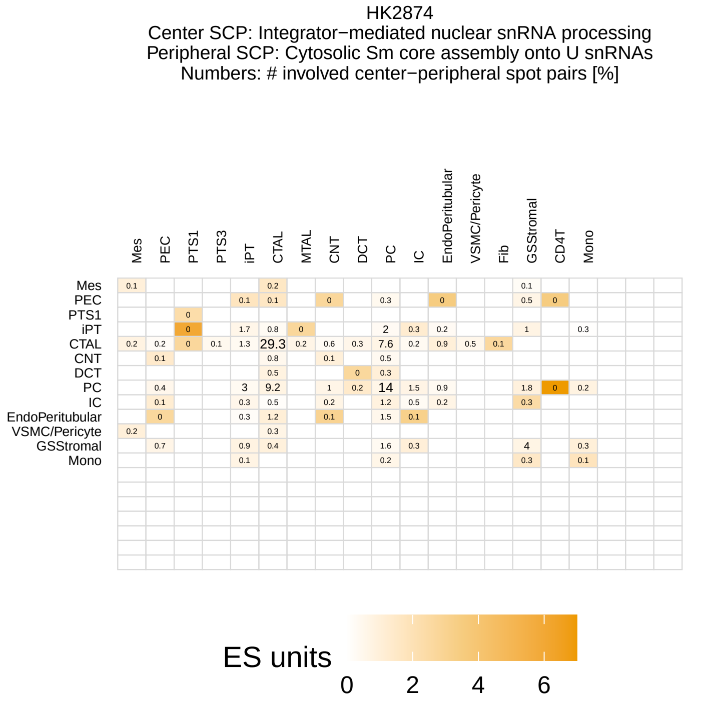
HK2874
Center SCP: Integrator−mediated nuclear snRNA processing
Peripheral SCP: Cytosolic Sm core assembly onto U snRNAs
Numbers: # involved center−peripheral spot pairs [%]
ES units
0 2 4 6

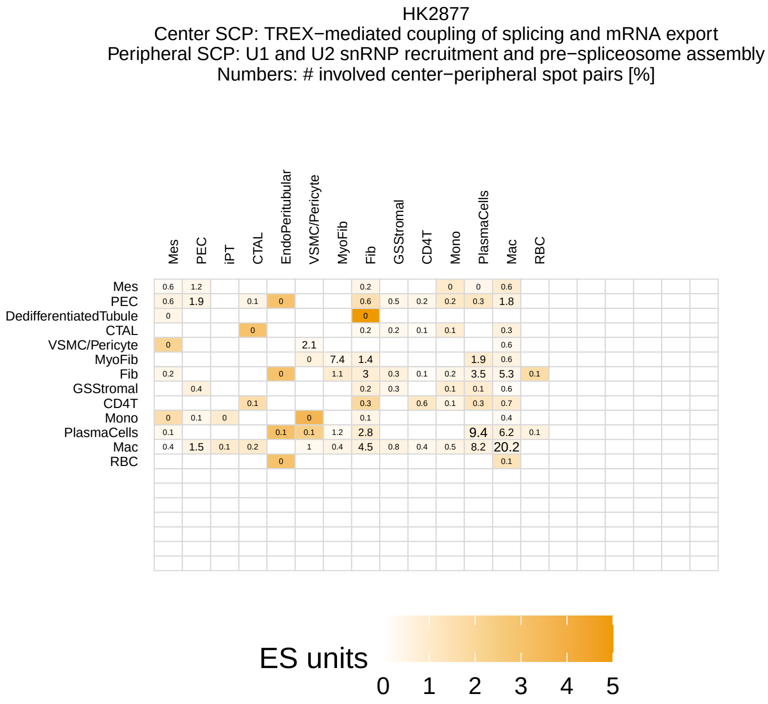
HK2877
Center SCP: TREX−mediated coupling of splicing and mRNA export
Peripheral SCP: U1 and U2 snRNP recruitment and pre−spliceosome assembly
Numbers: # involved center−peripheral spot pairs [%]
ES units
0 1 2 3 4 5

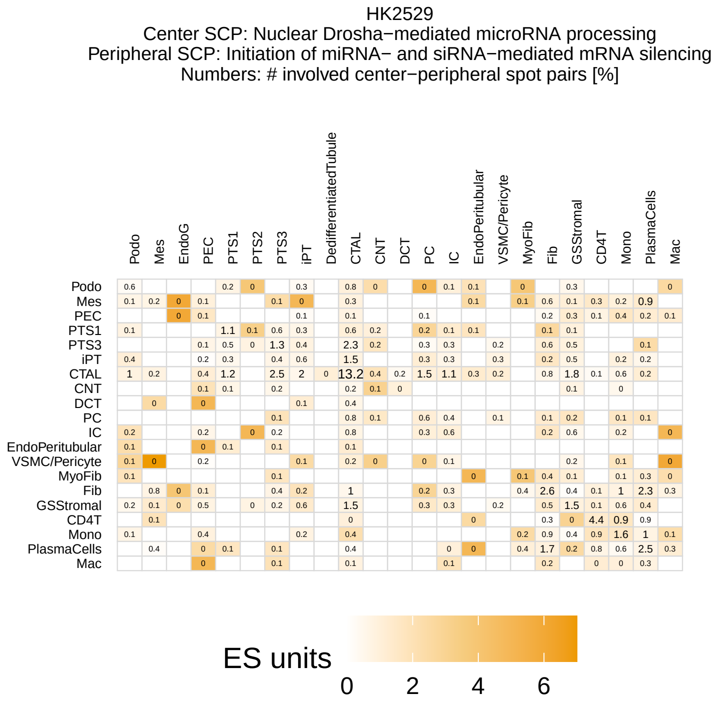
HK2529
Center SCP: Nuclear Drosha−mediated microRNA processing
Peripheral SCP: Initiation of miRNA− and siRNA−mediated mRNA silencing
Numbers: # involved center−peripheral spot pairs [%]
ES units
0 2 4 6

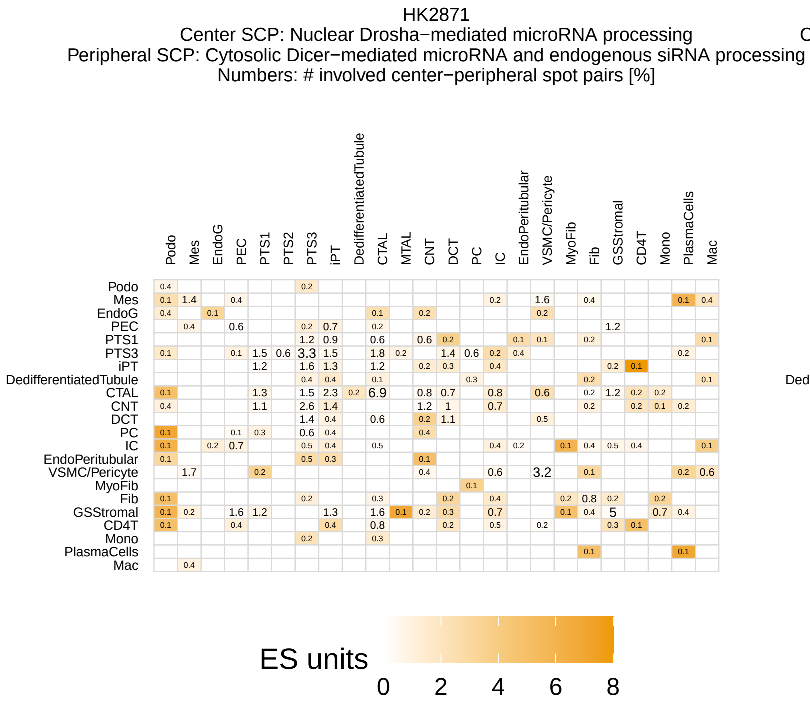
HK2871
Center SCP: Nuclear Drosha−mediated microRNA processing
Peripheral SCP: Cytosolic Dicer−mediated microRNA and endogenous siRNA processing
Numbers: # involved center−peripheral spot pairs [%]
ES units
0 2 4 6 8

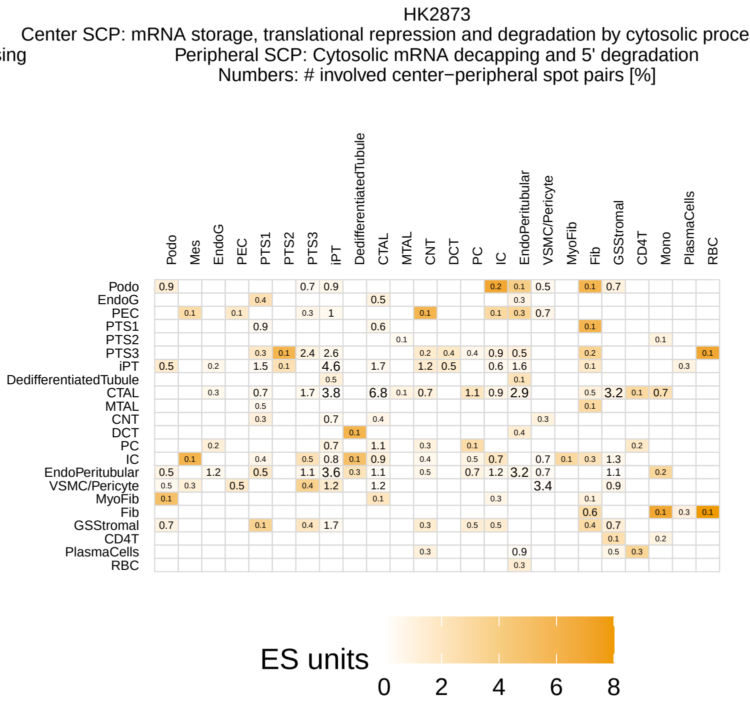
HK2873
Center SCP: mRNA storage, translational repression and degradation by cytosolic proce
Peripheral SCP: Cytosolic mRNA decapping and 5' degradation
Numbers: # involved center−peripheral spot pairs [%]
ES units
0 2 4 6 8

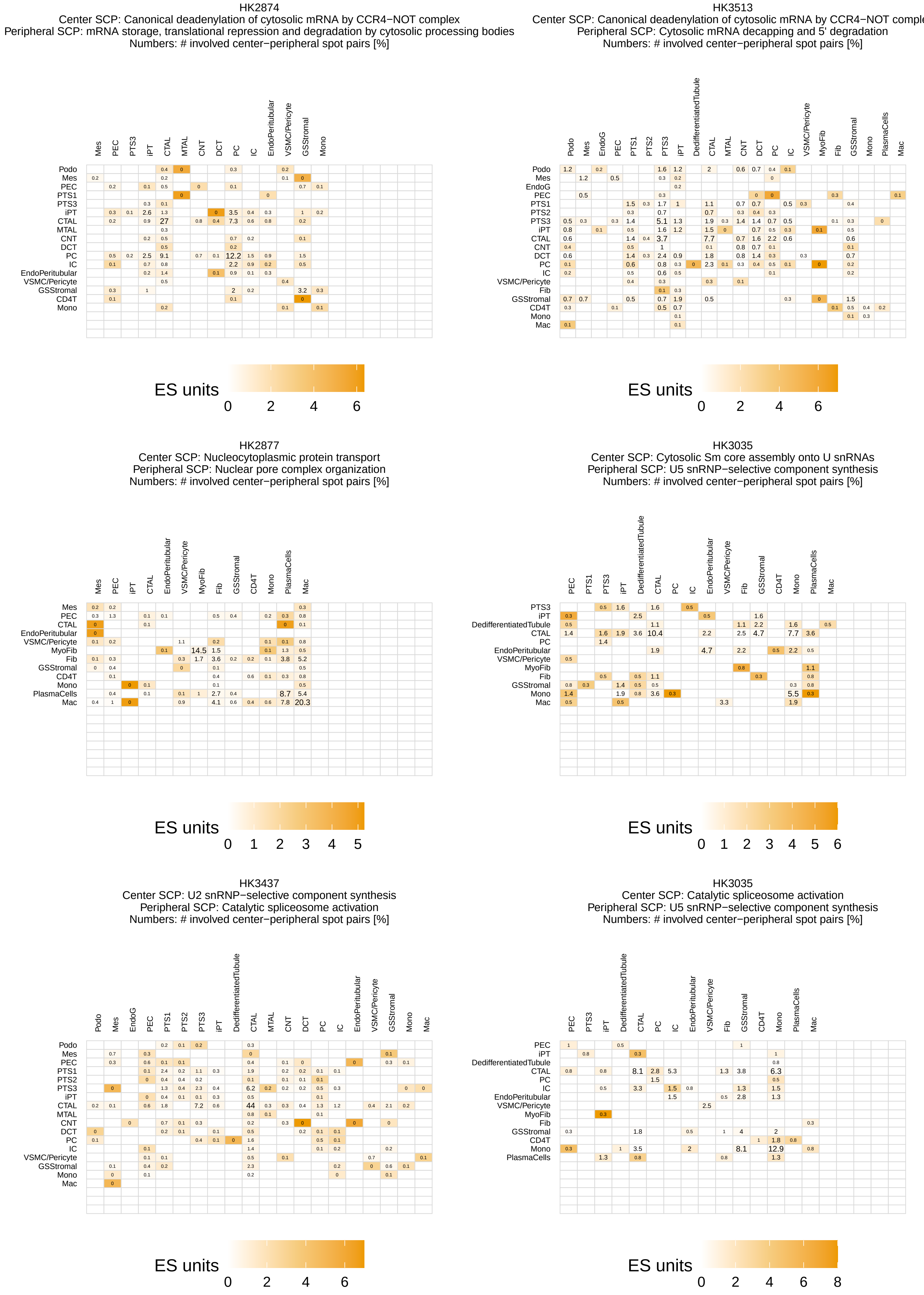
HK2874
Center SCP: Canonical deadenylation of cytosolic mRNA by CCR4−NOT complex
Peripheral SCP: mRNA storage, translational repression and degradation by cytosolic processing bodies
Numbers: # involved center−peripheral spot pairs [%]
ES units
0 2 4 6
HK3513
Center SCP: Canonical deadenylation of cytosolic mRNA by CCR4−NOT comple
Peripheral SCP: Cytosolic mRNA decapping and 5' degradation
Numbers: # involved center−peripheral spot pairs [%]
ES units
0 2 4 6
HK2877
Center SCP: Nucleocytoplasmic protein transport
Peripheral SCP: Nuclear pore complex organization
Numbers: # involved center−peripheral spot pairs [%]
ES units
0 1 2 3 4 5
HK3035
Center SCP: Cytosolic Sm core assembly onto U snRNAs
Peripheral SCP: U5 snRNP−selective component synthesis
Numbers: # involved center−peripheral spot pairs [%]
ES units
0 1 2 3 4 5 6
HK3437
Center SCP: U2 snRNP−selective component synthesis
Peripheral SCP: Catalytic spliceosome activation
Numbers: # involved center−peripheral spot pairs [%]
ES units
0 2 4 6
HK3035
Center SCP: Catalytic spliceosome activation
Peripheral SCP: U5 snRNP−selective component synthesis
Numbers: # involved center−peripheral spot pairs [%]
ES units
0 2 4 6 8

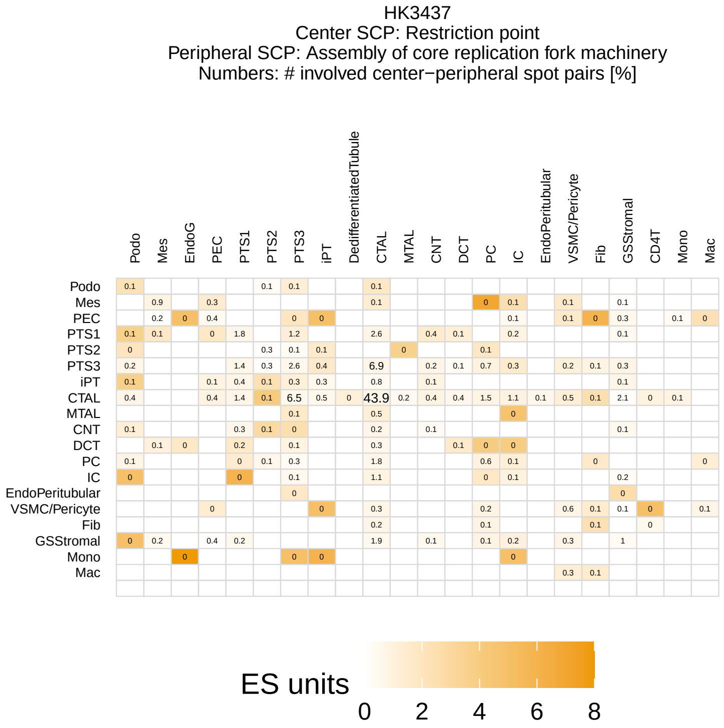
HK3437
Center SCP: Restriction point
Peripheral SCP: Assembly of core replication fork machinery
Numbers: # involved center−peripheral spot pairs [%]
Podo
Mes
EndoG
PEC
PTS1
PTS2
PTS3
iPT
DedifferentiatedTubule
CTAL
MTAL
CNT
DCT
PC
IC
EndoPeritubular
VSMC/Pericyte
Fib
GSStromal
CD4T
Mono
Mac
ES units
0 2 4 6 8

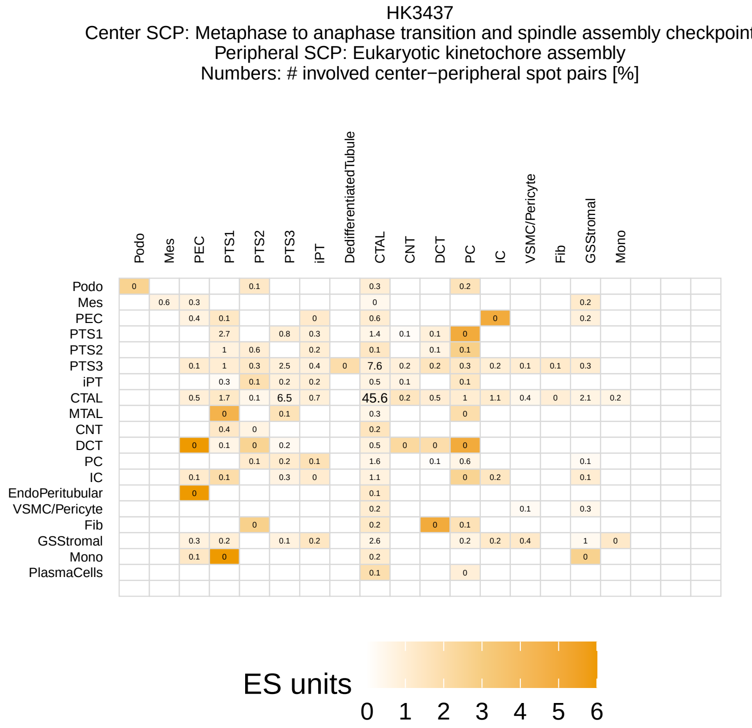
HK3437
Center SCP: Metaphase to anaphase transition and spindle assembly checkpoint
Peripheral SCP: Eukaryotic kinetochore assembly
Numbers: # involved center−peripheral spot pairs [%]
Podo
Mes
PEC
PTS1
PTS2
PTS3
iPT
DedifferentiatedTubule
CTAL
CNT
DCT
PC
IC
VSMC/Pericyte
Fib
GSStromal
Mono
MTAL
EndoPeritubular
PlasmaCells
ES units
0 1 2 3 4 5 6

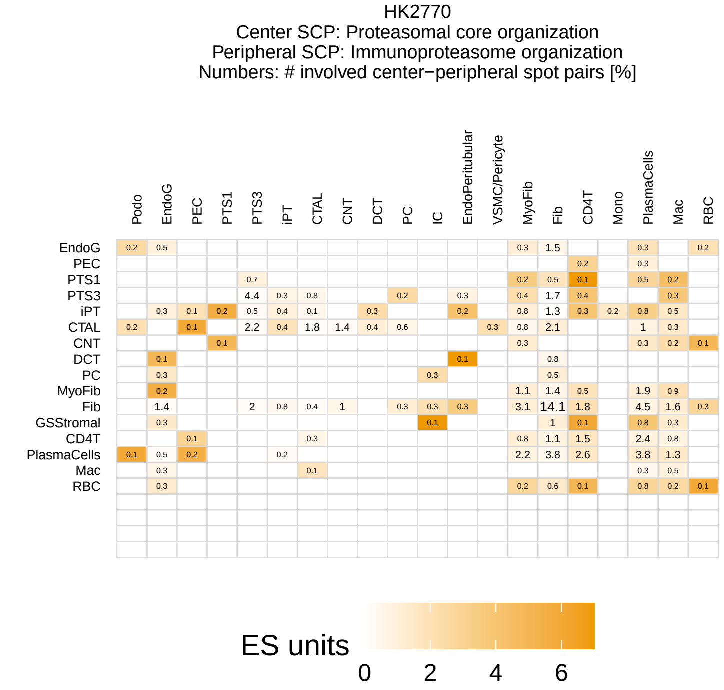
HK2770
Center SCP: Proteasomal core organization
Peripheral SCP: Immunoproteasome organization
Numbers: # involved center−peripheral spot pairs [%]
Podo
EndoG
PEC
PTS1
PTS3
iPT
CTAL
CNT
DCT
PC
IC
EndoPeritubular
VSMC/Pericyte
MyoFib
Fib
CD4T
Mono
PlasmaCells
Mac
RBC
GSStromal
ES units
0 2 4 6

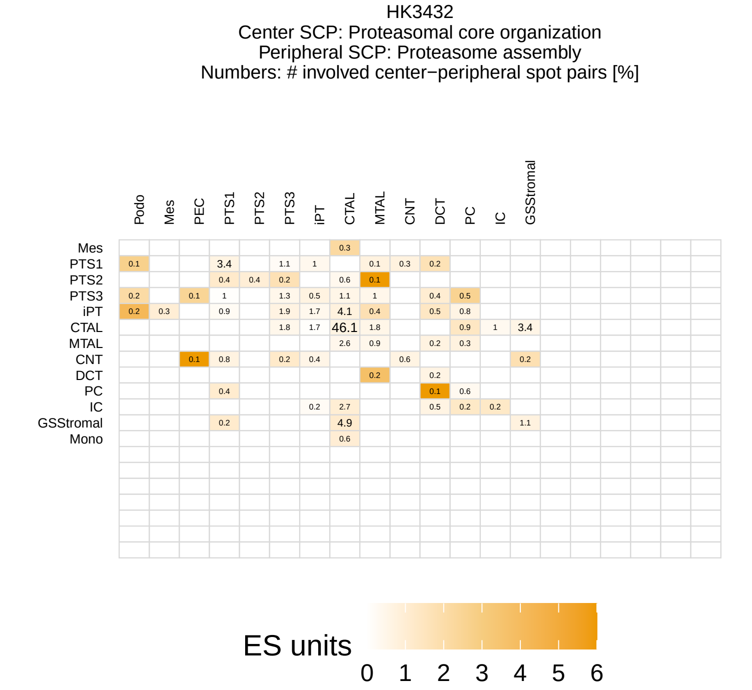
HK3432
Center SCP: Proteasomal core organization
Peripheral SCP: Proteasome assembly
Numbers: # involved center−peripheral spot pairs [%]
Podo
Mes
PEC
PTS1
PTS2
PTS3
iPT
CTAL
MTAL
CNT
DCT
PC
IC
GSStromal
Mono
ES units
0 1 2 3 4 5 6

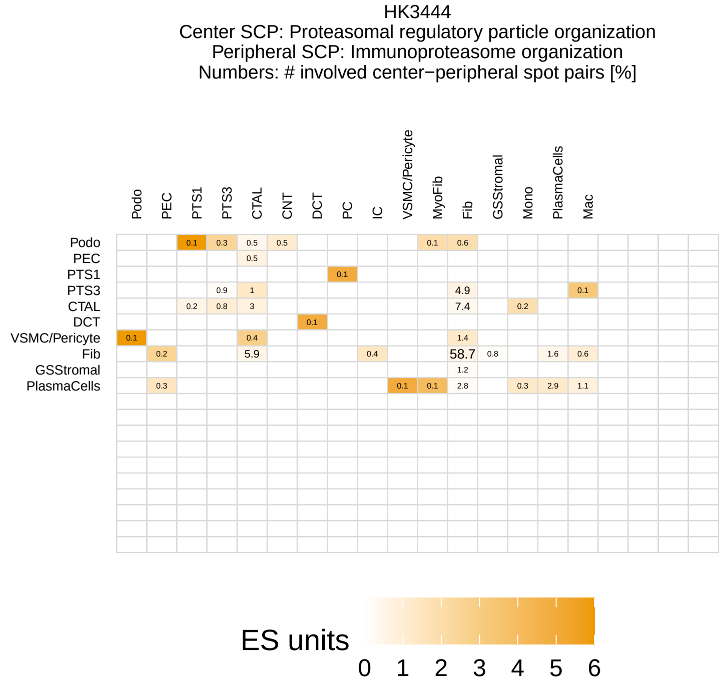
HK3444
Center SCP: Proteasomal regulatory particle organization
Peripheral SCP: Immunoproteasome organization
Numbers: # involved center−peripheral spot pairs [%]
Podo
PEC
PTS1
PTS3
CTAL
CNT
DCT
PC
IC
VSMC/Pericyte
MyoFib
Fib
GSStromal
Mono
PlasmaCells
Mac
ES units
0 1 2 3 4 5 6

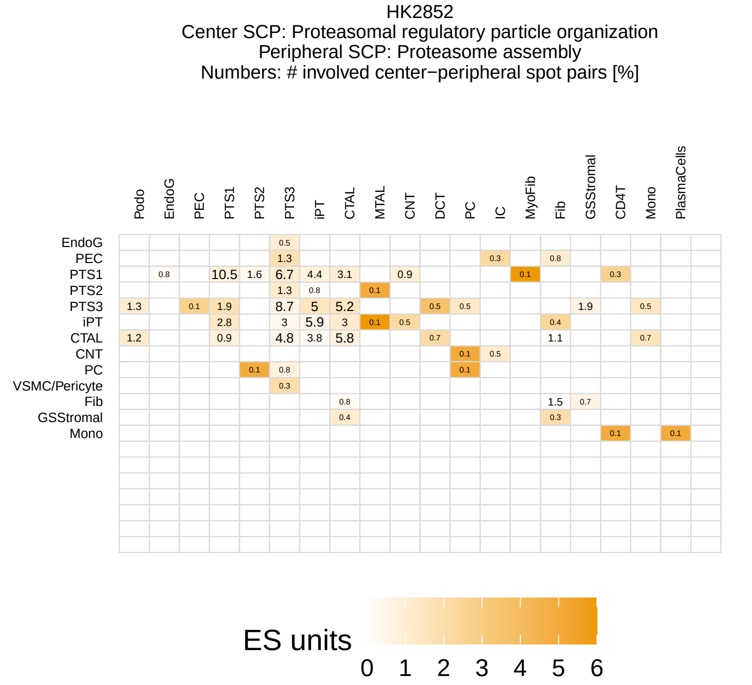
HK2852
Center SCP: Proteasomal regulatory particle organization
Peripheral SCP: Proteasome assembly
Numbers: # involved center−peripheral spot pairs [%]
Podo
EndoG
PEC
PTS1
PTS2
PTS3
iPT
CTAL
MTAL
CNT
DCT
PC
IC
MyoFib
Fib
GSStromal
CD4T
Mono
PlasmaCells
VSMC/Pericyte
ES units
0 1 2 3 4 5 6

HK2874
Center SCP: Antigen presentation via MHC class I molecules
Peripheral SCP: Immunoproteasome organization
Numbers: # involved center−peripheral spot pairs [%]

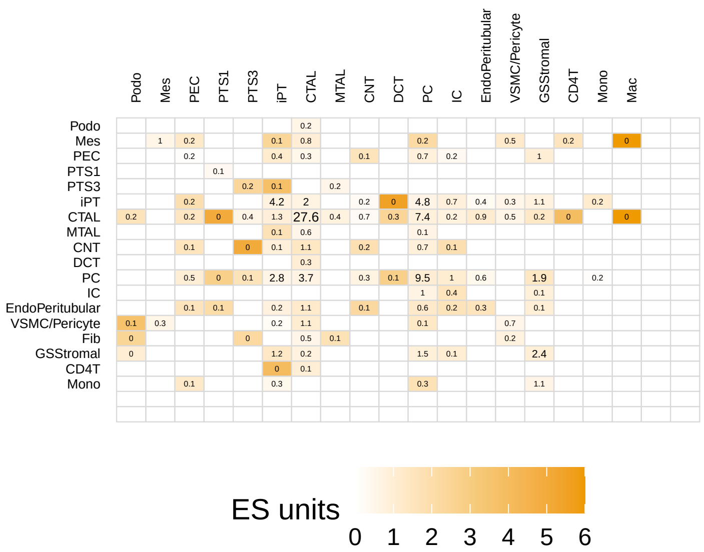


HK2873
Center SCP: Ras signaling pathway
Peripheral SCP: Fibroblast growth factor receptor signaling
Numbers: # involved center−peripheral spot pairs [%]

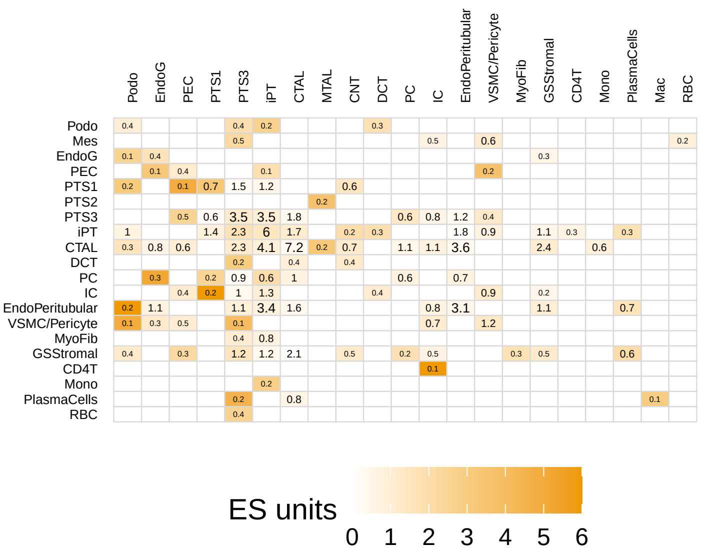


HK3432
Center SCP: Regulation of G−protein signaling by beta and gamma subunits
Peripheral SCP: GPCR desensitization and endocytic adaptor recruitment
Numbers: # involved center−peripheral spot pairs [%]

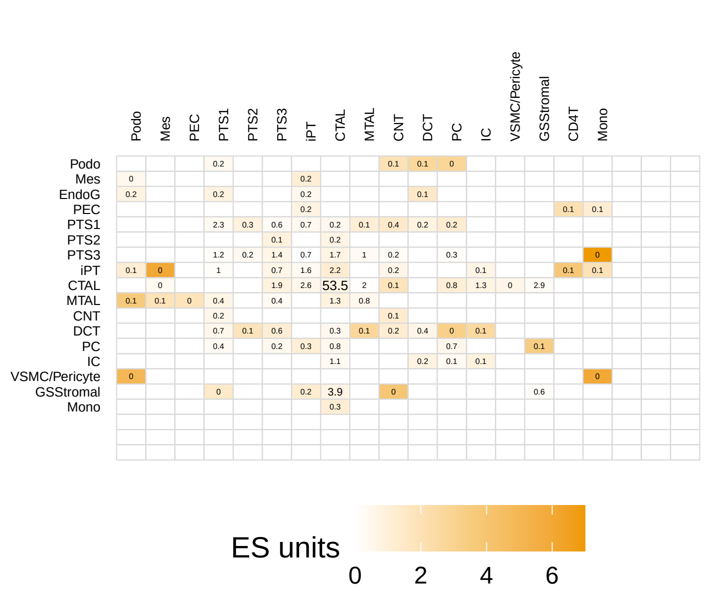


HK3035
Center SCP: Cofactor−mediated transcriptional regulation of multiple nuclear recep
Peripheral SCP: Retinoic acid receptor signaling
Numbers: # involved center−peripheral spot pairs [%]

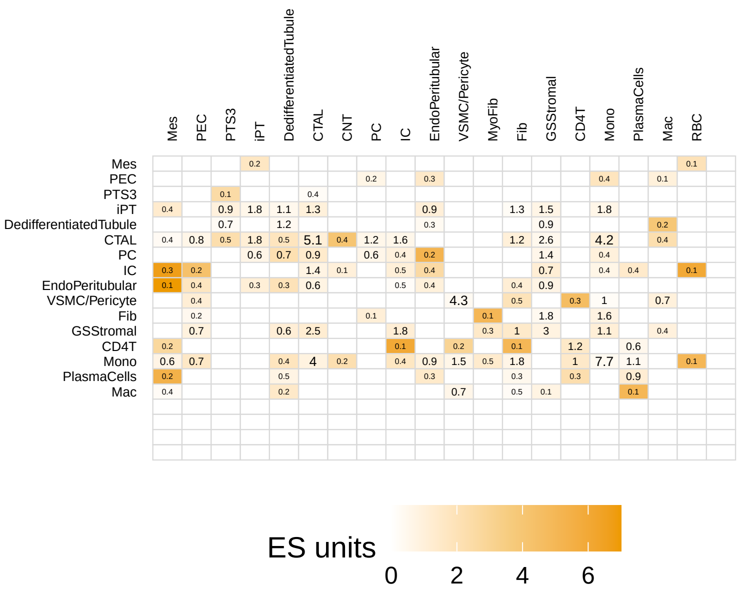


HK2877
Center SCP: Inhibition of apoptosis
Peripheral SCP: Extrinsic apoptosis pathway
Numbers: # involved center−peripheral spot pairs [%]

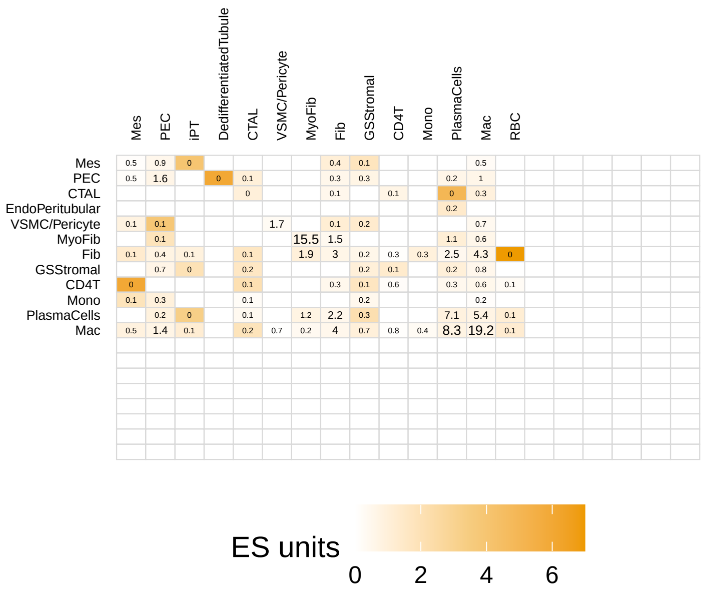


HK2877
Center SCP: Ceramide biosynthesis
Peripheral SCP: Extrinsic apoptosis pathway
Numbers: # involved center−peripheral spot pairs [%]

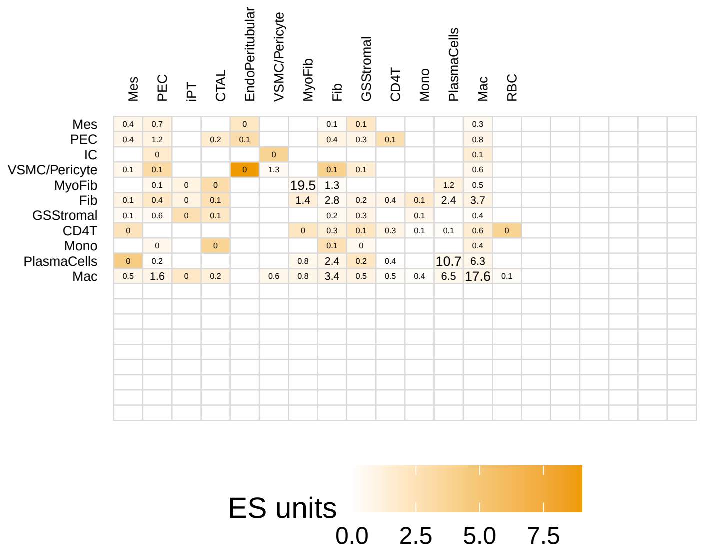

HK2873
Center SCP: Vesicle biogenesis at ER membrane
Peripheral SCP: Vesicle tethering at cis−Golgi
Numbers: # involved center−peripheral spot pairs [%]

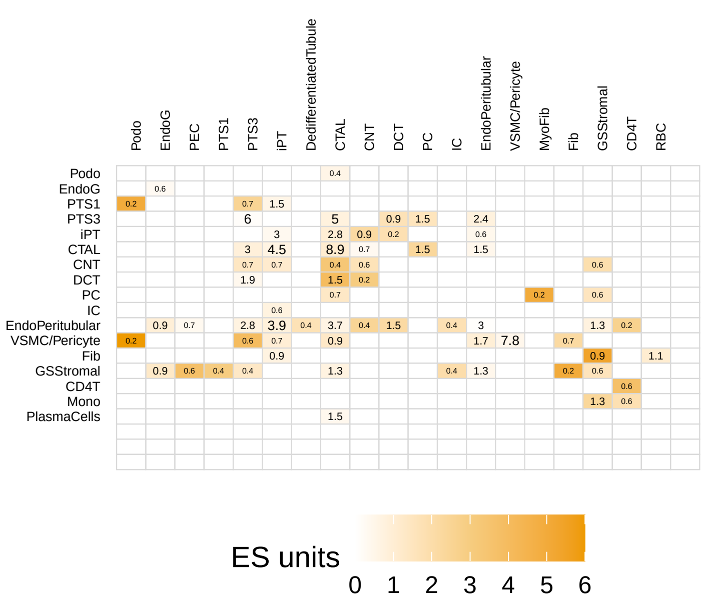


HK2529
Center SCP: Vesicle biogenesis at cis−Golgi
Peripheral SCP: Vesicle biogenesis at ER membrane
Numbers: # involved center−peripheral spot pairs [%]

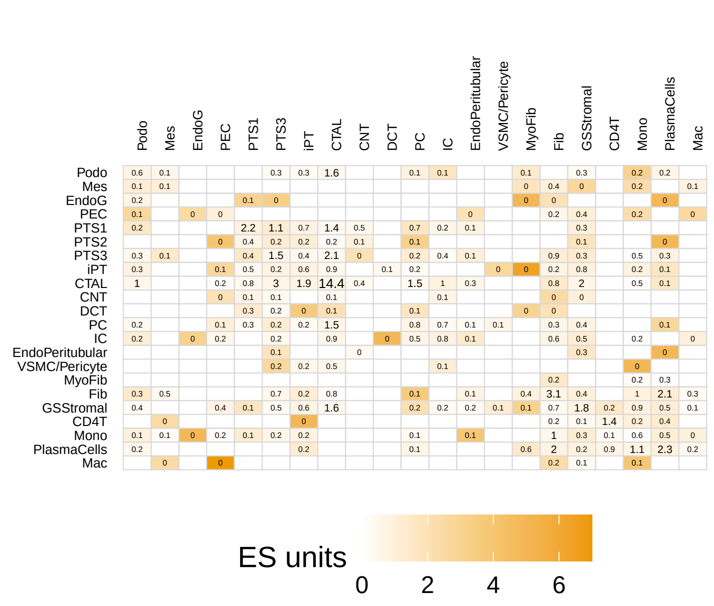


HK3444
Center SCP: Intra−golgi vesicle tethering
Peripheral SCP: Vesicle tethering at cis−Golgi
Numbers: # involved center−peripheral spot pairs [%]

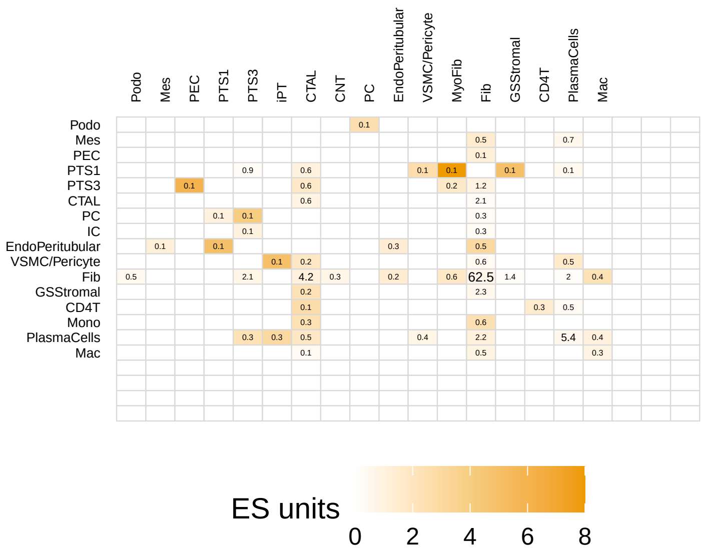


HK3432
Center SCP: Early endocytic vesicle and endosome tethering
Peripheral SCP: Early endocytic vesicle and endosome fusion
Numbers: # involved center−peripheral spot pairs [%]

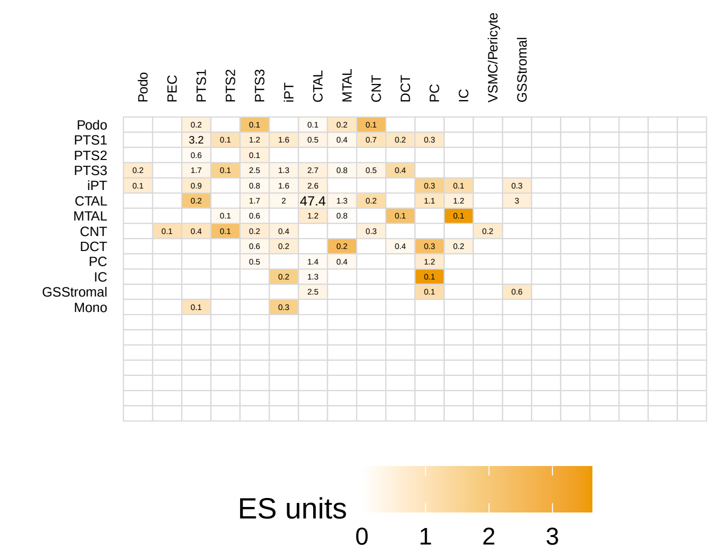


HK2874
Center SCP: Eukaryotic intraflagellar transport
Peripheral SCP: Mother centriole and distal appendage transformation into basal body
Numbers: # involved center−peripheral spot pairs [%]

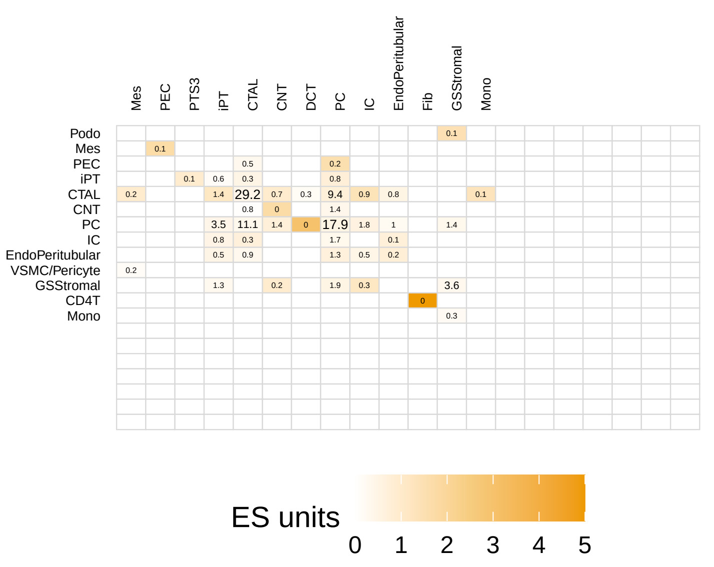


HK2871
Center SCP: Ciliary transition zone organization
Peripheral SCP: Eukaryotic motile axoneme organization
Numbers: # involved center−peripheral spot pairs [%]

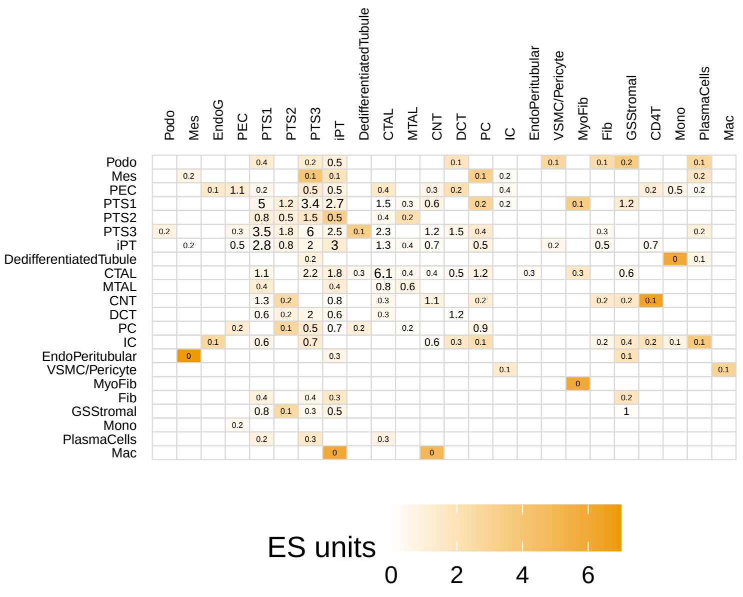

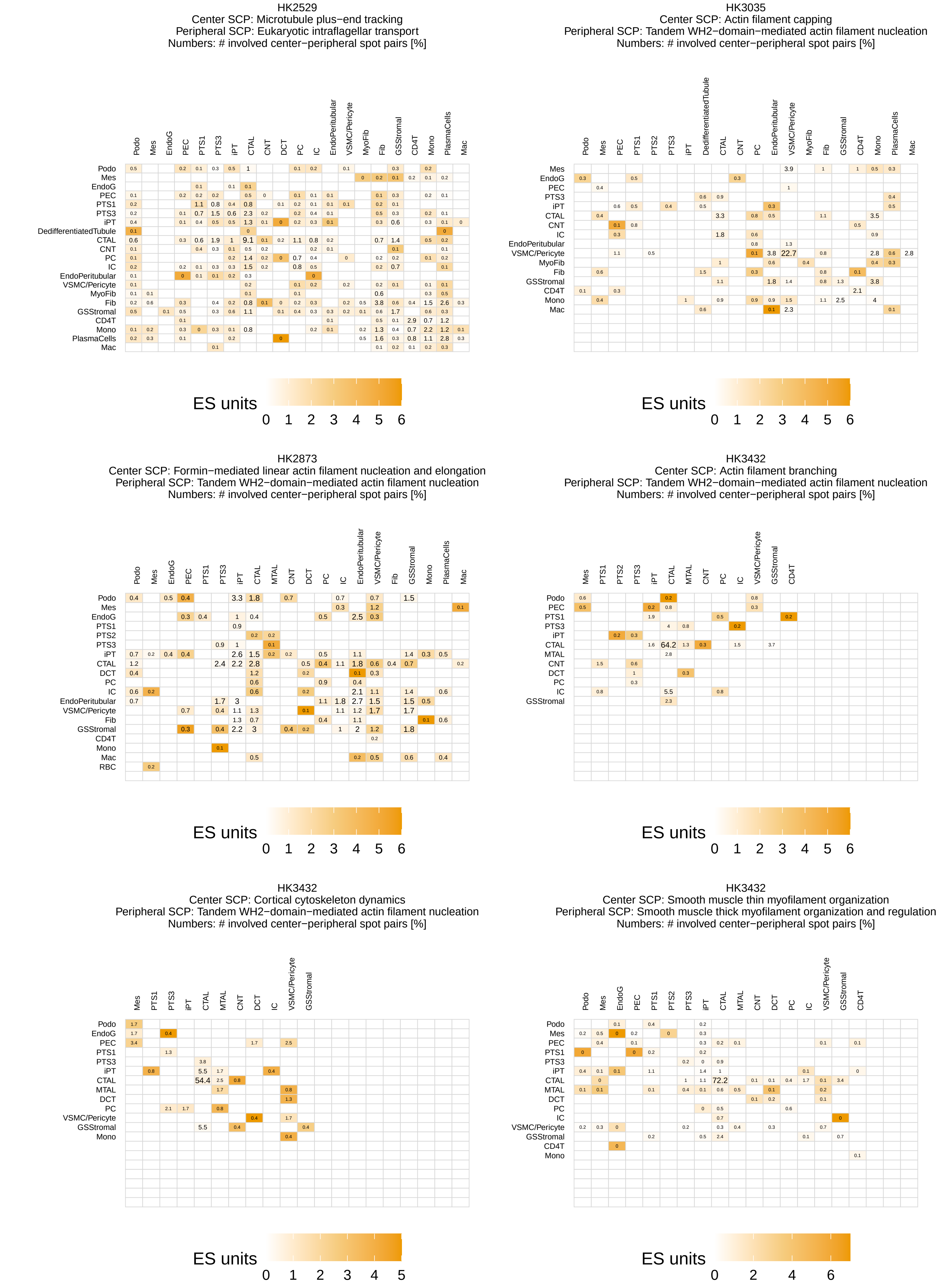
HK2529
Center SCP: Microtubule plus−end tracking
Peripheral SCP: Eukaryotic intraflagellar transport
Numbers: # involved center−peripheral spot pairs [%]
Podo Mes EndoG PEC PTS1 PTS3 iPT CTAL CNT DCT PC IC EndoPeritubular VSMC/Pericyte MyoFib Fib GSStromal CD4T Mono PlasmaCells Mac
Podo Mes EndoG PEC PTS1 PTS3 iPT DedifferentiatedTubule CTAL CNT PC IC EndoPeritubular VSMC/Pericyte MyoFib Fib GSStromal CD4T Mono PlasmaCells Mac
ES units
0 1 2 3 4 5 6
HK3035
Center SCP: Actin filament capping
Peripheral SCP: Tandem WH2−domain−mediated actin filament nucleation
Numbers: # involved center−peripheral spot pairs [%]
Podo Mes PEC PTS1 PTS2 PTS3 iPT DedifferentiatedTubule CTAL CNT PC EndoPeritubular VSMC/Pericyte MyoFib Fib GSStromal CD4T Mono PlasmaCells Mac
Mes EndoG PEC PTS3 iPT CTAL CNT IC EndoPeritubular VSMC/Pericyte MyoFib Fib GSStromal CD4T Mono Mac
ES units
0 1 2 3 4 5 6
HK2873
Center SCP: Formin−mediated linear actin filament nucleation and elongation
Peripheral SCP: Tandem WH2−domain−mediated actin filament nucleation
Numbers: # involved center−peripheral spot pairs [%]
Podo Mes EndoG PEC PTS1 PTS3 iPT CTAL MTAL CNT DCT PC IC EndoPeritubular VSMC/Pericyte Fib GSStromal Mono PlasmaCells Mac
Podo Mes EndoG PTS1 PTS2 PTS3 iPT CTAL DCT PC IC EndoPeritubular VSMC/Pericyte Fib GSStromal CD4T Mono Mac RBC
ES units
0 1 2 3 4 5 6
HK3432
Center SCP: Actin filament branching
Peripheral SCP: Tandem WH2−domain−mediated actin filament nucleation
Numbers: # involved center−peripheral spot pairs [%]
Mes PTS1 PTS2 PTS3 iPT CTAL MTAL CNT PC IC VSMC/Pericyte GSStromal CD4T
Podo PEC PTS1 PTS3 iPT CTAL MTAL CNT DCT PC IC GSStromal
ES units
0 1 2 3 4 5 6
HK3432
Center SCP: Cortical cytoskeleton dynamics
Peripheral SCP: Tandem WH2−domain−mediated actin filament nucleation
Numbers: # involved center−peripheral spot pairs [%]
Mes PTS1 PTS3 iPT CTAL MTAL CNT DCT IC VSMC/Pericyte GSStromal
Podo EndoG PEC PTS1 PTS3 iPT CTAL MTAL DCT PC VSMC/Pericyte GSStromal Mono
ES units
0 1 2 3 4 5
HK3432
Center SCP: Smooth muscle thin myofilament organization
Peripheral SCP: Smooth muscle thick myofilament organization and regulation
Numbers: # involved center−peripheral spot pairs [%]
Podo Mes EndoG PEC PTS1 PTS2 PTS3 iPT CTAL MTAL CNT DCT PC IC VSMC/Pericyte GSStromal CD4T
Podo Mes PEC PTS1 PTS3 iPT CTAL MTAL DCT PC IC VSMC/Pericyte GSStromal CD4T Mono
ES units
0 2 4 6

HK2844
Center SCP: Nuclear intermediate filaments
Peripheral SCP: LINC complex−mediated nuclear−cytoskeletal coupling
Numbers: # involved center−peripheral spot pairs [%]

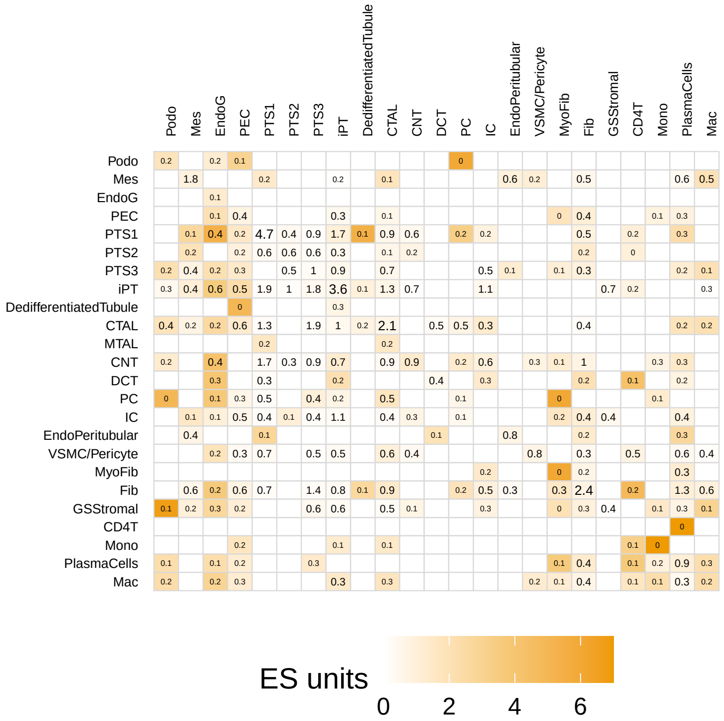


HK2852
Center SCP: Hemidesmosome organization
Peripheral SCP: Epithelial intermediate filament dynamics
Numbers: # involved center−peripheral spot pairs [%]

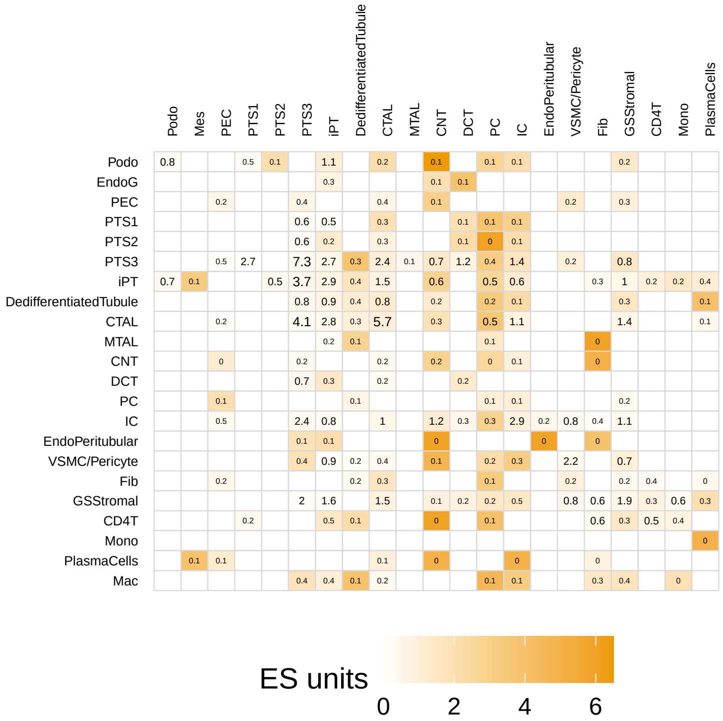


HK3035
Center SCP: Endoplasmic and sarcoplasmic calcium import, export and storage
Peripheral SCP: Endoplasmic reticulum−plasma membrane contact site formation
Numbers: # involved center−peripheral spot pairs [%]

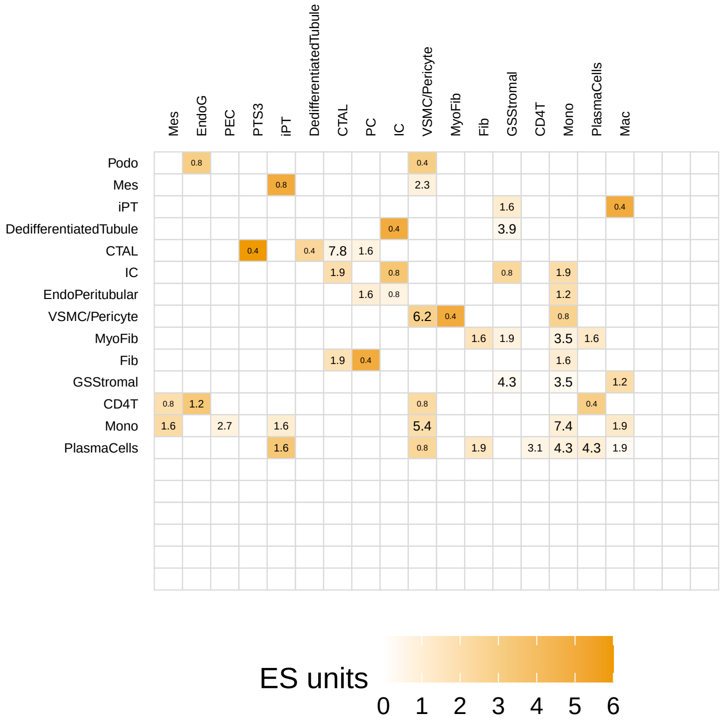

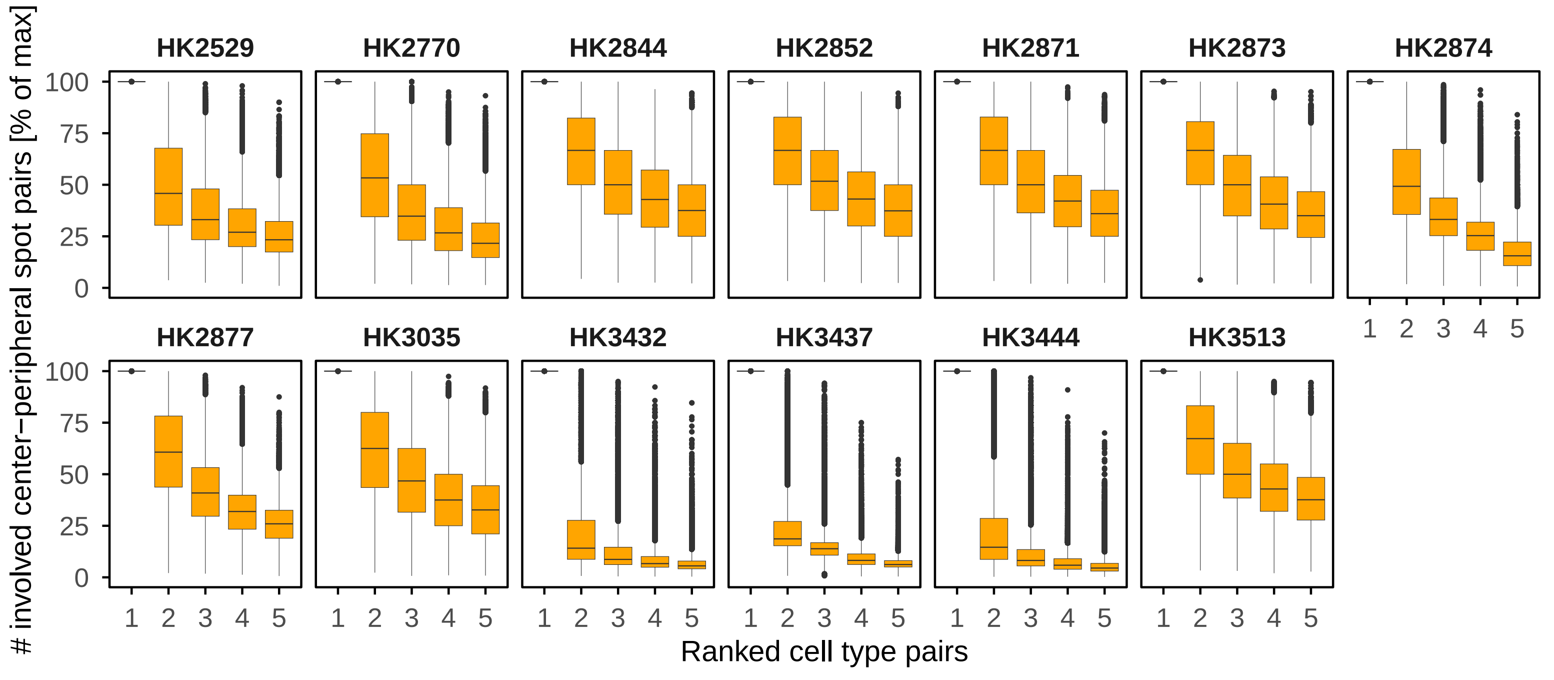

HK2529
HK2770
HK2844
HK2852
HK2871
HK2873
HK2874
HK2877
HK3035
HK3432
HK3437
HK3444
HK3513
### involved center−peripheral spot pairs [% of max]
Ranked cell type pairs


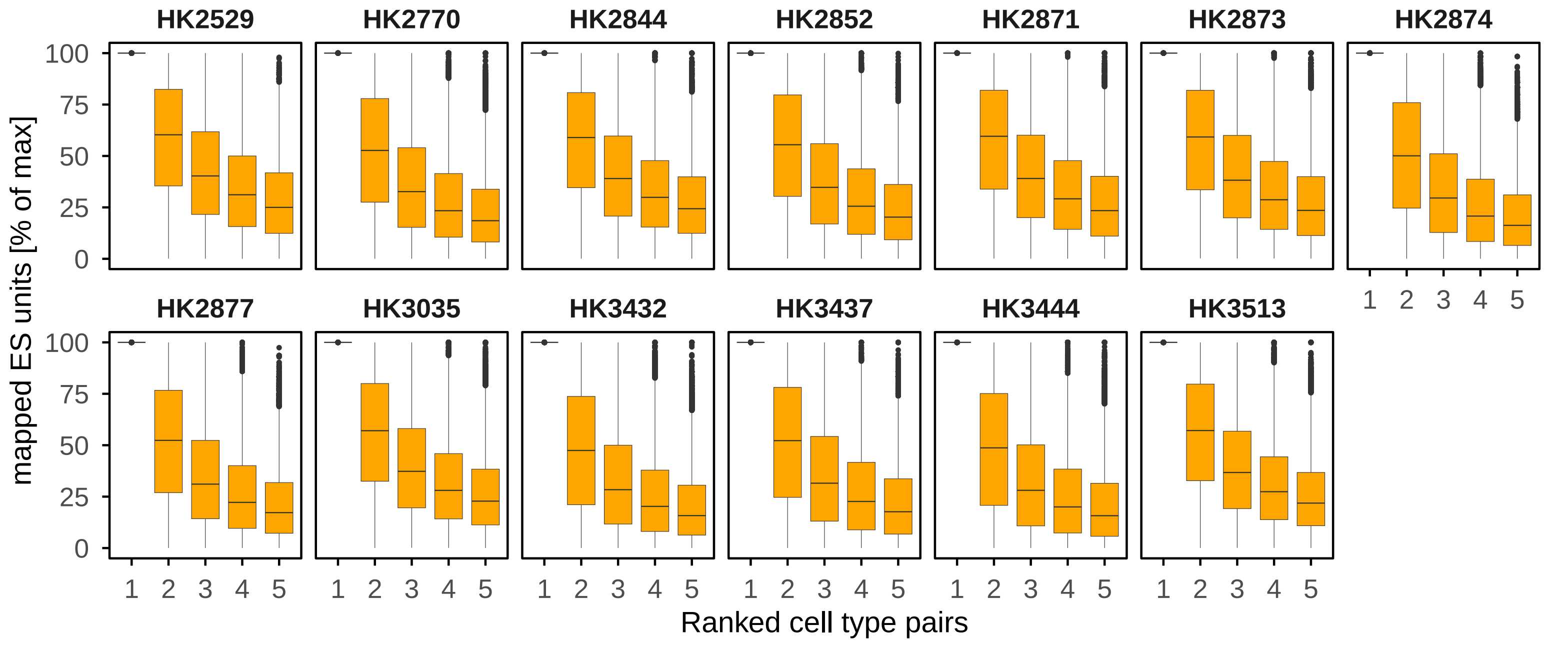

HK2529
HK2770
HK2844
HK2852
HK2871
HK2873
HK2874
HK2877
HK3035
HK3432
HK3437
HK3444
HK3513
mapped ES units [% of max]
Ranked cell type pairs


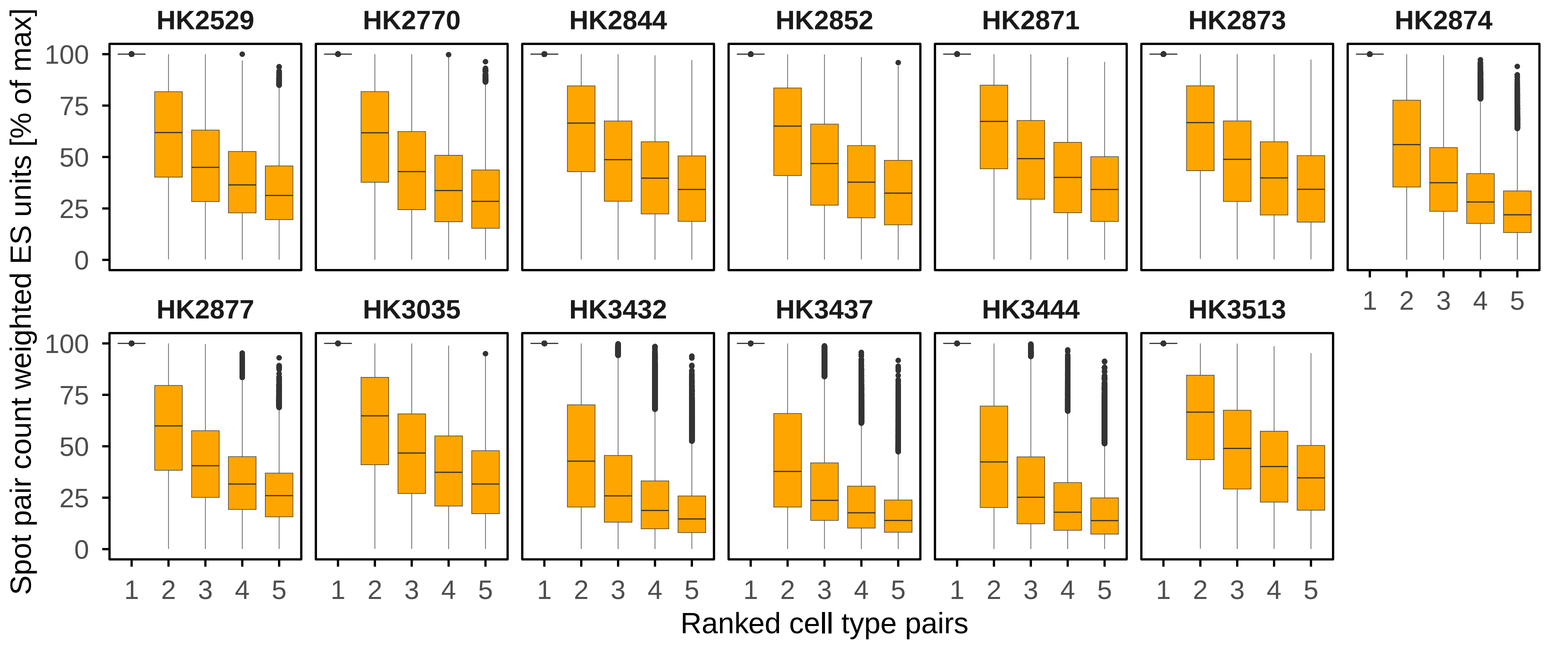

HK2529
HK2770
HK2844
HK2852
HK2871
HK2873
HK2874
HK2877
HK3035
HK3432
HK3437
HK3444
HK3513
Spot pair count weighted ES units [% of max]
Ranked cell type pairs

**Supplementary Figure 7. Missing cell-type-pair selectivity for identified SCP pairs. (A)** We repeated our analysis pipeline, but this time averaged the results for each combination of center and peripheral cell type annotations independently. Mapped (not normalized) ES units between all cell type pairs are shown for selected SCP pairs. Note that the ES units were averaged across the whole neighborhood, while higher numbers of mapped ES units can be obtained by averaging them only across spots that express the target SCP (see main figure. 8). The heatmap shown in figure 8A is part of these figures as well. **(B)** To systematically analyze potential cell-type selectivity, we identified that cell type pair for each SCP pair that has the highest number of center spots with the cross-cellular SCP pair mapping in question. We then normalized the number of center spots for all cell type pairs towards the identified highest number of center spots, followed by ranking of all cell type pairs --still for the same SCP pair- by the normalized center spot counts. For each tissue slice, we then combined the cell type pairs associated with all SCP pairs into groups of same ranks (ignoring which cell types are involved) to show the normalized center spot counts associated with the top 5 ranks (top panel). We repeated the same analysis with the mapped ES units (middle panel) and the product of the number of center spots and the mapped ES units (bottom panel and main figure 9B).

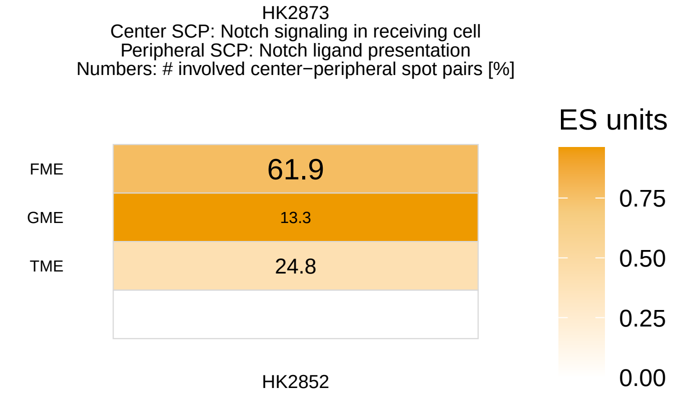
HK2873
Center SCP: Notch signaling in receiving cell
Peripheral SCP: Notch ligand presentation
Numbers: # involved center−peripheral spot pairs [%]
ES units
FME 61.9
GME 13.3
TME 24.8
0.75
0.50
0.25
0.00

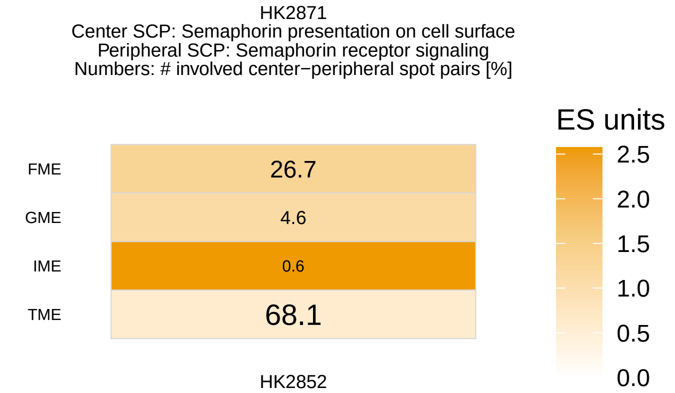
HK2871
Center SCP: Semaphorin presentation on cell surface
Peripheral SCP: Semaphorin receptor signaling
Numbers: # involved center−peripheral spot pairs [%]
ES units
FME 26.7
GME 4.6
IME 0.6
TME 68.1
2.5
2.0
1.5
1.0
0.5
0.0

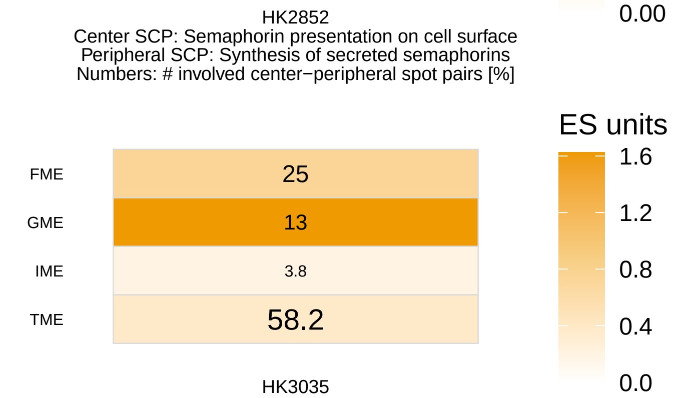
HK2852
Center SCP: Semaphorin presentation on cell surface
Peripheral SCP: Synthesis of secreted semaphorins
Numbers: # involved center−peripheral spot pairs [%]
ES units
FME 25
GME 13
IME 3.8
TME 58.2
1.6
1.2
0.8
0.4
0.0

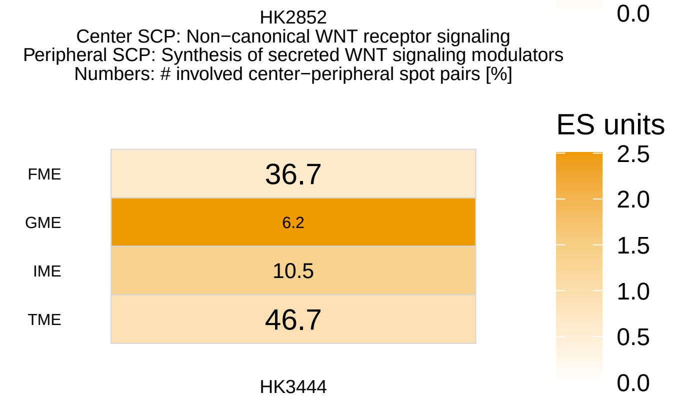
HK2852
Center SCP: Non−canonical WNT receptor signaling
Peripheral SCP: Synthesis of secreted WNT signaling modulators
Numbers: # involved center−peripheral spot pairs [%]
ES units
FME 36.7
GME 6.2
IME 10.5
TME 46.7
2.5
2.0
1.5
1.0
0.5
0.0

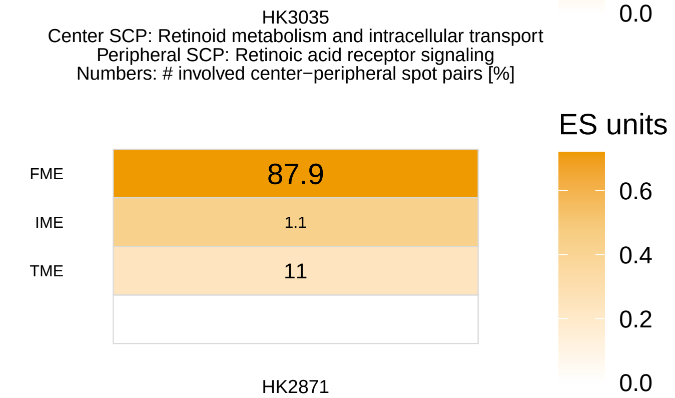
HK3035
Center SCP: Retinoid metabolism and intracellular transport
Peripheral SCP: Retinoic acid receptor signaling
Numbers: # involved center−peripheral spot pairs [%]
ES units
FME 87.9
IME 1.1
TME 11
0.6
0.4
0.2
0.0

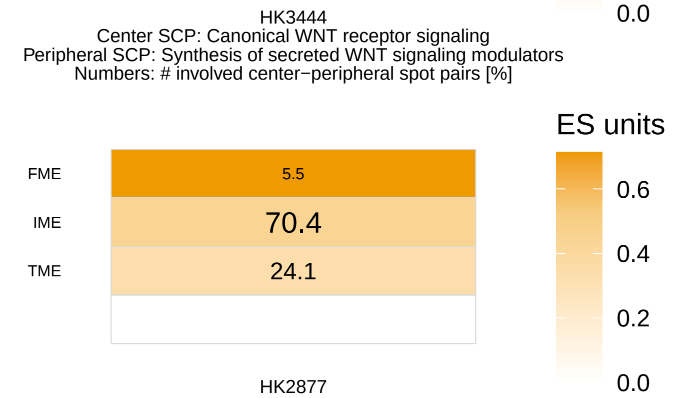
HK3444
Center SCP: Canonical WNT receptor signaling
Peripheral SCP: Synthesis of secreted WNT signaling modulators
Numbers: # involved center−peripheral spot pairs [%]
ES units
FME 5.5
IME 70.4
TME 24.1
0.6
0.4
0.2
0.0

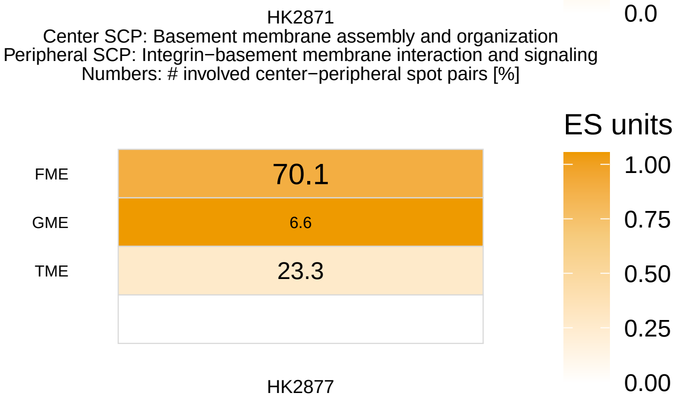
HK2871
Center SCP: Basement membrane assembly and organization
Peripheral SCP: Integrin−basement membrane interaction and signaling
Numbers: # involved center−peripheral spot pairs [%]
ES units
FME 70.1
GME 6.6
TME 23.3
1.00
0.75
0.50
0.25
0.00

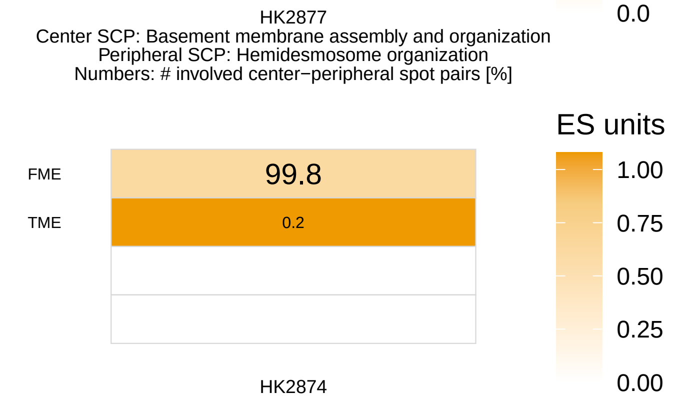
HK2877
Center SCP: Basement membrane assembly and organization
Peripheral SCP: Hemidesmosome organization
Numbers: # involved center−peripheral spot pairs [%]
ES units
FME 99.8
TME 0.2
1.00
0.75
0.50
0.25
0.00

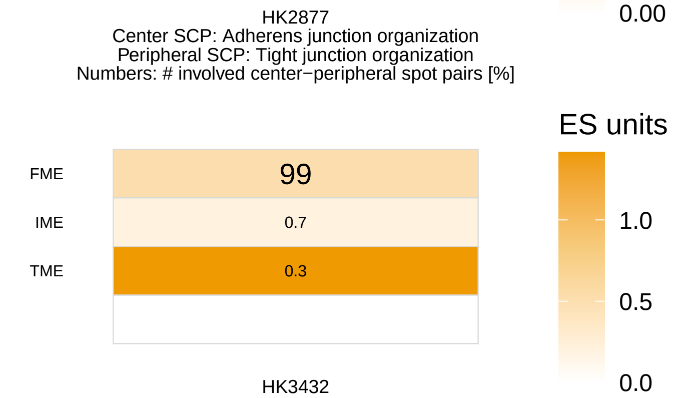
HK2877
Center SCP: Adherens junction organization
Peripheral SCP: Tight junction organization
Numbers: # involved center−peripheral spot pairs [%]
ES units
FME 99
IME 0.7
TME 0.3
1.0
0.5
0.0

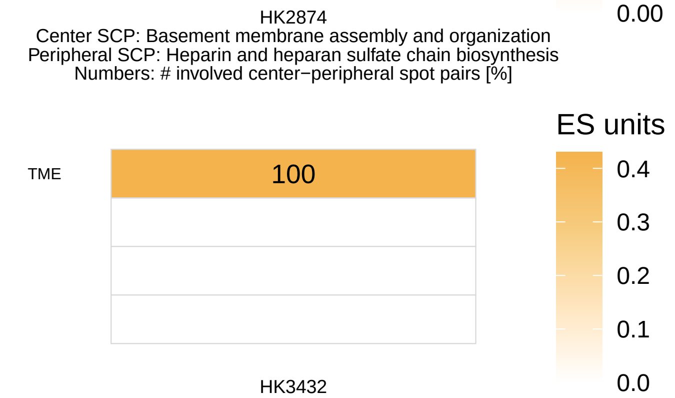
HK2874
Center SCP: Basement membrane assembly and organization
Peripheral SCP: Heparin and heparan sulfate chain biosynthesis
Numbers: # involved center−peripheral spot pairs [%]
ES units
TME 100
0.4
0.3
0.2
0.1
0.0

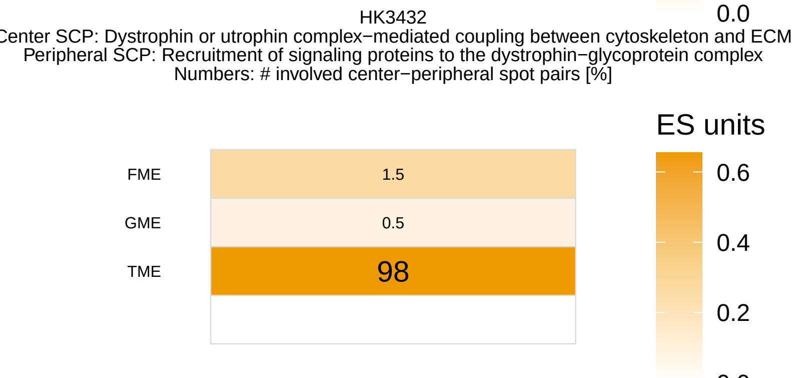
HK3432
Center SCP: Dystrophin or utrophin complex−mediated coupling between cytoskeleton and ECM
Peripheral SCP: Recruitment of signaling proteins to the dystrophin−glycoprotein complex
Numbers: # involved center−peripheral spot pairs [%]
ES units
FME 1.5
GME 0.5
TME 98
0.6
0.4
0.2

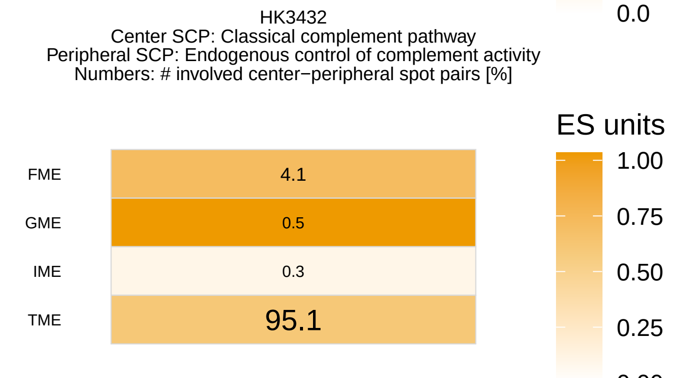
HK3432
Center SCP: Classical complement pathway
Peripheral SCP: Endogenous control of complement activity
Numbers: # involved center−peripheral spot pairs [%]
ES units
FME 4.1
GME 0.5
IME 0.3
TME 95.1
1.00
0.75
0.50
0.25

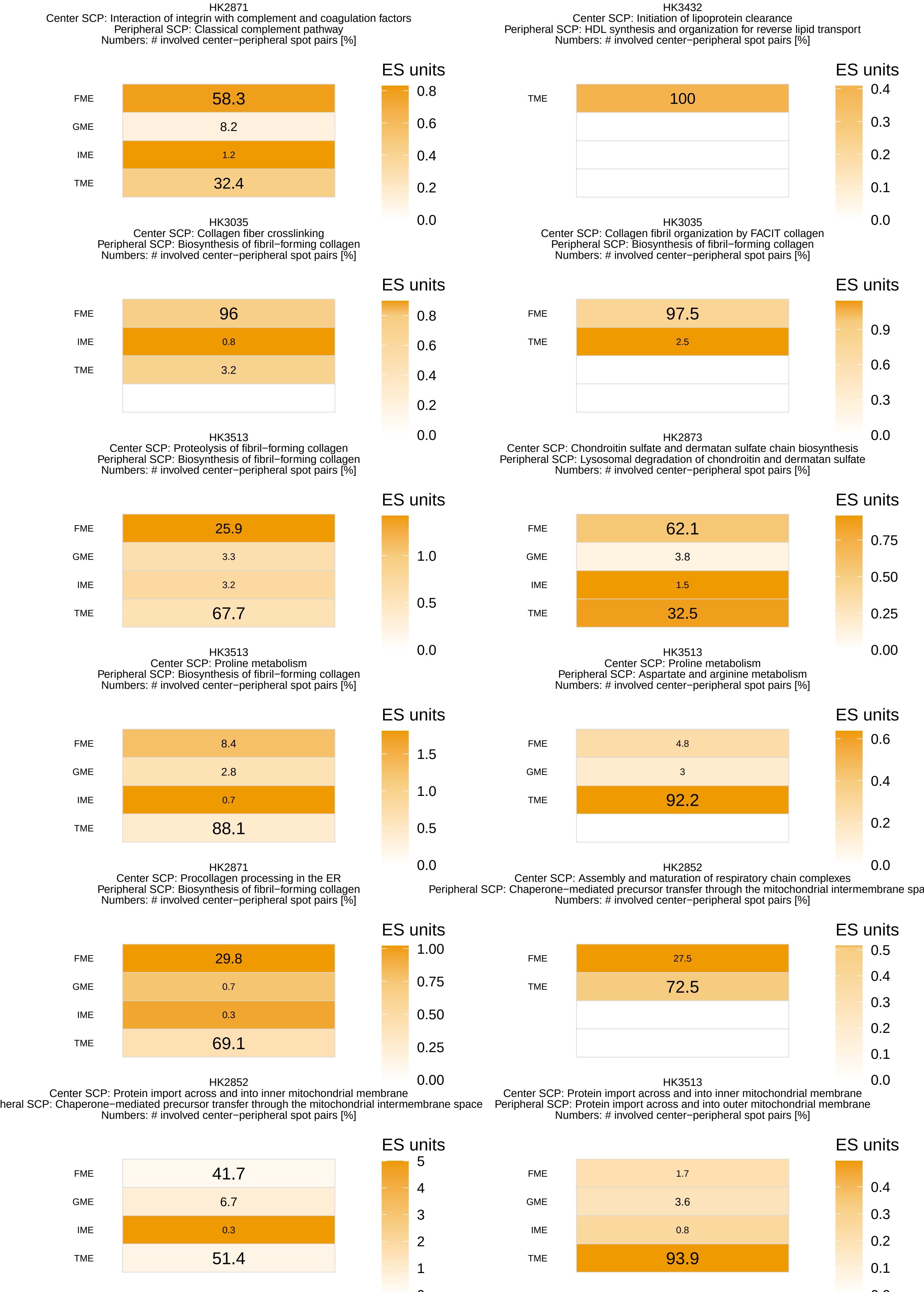
HK2871
Center SCP: Interaction of integrin with complement and coagulation factors
Peripheral SCP: Classical complement pathway
Numbers: # involved center−peripheral spot pairs [%]
ES units
FME 58.3
GME 8.2
IME 1.2
TME 32.4
0.8 0.6 0.4 0.2 0.0
HK3432
Center SCP: Initiation of lipoprotein clearance
Peripheral SCP: HDL synthesis and organization for reverse lipid transport
Numbers: # involved center−peripheral spot pairs [%]
ES units
TME 100
0.4 0.3 0.2 0.1 0.0
HK3035
Center SCP: Collagen fiber crosslinking
Peripheral SCP: Biosynthesis of fibril−forming collagen
Numbers: # involved center−peripheral spot pairs [%]
ES units
FME 96
IME 0.8
TME 3.2
0.8 0.6 0.4 0.2 0.0
HK3035
Center SCP: Collagen fibril organization by FACIT collagen
Peripheral SCP: Biosynthesis of fibril−forming collagen
Numbers: # involved center−peripheral spot pairs [%]
ES units
FME 97.5
TME 2.5
0.9 0.6 0.3 0.0
HK3513
Center SCP: Proteolysis of fibril−forming collagen
Peripheral SCP: Biosynthesis of fibril−forming collagen
Numbers: # involved center−peripheral spot pairs [%]
ES units
FME 25.9
GME 3.3
IME 3.2
TME 67.7
1.0 0.5 0.0
HK2873
Center SCP: Chondroitin sulfate and dermatan sulfate chain biosynthesis
Peripheral SCP: Lysosomal degradation of chondroitin and dermatan sulfate
Numbers: # involved center−peripheral spot pairs [%]
ES units
FME 62.1
GME 3.8
IME 1.5
TME 32.5
0.75 0.50 0.25 0.00
HK3513
Center SCP: Proline metabolism
Peripheral SCP: Biosynthesis of fibril−forming collagen
Numbers: # involved center−peripheral spot pairs [%]
ES units
FME 8.4
GME 2.8
IME 0.7
TME 88.1
1.5 1.0 0.5 0.0
HK3513
Center SCP: Proline metabolism
Peripheral SCP: Aspartate and arginine metabolism
Numbers: # involved center−peripheral spot pairs [%]
ES units
FME 4.8
GME 3
TME 92.2
0.6 0.4 0.2 0.0
HK2871
Center SCP: Procollagen processing in the ER
Peripheral SCP: Biosynthesis of fibril−forming collagen
Numbers: # involved center−peripheral spot pairs [%]
ES units
FME 29.8
GME 0.7
IME 0.3
TME 69.1
1.00 0.75 0.50 0.25 0.00
HK2852
Center SCP: Assembly and maturation of respiratory chain complexes
Peripheral SCP: Chaperone−mediated precursor transfer through the mitochondrial intermembrane spa
Numbers: # involved center−peripheral spot pairs [%]
ES units
FME 27.5
TME 72.5
0.5 0.4 0.3 0.2 0.1 0.0
HK2852
Center SCP: Protein import across and into inner mitochondrial membrane
heral SCP: Chaperone−mediated precursor transfer through the mitochondrial intermembrane space
Numbers: # involved center−peripheral spot pairs [%]
ES units
FME 41.7
GME 6.7
IME 0.3
TME 51.4
5 4 3 2 1
HK3513
Center SCP: Protein import across and into inner mitochondrial membrane
Peripheral SCP: Protein import across and into outer mitochondrial membrane
Numbers: # involved center−peripheral spot pairs [%]
ES units
FME 1.7
GME 3.6
IME 0.8
TME 93.9
0.4 0.3 0.2 0.1

HK3444
Center SCP: Protein import across and into inner mitochondrial membrane
Peripheral SCP: MICOS−dependent maintenance of mitochondrial cristae architecture
Numbers: # involved center−peripheral spot pairs [%]

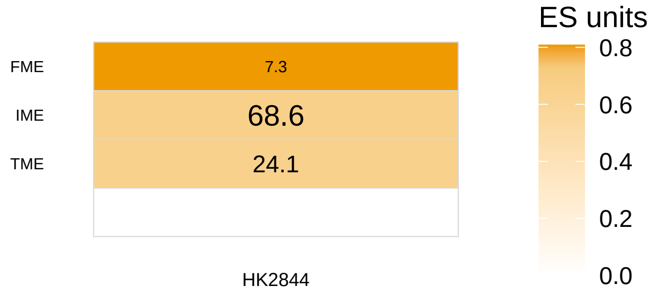


HK3035
Center SCP: Serine and glycine metabolism
Peripheral SCP: Mitochondrial translation elongation
Numbers: # involved center−peripheral spot pairs [%]

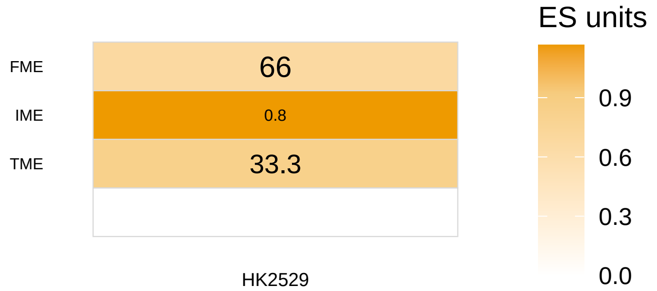


HK2844
Center SCP: Mitochondrial translation termination
Peripheral SCP: Mitochondrial translation elongation
Numbers: # involved center−peripheral spot pairs [%]

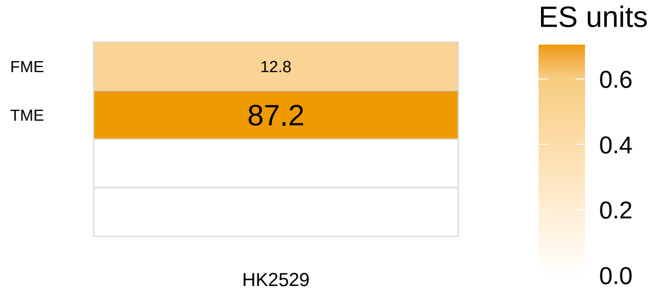


HK2529
Center SCP: Electron transport chain
Peripheral SCP: Mitochondrial ATP−synthesis coupled to proton gradient
Numbers: # involved center−peripheral spot pairs [%]

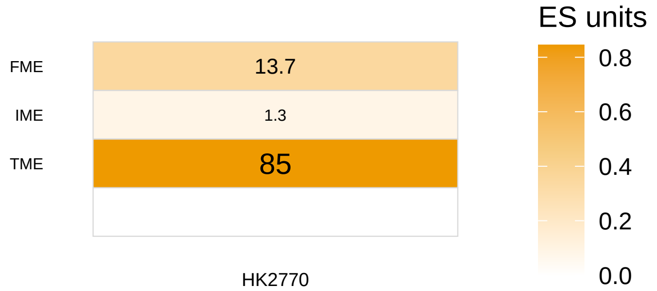


HK2529
Center SCP: Mitochondrial ATP−synthesis coupled to proton gradient
Peripheral SCP: Mitochondrial import and oxidative decarboxylation of pyruvate
Numbers: # involved center−peripheral spot pairs [%]

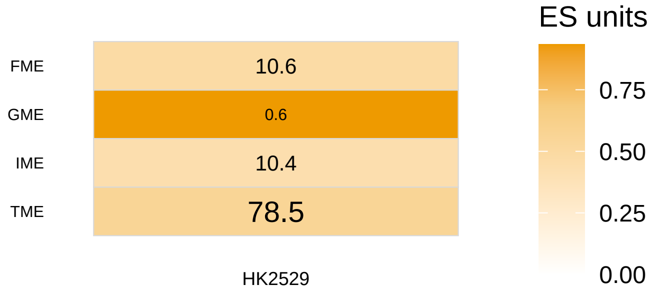


HK2770
Center SCP: MICOS−dependent maintenance of mitochondrial cristae architecture
Peripheral SCP: Mitochondrial ATP−synthesis coupled to proton gradient
Numbers: # involved center−peripheral spot pairs [%]

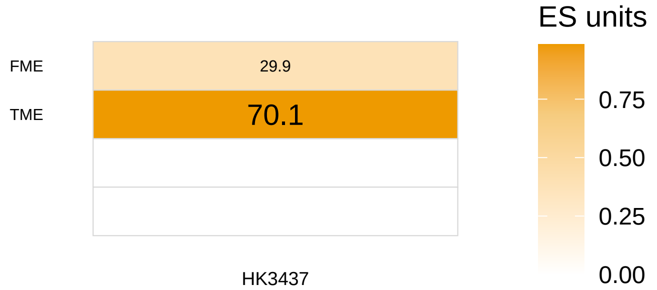


HK2529
Center SCP: Beta−oxidation
Peripheral SCP: Fatty acid activation
Numbers: # involved center−peripheral spot pairs [%]

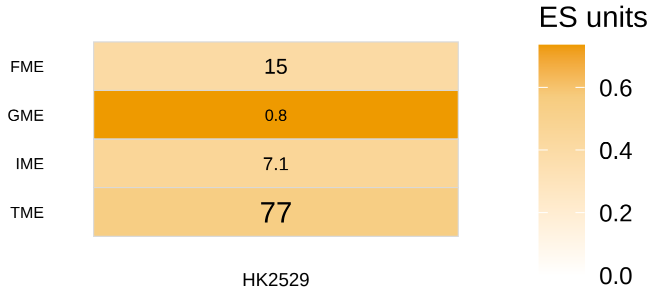


HK3437
Center SCP: Beta−oxidation
Peripheral SCP: Citric acid cycle
Numbers: # involved center−peripheral spot pairs [%]

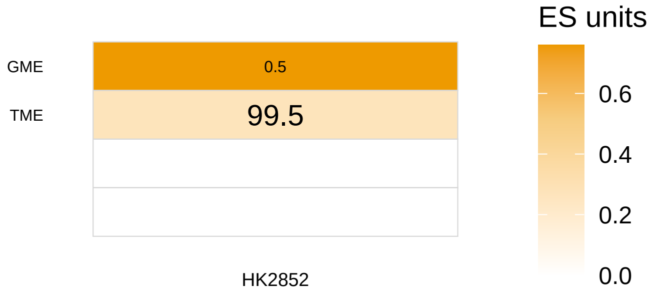


HK2529
Center SCP: Glycolysis and gluconeogenesis
Peripheral SCP: Mitochondrial import and oxidative decarboxylation of pyruvate
Numbers: # involved center−peripheral spot pairs [%]

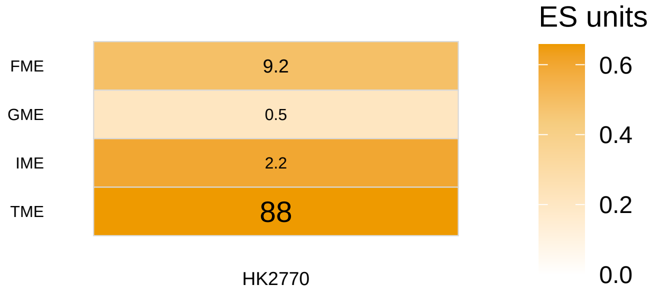


HK2852
Center SCP: Mitochondrial import and oxidative decarboxylation of pyruvate
Peripheral SCP: Malate aspartate shuttle
Numbers: # involved center−peripheral spot pairs [%]

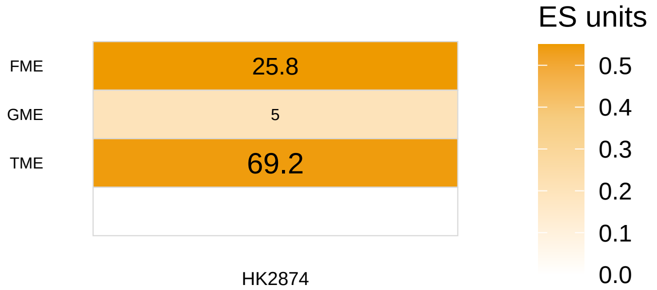


HK2770
Center SCP: Carnitine shuttle
Peripheral SCP: Carnitine biosynthesis and transmembrane transport
Numbers: # involved center−peripheral spot pairs [%]

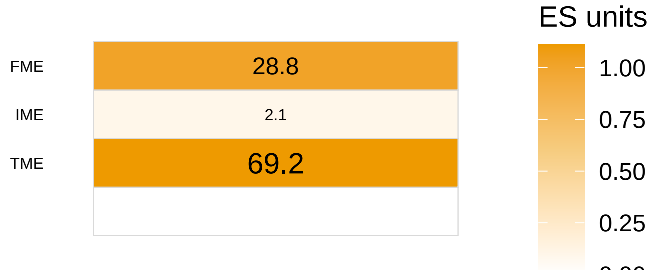


HK2874
Center SCP: Beta−oxidation
Peripheral SCP: Ketone body metabolism and transport
Numbers: # involved center−peripheral spot pairs [%]

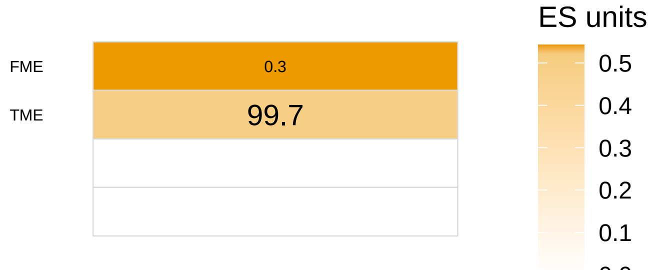

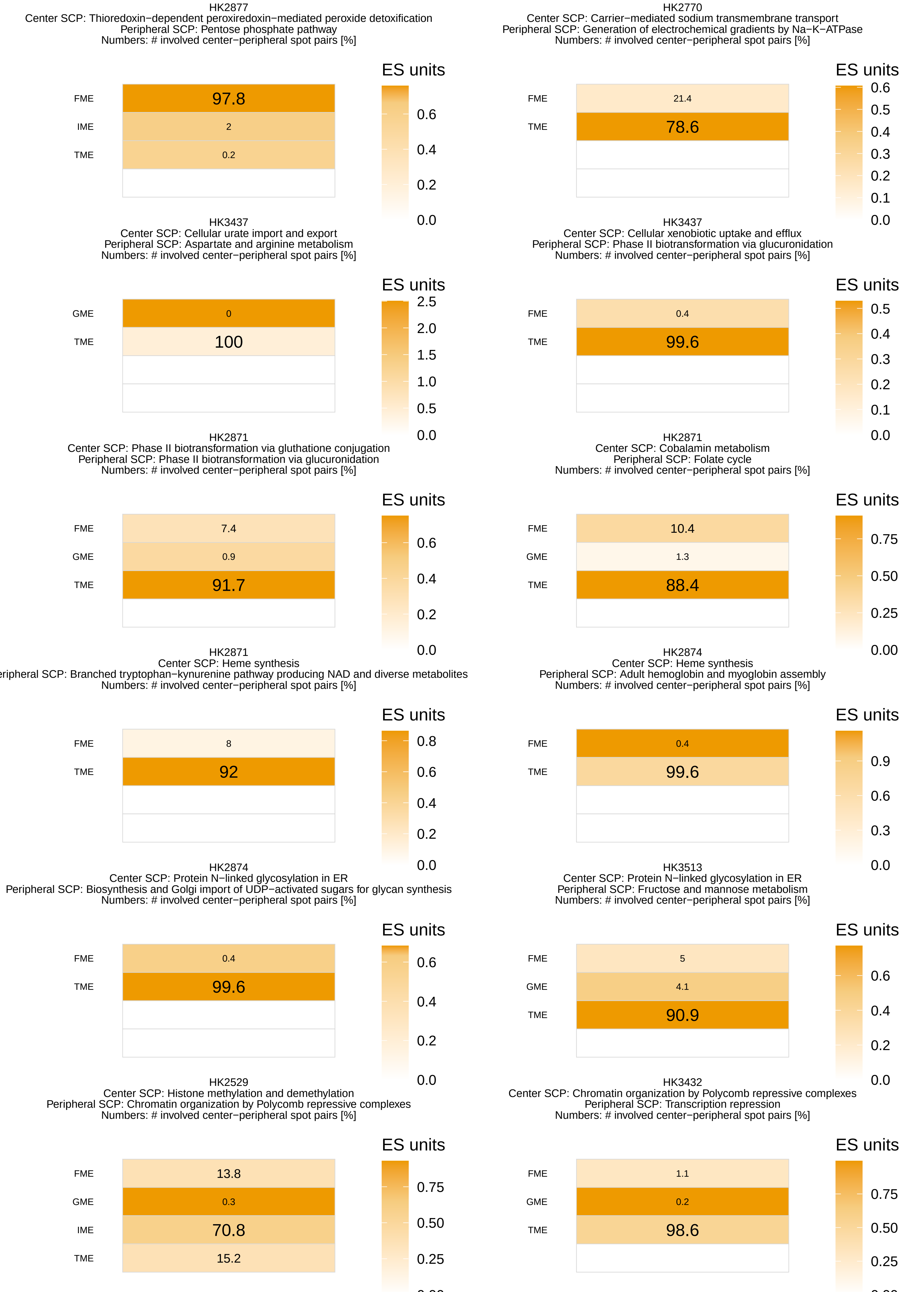
HK2877
Center SCP: Thioredoxin−dependent peroxiredoxin−mediated peroxide detoxification
Peripheral SCP: Pentose phosphate pathway
Numbers: # involved center−peripheral spot pairs [%]
FME 97.8
IME 2
TME 0.2
ES units 0.6 0.4 0.2 0.0
HK2770
Center SCP: Carrier−mediated sodium transmembrane transport
Peripheral SCP: Generation of electrochemical gradients by Na−K−ATPase
Numbers: # involved center−peripheral spot pairs [%]
FME 21.4
TME 78.6
ES units 0.6 0.5 0.4 0.3 0.2 0.1 0.0
HK3437
Center SCP: Cellular urate import and export
Peripheral SCP: Aspartate and arginine metabolism
Numbers: # involved center−peripheral spot pairs [%]
GME 0
TME 100
ES units 2.5 2.0 1.5 1.0 0.5 0.0
HK3437
Center SCP: Cellular xenobiotic uptake and efflux
Peripheral SCP: Phase II biotransformation via glucuronidation
Numbers: # involved center−peripheral spot pairs [%]
FME 0.4
TME 99.6
ES units 0.5 0.4 0.3 0.2 0.1 0.0
HK2871
Center SCP: Phase II biotransformation via gluthatione conjugation
Peripheral SCP: Phase II biotransformation via glucuronidation
Numbers: # involved center−peripheral spot pairs [%]
FME 7.4
GME 0.9
TME 91.7
ES units 0.6 0.4 0.2 0.0
HK2871
Center SCP: Cobalamin metabolism
Peripheral SCP: Folate cycle
Numbers: # involved center−peripheral spot pairs [%]
FME 10.4
GME 1.3
TME 88.4
ES units 0.75 0.50 0.25 0.00
HK2871
Center SCP: Heme synthesis
eripheral SCP: Branched tryptophan−kynurenine pathway producing NAD and diverse metabolites
Numbers: # involved center−peripheral spot pairs [%]
FME 8
TME 92
ES units 0.8 0.6 0.4 0.2 0.0
HK2874
Center SCP: Heme synthesis
Peripheral SCP: Adult hemoglobin and myoglobin assembly
Numbers: # involved center−peripheral spot pairs [%]
FME 0.4
TME 99.6
ES units 0.9 0.6 0.3 0.0
HK2874
Center SCP: Protein N−linked glycosylation in ER
Peripheral SCP: Biosynthesis and Golgi import of UDP−activated sugars for glycan synthesis
Numbers: # involved center−peripheral spot pairs [%]
FME 0.4
TME 99.6
ES units 0.6 0.4 0.2 0.0
HK3513
Center SCP: Protein N−linked glycosylation in ER
Peripheral SCP: Fructose and mannose metabolism
Numbers: # involved center−peripheral spot pairs [%]
FME 5
GME 4.1
TME 90.9
ES units 0.6 0.4 0.2 0.0
HK2529
Center SCP: Histone methylation and demethylation
Peripheral SCP: Chromatin organization by Polycomb repressive complexes
Numbers: # involved center−peripheral spot pairs [%]
FME 13.8
GME 0.3
IME 70.8
TME 15.2
ES units 0.75 0.50 0.25
HK3432
Center SCP: Chromatin organization by Polycomb repressive complexes
Peripheral SCP: Transcription repression
Numbers: # involved center−peripheral spot pairs [%]
FME 1.1
GME 0.2
TME 98.6
ES units 0.75 0.50 0.25

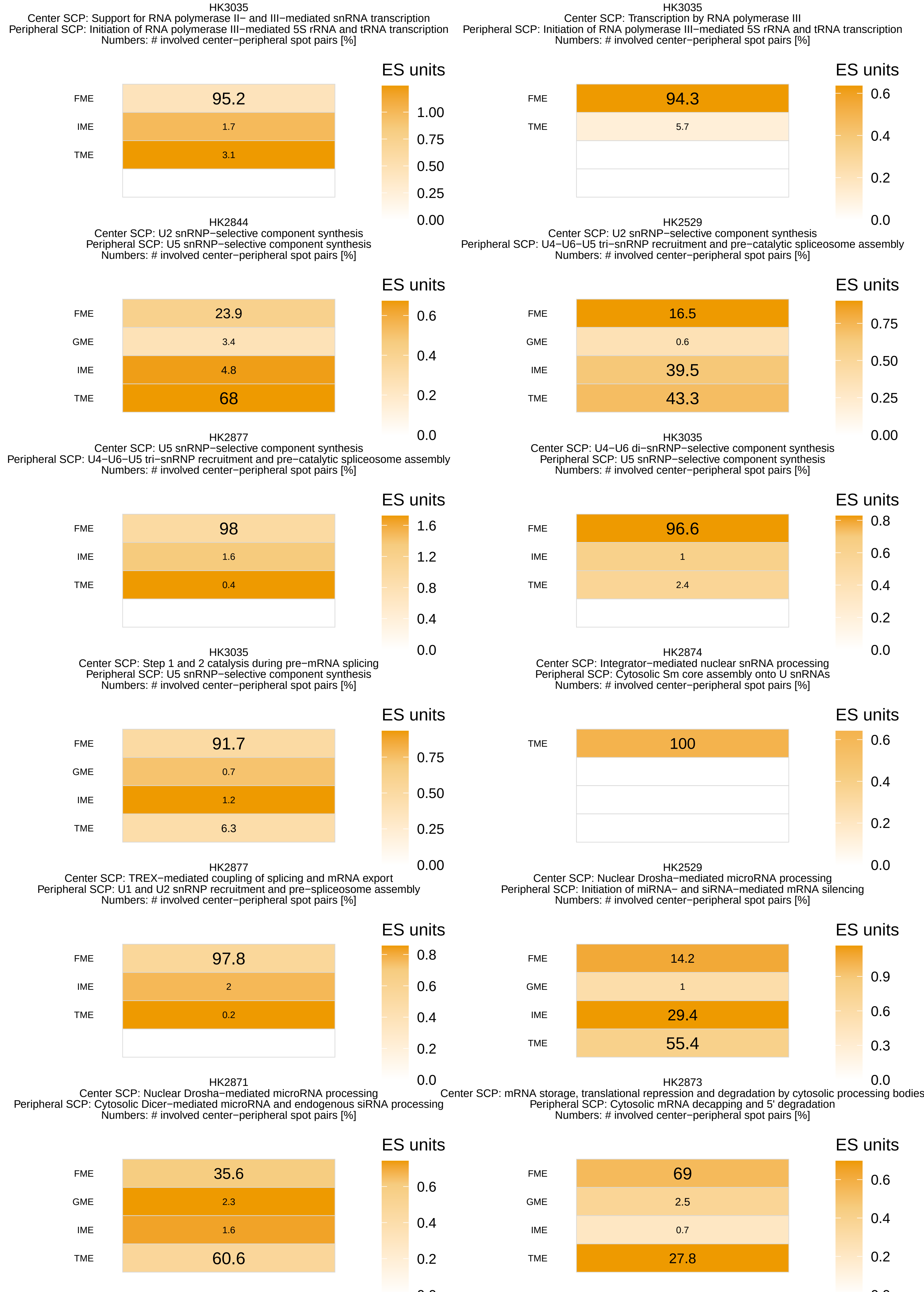
HK3035
Center SCP: Support for RNA polymerase II− and III−mediated snRNA transcription
Peripheral SCP: Initiation of RNA polymerase III−mediated 5S rRNA and tRNA transcription
Numbers: # involved center−peripheral spot pairs [%]
ES units
FME 95.2
IME 1.7
TME 3.1
1.00
0.75
0.50
0.25
0.00
HK3035
Center SCP: Transcription by RNA polymerase III
Peripheral SCP: Initiation of RNA polymerase III−mediated 5S rRNA and tRNA transcription
Numbers: # involved center−peripheral spot pairs [%]
ES units
FME 94.3
TME 5.7
0.6
0.4
0.2
0.0
HK2844
Center SCP: U2 snRNP−selective component synthesis
Peripheral SCP: U5 snRNP−selective component synthesis
Numbers: # involved center−peripheral spot pairs [%]
ES units
FME 23.9
GME 3.4
IME 4.8
TME 68
0.6
0.4
0.2
0.0
HK2529
Center SCP: U2 snRNP−selective component synthesis
Peripheral SCP: U4−U6−U5 tri−snRNP recruitment and pre−catalytic spliceosome assembly
Numbers: # involved center−peripheral spot pairs [%]
ES units
FME 16.5
GME 0.6
IME 39.5
TME 43.3
0.75
0.50
0.25
0.00
HK2877
Center SCP: U5 snRNP−selective component synthesis
Peripheral SCP: U4−U6−U5 tri−snRNP recruitment and pre−catalytic spliceosome assembly
Numbers: # involved center−peripheral spot pairs [%]
ES units
FME 98
IME 1.6
TME 0.4
1.6
1.2
0.8
0.4
0.0
HK3035
Center SCP: U4−U6 di−snRNP−selective component synthesis
Peripheral SCP: U5 snRNP−selective component synthesis
Numbers: # involved center−peripheral spot pairs [%]
ES units
FME 96.6
IME 1
TME 2.4
0.8
0.6
0.4
0.2
0.0
HK3035
Center SCP: Step 1 and 2 catalysis during pre−mRNA splicing
Peripheral SCP: U5 snRNP−selective component synthesis
Numbers: # involved center−peripheral spot pairs [%]
ES units
FME 91.7
GME 0.7
IME 1.2
TME 6.3
0.75
0.50
0.25
0.00
HK2874
Center SCP: Integrator−mediated nuclear snRNA processing
Peripheral SCP: Cytosolic Sm core assembly onto U snRNAs
Numbers: # involved center−peripheral spot pairs [%]
ES units
TME 100
0.6
0.4
0.2
0.0
HK2877
Center SCP: TREX−mediated coupling of splicing and mRNA export
Peripheral SCP: U1 and U2 snRNP recruitment and pre−spliceosome assembly
Numbers: # involved center−peripheral spot pairs [%]
ES units
FME 97.8
IME 2
TME 0.2
0.8
0.6
0.4
0.2
0.0
HK2529
Center SCP: Nuclear Drosha−mediated microRNA processing
Peripheral SCP: Initiation of miRNA− and siRNA−mediated mRNA silencing
Numbers: # involved center−peripheral spot pairs [%]
ES units
FME 14.2
GME 1
IME 29.4
TME 55.4
0.9
0.6
0.3
0.0
HK2871
Center SCP: Nuclear Drosha−mediated microRNA processing
Peripheral SCP: Cytosolic Dicer−mediated microRNA and endogenous siRNA processing
Numbers: # involved center−peripheral spot pairs [%]
ES units
FME 35.6
GME 2.3
IME 1.6
TME 60.6
0.6
0.4
0.2
HK2873
Center SCP: mRNA storage, translational repression and degradation by cytosolic processing bodies
Peripheral SCP: Cytosolic mRNA decapping and 5' degradation
Numbers: # involved center−peripheral spot pairs [%]
ES units
FME 69
GME 2.5
IME 0.7
TME 27.8
0.6
0.4
0.2

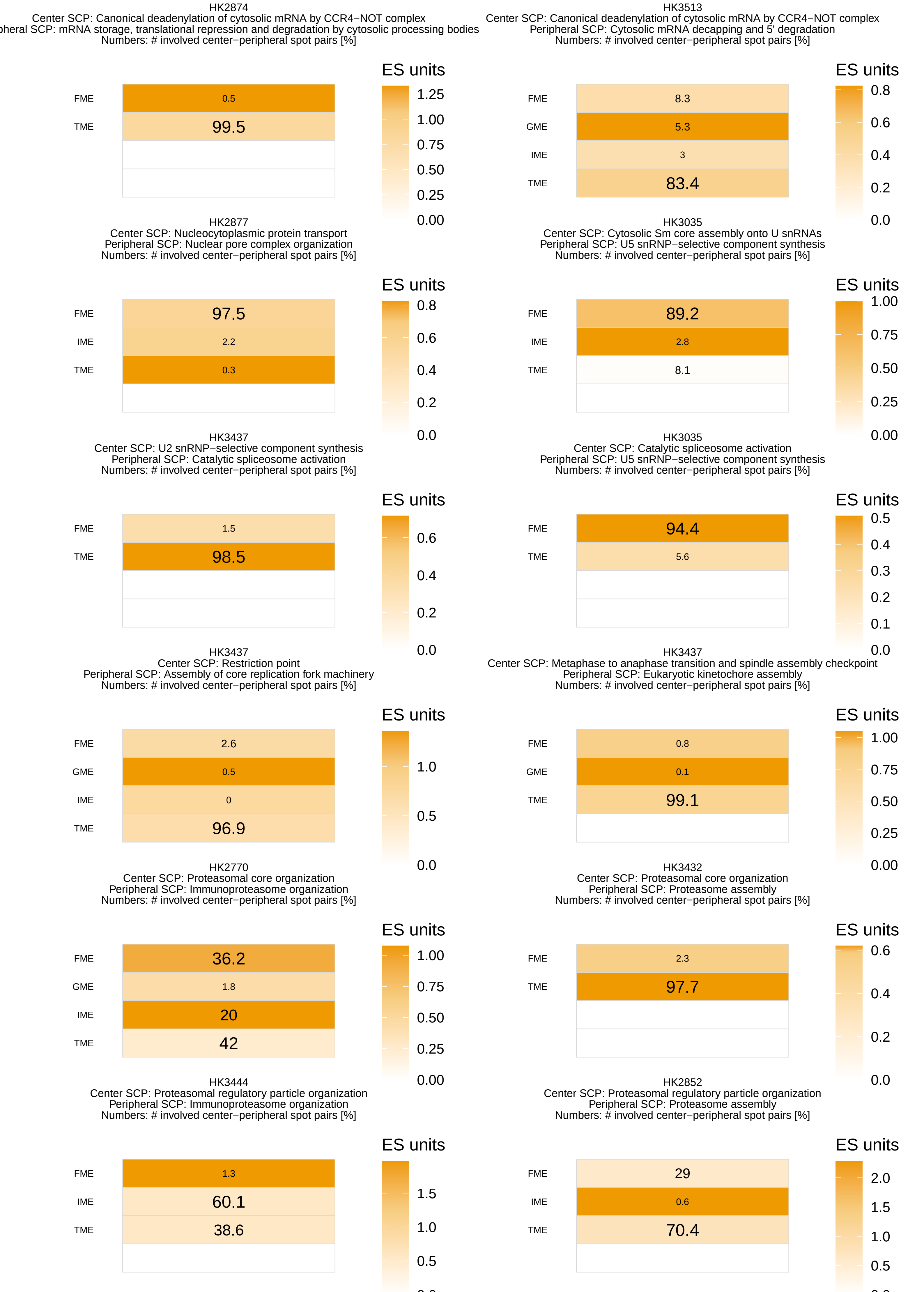
HK2874
Center SCP: Canonical deadenylation of cytosolic mRNA by CCR4−NOT complex
pheral SCP: mRNA storage, translational repression and degradation by cytosolic processing bodies
Numbers: # involved center−peripheral spot pairs [%]
ES units
FME 0.5
TME 99.5
1.25
1.00
0.75
0.50
0.25
0.00
HK3513
Center SCP: Canonical deadenylation of cytosolic mRNA by CCR4−NOT complex
Peripheral SCP: Cytosolic mRNA decapping and 5' degradation
Numbers: # involved center−peripheral spot pairs [%]
ES units
FME 8.3
GME 5.3
IME 3
TME 83.4
0.8
0.6
0.4
0.2
0.0
HK2877
Center SCP: Nucleocytoplasmic protein transport
Peripheral SCP: Nuclear pore complex organization
Numbers: # involved center−peripheral spot pairs [%]
ES units
FME 97.5
IME 2.2
TME 0.3
0.8
0.6
0.4
0.2
0.0
HK3035
Center SCP: Cytosolic Sm core assembly onto U snRNAs
Peripheral SCP: U5 snRNP−selective component synthesis
Numbers: # involved center−peripheral spot pairs [%]
ES units
FME 89.2
IME 2.8
TME 8.1
1.00
0.75
0.50
0.25
0.00
HK3437
Center SCP: U2 snRNP−selective component synthesis
Peripheral SCP: Catalytic spliceosome activation
Numbers: # involved center−peripheral spot pairs [%]
ES units
FME 1.5
TME 98.5
0.6
0.4
0.2
0.0
HK3035
Center SCP: Catalytic spliceosome activation
Peripheral SCP: U5 snRNP−selective component synthesis
Numbers: # involved center−peripheral spot pairs [%]
ES units
FME 94.4
TME 5.6
0.5
0.4
0.3
0.2
0.1
0.0
HK3437
Center SCP: Restriction point
Peripheral SCP: Assembly of core replication fork machinery
Numbers: # involved center−peripheral spot pairs [%]
ES units
FME 2.6
GME 0.5
IME 0
TME 96.9
1.0
0.5
0.0
HK3437
Center SCP: Metaphase to anaphase transition and spindle assembly checkpoint
Peripheral SCP: Eukaryotic kinetochore assembly
Numbers: # involved center−peripheral spot pairs [%]
ES units
FME 0.8
GME 0.1
TME 99.1
1.00
0.75
0.50
0.25
0.00
HK2770
Center SCP: Proteasomal core organization
Peripheral SCP: Immunoproteasome organization
Numbers: # involved center−peripheral spot pairs [%]
ES units
FME 36.2
GME 1.8
IME 20
TME 42
1.00
0.75
0.50
0.25
0.00
HK3432
Center SCP: Proteasomal core organization
Peripheral SCP: Proteasome assembly
Numbers: # involved center−peripheral spot pairs [%]
ES units
FME 2.3
TME 97.7
0.6
0.4
0.2
0.0
HK3444
Center SCP: Proteasomal regulatory particle organization
Peripheral SCP: Immunoproteasome organization
Numbers: # involved center−peripheral spot pairs [%]
ES units
FME 1.3
IME 60.1
TME 38.6
1.5
1.0
0.5
HK2852
Center SCP: Proteasomal regulatory particle organization
Peripheral SCP: Proteasome assembly
Numbers: # involved center−peripheral spot pairs [%]
ES units
FME 29
IME 0.6
TME 70.4
2.0
1.5
1.0
0.5

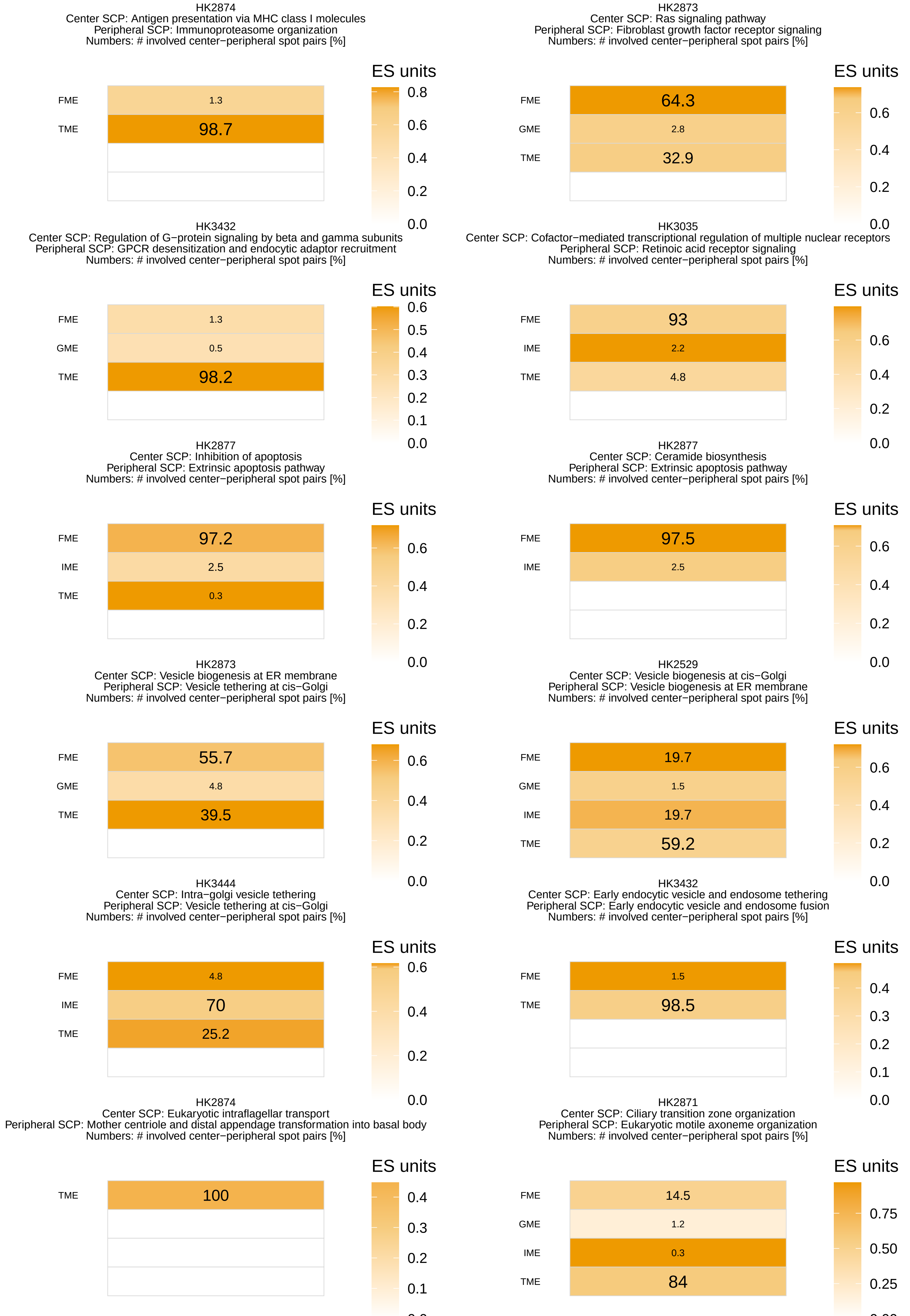
HK2874
Center SCP: Antigen presentation via MHC class I molecules
Peripheral SCP: Immunoproteasome organization
Numbers: # involved center−peripheral spot pairs [%]
FME 1.3
TME 98.7
ES units 0.8 0.6 0.4 0.2 0.0
HK2873
Center SCP: Ras signaling pathway
Peripheral SCP: Fibroblast growth factor receptor signaling
Numbers: # involved center−peripheral spot pairs [%]
FME 64.3
GME 2.8
TME 32.9
ES units 0.6 0.4 0.2 0.0
HK3432
Center SCP: Regulation of G−protein signaling by beta and gamma subunits
Peripheral SCP: GPCR desensitization and endocytic adaptor recruitment
Numbers: # involved center−peripheral spot pairs [%]
FME 1.3
GME 0.5
TME 98.2
ES units 0.6 0.5 0.4 0.3 0.2 0.1 0.0
HK3035
Center SCP: Cofactor−mediated transcriptional regulation of multiple nuclear receptors
Peripheral SCP: Retinoic acid receptor signaling
Numbers: # involved center−peripheral spot pairs [%]
FME 93
IME 2.2
TME 4.8
ES units 0.6 0.4 0.2 0.0
HK2877
Center SCP: Inhibition of apoptosis
Peripheral SCP: Extrinsic apoptosis pathway
Numbers: # involved center−peripheral spot pairs [%]
FME 97.2
IME 2.5
TME 0.3
ES units 0.6 0.4 0.2 0.0
HK2877
Center SCP: Ceramide biosynthesis
Peripheral SCP: Extrinsic apoptosis pathway
Numbers: # involved center−peripheral spot pairs [%]
FME 97.5
IME 2.5
ES units 0.6 0.4 0.2 0.0
HK2873
Center SCP: Vesicle biogenesis at ER membrane
Peripheral SCP: Vesicle tethering at cis−Golgi
Numbers: # involved center−peripheral spot pairs [%]
FME 55.7
GME 4.8
TME 39.5
ES units 0.6 0.4 0.2 0.0
HK2529
Center SCP: Vesicle biogenesis at cis−Golgi
Peripheral SCP: Vesicle biogenesis at ER membrane
Numbers: # involved center−peripheral spot pairs [%]
FME 19.7
GME 1.5
IME 19.7
TME 59.2
ES units 0.6 0.4 0.2 0.0
HK3444
Center SCP: Intra−golgi vesicle tethering
Peripheral SCP: Vesicle tethering at cis−Golgi
Numbers: # involved center−peripheral spot pairs [%]
FME 4.8
IME 70
TME 25.2
ES units 0.6 0.4 0.2 0.0
HK3432
Center SCP: Early endocytic vesicle and endosome tethering
Peripheral SCP: Early endocytic vesicle and endosome fusion
Numbers: # involved center−peripheral spot pairs [%]
FME 1.5
TME 98.5
ES units 0.4 0.3 0.2 0.1 0.0
HK2874
Center SCP: Eukaryotic intraflagellar transport
Peripheral SCP: Mother centriole and distal appendage transformation into basal body
Numbers: # involved center−peripheral spot pairs [%]
TME 100
ES units 0.4 0.3 0.2 0.1
HK2871
Center SCP: Ciliary transition zone organization
Peripheral SCP: Eukaryotic motile axoneme organization
Numbers: # involved center−peripheral spot pairs [%]
FME 14.5
GME 1.2
IME 0.3
TME 84
ES units 0.75 0.50 0.25

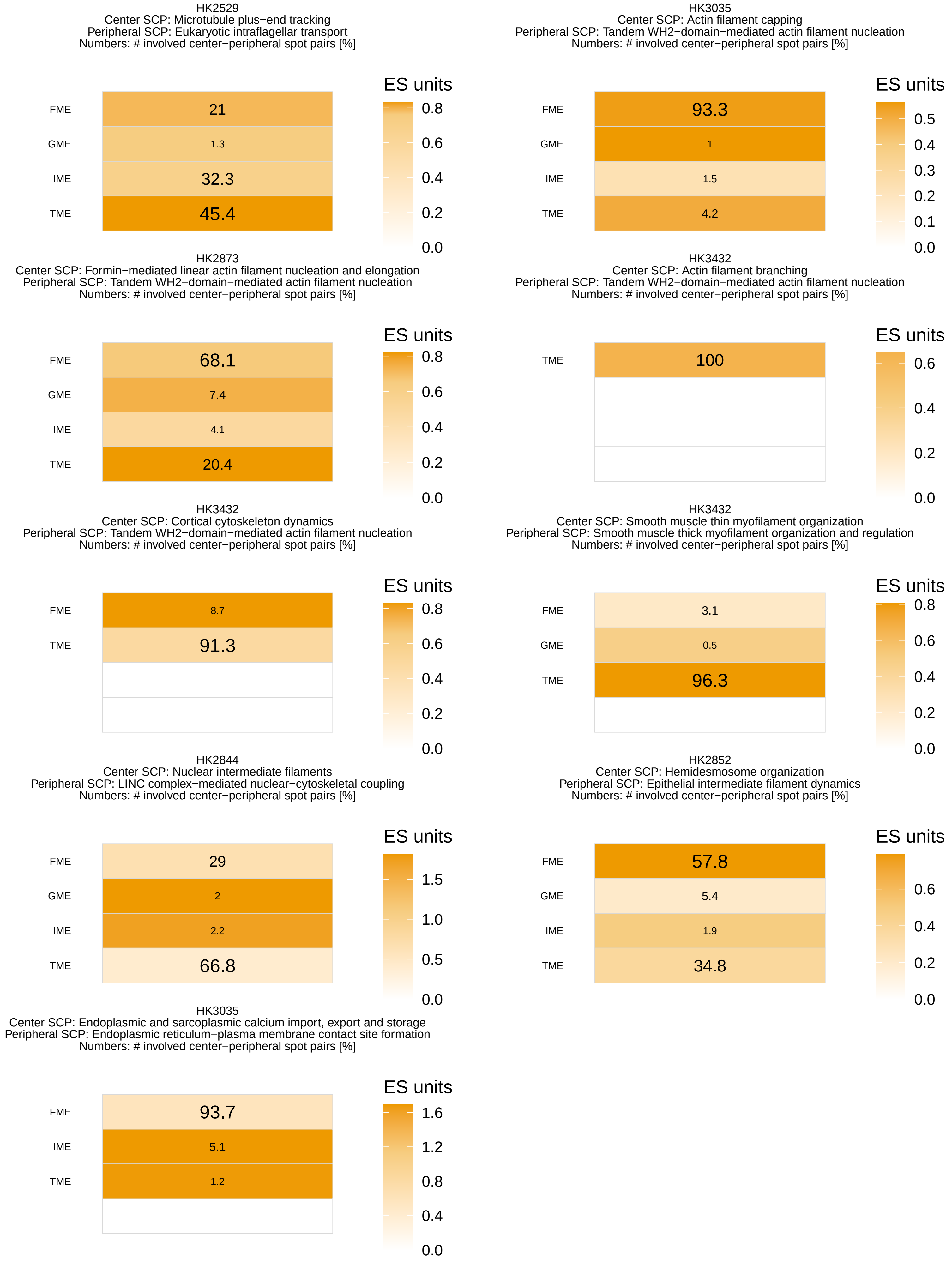
HK2529
Center SCP: Microtubule plus−end tracking
Peripheral SCP: Eukaryotic intraflagellar transport
Numbers: # involved center−peripheral spot pairs [%]
FME 21
GME 1.3
IME 32.3
TME 45.4
ES units
0.8
0.6
0.4
0.2
0.0
HK3035
Center SCP: Actin filament capping
Peripheral SCP: Tandem WH2−domain−mediated actin filament nucleation
Numbers: # involved center−peripheral spot pairs [%]
FME 93.3
GME 1
IME 1.5
TME 4.2
ES units
0.5
0.4
0.3
0.2
0.1
0.0
HK2873
Center SCP: Formin−mediated linear actin filament nucleation and elongation
Peripheral SCP: Tandem WH2−domain−mediated actin filament nucleation
Numbers: # involved center−peripheral spot pairs [%]
FME 68.1
GME 7.4
IME 4.1
TME 20.4
ES units
0.8
0.6
0.4
0.2
0.0
HK3432
Center SCP: Actin filament branching
Peripheral SCP: Tandem WH2−domain−mediated actin filament nucleation
Numbers: # involved center−peripheral spot pairs [%]
TME 100
ES units
0.6
0.4
0.2
0.0
HK3432
Center SCP: Cortical cytoskeleton dynamics
Peripheral SCP: Tandem WH2−domain−mediated actin filament nucleation
Numbers: # involved center−peripheral spot pairs [%]
FME 8.7
TME 91.3
ES units
0.8
0.6
0.4
0.2
0.0
HK3432
Center SCP: Smooth muscle thin myofilament organization
Peripheral SCP: Smooth muscle thick myofilament organization and regulation
Numbers: # involved center−peripheral spot pairs [%]
FME 3.1
GME 0.5
TME 96.3
ES units
0.8
0.6
0.4
0.2
0.0
HK2844
Center SCP: Nuclear intermediate filaments
Peripheral SCP: LINC complex−mediated nuclear−cytoskeletal coupling
Numbers: # involved center−peripheral spot pairs [%]
FME 29
GME 2
IME 2.2
TME 66.8
ES units
1.5
1.0
0.5
0.0
HK2852
Center SCP: Hemidesmosome organization
Peripheral SCP: Epithelial intermediate filament dynamics
Numbers: # involved center−peripheral spot pairs [%]
FME 57.8
GME 5.4
IME 1.9
TME 34.8
ES units
0.6
0.4
0.2
0.0
HK3035
Center SCP: Endoplasmic and sarcoplasmic calcium import, export and storage
Peripheral SCP: Endoplasmic reticulum−plasma membrane contact site formation
Numbers: # involved center−peripheral spot pairs [%]
FME 93.7
IME 5.1
TME 1.2
ES units
1.6
1.2
0.8
0.4
0.0

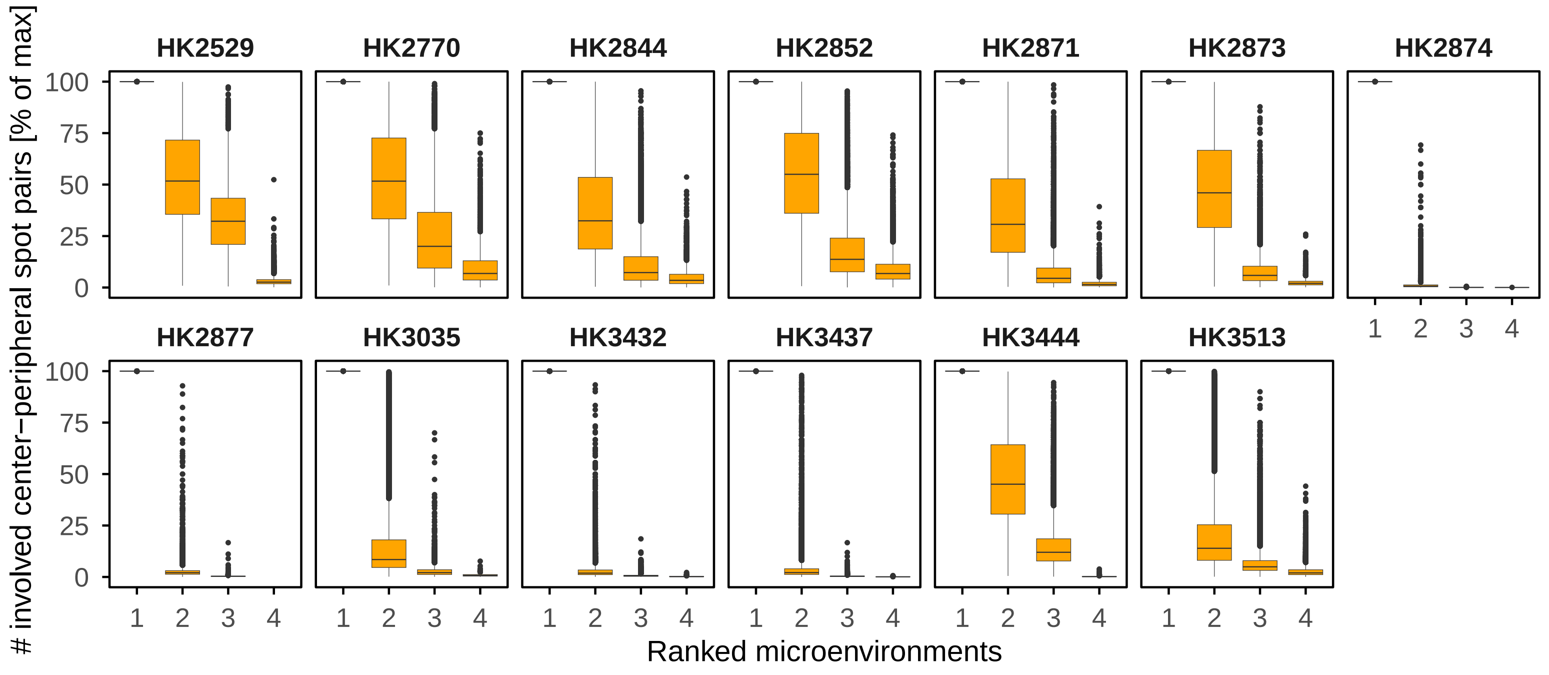

HK2529
HK2770
HK2844
HK2852
HK2871
HK2873
HK2874
HK2877
HK3035
HK3432
HK3437
HK3444
HK3513
### involved center−peripheral spot pairs [% of max]
0
25
50
75
100
1
2
3
4
Ranked microenvironments


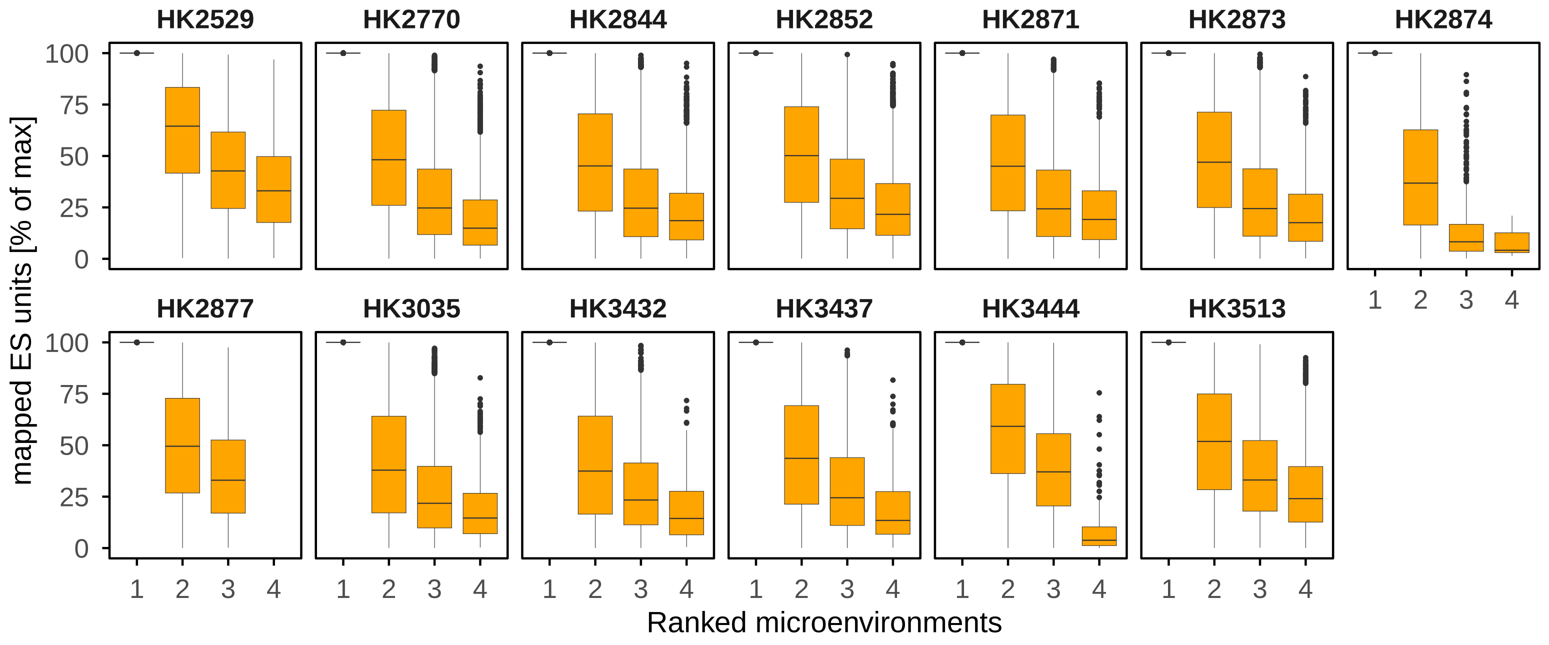

HK2529
HK2770
HK2844
HK2852
HK2871
HK2873
HK2874
HK2877
HK3035
HK3432
HK3437
HK3444
HK3513
mapped ES units [% of max]
0
25
50
75
100
1
2
3
4
Ranked microenvironments


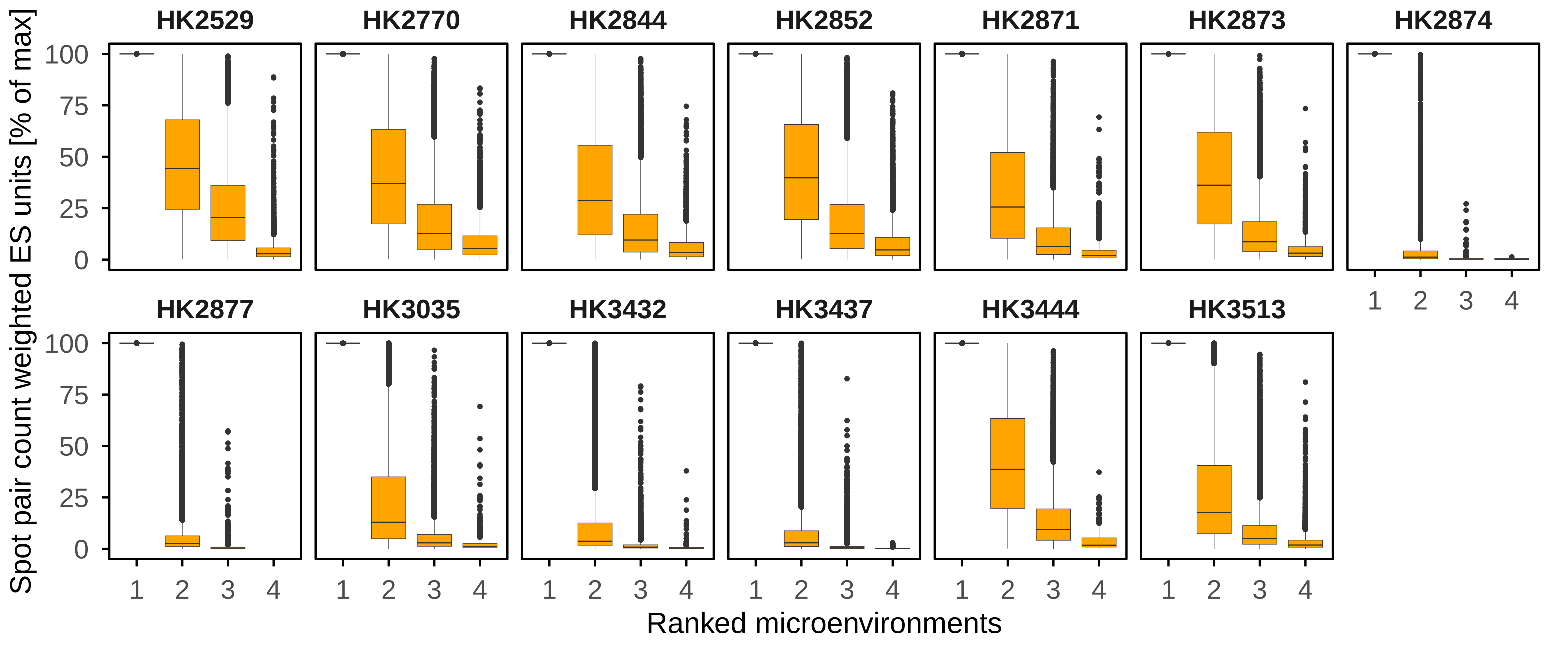

HK2529
HK2770
HK2844
HK2852
HK2871
HK2873
HK2874
HK2877
HK3035
HK3432
HK3437
HK3444
HK3513
Spot pair count weighted ES units [% of max]
0
25
50
75
100
1
2
3
4
Ranked microenvironments

**Supplementary Figure 8. Documentation of microenvironment-selectivity for identified SCP pairs. (A)** We repeated our analysis pipeline, but this time averaged the results only across center and peripheral spots that belonged to the same microenvironment. Mapped (not normalized) ES units within each microenvironment are shown for selected SCP pairs. **(B)** To systematically analyze potential microenvironment selectivity, we identified that microenvironment for each SCP pair that contains the highest number of center spots with the cross-cellular SCP pair mapping in question. We then normalized the number of center spots for all cell type pairs towards the identified highest number of center spots, followed by ranking of all microenvironments --still for the same SCP pair- by the normalized center spot counts. For each tissue slice, we then combined the microenvironments associated with all SCP pairs into groups of same ranks (ignoring which microenvironments share ranks) to show the normalized center spot counts associated with each of the four microenvironments (top panel). We repeated the same analysis with the mapped ES units (middle panel) and the product of the number of center spots and the mapped ES units (bottom panel and main figure 9D).
